# Community Resource: A Genome-Based Extension of Large-Scale Wheat Proteogenomics

**DOI:** 10.64898/2026.06.17.733048

**Authors:** Delphine Vincent, Rudi Appels

**Affiliations:** Faculty of Science, University of Melbourne, Parkville, VIC 3010, Australia

**Keywords:** *Triticum aestivum*, proteogenomics, genome annotation, exon-resolved peptide projection, FragPipe, MSFragger, GFF3, BED, Apollo JBrowse

## Abstract

Bread wheat (*Triticum aestivum* L.) possesses a large and highly repetitive allohexaploid genome and annotation requires extensive protein-level validation. We developed a genome-based wheat proteogenomics workflow integrating large-scale MS/MS reanalysis, GFF3-based peptide coordinate reconstruction, thorough validation, and genome browser-compatible peptide deployment against the IWGSC RefSeq v2.1 reference genome. Public wheat proteomics datasets comprising 577 raw mass spectrometry files (∼1.0 TB) from 32 tissues were reprocessed using FragPipe/MSFragger, generating 2,226,779 non-redundant peptides and 1,648,740 unique protein accessions. Peptide-to-genome projections using GFF3 annotation files produced 8,291,056 genomic peptide projected rows, of which 98.14% passed validation procedures. Overall, peptide evidence supported 103,095 high-confidence (HC) and 135,495 low-confidence (LC) wheat gene models, corresponding to 96.4% and 84.7% of all parsed HC and LC annotations, respectively. In total, 238,590 wheat gene models (89.4% of all parsed annotations) received protein-level support. Apollo/JBrowse-compatible BED tracks enabled exon-resolved visualisation of peptide evidence across wheat chromosomes. Together, this study establishes a scalable GFF3-based proteogenomics framework for complex polyploid plant genomes and provides an extensive community resource for wheat genome annotation refinement and visual exploration (https://bread-wheat-um.genome.edu.au/apollo/49826/jbrowse/index.html).

**Graphical abstract:** 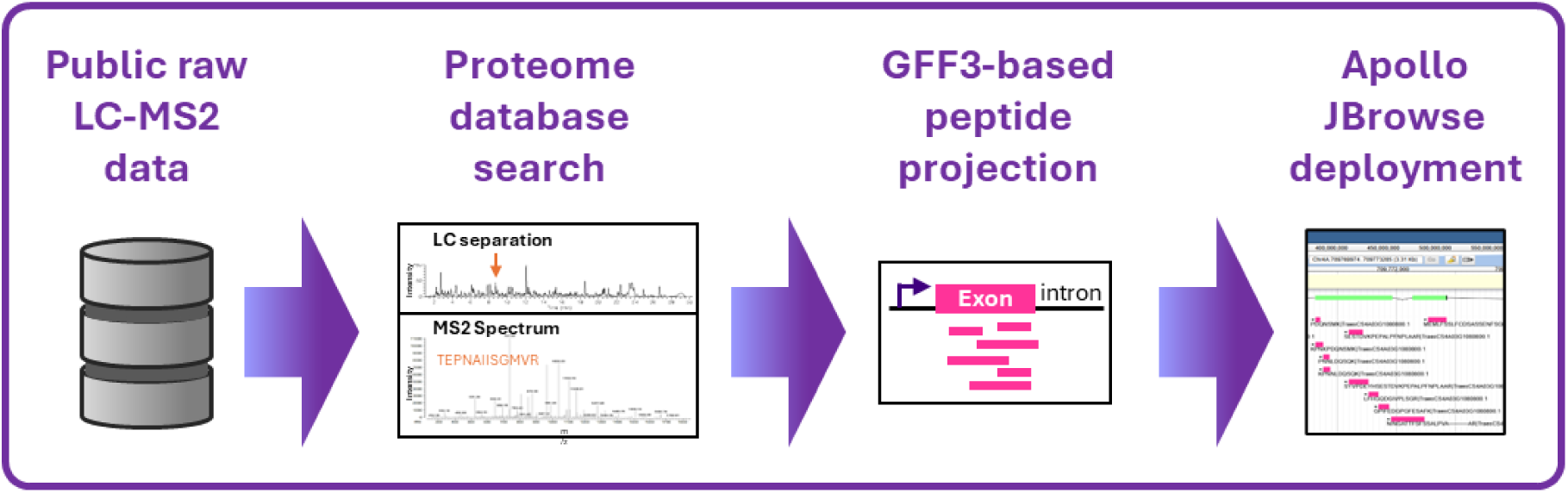

## 1. INTRODUCTION

Bread wheat (*Triticum aestivum* L.) is one of the world’s most important staple crops and a major source of human calories and protein^1^. The release of chromosome-scale wheat reference assemblies by the International Wheat Genome Sequencing Consortium (IWGSC)^2–5^ represented a major breakthrough for wheat genomics by providing high-quality genome sequences together with extensive high-confidence (HC) and low-confidence (LC) gene annotations. These genomic resources have substantially accelerated wheat functional genomics, breeding, and systems biology research by enabling detailed investigation of gene structure, expression, and genome organisation^6–9^.

Despite these advances, accurate annotation of large and highly repetitive polyploid genomes remains challenging^10^. Bread wheat possesses a complex allohexaploid genome rich in duplicated homeologous loci, repetitive sequences, alternative splice isoforms, and fragmented transcript evidence, all of which complicate reliable gene-model prediction and validation^2^. Consequently, independent experimental evidence, including peptide-level mass-spectrometry support, is required to confirm protein-coding potential and refine existing genome annotations^11^.

At the interface of proteomics and genomics, proteogenomics provides an important complementary strategy by integrating mass spectrometry-derived peptide evidence with genomic and transcriptomic resources to annotate protein-coding genes^12–14^. This methodology provides orthogonal information to traditional forms of evidence used for genome annotation. In plants, proteogenomics has been used to validate predicted coding regions, refine gene models, and support weakly annotated loci, while peptide evidence directly confirms protein-level expression^15–19^. In wheat, large-scale proteomics resources across multiple tissues and developmental stages have created valuable opportunities for genome-scale annotation refinement^20–24^.

In our previous study, we developed a workflow based on direct tBLASTn alignment of public wheat peptide sequences against the IWGSC RefSeq v2.1 genome, demonstrating large-scale peptide mapping and annotation support across wheat chromosomes^25^. However, direct peptide genomic alignment remained computationally restrictive and sensitive to peptide length, splice junctions, and frame-disrupted regions.

In the present study, we extend our previous work by developing a genome-based proteogenomics workflow integrating raw data reprocessing, peptide genomic coordinate reconstruction based on Generic Feature Format Version 3 (GFF3), four-pronged projection validations, and genome browser-compatible deployment via Browser Extensible Data (BED) exports. This peptide-to-genome framework relying on genomic annotations improves peptide recovery and proteogenomic coverage while providing a scalable resource for wheat gene model refinement and Apollo/JBrowse visualisation.

## 2. MATERIALS AND METHODS

### 2.1. Study overview

The study proteogenomic strategy is outlined in Figure 1.

**Figure 1.**
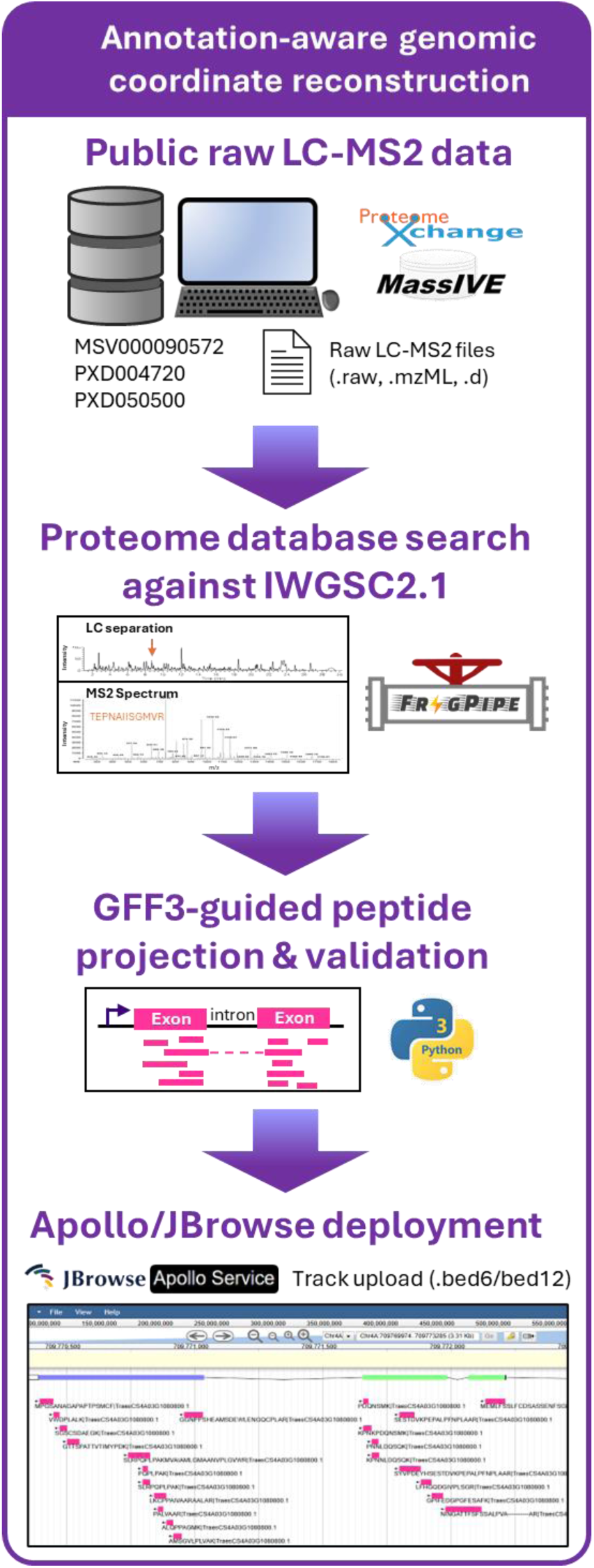
Overview of the wheat proteogenomic workflow. The genome-based workflow developed in this study involved raw MS/MS reanalysis against the IWGSC RefSeq v2.1 proteome, followed by GFF3-based peptide projection and generation of browser-ready BED6/BED12 tracks for Apollo JBrowse visualization; details are available in Suppl. Fig. S1 and Jupyter Notebook.

Following peptide search on raw tandem mass spectrometry data (MS/MS) using FragPipe, peptide genomic coordinates were programmatically reconstructed in Python using genome annotation-based peptide projection, using custom Python scripts^26^ implemented within JupyterLab Notebook environment^27^. Peptide locations within proteins were mapped back to chromosomal coordinates using the transcript, exon, and coding sequence information contained within the IWGSC RefSeq v2.1 GFF3^28^ files. Peptide-to-genome projections were thoroughly checked and only validated projections were exported as BED files for JBrowse^29^ deployment. The proteogenomic workflow is summarised in Suppl. Fig. S1 and all the steps involved are fully detailed in the notebook (Suppl. File S1). FragPipe^30^ processing parameters are supplied in Suppl. File S2. Peptide genomic coordinate reconstruction and validations are explained in Suppl. File S3. Python outputs are presented in Fig. 2, Suppl. Fig. S2-4, Table 1 and Suppl. Tables S1-S6. Peptide alignments along the wheat genome are exemplified in Fig. 3. BED files are available via GitHub (https://github.com/dlf2024/Python-wheat-proteogenomics_2026) and publicly accessible on Apollo/JBrowse server (https://bread-wheat-um.genome.edu.au/apollo/49826/jbrowse/index.html).

**Figure 2.**
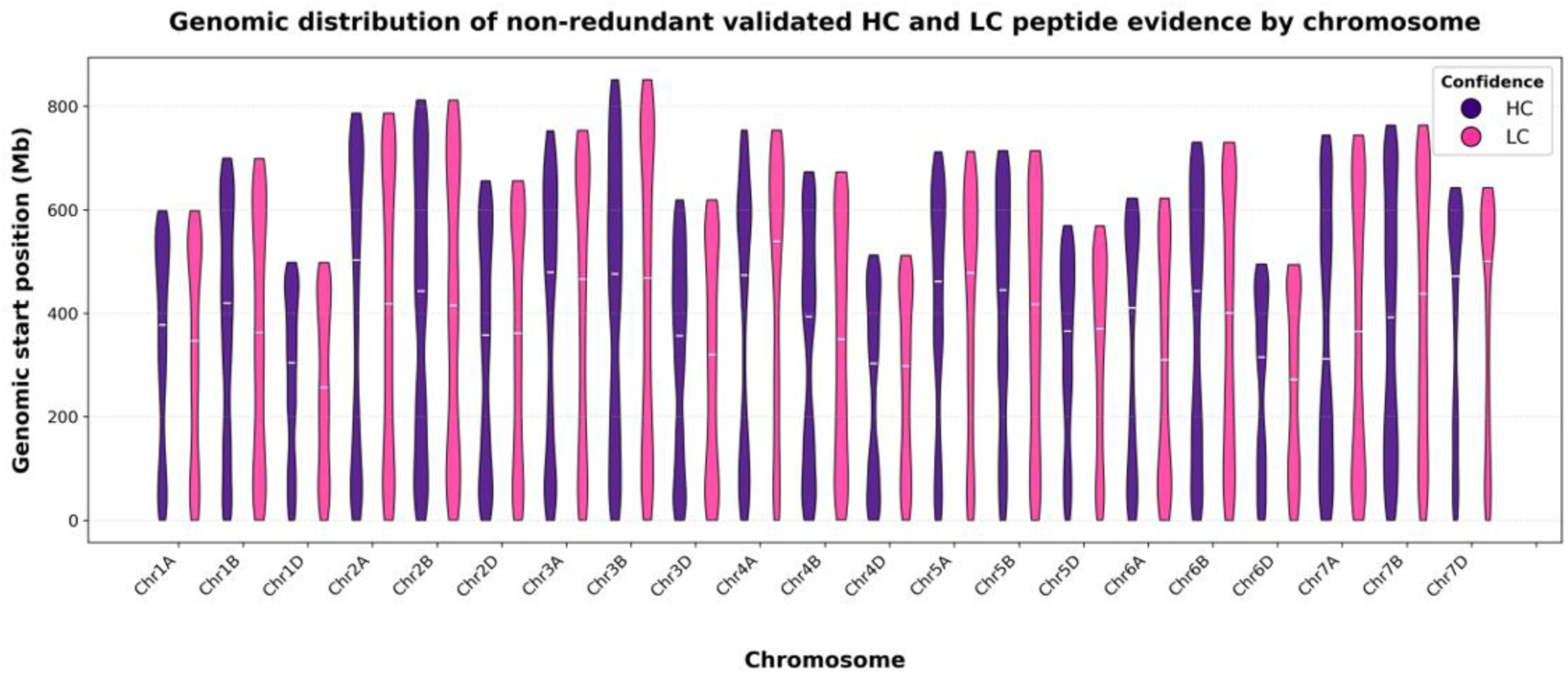
Genome-wide distribution of projected peptide evidence across wheat chromosomes. Violin plot shows genomic coordinate distributions of HC and LC peptide projections for each chromosome. The white horizontal bar denotes the median.

**Figure 3.**
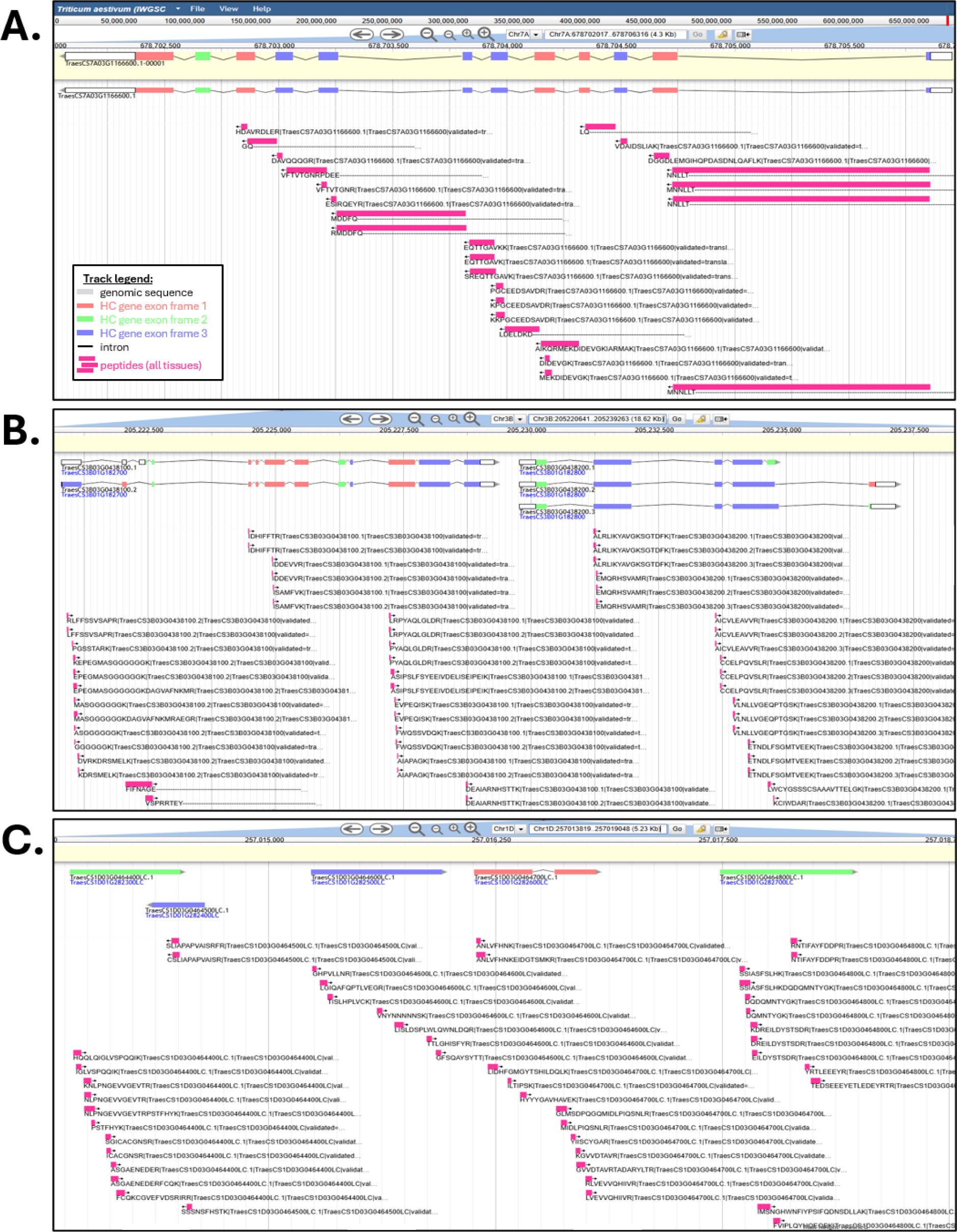
Apollo/JBrowse visualisation of GFF3-based peptide projections along wheat gene models. (A) Example of HC gene model validation supported by extensive projected peptide evidence including intron-spanning peptides (Chr7A:678700836..678707865). (B) Examples of projected peptides aligning with isoforms (Chr3B:205220641..205239263). (C) Examples of mapped peptides supporting LC gene models (Chr1D:257013819..257019048), suggesting potential promotion to HC status.

**Table 1.**
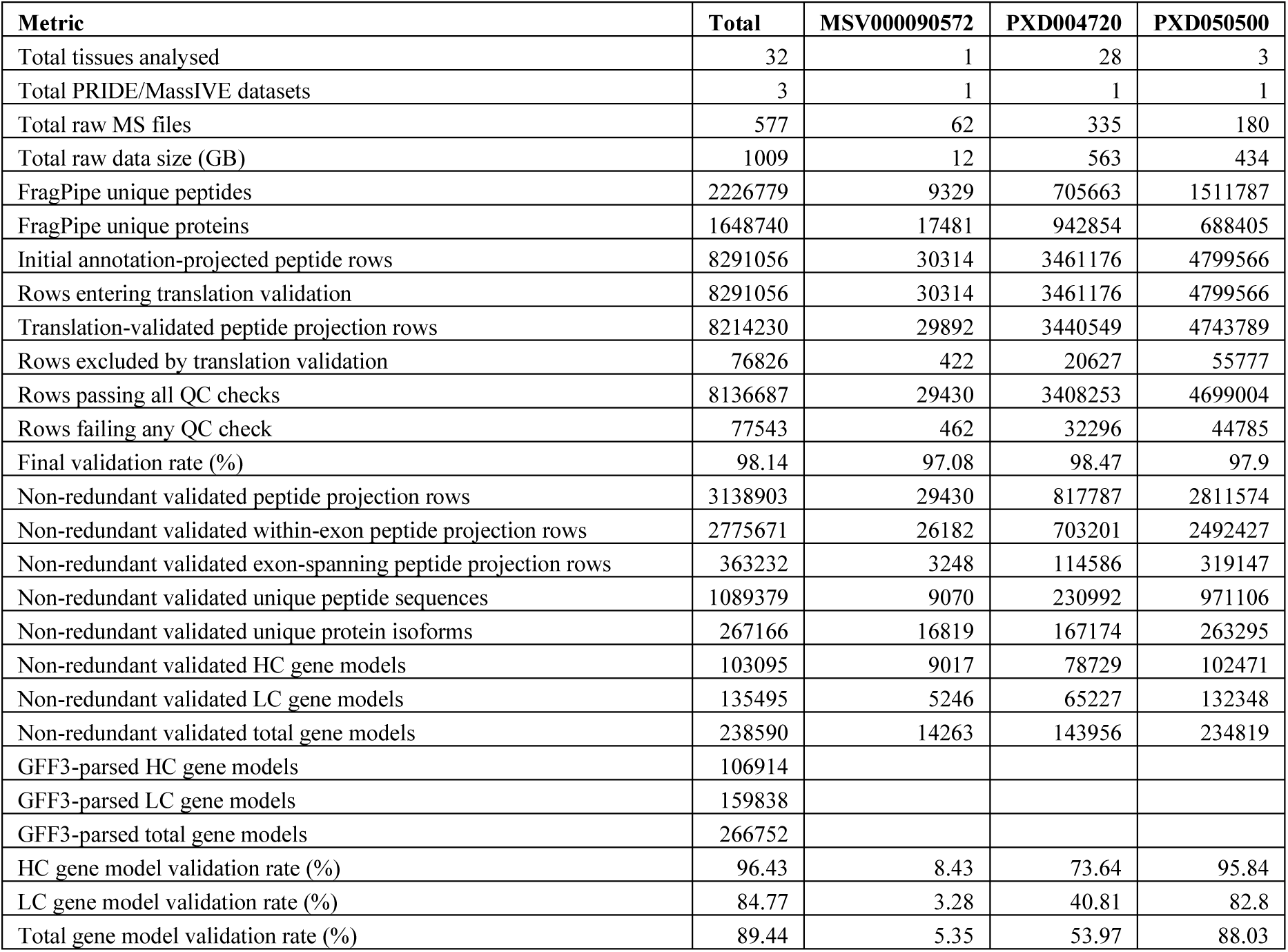
Summary of proteogenomic workflow outputs across public wheat datasets.

| Metric | Total | MSV000090572 | PXD004720 | PXD050500 |
| --- | --- | --- | --- | --- |
| Total tissues analysed | 32 | 1 | 28 | 3 |
| Total PRIDE/MassIVE datasets | 3 | 1 | 1 | 1 |
| Total raw MS files | 577 | 62 | 335 | 180 |
| Total raw data size (GB) | 1009 | 12 | 563 | 434 |
| FragPipe unique peptides | 2226779 | 9329 | 705663 | 1511787 |
| FragPipe unique proteins | 1648740 | 17481 | 942854 | 688405 |
| Initial annotation-projected peptide rows | 8291056 | 30314 | 3461176 | 4799566 |
| Rows entering translation validation | 8291056 | 30314 | 3461176 | 4799566 |
| Translation-validated peptide projection rows | 8214230 | 29892 | 3440549 | 4743789 |
| Rows excluded by translation validation | 76826 | 422 | 20627 | 55777 |
| Rows passing all QC checks | 8136687 | 29430 | 3408253 | 4699004 |
| Rows failing any QC check | 77543 | 462 | 32296 | 44785 |
| Final validation rate (%) | 98.14 | 97.08 | 98.47 | 97.9 |
| Non-redundant validated peptide projection rows | 3138903 | 29430 | 817787 | 2811574 |
| Non-redundant validated within-exon peptide projection rows | 2775671 | 26182 | 703201 | 2492427 |
| Non-redundant validated exon-spanning peptide projection rows | 363232 | 3248 | 114586 | 319147 |
| Non-redundant validated unique peptide sequences | 1089379 | 9070 | 230992 | 971106 |
| Non-redundant validated unique protein isoforms | 267166 | 16819 | 167174 | 263295 |
| Non-redundant validated HC gene models | 103095 | 9017 | 78729 | 102471 |
| Non-redundant validated LC gene models | 135495 | 5246 | 65227 | 132348 |
| Non-redundant validated total gene models | 238590 | 14263 | 143956 | 234819 |
| GFF3-parsed HC gene models | 106914 |  |  |  |
| GFF3-parsed LC gene models | 159838 |  |  |  |
| GFF3-parsed total gene models | 266752 |  |  |  |
| HC gene model validation rate (%) | 96.43 | 8.43 | 73.64 | 95.84 |
| LC gene model validation rate (%) | 84.77 | 3.28 | 40.81 | 82.8 |
| Total gene model validation rate (%) | 89.44 | 5.35 | 53.97 | 88.03 |

### 2.2. Genome annotation resources

Proteogenomic analyses were performed against *Triticum aestivum* reference genome IWGSC RefSeq v2.1 and associated gene model annotations, including GFF3 genome annotation files, gene model functional annotations as well as HC and LC protein sequences as FASTA format^2–5^ (https://urgi.versailles.inrae.fr/download/iwgsc/IWGSC_RefSeq_Annotations/v2.1/). The IWGSC RefSeq v2.1 assembly and its associated high-confidence (HC) and low-confidence (LC) annotations provide a refined chromosome-scale representation of the allohexaploid wheat genome, comprising 266,752 annotated genes^5^, and were therefore selected as the reference framework for peptide projection in this study. Using Python, the 106,914 HC and 159,838 LC GFF3 annotation files were parsed to generate protein-, transcript-, exon-, Coding DNA Sequence (CDS)-, and gene-level coordinate tables required for peptide-to-genome projection analyses. Protein-to-gene relationships and transcript exon structures were reconstructed directly from the annotation files (Suppl. File S1).

### 2.3. Public proteomics datasets

Large-scale wheat proteomics studies and proteome atlas efforts were used as a framework for comparison and interpretation of tissue-level protein evidence across the wheat genome^20,22,31^. From these studies, publicly available wheat proteomics datasets covering multiple tissues and developmental stages were retrieved from the PRIDE^32^ and MassIVE^33^ repositories (PXD004720, PXD050500, MSV000090572) to maximise proteogenomic coverage across the wheat genome. Raw mass spectrometry data were downloaded in vendor-specific formats and converted to mzML format where required using ProteoWizard MSConvert^34^. Across all datasets, peptide identification workflows were standardised to ensure consistency of downstream proteogenomic analyses. Tissue-specific outputs were retained separately to enable comparative assessment of peptide support across wheat tissues and developmental stages (Suppl. File S1).

### 2.4. Protein database

A protein search database was generated from the IWGSC RefSeq v2.1 HC and LC peptide FASTA files using the Galaxy platform^35^. HC and LC protein sequences were concatenated together with common laboratory contaminant proteins (cRAP database) and decoy reversed sequences for false discovery rate (FDR) estimation^36^. Technical details are provided in Suppl. File S1. This protein database enabled simultaneous assessment of peptide support for both HC and LC annotations; it can be retrieved via our GitHub repository (https://github.com/dlf2024/Python-wheat-proteogenomics_2026/tree/main/protein_database).

### 2.5. Peptide identification

Searches were performed against the combined wheat HC–LC target-decoy protein database using FragPipe 24.0 with MSFragger 4.4.1, PeptideProphet, ProteinProphet, and Philosopher reporting^30^. In this context, peptide identification refers to matching experimentally observed MS/MS fragmentation spectra against theoretically predicted peptide sequences derived from the wheat protein database. The search used Data-dependent acquisition (DDA) mode, Trypsin/P specificity, fully enzymatic cleavage, and up to 10 missed cleavages. Precursor and true precursor mass tolerances were set to ±10 ppm and ±20 ppm, respectively, with a fragment mass tolerance of 0.05 Da. Static cysteine carbamidomethylation (+57.02146 Da) was applied, while methionine oxidation (+15.9949 Da), protein N-terminal acetylation (+42.0106 Da), and protein N-terminal methionine clipping were treated as variable modifications. Peptides were restricted to 6–100 amino acids and 300–10,000 Da. Peptide-spectrum match (PSM) validation was enabled, and final reports were generated using sequential filtering at 1% protein FDR, with decoys excluded and contaminants removed from the final reports. The complete workflow parameters are available in Suppl. File S2. FragPipe was selected for its rapid and sensitive large-scale peptide search capacity^30^ suitable for complex polyploid proteomes. Processing parameters were harmonised across all datasets to minimise technical variation between tissues and studies.

### 2.6. Peptide-to-genome projection

Proteogenomic mapping was performed using a custom workflow implemented in Python^26^ and employed genome annotations (Suppl. Fig. S1, Suppl. File S3). These GFF3 files describe the genomic organisation of genes, transcripts, exons, and coding regions along chromosomes^28^ and therefore provide the structural framework required for peptide-to-genome projection. Identified peptides were first linked to their corresponding wheat protein identifiers. Protein identifiers were then associated with transcript and CDS coordinates extracted from the HC and LC GFF3 files. For each peptide, amino acid (AA) coordinates within the parent protein sequence were converted to genomic coordinates by mapping the corresponding coding nucleotide interval onto the annotated CDS structure of the parent transcript, thereby accommodating both single-exon and multi-exon gene models.

### 2.7. Projection validation

Projected peptides were subsequently subjected to four independent validation procedures using Python scripts (Suppl. Fig. S1, Suppl. File S3). First, genomic intervals were translated back into AA sequences following exon reconstruction and compared with the original experimentally identified peptide sequence. Second, BED geometry was examined to ensure valid block structures and coordinate ranges. Third, chromosome and strand assignments were verified for consistency with the parent transcript model. Fourth, protein-coordinate relationships were checked to confirm that projected genomic intervals corresponded to the correct peptide location within the source protein. Only projections passing all validation checks were retained for export and downstream analyses. This procedure enabled peptide projection across exon junctions and accounted for transcript strand orientation.

### 2.8. Projection exportation

Using Python, projected peptides were exported as BED6 and BED12 tracks compatible with JBrowse genome visualisation tool^29^, made available via GitHub (https://github.com/dlf2024/Python-wheat-proteogenomics_2026/tree/main/BED_files) and permanently uploaded on the Apollo server^37^ (https://bread-wheat-um.genome.edu.au/apollo/49826/jbrowse/index.html). Peptide labels embedded within BED tracks incorporated peptide sequence information together with dashes as visual indicators of exon-spanning structure (Suppl. File S1).

### 2.9. Exploratory data analysis

Exploratory data analysis (EDA) was performed using Python and Matplotlib package^38^. Tissue-specific and combined peptide projection outputs were summarised using violin plot, scatterplot, boxplot, histogram, cumulated barplot, UpSet plot and pie charts (Suppl. File S1). Those charts helped quantify HC and LC gene model support, tissue-specific proteogenomic coverage, proportions of intron-spanning peptides and chromosome-level peptide distributions.

### 2.10. Reproducible computational workflow available on GitHub

All data processing, peptide projection, filtering, summarisation, and visualisation workflows were implemented using Python within Jupyter Notebook environments. The notebook structure was designed to provide a fully reproducible end-to-end proteogenomic workflow spanning raw data retrieval and processing, peptide-to-genome mapping with projection validation and export, as well as EDA for data interpretation (Suppl. File S1). The use of notebook-based computational workflows aligns with current recommendations promoting reproducibility and transparency in large-scale omics analyses^27^. Such a platform is widely used in programming as it offers a robust framework for documenting and sharing all activities of research projects^39^. Workflow modularity also enabled iterative optimisation of peptide projection strategies and quality-control procedures during development of the proteogenomic framework. The full notebook and most associated files are publicly available via GitHub (https://github.com/dlf2024/Python-wheat-proteogenomics_2026).

## 3. RESULTS AND DISCUSSION

### 3.1. Genome-based wheat proteogenomic workflow

A proteogenomic workflow based on by *T aestivum* genome annotations was developed to assign experimentally identified wheat peptides to precise chromosomal positions while preserving exon–intron gene structure (Fig. 1; Suppl. Fig. S1). Unlike our previous pipeline, which relied on tBLASTn alignments of public peptide identifications against the bread wheat genome^25^, the present strategy reprocessed the original raw MS/MS files using a standardised FragPipe workflow and searched them against the IWGSC RefSeq v2.1 HC+LC protein database (Suppl. File S2). This generated a consistent peptide-identification framework across all datasets and the various wheat tissues and developmental stages they covered^20,22,23^.

Identified peptides were then linked to their parent wheat proteins, transcripts, and gene models using the IWGSC RefSeq v2.1 GFF3 annotations. Because these annotations define exon, CDS, transcript, and gene coordinates^28^, peptide amino-acid positions within proteins could be converted into coding nucleotide intervals and projected onto chromosome coordinates through exon-resolved CDS reconstruction followed by thorough validation checks (Suppl. File S3). This enabled both single-exon and exon-spanning peptides to be represented correctly, with multi-exon peptides exported as discontinuous genomic blocks reflecting the underlying exon–intron structure. Final validated projections were exported as browser-compatible BED6/BED12 tracks for Apollo/JBrowse visualisation^29,40^.

Annotation-based proteogenomics provides an orthogonal layer of evidence for validating and refining gene annotations, particularly in complex plant genomes where duplicated loci, large gene families, repetitive content, and incomplete transcript evidence complicate annotation^2,4,5,7^. In wheat, the IWGSC RefSeq v2.1 assembly and HC/LC annotations provide a chromosome-scale reference framework^2,5^, but protein-level evidence remains important for confirming coding potential and supporting annotation review^12,13^. In plants, proteogenomics has been successfully applied to improve genome annotation and identify novel coding features in species including *Arabidopsis thaliana*^41^, grapevine^17^, sweet potato^15^, pear^19^, pineapple^16^, and wheat ^20,25^. These studies primarily employed proteogenomic database-expansion strategies in which peptide identifications were interpreted against translated genomic, transcriptomic, or customised protein sequence resources to discover novel coding loci, splice variants, exon-boundary corrections, and annotation discrepancies. In contrast, relatively few proteogenomic workflows have focused on direct annotation-based peptide coordinate reconstruction using existing genome annotations. Methods such as PGx^42^, PoGo^43^, and Peptimapper^44^ were specifically developed to project experimentally identified peptides onto genomic coordinates using transcript and coding-sequence annotations, enabling visualisation within genome browsers and facilitating annotation validation. The present workflow extends this annotation-based paradigm to bread wheat by leveraging IWGSC RefSeq v2.1 GFF3 annotations to generate exon-resolved peptide projections and browser-compatible Apollo/JBrowse tracks at genome scale.

### 3.2. Large-scale peptide-to-genome projection and validation

Across the three public wheat proteomics datasets analysed^20,22,23^, capturing 32 tissues and developmental stages, a total of 577 raw MS files (∼1.0 TB) were reprocessed and generated 2,226,779 unique peptides and 1,648,740 unique proteins following contaminant filtering and protein inference (Table 1). GFF3-based peptide-to-genome mapping subsequently produced 8,291,056 projection rows (*i.e.* unique protein/peptide pairs), reflecting a vast peptide redundancy across tissues, transcript isoforms, and homeologous gene copies within the allohexaploid wheat genome.

No orphan peptides were detected among the annotation-supported peptide evidence entering genome projection. A small number of peptide-protein pairs (932/8,291,988, ∼0.01%, Suppl. Table S1) lacked a GFF3-derived gene-model mapping and were therefore excluded; these should be distinguished from failed genome projections. Genome projection was complete for all annotation-supported rows with 100% peptide-protein-gene rows mapped to genomic coordinates prior to validation. Failed projections therefore represent conservative exclusions of initially assigned peptide/protein pairs that did not meet sequence or coordinate-provenance criteria, rather than evidence for novel loci or missing annotations.

To ensure that GFF3-based peptide projections were biologically valid and genome-browser compatible, each projected row was subjected to one translation-based check and three coordinate-level quality control checks. Translation validation reconstructed the projected genomic CDS sequence, translated it in the expected reading frame, and compared it with the original FragPipe-identified peptide, consistent with proteogenomic approaches that use MS/MS evidence to validate translated coding regions and refine gene annotation^11,12^. Coordinate-level checks assessed BED geometry, exon-resolved block lengths, and concordance of chromosome, strand, and protein-coordinate assignments with the source GFF3 annotation, in line with established peptide-to-genome mapping principles for genome-browser-compatible.visualisation^42,43^.

Translation validation excluded 76,826 projection rows (0.93%), including 76,811 translation mismatches and 15 rows containing ambiguous nucleotide characters (Table 1; Suppl. Table S2). After isoleucine/leucine normalisation, 8,214,230 of 8,291,056 projected rows (99.07%) were translation-validated, with tissue-level rates ranging from 98.61% to 99.65% (Suppl. Table S2). Validation covered all projected peptide rows, including single-block, multi-block, positive-strand, and negative-strand projections. The high validation rate therefore supports robust reconstruction across both simple and structurally complex projections, including exon-spanning and reverse-strand gene models. This is consistent with proteogenomic studies showing that peptide evidence can validate exons, splice junctions, and coding structures at the level of translation^13,41,45^.

Following translation validation, 8,136,687 rows passed all coordinate-level quality control checks, whereas 77,543 rows failed at least one check and were excluded from the final genome-browser exports (Table 1; Suppl. Table S1). No projections against ChrUnknown were retained. Quality control check failures were attributable to chromosome/strand concordance inconsistencies, while BED geometry, nucleotide block-length, and protein-coordinate checks were satisfied for all translation-validated rows (Suppl. Table S3). Thus, excluded rows primarily reflected unresolved chromosome/strand provenance rather than failure of exon-resolved coordinate reconstruction or BED feature construction^42^.

Overall, 8,136,687 of the 8,291,056 initial annotation-projected rows passed both translation validation and all coordinate-level quality control checks, corresponding to a final validation rate of 98.14% (Table 1). Dataset-level validation rates were similarly high: 97.08% for MSV000090572, 98.47% for PXD004720, and 97.90% for PXD050500. These results indicate that the workflow reconstructed valid peptide genomic coordinates for 98.14% of projected peptide/protein pairs across datasets, tissues, exon structures, and strand orientations.

After removal of failed projections and collapse of redundant evidence, the final validated dataset comprised 3,138,903 non-redundant peptide projection rows, including 2,775,671 within-exon and 363,232 intron-spanning projections (Table 1). These represented 1,089,379 unique peptide sequences, 267,166 protein isoforms, and 238,590 gene models. This substantially extends previous wheat proteome-mapping efforts, including the 89,754 unique peptides and 46,016 inferred gene loci reported by Duncan *et al.*^20^ , and the 29,902 HC and 955 LC peptide-supported proteins reported by Liu *et al.*^22^. Our strategy also addressed key limitations of direct peptide-to-genome alignment. Short peptide sequences are poorly suited to BLAST-based alignment because statistical confidence declines with query length^46^, particularly across exon junctions or frame- disrupted regions^47^. In our previous tBLASTn-based workflow^25^, only 92,719 genomic hits were recovered from 861,759 unique peptides, leaving approximately 89% of peptides without genomic assignment. In contrast to discovery-oriented proteogenomic strategies that identify novel peptides, splice events, alternative open reading frames, or unannotated coding regions^12,13,42,43,45,48–51^, the present workflow conservatively validates peptide support for existing IWGSC RefSeq v2.1 HC and LC gene models while preserving peptide-to-gene coordinate relationships for genome-browser visualisation.

### 3.3. Proteogenomic support for wheat HC and LC gene models

The main objective of this study was to provide extensive protein-level support for both HC and LC bread wheat annotations. Overall, 103,095 non-redundant HC gene models and 135,495 LC gene models received peptide-supported projections (Table 1), corresponding to 96.4% and 84.8% of all parsed HC and LC annotations, respectively. Globally, peptide evidence supported 238,590 annotated wheat gene models, representing 89.4% of annotations from IWGSC RefSeq v2.1. This suggests that many LC annotations represent genuinely expressed coding loci that remain classified as low confidence because of incomplete transcript support or unresolved structural ambiguities during original annotation^2^. These loci therefore represent strong candidates for future IWGSC RefSeq curation and potential reclassification to HC status. Similar observations have been reported in other large-scale proteogenomics studies where peptide evidence supports putative or weakly annotated loci lacking robust transcript-level validation^14,50^. Our genome annotation-based projection rates were substantially higher than those obtained in our previous proteogenomics study^25^, where only 31.4% of HC annotations and 2.3% of LC annotations received peptide support through tBLASTn alignment. The dramatic increase observed here likely reflects direct raw-data reprocessing, expanded dataset coverage, GFF3-based coordinate reconstruction, and removal of BLAST-related short-peptide alignment limitations.

Peptide-support distributions further highlighted marked differences between HC and LC annotations (Suppl. Fig. S2). HC gene models generally exhibited broader and more abundant peptide support than LC gene models, with stronger representation among highly supported loci. The relationship between protein length and projected unique peptide count was positive for both annotation classes, yet HC proteins showed greater peptide accumulation across the length range than LC proteins (Suppl. Fig. S2A). Boxplot and histogram distributions confirmed this pattern, with HC gene models more frequently represented among highly supported loci, whereas LC models were enriched among lower-support categories (Suppl. Fig. S2B–C). In particular, 56,117 (54.4%) HC gene models were supported by more than ten unique peptides, compared with 17,439 (12.9%) LC gene models (Suppl. Tables S4). Conversely, 60.6% of LC annotations were supported by only one to four peptides, consistent with their lower-confidence status, compared with 20.3% HC annotations. Of the 3,138,903 non-redundant validated peptide projection rows, 2,409,255 (76.8%) were assigned to HC annotations and 729,648 (23.2%) to LC annotations (Suppl. Fig. S2D; Suppl. Table S5). Most projections in both confidence classes were within-exon, but intron-spanning evidence was proportionally more frequent among HC projections (329,096, 13.7%) than LC projections (34,136, 4.7%) (Suppl. Fig. S2E-F). This suggests that LC loci are often supported by within-exon peptide evidence, but less frequently by peptides spanning annotated exon junctions, which may reflect shorter or less structurally resolved gene models, incomplete transcript support, or remaining uncertainty in LC gene-model architecture^11,52^.

Together, these observations are consistent with the stronger structural and transcriptomic evidence underlying HC annotations, albeit demonstrating that many LC loci possess reproducible protein-level evidence. These loci therefore provide a prioritised set of candidates for future *T. aestivum* genome annotation curation and potential confidence-level reclassification. This is particularly relevant for loci where peptide evidence can be combined with transcriptomic, comparative-genomic, or gene-structure information. Such integration is consistent with high-stringency proteogenomic frameworks, which recommend using peptide evidence alongside orthogonal annotation support for gene-model refinement^41,52^.

Tissue-level analyses further highlighted substantial variation in proteogenomic coverage across datasets and organs (Suppl. Fig. S3; Suppl. Table S6). The three PXD050500 seedling tissues^22^ contributed the highest protein-isoform coverage, with node, coleoptile, and radicle supporting 78.7%, 77.6%, and 66.9% total annotated protein-isoform coverage, respectively. Among the 28 PXD004720 tissues and development stages^20^, the most extensive coverage was observed for grain Zadoks 71 and coleoptile, which supported 15.5% and 15.3% total protein-isoform coverage, respectively. The single MSV000090572 stored-grain dataset supported 5.7%. These coverage differences likely reflect variation in tissue type, proteome complexity, experimental depth, fractionation strategy, instrument platform, and total MS/MS acquisition effort, all of which are known to influence proteome coverage in bottom-up MS-based proteomics^20,53,54^.

UpSet analyses further demonstrated that projected protein isoforms included both shared and tissue-specific components across the datasets (Suppl. Fig. S4; Suppl. Table S6). The largest intersections were dominated by individual tissues and small tissue combinations, whereas intersections shared across many tissues were comparatively smaller. This reinforces that bread wheat proteome detection is strongly influenced by tissue context and analytical depth, and that broad annotation support requires sampling across multiple organs, developmental stages, and proteomic acquisition strategies^20–24^.

Overall, these results show that large-scale peptide projection provides an effective protein-level framework for evaluating HC and LC wheat gene models. The stronger peptide support observed for HC annotations is consistent with their higher-confidence structural classification, whereas the substantial validated support for LC loci indicates that many low-confidence models encode experimentally observed proteins and deserve future annotation review. Peptide evidence have been used to support gene-model refinement, identify translated coding regions, and improve the functional interpretation of complex genomes^20,22,45^. The present workflow extends this principle by linking validated peptide evidence directly to RefSeq v2.1 HC and LC gene models and by providing genome-browser-compatible coordinates for community inspection and curation.

### 3.4. Genome-wide visualisation of peptide projections

Violin plots of peptide genomic coordinates revealed broad chromosomal representation of validated peptide projections across all 21 wheat chromosomes for both HC and LC annotations (Fig. 2). No major chromosome-scale absence of peptide-supported annotations was observed. This genome-wide coverage likely reflects the diversity of the selected public proteomics datasets, which collectively sampled multiple wheat tissues and developmental stages, thereby capturing protein-coding evidence from a broad range of genomic loci. The thinnest regions of the violin profiles generally coincided with chromosome intervals expected to be gene-poor, including centromeric/pericentromeric regions, consistent with the reduced density of protein-coding loci in these regions^2^.

Apollo/JBrowse visualisation confirmed that the validated BED6/BED12 tracks provide interpretable, exon-resolved genomic representations of peptide evidence along annotated *T. aestivum* gene models (Fig. 3). Peptide labels are displayed in their original MS/MS identification orientation from N- to C-terminus, irrespective of genomic strand. Consequently, peptides mapped to reverse-strand loci may appear visually reversed relative to left-to-right genomic coordinates in Apollo/JBrowse; this relates to display orientation rather than incorrect projection. Peptides spanning splice junctions generated continuous BED block patterns corresponding to exon–intron organisation, while dashes preserved intron-spanning peptide labels for manual inspection (Fig. 3A). This is consistent with the summary of validated projection structures, in which exon-spanning peptides represented a substantial subset of HC projections and a smaller but still informative subset of LC projections (Suppl. Fig. S2E–F).

The browser views also illustrated how projected peptide evidence can support different annotation-review scenarios. Several loci showed dense peptide coverage across HC gene models (Fig. 3A), while others demonstrated isoform-specific peptide support, indicating that the workflow can distinguish closely related transcript variants where peptides map to isoform-specific exon structures or protein sequences (Fig. 3B). In addition, LC gene models with extensive peptide support were identified (Fig. 3C), reinforcing the conclusion that some annotations currently classified as low confidence likely represent *bona fide* expressed protein-coding loci and warrant reclassification. MS/MS-derived peptides have confirmed translation of predicted coding regions, support exon and splice-junction annotation, distinguish protein isoforms, and guide gene-model revision^11,52,55^.

Peptides aligning along LC genes provide experimentally validated protein-level support that may assist future annotation review, confidence reassessment, and community curation. This genome-browser visualisation follows previous proteogenomic resources in which MS-derived peptides were mapped to genomic coordinates and displayed as interpretable evidence tracks, including peptide-to-BED tools and web-based viewers linking peptide identifications to exon usage, isoform structure, genomic alignments, and MS/MS evidence^29,40,42,55^. More broadly, deployment within JBrowse/Apollo follows established genome-annotation practice, where browser-based evidence tracks provide a stable framework for visual inspection, collaborative curation, and refinement of gene models.

Together, the genome-wide and locus-level visualisations demonstrate that our strategy converts large-scale MS/MS peptide evidence into a reusable annotation resource. By combining validated peptide-to-genome projections, exon-aware BED structure, intron-spanning labels, and permanent Apollo/JBrowse deployment, the study extends beyond static proteogenomic summary tables to provide the community with a genome-browser tool for inspecting wheat gene-model support, evaluating HC and LC annotations, and prioritising loci for future curation.

## CONCLUSION

This study presents a genome-based wheat proteogenomics workflow integrating large-scale MS/MS reanalysis with GFF3-based peptide projection against the IWGSC RefSeq v2.1 genome. By combining FragPipe/MSFragger peptide identification, GFF3-based coordinate reconstruction, and rigorous projection validation, the workflow provided extensive protein-level support for 238,590 *T. aestivum* gene models, representing 89.4% of all parsed HC and LC annotations. The resulting validated BED tracks deployed in Apollo/JBrowse and associated exploratory analyses provide a reusable community resource for wheat genome annotation refinement. More broadly, this annotation-based framework offers a scalable strategy for proteogenomic analysis in large and complex polyploid plant genomes.

## ASSOCIATED CONTENT

### Supporting Information

- Supplementary File S1: Jupyter Notebook markdown file
- Supplementary File S2: FragPipe workflow and MSFragger search parameters
- Supplementary File S3: Peptide genomic coordinate reconstruction and validation steps
- Supplementary Table S1: Complete validated peptide projection resource summary
- Supplementary Table S2: Translation validation summary
- Supplementary Table S3: Quality control check failures
- Supplementary Table S4: HC and LC gene-model peptide support statistics
- Supplementary Table S5: Within-exon and intron-spanning peptide support statistics
- Supplementary Table S6: Tissue-level HC/LC proteogenomic coverage summary
- Supplementary Figure S1: Computational workflow of the genome-based wheat proteogenomics approach devised in this study.
- Supplementary Figure S2: Peptide support distributions across HC and LC wheat gene models.
- Supplementary Figure S3: Proteogenomic coverage across wheat tissues
- Supplementary Figure S4: UpSet plot of projected protein intersections across wheat tissues.

### Data availability

The complete reproducible wheat proteogenomics workflow, including JupyterLab notebooks, Python scripts, BED6/BED12 genome browser tracks, workflow documentation, and computational environment specifications, is publicly available at: https://github.com/dlf2024/Python-wheat-proteogenomics_2026

Apollo/JBrowse server: https://bread-wheat-um.genome.edu.au/apollo/49826/jbrowse/index.html AUTHOR INFORMATION

### Author Contributions

Conceptualization, D.V.; methodology, D.V.; software and resources, D.V.; formal analysis, D.V.;validation, D.V.; visualization, D.V.; writing—original draft preparation, D.V.; writing—review and editing, D.V. and R.A.; supervision, D.V.; project administration, D.V.; permanent upload of proteogenomic peptides to Apollo Jbrowse: R.A.; funding: R.A. The manuscript was written through contributions of all authors. All authors have given approval to the final version of the manuscript.

### Funding Sources

Private funds were used in this study.

## ACKNOWLEDGMENT

The authors acknowledge the valuable assistance provided by colleagues in the Australia BioCommons (https://www.biocommons.org.au/team), in particular we thank Mike Thang for permanently uploading peptide BED12 tracks to bread wheat Apollo/Jbrowse server and thus enabling the viewing of proteogenomic results to the wider community.

AA: Amino Acid
BED: Browser Extensible Data
BLAST: Basic Local Alignment Search Tool
CDS: Coding DNA Sequence
cRAP: Common Repository of Adventitious Proteins
DDA: Data-dependent acquisition
EDA: Exploratory data analysis
FDR: False discovery rate
FTP: File Transfer Protocol
GFF3: General Feature Format version 3
HC: High confidence
IWGSC: International Wheat Genome Sequencing Consortium
JBrowse: JavaScript genome browser
QC: Quality Control
LC: Low confidence
MassIVE: Mass Spectrometry Interactive Virtual Environment repository
MS/MS: tandem mass spectrometry
mzML: Mass spectrometry markup language
PRIDE: Proteomics Identifications Database
PSM: Peptide-spectrum match
RefSeq: Reference Sequence
TB: Terabyte
tBLASTn: Translated Basic Local Alignment Search Tool against nucleotide database

