## Supplementary material for "Community Resource: A Genome-Based Extension of Large-Scale Wheat Proteogenomics": Vincent_wheat-proteogenomics_technical-note_2026-06-16_Suppl-Figures.pdf

Graphical Abstract

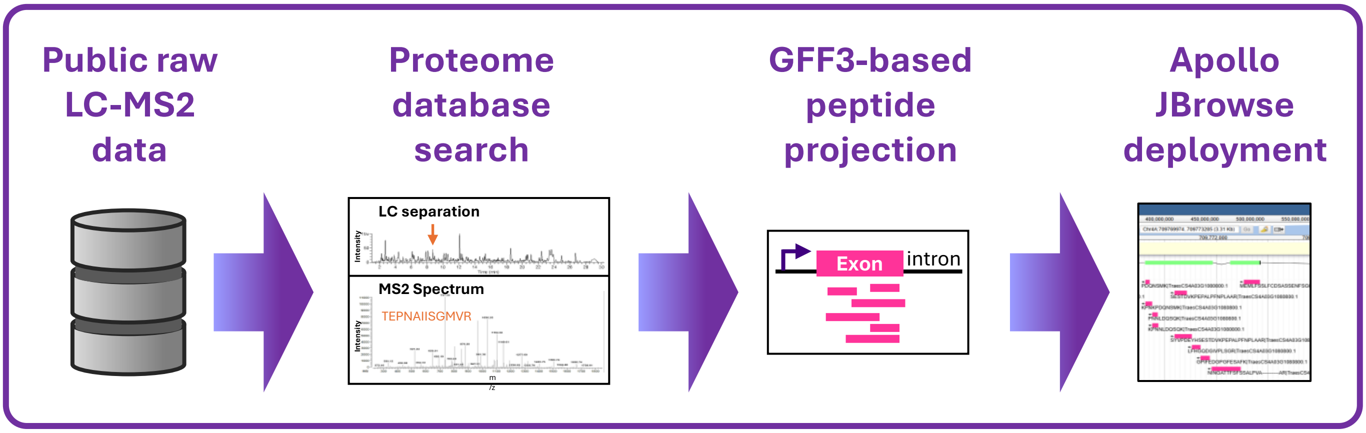

**Supplementary Figure S1.** Computational workflow of the genome annotation-based wheat proteogenomics approach devised in this study. Public LC–MS/MS datasets from PRIDE and MassIVE were retrieved and reprocessed against the IWGSC RefSeq v2.1 HC+LC wheat proteome database using FragPipe/MSFragger. GFF3 annotations were parsed to reconstruct protein–transcript–gene relationships and exon/CDS coordinates. Non-contaminant peptide evidence was integrated with GFF3-derived annotation frameworks for genome projection, whereby peptide amino-acid coordinates were converted into exon-aware genomic coordinates. Projection validation involved translation-based peptide reconstruction together with independent BED geometry, chromosome/strand, and protein-coordinate as further quality control checks. Validated peptide projectons were used for genome-scale exploratory data analysis (EDA) and exported as BED6/BED12 tracks for Apollo/JBrowse deployment. Detailed computational steps and Python code are provided in the accompanying Jupyter Notebook and GitHub repository, as well as Supplementary File S3.

### Genome-guided wheat proteogenomics workflow

#### 1. DATA RETRIEVAL + REFERENCE GENOME + PROTEOME SEARCH

*Steps 1–4*

FTP download of raw LC–MS2 proteomic and GFF3 genomic files (*Unix*)
Data conversion (*ProteoWizard MSConvert*)
Protein database creation with decoy (*Galaxy)*
Proteome search against IWGSC v2.1 database (*FragPipe: MSFragger → PeptideProphet → ProteinProphet*)

#### 2. GFF3 ANNOTATION FRAMEWORK

*Step 5*

ProteinID → TranscriptID → GeneModel → CDS blocks (*Python*)

#### 3. PEPTIDE EVIDENCE CONSTRUCTION

*Steps 6–8*

FragPipe output importation (*Python*)
Non-contaminant peptide–protein–gene evidence (*Python*)
Protein to gene model projections using GFF3-derived table (*Python*)

#### 4. GFF3-BASED PEPTIDE PROJECTION

*Step 9*

Peptide in protein → coding nucleotide coordinates → exon-aware genomic blocks (*Python*)

**Protein sequence**
MKWVTFPEPTIDEISLLFSSAY…
↓
**Peptide CDS coordinates**
nt 241–312
↓
**GFF3 exon blocks**
Exon1 – Intron – Exon2
↓
**Genomic peptide projection**
▬▬-----▬▬

1. Locates the peptide amino acid sequence within the corresponding protein sequence.
2. Converts the peptide amino acid interval into coding nucleotide positions.
3. Retrieves the CDS blocks associated with the corresponding transcript.
4. Projects coding nucleotide positions onto genomic coordinates.
5. Collapses projected nucleotide positions into genomic blocks.
6. Generates an intron-aware peptide display label (exon-spanning junctions shown using dash characters).

#### 5. PEPTIDE PROJECTION VALIDATION

*Steps 10, 11*

Translation validation: projected genomic blocks → translated peptide → match/mismatch QC (*Python*)
Independent sanity checks (*Python*)

**Projected genomic blocks**
▬▬-----▬▬
↓
**Retrieved genomic DNA**
ATGGCC…GT…AAGTAA
↓
**Translated peptide**
NINGATTFFSSALPVA
↓
**Original peptide**
= NINGATTFFSSALPVA
↓
**Evaluation**
☑ match

Beside the translation validation, 3 independent sanity checks are performed:

1. BED geometry check: confirms BED coordinates by checking start coordinates < end coordinates, and that BED block sizes and block starts are consistent with each peptide genomic interval.
2. Chromosome and strand check: confirms that projected peptides are assigned only to expected wheat chromosomes and valid strand values (`+` or `-`).
3. Protein-coordinate consistency check: confirms that each peptide projection has coherent amino-acid coordinates, including matching peptide length and valid protein coordinate intervals.

**Evaluation**:

- translated sequence match + successful checks → peptide validated ☑
- translated sequence mismatch and/or unsuccessful check → peptide rejected ☒

#### 6. RESOURCE OUTPUT + INTERPRETATION

*Steps 12-25*

validated projections exported as BED tracks (*Python*)
BED tracks permanent upload (*Apollo Jbrowse)*
EDA (scatter, box, violin, bar, UpSet, circular plots, histograms, Venn diagram, pie chart) (*Python*)
HC/LC support (*Python*)

**Supplementary Figure S2:** Peptide support distributions across HC and LC wheat gene models. (A) Scatterplot showing the relationship between protein length and the number of projected unique peptides per protein isoform, separated by HC and LC annotation confidence. (B) Boxplot comparing projected peptide support between HC and LC gene models. (C) Distribution of projected unique peptides per gene model showing broader and more abundant support for HC annotations. (D) Proportion of non-redundant validated peptide projections assigned to HC and LC gene models. (E) Proportion of HC peptide projections classified as within-exon or exon-spanning. (F) Proportion of LC peptide projections classified as within-exon or exon-spanning.

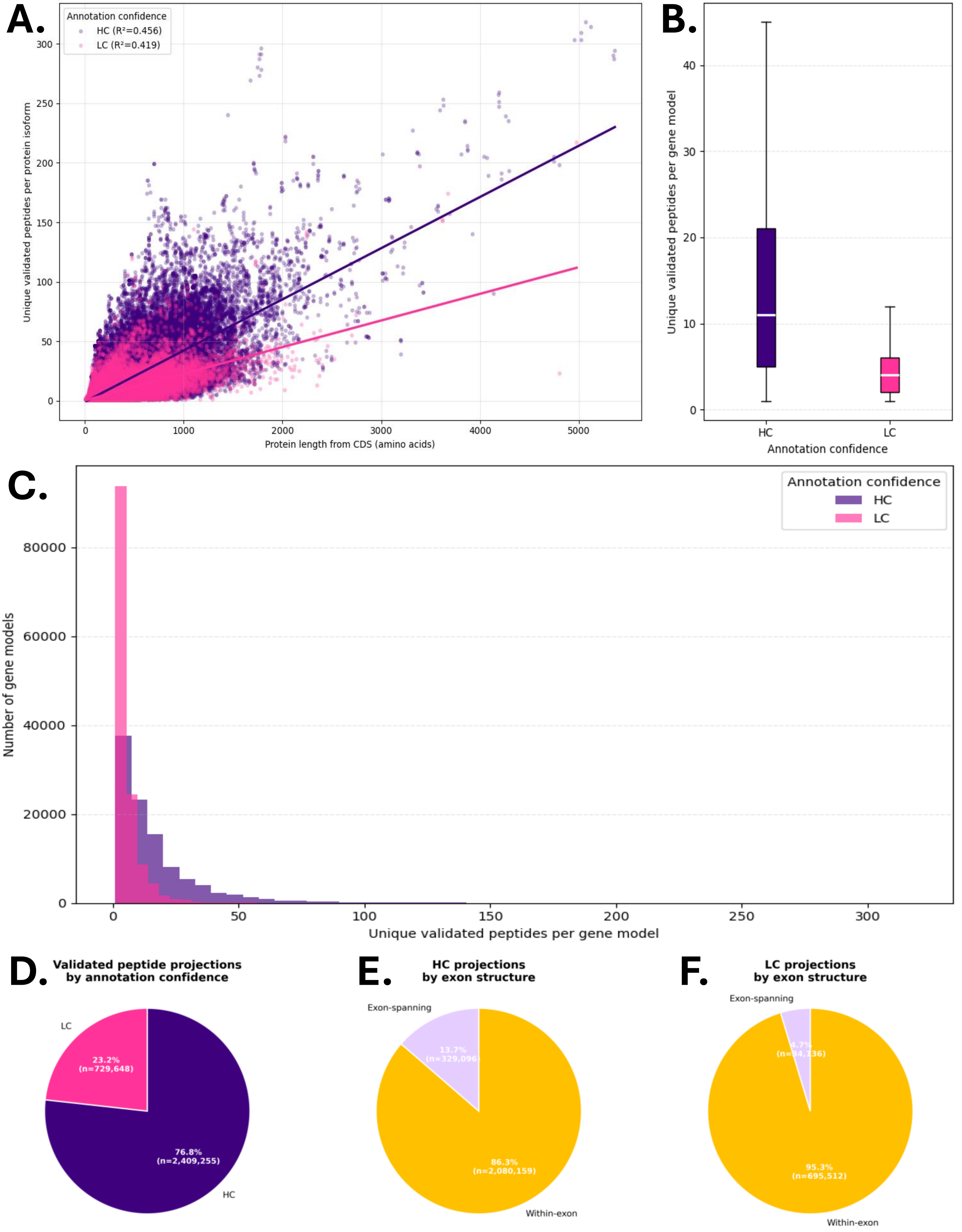

**Supplementary Figure S3.** Proteogenomic coverage across wheat tissues. Barplot of protein-level coverage showing the percentage of annotated HC and LC gene models supported by projected protein accessions.

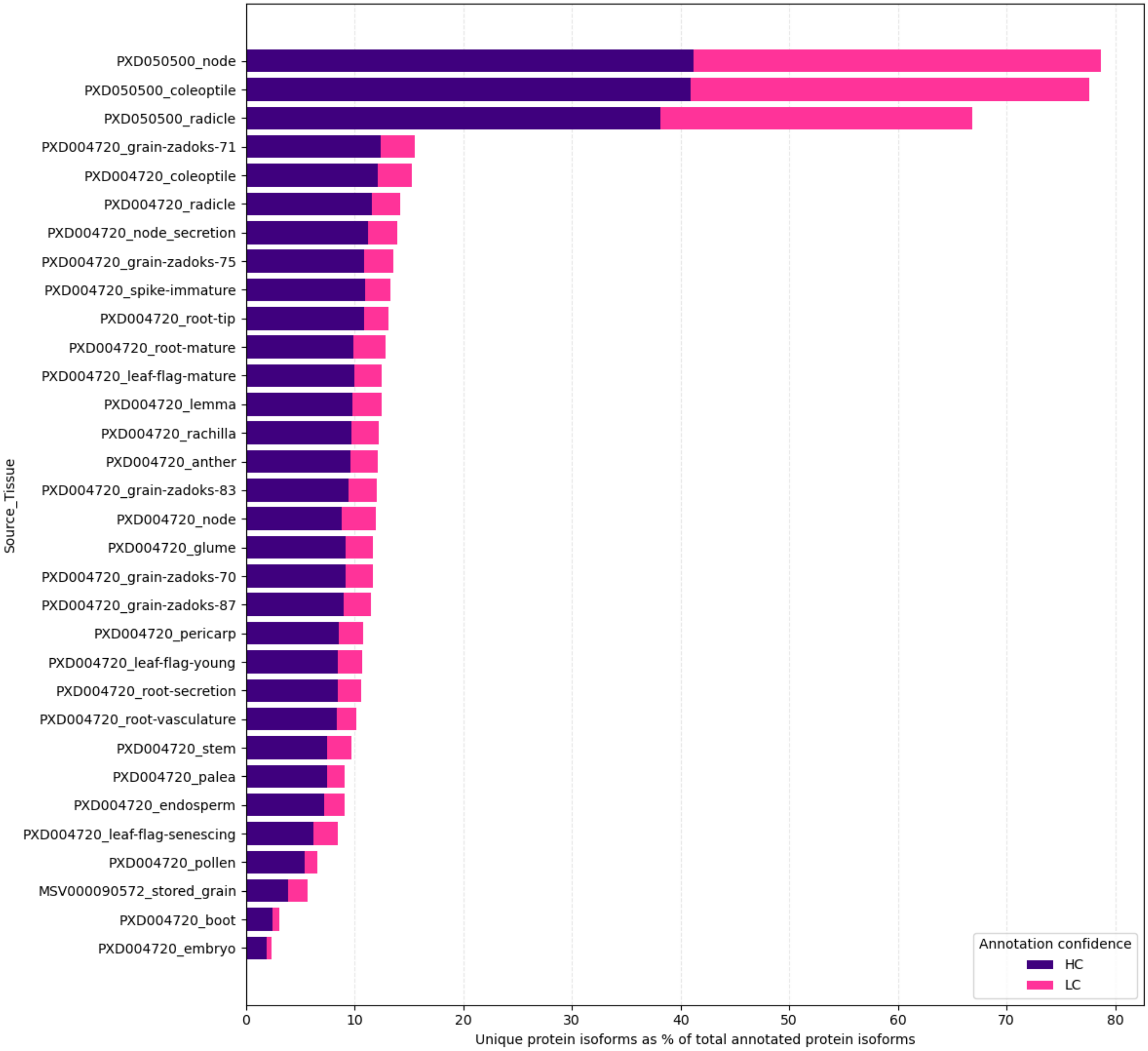

**Supplementary Figure S4:** UpSet plot of projected protein intersections across wheat tissues. Vertical bars indicate the number of shared projected proteins among tissue combinations, while horizontal bars show the total number of projected proteins identified per tissue. The analysis highlights both broadly shared and tissue-specific proteogenomic signatures across the wheat datasets.

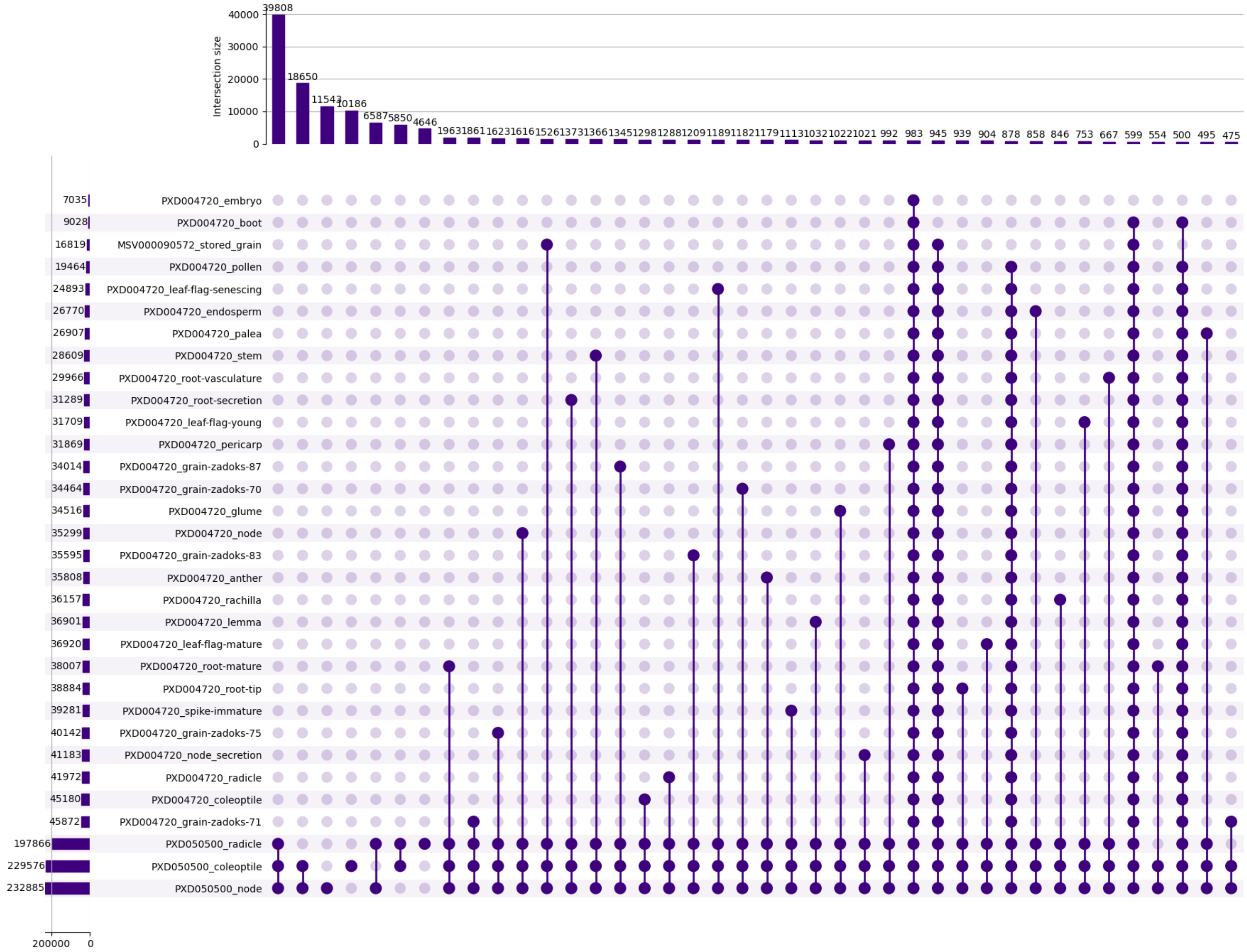
