## Supplementary material for "Community Resource: A Genome-Based Extension of Large-Scale Wheat Proteogenomics": Vincent_wheat-proteogenomics_technical-note_2026-06-16_Suppl-File-S1.md

# Community Resource: A Genome-Guided Extension of Large-Scale Wheat Proteogenomics

---

## Author   
Dr Delphine Vincent  
_website:_ https://dlf2024.github.io/  
_github:_ https://github.com/dlf2024      
_Date:_ 02/06/2026

---

## Rationale  
This project aims at validating gene model annotations in bread wheat (_Triticum aestivum_, IWGSC RefSeq v2.1) using a proteogenomics strategy by identifying peptides/proteins in various wheat tissues from public proteomics experiments and aligning them along the genome.     
This study follows on our previous proteogenomics project from 2024 in which peptides identified in public repositories MSV000090572 and PXD004720 were aligned along the wheat genome using a tBLASTn strategy (Citation: Vincent, D.; Appels, R. (2024) Community Resource: Large-Scale Proteogenomics to Refine Wheat Genome Annotations. Int. J. Mol. Sci. 2024, 25, 8614. https://doi.org/10.3390/ijms25168614).    
The peptide alignments along bread wheat genome are publicly available from the Apollo JBrowse service (https://bread-wheat-um.genome.edu.au/apollo/49826/jbrowse/index.html).   

---

## Steps  
- 1: Data retrieval and extraction (LFTP via Ubuntu Unix outside of this notebook)
- 2: Data conversion (using MSConvert GUI outside of this notebook)
- 3: Protein database creation with decoy (using Galaxy Proteomics platform outside of this notebook)
- 4: Peptide/Protein search (using FragPipe program outside of this notebook)
- 5: Build a protein-to-gene mapping table (this notebook)
- 6: FragPipe Output Annotation and Quality Summary (this notebook)
- 7: Build non-contaminant peptide–protein evidence tables (this notebook)
- 8: Map proteins to gene models using the GFF3-derived mapping table (this notebook)
- 9: Project peptide positions from proteins onto genomic coordinates (this notebook)
- 10: Translation validation of annotation-guided peptide genome projections (this notebook)
- 11: Sanity Checks for translation-validated peptide Genome Projections (this notebook)
- 12: Export all tissues BED6/BED12 files for JBrowse (this notebook)
- 13: Create a single non-redundant combined BED track (this notebook)
- 14: Prepare BED Files for Apollo/JBrowse Public Upload (this notebook)
- 15: Generate tissue/protein/gene/isoform summary tables (this notebook)
- 16: EDA: Barplot of HC and LC Proteogenomic Coverage by Tissue (this notebook)
- 17: EDA: Tissue Overlap Using UpSet Plots (this notebook)
- 18: EDA: Distributions of Peptide Support per Gene Model (this notebook)
- 19: EDA: Scatterplot of Peptide Length and Probability by Annotation Confidence  (this notebook)
- 20: EDA: Scatterplot of Protein Length versus Peptide Support (this notebook)
- 21: EDA: Violin plot of Chromosomal Distribution of Peptide Genomic Start Positions (this notebook)
- 22: EDA: Circular plot of Tissue-Level Peptide Genome Map (this notebook)
- 23: EDA: Pie chart of HC and LC Proteogenomic Coverage with Within-Exon/Exon-spanning Peptides (this notebook)
- 24: Combine Python workflow Summary Tables at Source/Tissue Level (this notebook)
- 25: Prepare manuscript Table 1 summary statistics (this notebook)

---

## Wheat Genome Annotation Source  
- Citation: The International Wheat Genome Sequencing Consortium (IWGSC); Appels, R. et al. (2018) Shifting the Limits in Wheat Research and Breeding Using a Fully Annotated Reference Genome. Science, 361, eaar7191. https://doi.org/10.1126/science.aar7191  
- IWGSC URL: https://urgi.versailles.inrae.fr/download/iwgsc/IWGSC_RefSeq_Annotations/v2.1/
- Data files:
  - iwgsc_refseqv2.1_annotation_200916_LC.gff3 (genome annotations of low confidence (LC) gene models)
  - iwgsc_refseqv2.1_annotation_200916_HC.gff3 (genome annotation of high confidence (HC) gene models)
  - iwgsc_refseqv2.1_functional_annotation.csv (gene model functional annotations)
  - iwgsc_refseqv2.1_annotation_200916_HC_pep.fasta (AA sequences of HC gene models)
  - iwgsc_refseqv2.1_annotation_200916_LC_pep.fasta (AA sequences of LC gene models)

---

## MS-Proteomics Raw Data Sources 

### Liu et al 2025 (PXD050500)  
- Citation: Liu S, et al (2025) A telomere-to-telomere genome assembly coupled with multi-omic data provides insights into the evolution of hexaploidy bread wheat. Nature Genetics, 57: 1008-1020. https://doi.org/10.1038/s41588-025-02137-x   
- ProteomeXchange URL: https://proteomecentral.proteomexchange.org/cgi/GetDataset?ID=PXD050500   
- PRIDE URL: https://www.ebi.ac.uk/pride/archive/projects/PXD050500    
- FTP URL: https://ftp.pride.ebi.ac.uk/pride/data/archive/2024/12/PXD050500/   
- Data files: 180 .d files (timsTOF Pro mass spectrometer, Bruker Daltonics) - 400 GB
- Tissues: 
  - node (60 fractions)
  - coleoptile (60 fractions)
  - radicle (60 fractions)

### Vincent et al 2023 (MSV000090572)    
- Citation: Vincent, D. et al. (2023) A Community Resource to Mass Explore the Wheat Grain Proteome and Its Application to the Late-Maturity Alpha-Amylase (LMA) Problem. GigaScience, 12, giad084. https://doi.org/10.1093/gigascience/giad084     
- MassIVE URL: https://massive.ucsd.edu/ProteoSAFe/result.jsp?task=9e8c4c3c9d924de8800237e7e828e1d9&view=advanced_view#%7B%7D   
- FTP URL: ftp://massive-ftp.ucsd.edu/v05/MSV000090572/
- Data files: 62 .raw files (LTQ Orbitrap Elite mass analyser, Thermo Scientific) - 12 GB
- Tissues:
  - stored grains (mature, harvested grains stored in optimum conditions)

### Duncan et al 2017 (PXD004720)     
- Citation: Duncan, O. et al (2017) Resource: Mapping the Triticum aestivum Proteome. Plant J., 89, 601-616. https://doi.org/10.1111/tpj.13402      
- ProteomeXchange URL: https://proteomecentral.proteomexchange.org/ui?pxid=PXD004720    
- PRIDE URL: https://www.ebi.ac.uk/pride/archive/projects/PXD004720   
- FTP URL: https://ftp.pride.ebi.ac.uk/pride/data/archive/2016/11/PXD004720/
- Data files: 335 .mzXML files (QTOF, Agilent) - 600 GB
- Tissues:
  - anther (12 fractions)
  - boot (12 fractions)
  - coleoptile (12 fractions)
  - embryo (12 fractions)
  - endosperm (12 fractions)
  - glume (12 fractions)
  - grain-zadoks-70 (12 fractions)
  - grain-zadoks-71 (12 fractions)
  - grain-zadoks-75 (12 fractions)
  - grain-zadoks-83 (12 fractions)
  - grain-zadoks-87 (12 fractions)
  - leaf-flag-mature (12 fractions)
  - leaf-flag-senescing (12 fractions)
  - leaf-flag-young (12 fractions)
  - lemma (12 fractions)
  - node (12 fractions)
  - node_secretion (12 fractions)
  - palea (11 fractions)
  - pericarp (12 fractions)
  - pollen (12 fractions)
  - rachilla (12 fractions)
  - radicle (12 fractions)
  - root-mature (12 fractions)
  - root-secretion (12 fractions)
  - root-tip (12 fractions)
  - root-vasculature (12 fractions)
  - spike-immature (12 fractions)
  - stem (12 fractions)

---

## Computational environment  
- Windows 10
- WSL2 Ubuntu 24.04
- Python 3.12
- JupyterLab 4.3.4
- FragPipe 24.0
- MSFragger 4.4.1
- ProteoWizard MSConvert 3.0
- Galaxy Proteomics platform

---

## Folder architecture  
project/   
├── raw_data/    
├── mzML/    
├── genome_annotation/    
├── protein_database/    
├── fragpipe_results/    
├── tblastn_results/  
├── python_outputs/    
│      ├── bed/    
│      ├── tables/    
│      └── figures/  

---

## AI-assisted workflow development   
Portions of the Python workflow, code optimisation, debugging assistance, and markdown documentation were developed with the assistance of OpenAI ChatGPT (GPT-5 series) under the direction and scientific supervision of the author.   
All computational workflows, parameter selections, biological interpretations, quality control, and final validation were performed and verified by the author.  

---

## Apollo JBrowse Deployment

Projected peptide alignments generated from this workflow are publicly available through the Apollo JBrowse genome browser:  

https://bread-wheat-um.genome.edu.au/apollo/49826/jbrowse/index.html

Tracks include:
- annotation-guided peptide genome projections
- BED6 and BED12 visualisation tracks
- tissue-specific peptide mappings
- non-redundant combined peptide tracks 

# Step 1 — Data Retrieval and Extraction

Large-scale public proteomics datasets were retrieved from the PRIDE and MassIVE repositories using the Linux command-line utility `lftp` executed within a Ubuntu WSL2 environment under Windows 11. Due to the large data volume (>1 TB total), data management and storage optimisation were critical considerations throughout the workflow.

---

## Computational environment

### Operating system
- Windows 11 Pro
- WSL2 Ubuntu Linux

### Software
- `lftp`
- `7zip`
- Windows PowerShell

### Install required Linux utilities

```bash
sudo apt update
sudo apt install lftp p7zip-full
```

---

## Data retrieval using lftp (e.g. PXD050500)

### Create new directory to store files

```bash
mkdir -p /mnt/e/wheat_proteogenomics/PXD050500
cd /mnt/e/wheat_proteogenomics/PXD050500
```

### Connect to public repository (PRIDE)

```bash
lftp ftp.pride.ebi.ac.uk
```

### Inside the `lftp` session, download files

```bash
cd /pride/data/archive/2024/12/PXD050500
mirror --verbose .
exit
```

---

## Monitoring storage usage in WSL2

Large downloads can rapidly consume WSL2 virtual disk space.

### Check Linux folder size

```bash
du -sh /root/wheat_pride/PXD004720
```

### Check available disk space

```bash
df -h
```

### Cleaning temporary files

```bash
rm -rf /tmp/*
```

---

## Archive extraction

### Test archive integrity before extraction

```bash
7z t filename.rar
```

### Extract archive

```bash
7z x filename.rar
```

### Remove extracted archives no longer needed

```bash
rm filename.rar
```

---

## WSL2 disk cleanup

After deleting large datasets, WSL2 virtual disks may still occupy substantial space on the Windows drive.

### Shutdown WSL2

```powershell
wsl --shutdown
```

### Locate the virtual disk

```powershell
Get-ChildItem "$env:LOCALAPPDATA\Packages" -Recurse -Filter ext4.vhdx
```

---

## Notes

- Download times varied from several hours to multiple days depending on repository speed and internet bandwidth.
- Stable internet connection and uninterrupted power supply were strongly recommended.
- External SSD storage substantially improved workflow efficiency compared to mechanical HDD devices.

# Step 2 — Data Conversion using MSConvert

Some raw mass spectrometry files required conversion into the open standard `.mzML` format prior to peptide identification analysis in FragPipe. File conversion was performed using the ProteoWizard software suite (`MSConvert GUI`) under Windows 11.

### Software
- ProteoWizard MSConvert
- URL: https://proteowizard.sourceforge.io/

---

## Input formats

### PXD004720
- Input format: `.mzXML`
- Instrument: Agilent QTOF

### MSV000090572
- Input format: `.raw`
- Instrument: Thermo Orbitrap Elite

### PXD050500
- Input format: `.d`
- Instrument: Bruker timsTOF Pro
- No conversion required for FragPipe analysis

---

## MSConvert GUI workflow

### 1. Launch MSConvert GUI

Open:
```text
ProteoWizard → MSConvert
```

### 2. Select input files

- Click `Browse`
- Select raw data files
- Add files to processing queue

### 3. Select output directory

Choose destination folder for converted files.

Example:
```text
E:\wheat_proteogenomics\PXD004720\mzML\
```

---

## Recommended conversion settings

### Output format
```text
mzML
```

### Binary encoding precision
```text
64-bit
```

### Compression
```text
zlib compression
```

### Peak picking
```text
Vendor peak picking
```

### Use zlib compression
```text
Enabled
```

### Write index
```text
Enabled
```

### TPP compatibility
```text
Enabled
```

---

## Start conversion

Click:
```text
Start
```

Converted `.mzML` files were subsequently used as input for FragPipe peptide identification workflows.

---

## Notes

- File conversion duration depended on instrument type and file size.
- Some Agilent `.mzXML` files from PXD004720 were unreadable by FragPipe and required reconversion using MSConvert.
- Output `.mzML` files substantially increased compatibility and stability during downstream peptide identification analyses.

# Step 3 — FASTA Database Creation

A combined target-decoy protein database was created using Galaxy Proteomics by merging high-confidence (HC) and low-confidence (LC) wheat protein annotations together with the common Repository of Adventitious Proteins (cRAP) contaminant database.

## Wheat protein sequence source

- IWGSC URL:
  https://urgi.versailles.inrae.fr/download/iwgsc/IWGSC_RefSeq_Annotations/v2.1/

### Input FASTA files

```text
iwgsc_refseqv2.1_annotation_200916_HC_pep.fasta
iwgsc_refseqv2.1_annotation_200916_LC_pep.fasta
```

---

## Software

### Galaxy Proteomics platform

- Galaxy URL:
  https://proteomics.usegalaxy.eu/


---

## Upload wheat FASTA files

### Open tool

```text
Upload Data
```

### Workflow

- Click `Choose local file`
- Select HC and LC FASTA files
- Click `Start`

---

## Download cRAP contaminant database

The cRAP database contains common mass spectrometry contaminants such as keratins and trypsin.

### Open tool

```text
Protein Database Downloader
```

### Settings

```text
Database = cRAP (contaminants)
```

### Run tool

Click:

```text
Run Tool
```

---

## Merge FASTA databases and remove duplicate entries

### Open tool

```text
FASTA Merge Files and Filter Unique Sequences
```

### Settings

#### Merge strategy

```text
Merge all FASTAs (always output a single FASTA)
```

#### Input files

```text
HC + LC + cRAP FASTA files
```

#### Duplicate filtering mode

```text
Accession Only
```

#### Parsing rule

```text
^>([^ ]+).*$
```

### Run tool

Click:

```text
Run Tool
```

---

## Create decoy database

A reverse-sequence decoy database was generated for false discovery rate (FDR) estimation during peptide identification.

### Open tool

```text
Create Decoy Database (reverse)
```

### Settings

#### Include original sequences

```text
Yes
```

#### Decoy prefix

```text
decoy_
```

### Run tool

Click:

```text
Run Tool
```

---

## Download final protein database

Download the generated target-decoy FASTA database locally.

### Final database filename

```text
iwgsc_refseqv2.1_annotation_200916_HC_LC_pep_cRAP_with_DECOY.fasta
```

---

## Notes

- HC = high-confidence gene models.
- LC = low-confidence gene models.
- Reverse decoy sequences were used for downstream FDR estimation in FragPipe.
- Inclusion of the cRAP database improved contaminant identification and filtering during peptide searches.

# Step 4 — Peptide and Protein Search using FragPipe

Peptide and protein identification was performed using FragPipe with MSFragger against the combined wheat HC/LC target-decoy protein database containing cRAP contaminant sequences. All tissues were processed using the same FragPipe workflow and search parameters.

---

## Software

### FragPipe workflow version

```text
FragPipe version: 24.0
MSFragger version: 4.4.1
Philosopher version: 5.1.3-RC9
ProteinProphet: enabled
PeptideProphet: enabled
Percolator: disabled
IonQuant: disabled
```

---

## Input files

FragPipe searches were performed on the converted or native mass spectrometry files for each tissue.

### Input formats

```text
PXD050500    Bruker .d folders
MSV000090572 Thermo .raw files
PXD004720    mzML files converted from mzXML
```

---

## Protein database

The following combined target-decoy FASTA database was used:

```text
iwgsc_refseqv2.1_annotation_200916_HC_LC_pep_cRAP_with_DECOY.fasta
```

### Decoy prefix

```text
decoy_
```

---

## Workflow setup

For each tissue, a separate FragPipe run was created.

### Example output directory

```text
C:\Users\Delphine_2026\FragPipe_results\PDX004720_stem_run01
```

### Computational settings

```text
Threads: 4
RAM: 32 GB
Data type: DDA
```

---

## MSFragger search parameters

### Enzyme specificity

```text
Enzyme: Trypsin/P
Cleavage residues: K, R
Cleavage direction: C-terminal
Number of enzymatic termini: 2
Allowed missed cleavages: 10
```

### Peptide length and mass range

```text
Minimum peptide length: 6 aa
Maximum peptide length: 100 aa
Peptide mass range: 300–10,000 Da
```

### Precursor mass tolerance

```text
Precursor mass tolerance: ±10 ppm
True precursor mass tolerance: ±20 ppm
Isotope error: 0/1/2
Precursor mass mode: selected
```

### Fragment mass tolerance

```text
Fragment mass tolerance: ±0.05 Da
Fragment ion series: b, y
Maximum fragment charge: 10
```

### Precursor charge

```text
Charge range: 1–4
Override charge: disabled
```

---

## Modifications

### Fixed modification

```text
Cysteine carbamidomethylation: +57.02146 Da
```

### Variable modifications

```text
Methionine oxidation: +15.9949 Da
Protein N-terminal acetylation: +42.0106 Da
Protein N-terminal methionine clipping: enabled
```

### Variable modification limits

```text
Maximum variable modifications per peptide: 3
Maximum modified forms per peptide: 5
```

---

## Spectrum processing

```text
Deisotoping: enabled
Neutral loss removal: enabled
Remove precursor peak: enabled
Precursor removal range: -1.5 to +1.5 Th
Minimum peaks per spectrum: 15
Top N peaks used: 150
Minimum matched fragments: 4
Minimum fragments for modelling: 2
```

---

## Validation and reporting

### Validation tools

```text
PeptideProphet: enabled
ProteinProphet: enabled
Percolator: disabled
```

### Report settings

```text
Protein FDR: 1%
Sequential filtering: enabled
Peptide-level summary: enabled
Protein-level summary: enabled
Decoys printed in final report: no
Contaminants removed in final report: yes
```

---

## FragPipe outputs used downstream

The main FragPipe output files used for downstream annotation and genome projection were:

```text
psm.tsv
peptide.tsv
protein.tsv
combined_protein.tsv
experiment_annotation.tsv / sdrf.tsv
```

These files were subsequently imported into Python for annotation, contaminant filtering, peptide–protein association, protein-to-gene mapping, and peptide genomic coordinate projection.

---

## Notes

- Each tissue was processed independently to simplify batch management and avoid excessive memory usage.
- The same FragPipe workflow and MSFragger parameters were applied across all 32 tissues.
- The workflow was optimised for peptide/protein identification rather than label-free quantification.
- Matched fragment export was disabled to reduce output size and improve run stability.
- Output files are captured in a file manifest "wheat_tissues_FragPipe-result-manifest_2026-05-11.csv"

# Step 5 — Build a protein-to-gene mapping table
_inputs:_   
- iwgsc_refseqv2.1_annotation_200916_HC.gff3
- iwgsc_refseqv2.1_annotation_200916_LC.gff3
- iwgsc_refseqv2.1_functional_annotation.csv

__Aim:__  
Map each protein accession / gene model to:  
- chromosome
- gene coordinates
- transcript/CDS model
- HC/LC status
- functional annotation

__Summary:__  
_Parse the wheat GFF3 annotations and build a protein-to-gene mapping table_  
This step parses the high-confidence (HC) and low-confidence (LC) wheat GFF3 annotation files and extracts the structural relationships among genes, transcripts, and CDS features.   
The goal is to create a clean annotation table linking protein accessions and gene models to chromosome, strand, genomic coordinates, and annotation confidence level.   
This table will then be merged into the FASTA-validated peptide–protein evidence in the next step.


```python
#################### Wheat genome data ########################

# ============================================================
# Parse HC + LC GFF3 files and build protein-to-gene mapping table (takes 5 min)
# ============================================================

import pandas as pd
import numpy as np
import re

# -----------------------------
# 1. Input / output file names
# -----------------------------
hc_gff3 = "genome_annotation/iwgsc_refseqv2.1_annotation_200916_HC.gff3"
lc_gff3 = "genome_annotation/iwgsc_refseqv2.1_annotation_200916_LC.gff3"

features_out = "python_outputs/tables/wheat_gff3_parsed_features_HC_LC.csv"
mapping_out  = "python_outputs/tables/wheat_protein_gene_mapping_HC_LC.csv"
summary_out  = "python_outputs/tables/wheat_gff3_parsing_summary_HC_LC.csv"

# -----------------------------
# 2. Helper functions
# -----------------------------
def parse_attributes(attr_text):
    attrs = {}
    if pd.isna(attr_text):
        return attrs

    for item in str(attr_text).strip().split(";"):
        if not item.strip():
            continue
        if "=" in item:
            key, value = item.split("=", 1)
            attrs[key.strip()] = value.strip()
        else:
            attrs[item.strip()] = ""
    return attrs

def strip_prefix(value):
    if value is None or pd.isna(value):
        return np.nan
    value = str(value).strip()
    value = re.sub(r"^[A-Za-z_]+:", "", value)
    return value

def split_multi(value):
    if value is None or pd.isna(value):
        return []
    return [strip_prefix(x.strip()) for x in str(value).split(",") if x.strip()]

def gene_model_from_protein(protein_id):
    if protein_id is None or pd.isna(protein_id):
        return np.nan
    protein_id = str(protein_id).strip()
    return re.sub(r"\.\d+$", "", protein_id)

def normalise_feature_type(ftype):
    ftype = str(ftype)
    if ftype in {"mRNA", "transcript"}:
        return "transcript"
    return ftype

# -----------------------------
# 3. GFF3 parser
# -----------------------------
def parse_gff3(filepath, confidence_label):
    rows = []

    with open(filepath, "r", encoding="utf-8", errors="ignore") as f:
        for line in f:
            if not line.strip() or line.startswith("#"):
                continue

            parts = line.rstrip("\n").split("\t")
            if len(parts) != 9:
                continue

            seqid, source, feature_type, start, end, score, strand, phase, attributes = parts
            feature_type = normalise_feature_type(feature_type)
            attrs = parse_attributes(attributes)

            rows.append({
                "SeqID": seqid,
                "Source_gff": source,
                "FeatureType": feature_type,
                "Start": int(start),
                "End": int(end),
                "Score": score if score != "." else np.nan,
                "Strand": strand,
                "Phase": phase if phase != "." else np.nan,
                "Attributes": attributes,
                "Annotation_confidence": confidence_label,
                "ID_raw": attrs.get("ID", np.nan),
                "Parent_raw": attrs.get("Parent", np.nan),
                "Name": attrs.get("Name", np.nan),
                "biotype": attrs.get("biotype", attrs.get("gene_biotype", np.nan)),
                "protein_id_raw": attrs.get("protein_id", np.nan),
                "Derives_from_raw": attrs.get("Derives_from", np.nan),
            })

    df = pd.DataFrame(rows)

    df["ID"] = df["ID_raw"].apply(strip_prefix)
    df["Parent"] = df["Parent_raw"].apply(lambda x: ", ".join(split_multi(x)) if pd.notna(x) else np.nan)
    df["protein_id"] = df["protein_id_raw"].apply(strip_prefix)
    df["Derives_from"] = df["Derives_from_raw"].apply(lambda x: ", ".join(split_multi(x)) if pd.notna(x) else np.nan)

    return df

# -----------------------------
# 4. Parse HC + LC GFF3 files
# -----------------------------
hc_df = parse_gff3(hc_gff3, "HC")
lc_df = parse_gff3(lc_gff3, "LC")
features = pd.concat([hc_df, lc_df], ignore_index=True)

print(f"Parsed HC features: {hc_df.shape[0]:,}")
print(f"Parsed LC features: {lc_df.shape[0]:,}")
print(f"Combined features:  {features.shape[0]:,}")

features.to_csv(features_out, index=False)
print(f"Saved parsed feature table to: {features_out}")

# -----------------------------
# 5. Extract genes
# -----------------------------
genes = features[features["FeatureType"] == "gene"].copy()

genes = genes.rename(columns={
    "ID": "GeneModel",
    "SeqID": "Gene_Chromosome",
    "Start": "Gene_start",
    "End": "Gene_end",
    "Strand": "Gene_strand",
    "Name": "Gene_Name",
    "biotype": "Gene_biotype",
})

gene_cols = [
    "GeneModel", "Gene_Chromosome", "Gene_start", "Gene_end", "Gene_strand",
    "Annotation_confidence", "Gene_Name", "Gene_biotype"
]
genes = genes[gene_cols].drop_duplicates()

print(f"Gene rows: {genes.shape[0]:,}")

# -----------------------------
# 6. Extract transcripts
# -----------------------------
transcripts = features[features["FeatureType"] == "transcript"].copy()

transcripts = transcripts.rename(columns={
    "ID": "TranscriptID",
    "SeqID": "Transcript_Chromosome",
    "Start": "Transcript_start",
    "End": "Transcript_end",
    "Strand": "Transcript_strand",
    "Parent": "GeneModel",
    "Name": "Transcript_Name",
    "biotype": "Transcript_biotype",
})

tx_cols = [
    "TranscriptID", "GeneModel", "Transcript_Chromosome",
    "Transcript_start", "Transcript_end", "Transcript_strand",
    "Annotation_confidence", "Transcript_Name", "Transcript_biotype"
]
transcripts = transcripts[tx_cols].drop_duplicates()

print(f"Transcript rows: {transcripts.shape[0]:,}")

# -----------------------------
# 7. Extract CDS features and summarise by transcript
# -----------------------------
cds = features[features["FeatureType"] == "CDS"].copy()

# Parent of CDS is usually transcript ID
cds["TranscriptID"] = cds["Parent"].apply(
    lambda x: split_multi(x)[0] if pd.notna(x) and len(split_multi(x)) > 0 else np.nan
)

# Ensure compatible dtype before string assignment
cds["ProteinID"] = cds["protein_id"].astype("object")

missing_prot = cds["ProteinID"].isna() | (cds["ProteinID"].astype(str).str.strip() == "")
cds.loc[missing_prot, "ProteinID"] = cds.loc[missing_prot, "TranscriptID"].astype("object")

cds["GeneModel_from_protein"] = cds["ProteinID"].apply(gene_model_from_protein)
cds["CDS_nt_len"] = cds["End"] - cds["Start"] + 1

cds_summary = (
    cds
    .dropna(subset=["TranscriptID"])
    .groupby(["TranscriptID", "ProteinID", "GeneModel_from_protein", "Annotation_confidence"], dropna=False)
    .agg(
        CDS_feature_count=("FeatureType", "size"),
        CDS_start=("Start", "min"),
        CDS_end=("End", "max"),
        CDS_total_nt_length=("CDS_nt_len", "sum"),
        CDS_Chromosome=("SeqID", "first"),
        CDS_strand=("Strand", "first"),
        CDS_phase_values=("Phase", lambda s: "; ".join(sorted({str(x) for x in s if pd.notna(x)}))),
    )
    .reset_index()
)

print(f"CDS summary rows: {cds_summary.shape[0]:,}")

# -----------------------------
# 8. Merge transcript + gene + CDS information
# -----------------------------
mapping = cds_summary.merge(
    transcripts,
    on=["TranscriptID", "Annotation_confidence"],
    how="left"
)

# Prefer transcript-linked GeneModel, fallback to protein-derived GeneModel
mapping["GeneModel"] = np.where(
    mapping["GeneModel"].notna() & (mapping["GeneModel"].astype(str).str.strip() != ""),
    mapping["GeneModel"],
    mapping["GeneModel_from_protein"]
)

mapping = mapping.merge(
    genes,
    on=["GeneModel", "Annotation_confidence"],
    how="left"
)

# Build final chromosome/strand columns explicitly
mapping["Chromosome"] = np.where(
    mapping["Gene_Chromosome"].notna(),
    mapping["Gene_Chromosome"],
    np.where(
        mapping["Transcript_Chromosome"].notna(),
        mapping["Transcript_Chromosome"],
        mapping["CDS_Chromosome"]
    )
)

mapping["Strand"] = np.where(
    mapping["Gene_strand"].notna(),
    mapping["Gene_strand"],
    np.where(
        mapping["Transcript_strand"].notna(),
        mapping["Transcript_strand"],
        mapping["CDS_strand"]
    )
)

mapping["Protein_length_aa_from_CDS"] = np.where(
    pd.to_numeric(mapping["CDS_total_nt_length"], errors="coerce").notna(),
    (pd.to_numeric(mapping["CDS_total_nt_length"], errors="coerce") / 3).round(2),
    np.nan
)

final_cols = [
    "ProteinID",
    "GeneModel",
    "TranscriptID",
    "Chromosome",
    "Strand",
    "Gene_start",
    "Gene_end",
    "Transcript_start",
    "Transcript_end",
    "CDS_start",
    "CDS_end",
    "CDS_feature_count",
    "CDS_total_nt_length",
    "Protein_length_aa_from_CDS",
    "CDS_phase_values",
    "Annotation_confidence",
    "Gene_Name",
    "Gene_biotype",
    "Transcript_Name",
    "Transcript_biotype",
]
mapping = mapping[final_cols].drop_duplicates()

# -----------------------------
# 9. Add mapping status flags
# -----------------------------
mapping["ProteinID_found_in_GFF3"] = mapping["ProteinID"].notna()
mapping["GeneModel_found_in_GFF3"] = mapping["GeneModel"].notna()
mapping["Has_gene_coordinates"] = mapping["Gene_start"].notna() & mapping["Gene_end"].notna()
mapping["Has_transcript_coordinates"] = mapping["Transcript_start"].notna() & mapping["Transcript_end"].notna()

sort_cols = [c for c in ["Annotation_confidence", "Chromosome", "Gene_start", "ProteinID"] if c in mapping.columns]
mapping = mapping.sort_values(sort_cols).reset_index(drop=True)

mapping.to_csv(mapping_out, index=False)
print(f"Saved protein-to-gene mapping table to: {mapping_out}")

# -----------------------------
# 10. Summary tables
# -----------------------------
summary_rows = [
    {"Metric": "Parsed_HC_features", "Value": int(hc_df.shape[0])},
    {"Metric": "Parsed_LC_features", "Value": int(lc_df.shape[0])},
    {"Metric": "Combined_features", "Value": int(features.shape[0])},
    {"Metric": "Unique_genes", "Value": int(genes["GeneModel"].nunique())},
    {"Metric": "Unique_transcripts", "Value": int(transcripts["TranscriptID"].nunique())},
    {"Metric": "Unique_proteins_in_mapping", "Value": int(mapping["ProteinID"].nunique())},
    {"Metric": "Unique_gene_models_in_mapping", "Value": int(mapping["GeneModel"].nunique())},
    {"Metric": "Rows_with_gene_coordinates", "Value": int(mapping["Has_gene_coordinates"].sum())},
    {"Metric": "Rows_with_transcript_coordinates", "Value": int(mapping["Has_transcript_coordinates"].sum())},
]

if "Annotation_confidence" in mapping.columns:
    conf_counts = mapping["Annotation_confidence"].value_counts(dropna=False).to_dict()
    for k, v in conf_counts.items():
        summary_rows.append({"Metric": f"Mapping_rows_{k}", "Value": int(v)})

summary_df = pd.DataFrame(summary_rows)
summary_df.to_csv(summary_out, index=False)

# -----------------------------
# 11. Print summary
# -----------------------------
print("\n===== GFF3 PARSING SUMMARY =====")
display(summary_df)

print("\nExample protein-to-gene mappings:")
display(mapping.head(10))

print("\nExample parsed gene features:")
display(genes.head(5))

print("\nExample parsed transcript features:")
display(transcripts.head(5))

print("\nExample CDS summary:")
display(cds_summary.head(5))
```

    Parsed HC features: 1,874,236
    Parsed LC features: 967,070
    Combined features:  2,841,306
    Saved parsed feature table to: python_outputs/tables/wheat_gff3_parsed_features_HC_LC.csv
    Gene rows: 266,760
    Transcript rows: 295,922
    CDS summary rows: 295,914
    Saved protein-to-gene mapping table to: python_outputs/tables/wheat_protein_gene_mapping_HC_LC.csv
    
    ===== GFF3 PARSING SUMMARY =====
    


<div>
<style scoped>
    .dataframe tbody tr th:only-of-type {
        vertical-align: middle;
    }

    .dataframe tbody tr th {
        vertical-align: top;
    }

    .dataframe thead th {
        text-align: right;
    }
</style>
<table border="1" class="dataframe">
  <thead>
    <tr style="text-align: right;">
      <th></th>
      <th>Metric</th>
      <th>Value</th>
    </tr>
  </thead>
  <tbody>
    <tr>
      <th>0</th>
      <td>Parsed_HC_features</td>
      <td>1874236</td>
    </tr>
    <tr>
      <th>1</th>
      <td>Parsed_LC_features</td>
      <td>967070</td>
    </tr>
    <tr>
      <th>2</th>
      <td>Combined_features</td>
      <td>2841306</td>
    </tr>
    <tr>
      <th>3</th>
      <td>Unique_genes</td>
      <td>266760</td>
    </tr>
    <tr>
      <th>4</th>
      <td>Unique_transcripts</td>
      <td>295922</td>
    </tr>
    <tr>
      <th>5</th>
      <td>Unique_proteins_in_mapping</td>
      <td>295914</td>
    </tr>
    <tr>
      <th>6</th>
      <td>Unique_gene_models_in_mapping</td>
      <td>266752</td>
    </tr>
    <tr>
      <th>7</th>
      <td>Rows_with_gene_coordinates</td>
      <td>295914</td>
    </tr>
    <tr>
      <th>8</th>
      <td>Rows_with_transcript_coordinates</td>
      <td>295914</td>
    </tr>
    <tr>
      <th>9</th>
      <td>Mapping_rows_LC</td>
      <td>163290</td>
    </tr>
    <tr>
      <th>10</th>
      <td>Mapping_rows_HC</td>
      <td>132624</td>
    </tr>
  </tbody>
</table>
</div>


    
    Example protein-to-gene mappings:
    


<div>
<style scoped>
    .dataframe tbody tr th:only-of-type {
        vertical-align: middle;
    }

    .dataframe tbody tr th {
        vertical-align: top;
    }

    .dataframe thead th {
        text-align: right;
    }
</style>
<table border="1" class="dataframe">
  <thead>
    <tr style="text-align: right;">
      <th></th>
      <th>ProteinID</th>
      <th>GeneModel</th>
      <th>TranscriptID</th>
      <th>Chromosome</th>
      <th>Strand</th>
      <th>Gene_start</th>
      <th>Gene_end</th>
      <th>Transcript_start</th>
      <th>Transcript_end</th>
      <th>CDS_start</th>
      <th>...</th>
      <th>CDS_phase_values</th>
      <th>Annotation_confidence</th>
      <th>Gene_Name</th>
      <th>Gene_biotype</th>
      <th>Transcript_Name</th>
      <th>Transcript_biotype</th>
      <th>ProteinID_found_in_GFF3</th>
      <th>GeneModel_found_in_GFF3</th>
      <th>Has_gene_coordinates</th>
      <th>Has_transcript_coordinates</th>
    </tr>
  </thead>
  <tbody>
    <tr>
      <th>0</th>
      <td>TraesCS1A03G0000200.1</td>
      <td>TraesCS1A03G0000200</td>
      <td>TraesCS1A03G0000200.1</td>
      <td>Chr1A</td>
      <td>-</td>
      <td>40098</td>
      <td>70338</td>
      <td>40098</td>
      <td>70338</td>
      <td>58508</td>
      <td>...</td>
      <td>0</td>
      <td>HC</td>
      <td>TraesCS1A03G0000200</td>
      <td>NaN</td>
      <td>TraesCS1A03G0000200.1</td>
      <td>NaN</td>
      <td>True</td>
      <td>True</td>
      <td>True</td>
      <td>True</td>
    </tr>
    <tr>
      <th>1</th>
      <td>TraesCS1A03G0000400.1</td>
      <td>TraesCS1A03G0000400</td>
      <td>TraesCS1A03G0000400.1</td>
      <td>Chr1A</td>
      <td>+</td>
      <td>70239</td>
      <td>89245</td>
      <td>70239</td>
      <td>89245</td>
      <td>70239</td>
      <td>...</td>
      <td>0</td>
      <td>HC</td>
      <td>TraesCS1A03G0000400</td>
      <td>NaN</td>
      <td>TraesCS1A03G0000400.1</td>
      <td>NaN</td>
      <td>True</td>
      <td>True</td>
      <td>True</td>
      <td>True</td>
    </tr>
    <tr>
      <th>2</th>
      <td>TraesCS1A03G0000600.1</td>
      <td>TraesCS1A03G0000600</td>
      <td>TraesCS1A03G0000600.1</td>
      <td>Chr1A</td>
      <td>+</td>
      <td>95906</td>
      <td>104903</td>
      <td>95906</td>
      <td>104903</td>
      <td>104607</td>
      <td>...</td>
      <td>0</td>
      <td>HC</td>
      <td>TraesCS1A03G0000600</td>
      <td>NaN</td>
      <td>TraesCS1A03G0000600.1</td>
      <td>NaN</td>
      <td>True</td>
      <td>True</td>
      <td>True</td>
      <td>True</td>
    </tr>
    <tr>
      <th>3</th>
      <td>TraesCS1A03G0000800.1</td>
      <td>TraesCS1A03G0000800</td>
      <td>TraesCS1A03G0000800.1</td>
      <td>Chr1A</td>
      <td>+</td>
      <td>102794</td>
      <td>122504</td>
      <td>102794</td>
      <td>122504</td>
      <td>121263</td>
      <td>...</td>
      <td>0</td>
      <td>HC</td>
      <td>TraesCS1A03G0000800</td>
      <td>NaN</td>
      <td>TraesCS1A03G0000800.1</td>
      <td>NaN</td>
      <td>True</td>
      <td>True</td>
      <td>True</td>
      <td>True</td>
    </tr>
    <tr>
      <th>4</th>
      <td>TraesCS1A03G0001000.1</td>
      <td>TraesCS1A03G0001000</td>
      <td>TraesCS1A03G0001000.1</td>
      <td>Chr1A</td>
      <td>-</td>
      <td>149490</td>
      <td>154559</td>
      <td>149490</td>
      <td>154559</td>
      <td>149490</td>
      <td>...</td>
      <td>0; 2</td>
      <td>HC</td>
      <td>TraesCS1A03G0001000</td>
      <td>NaN</td>
      <td>TraesCS1A03G0001000.1</td>
      <td>NaN</td>
      <td>True</td>
      <td>True</td>
      <td>True</td>
      <td>True</td>
    </tr>
    <tr>
      <th>5</th>
      <td>TraesCS1A03G0001200.1</td>
      <td>TraesCS1A03G0001200</td>
      <td>TraesCS1A03G0001200.1</td>
      <td>Chr1A</td>
      <td>+</td>
      <td>162313</td>
      <td>162609</td>
      <td>162313</td>
      <td>162609</td>
      <td>162313</td>
      <td>...</td>
      <td>0</td>
      <td>HC</td>
      <td>TraesCS1A03G0001200</td>
      <td>NaN</td>
      <td>TraesCS1A03G0001200.1</td>
      <td>NaN</td>
      <td>True</td>
      <td>True</td>
      <td>True</td>
      <td>True</td>
    </tr>
    <tr>
      <th>6</th>
      <td>TraesCS1A03G0001300.1</td>
      <td>TraesCS1A03G0001300</td>
      <td>TraesCS1A03G0001300.1</td>
      <td>Chr1A</td>
      <td>-</td>
      <td>169592</td>
      <td>169969</td>
      <td>169592</td>
      <td>169969</td>
      <td>169592</td>
      <td>...</td>
      <td>0; 2</td>
      <td>HC</td>
      <td>TraesCS1A03G0001300</td>
      <td>NaN</td>
      <td>TraesCS1A03G0001300.1</td>
      <td>NaN</td>
      <td>True</td>
      <td>True</td>
      <td>True</td>
      <td>True</td>
    </tr>
    <tr>
      <th>7</th>
      <td>TraesCS1A03G0001400.1</td>
      <td>TraesCS1A03G0001400</td>
      <td>TraesCS1A03G0001400.1</td>
      <td>Chr1A</td>
      <td>-</td>
      <td>180175</td>
      <td>180552</td>
      <td>180175</td>
      <td>180552</td>
      <td>180175</td>
      <td>...</td>
      <td>0</td>
      <td>HC</td>
      <td>TraesCS1A03G0001400</td>
      <td>NaN</td>
      <td>TraesCS1A03G0001400.1</td>
      <td>NaN</td>
      <td>True</td>
      <td>True</td>
      <td>True</td>
      <td>True</td>
    </tr>
    <tr>
      <th>8</th>
      <td>TraesCS1A03G0001500.1</td>
      <td>TraesCS1A03G0001500</td>
      <td>TraesCS1A03G0001500.1</td>
      <td>Chr1A</td>
      <td>+</td>
      <td>233818</td>
      <td>245051</td>
      <td>233818</td>
      <td>245051</td>
      <td>243614</td>
      <td>...</td>
      <td>0</td>
      <td>HC</td>
      <td>TraesCS1A03G0001500</td>
      <td>NaN</td>
      <td>TraesCS1A03G0001500.1</td>
      <td>NaN</td>
      <td>True</td>
      <td>True</td>
      <td>True</td>
      <td>True</td>
    </tr>
    <tr>
      <th>9</th>
      <td>TraesCS1A03G0001700.1</td>
      <td>TraesCS1A03G0001700</td>
      <td>TraesCS1A03G0001700.1</td>
      <td>Chr1A</td>
      <td>-</td>
      <td>269472</td>
      <td>289602</td>
      <td>269472</td>
      <td>289602</td>
      <td>278951</td>
      <td>...</td>
      <td>0</td>
      <td>HC</td>
      <td>TraesCS1A03G0001700</td>
      <td>NaN</td>
      <td>TraesCS1A03G0001700.1</td>
      <td>NaN</td>
      <td>True</td>
      <td>True</td>
      <td>True</td>
      <td>True</td>
    </tr>
  </tbody>
</table>
<p>10 rows × 24 columns</p>
</div>


    
    Example parsed gene features:
    


<div>
<style scoped>
    .dataframe tbody tr th:only-of-type {
        vertical-align: middle;
    }

    .dataframe tbody tr th {
        vertical-align: top;
    }

    .dataframe thead th {
        text-align: right;
    }
</style>
<table border="1" class="dataframe">
  <thead>
    <tr style="text-align: right;">
      <th></th>
      <th>GeneModel</th>
      <th>Gene_Chromosome</th>
      <th>Gene_start</th>
      <th>Gene_end</th>
      <th>Gene_strand</th>
      <th>Annotation_confidence</th>
      <th>Gene_Name</th>
      <th>Gene_biotype</th>
    </tr>
  </thead>
  <tbody>
    <tr>
      <th>0</th>
      <td>TraesCS1A03G0000200</td>
      <td>Chr1A</td>
      <td>40098</td>
      <td>70338</td>
      <td>-</td>
      <td>HC</td>
      <td>TraesCS1A03G0000200</td>
      <td>NaN</td>
    </tr>
    <tr>
      <th>10</th>
      <td>TraesCS1A03G0000400</td>
      <td>Chr1A</td>
      <td>70239</td>
      <td>89245</td>
      <td>+</td>
      <td>HC</td>
      <td>TraesCS1A03G0000400</td>
      <td>NaN</td>
    </tr>
    <tr>
      <th>17</th>
      <td>TraesCS1A03G0000600</td>
      <td>Chr1A</td>
      <td>95906</td>
      <td>104903</td>
      <td>+</td>
      <td>HC</td>
      <td>TraesCS1A03G0000600</td>
      <td>NaN</td>
    </tr>
    <tr>
      <th>24</th>
      <td>TraesCS1A03G0000800</td>
      <td>Chr1A</td>
      <td>102794</td>
      <td>122504</td>
      <td>+</td>
      <td>HC</td>
      <td>TraesCS1A03G0000800</td>
      <td>NaN</td>
    </tr>
    <tr>
      <th>32</th>
      <td>TraesCS1A03G0001000</td>
      <td>Chr1A</td>
      <td>149490</td>
      <td>154559</td>
      <td>-</td>
      <td>HC</td>
      <td>TraesCS1A03G0001000</td>
      <td>NaN</td>
    </tr>
  </tbody>
</table>
</div>


    
    Example parsed transcript features:
    


<div>
<style scoped>
    .dataframe tbody tr th:only-of-type {
        vertical-align: middle;
    }

    .dataframe tbody tr th {
        vertical-align: top;
    }

    .dataframe thead th {
        text-align: right;
    }
</style>
<table border="1" class="dataframe">
  <thead>
    <tr style="text-align: right;">
      <th></th>
      <th>TranscriptID</th>
      <th>GeneModel</th>
      <th>Transcript_Chromosome</th>
      <th>Transcript_start</th>
      <th>Transcript_end</th>
      <th>Transcript_strand</th>
      <th>Annotation_confidence</th>
      <th>Transcript_Name</th>
      <th>Transcript_biotype</th>
    </tr>
  </thead>
  <tbody>
    <tr>
      <th>1</th>
      <td>TraesCS1A03G0000200.1</td>
      <td>TraesCS1A03G0000200</td>
      <td>Chr1A</td>
      <td>40098</td>
      <td>70338</td>
      <td>-</td>
      <td>HC</td>
      <td>TraesCS1A03G0000200.1</td>
      <td>NaN</td>
    </tr>
    <tr>
      <th>11</th>
      <td>TraesCS1A03G0000400.1</td>
      <td>TraesCS1A03G0000400</td>
      <td>Chr1A</td>
      <td>70239</td>
      <td>89245</td>
      <td>+</td>
      <td>HC</td>
      <td>TraesCS1A03G0000400.1</td>
      <td>NaN</td>
    </tr>
    <tr>
      <th>18</th>
      <td>TraesCS1A03G0000600.1</td>
      <td>TraesCS1A03G0000600</td>
      <td>Chr1A</td>
      <td>95906</td>
      <td>104903</td>
      <td>+</td>
      <td>HC</td>
      <td>TraesCS1A03G0000600.1</td>
      <td>NaN</td>
    </tr>
    <tr>
      <th>25</th>
      <td>TraesCS1A03G0000800.1</td>
      <td>TraesCS1A03G0000800</td>
      <td>Chr1A</td>
      <td>102794</td>
      <td>122504</td>
      <td>+</td>
      <td>HC</td>
      <td>TraesCS1A03G0000800.1</td>
      <td>NaN</td>
    </tr>
    <tr>
      <th>33</th>
      <td>TraesCS1A03G0001000.1</td>
      <td>TraesCS1A03G0001000</td>
      <td>Chr1A</td>
      <td>149490</td>
      <td>154559</td>
      <td>-</td>
      <td>HC</td>
      <td>TraesCS1A03G0001000.1</td>
      <td>NaN</td>
    </tr>
  </tbody>
</table>
</div>


    
    Example CDS summary:
    


<div>
<style scoped>
    .dataframe tbody tr th:only-of-type {
        vertical-align: middle;
    }

    .dataframe tbody tr th {
        vertical-align: top;
    }

    .dataframe thead th {
        text-align: right;
    }
</style>
<table border="1" class="dataframe">
  <thead>
    <tr style="text-align: right;">
      <th></th>
      <th>TranscriptID</th>
      <th>ProteinID</th>
      <th>GeneModel_from_protein</th>
      <th>Annotation_confidence</th>
      <th>CDS_feature_count</th>
      <th>CDS_start</th>
      <th>CDS_end</th>
      <th>CDS_total_nt_length</th>
      <th>CDS_Chromosome</th>
      <th>CDS_strand</th>
      <th>CDS_phase_values</th>
    </tr>
  </thead>
  <tbody>
    <tr>
      <th>0</th>
      <td>TraesCS1A03G0000100LC.1</td>
      <td>TraesCS1A03G0000100LC.1</td>
      <td>TraesCS1A03G0000100LC</td>
      <td>LC</td>
      <td>1</td>
      <td>41202</td>
      <td>41522</td>
      <td>321</td>
      <td>Chr1A</td>
      <td>+</td>
      <td>0</td>
    </tr>
    <tr>
      <th>1</th>
      <td>TraesCS1A03G0000200.1</td>
      <td>TraesCS1A03G0000200.1</td>
      <td>TraesCS1A03G0000200</td>
      <td>HC</td>
      <td>1</td>
      <td>58508</td>
      <td>58768</td>
      <td>261</td>
      <td>Chr1A</td>
      <td>-</td>
      <td>0</td>
    </tr>
    <tr>
      <th>2</th>
      <td>TraesCS1A03G0000300LC.1</td>
      <td>TraesCS1A03G0000300LC.1</td>
      <td>TraesCS1A03G0000300LC</td>
      <td>LC</td>
      <td>1</td>
      <td>42451</td>
      <td>42645</td>
      <td>195</td>
      <td>Chr1A</td>
      <td>-</td>
      <td>0</td>
    </tr>
    <tr>
      <th>3</th>
      <td>TraesCS1A03G0000400.1</td>
      <td>TraesCS1A03G0000400.1</td>
      <td>TraesCS1A03G0000400</td>
      <td>HC</td>
      <td>1</td>
      <td>70239</td>
      <td>70556</td>
      <td>318</td>
      <td>Chr1A</td>
      <td>+</td>
      <td>0</td>
    </tr>
    <tr>
      <th>4</th>
      <td>TraesCS1A03G0000500LC.1</td>
      <td>TraesCS1A03G0000500LC.1</td>
      <td>TraesCS1A03G0000500LC</td>
      <td>LC</td>
      <td>3</td>
      <td>94185</td>
      <td>110830</td>
      <td>309</td>
      <td>Chr1A</td>
      <td>+</td>
      <td>0; 1</td>
    </tr>
  </tbody>
</table>
</div>


# Step 6 — FragPipe Output Annotation and Quality Summary

FragPipe peptide and protein identification tables were standardised and annotated in Python to improve traceability, facilitate downstream batch processing, and enable dataset-wide quality assessment across all wheat tissues.

Annotated FragPipe output tables were generated for both peptide- and protein-level results by importing the original `.tsv` files and appending standardised metadata columns derived from the experimental manifest file.

---

## Input files

### FragPipe result tables

```text
FragPipe_FirstAuthor_Source_Tissue-Raw-Code_peptide.tsv
FragPipe_FirstAuthor_Source_Tissue-Raw-Code_protein.tsv
```

### Example

```text
FragPipe_Duncan_PXD004720_anther_peptide.tsv
FragPipe_Duncan_PXD004720_anther_protein.tsv
```

### Manifest file

```text
wheat_tissues_FragPipe-result-manifest_2026-05-11.csv
```

---

## Metadata annotation

The following traceability columns were added to each FragPipe result table:

| Column | Description |
|---|---|
| Index | Sequential row identifier |
| Source | Proteomics repository/project identifier |
| Species | Species name |
| Tissue | Wheat tissue name/code |
| Batch | FragPipe processing batch |
| FragPipe_result | Result type (`peptide` or `protein`) |
| Contaminant | Contaminant assignment (`yes` or `no`) |

---

## Contaminant assignment

Contaminant status was inferred directly from the FragPipe `Protein` column.

### Rule

```text
Protein accession starts with "Traes" → Contaminant = no
Otherwise → Contaminant = yes
```

This strategy enabled rapid separation of wheat-derived peptide/protein identifications from common contaminant proteins originating from keratins, trypsin, laboratory handling, or external contaminant databases.

---

## Output files

Annotated tables were exported as `.csv` files into:

```text
python_outputs/tables/
```

### Example outputs

```text
FragPipe_Duncan_PXD004720_anther_peptide_annotated.csv
FragPipe_Duncan_PXD004720_anther_protein_annotated.csv
```

---

## Batch processing

All wheat tissues were processed automatically using generic Python functions driven by the experimental manifest file.

### Total processed outputs

```text
32 tissues × 2 result types = 64 annotated tables
```

---

## Quality summary table

A global summary table was additionally generated to capture key identification and contaminant statistics for each annotated FragPipe output file.

### Summary metrics

| Metric | Description |
|---|---|
| peptide_count | Number of peptide rows |
| protein_count | Number of protein rows |
| Contaminant_count | Number of contaminant entries |
| Non_contaminant_count | Number of wheat-derived entries |

### Output summary file

```text
wheat_fragpipe_annotation_summary_step6.csv
```

This summary table served as the foundation for subsequent filtering, peptide-to-protein mapping, genome projection, and downstream proteogenomics analyses.


```python
# ============================================================
# Step 6 — Annotate FragPipe peptide/protein output tables
# ============================================================

import pandas as pd
from pathlib import Path

# -----------------------------
# 1. Input / output paths
# -----------------------------
fragpipe_dir = Path("FragPipe_results")
manifest_file = fragpipe_dir / "wheat_tissues_FragPipe-result-manifest_2026-05-11.csv"

output_dir = Path("python_outputs/tables")
output_dir.mkdir(parents=True, exist_ok=True)

# -----------------------------
# 2. Load manifest
# -----------------------------
manifest = pd.read_csv(manifest_file, encoding="utf-8-sig")

display(manifest.head())


# -----------------------------
# 3. Helper function
# -----------------------------
def annotate_fragpipe_output(source, tissue_raw_code, result_type):
    """
    Annotate one FragPipe peptide or protein output table with traceability columns.

    Parameters
    ----------
    source : str
        Proteomics dataset source, e.g. "PXD004720", "PXD050500", "MSV000090572".

    tissue_raw_code : str
        Tissue raw code as listed in the manifest, e.g. "anther", "node", "stored-grain".

    result_type : str
        Either "peptide" or "protein".

    Returns
    -------
    pd.DataFrame
        Annotated FragPipe table.
    """

    result_type = result_type.lower()

    if result_type not in ["peptide", "protein"]:
        raise ValueError("result_type must be either 'peptide' or 'protein'.")

    # Find matching manifest row
    match = manifest[
        (manifest["Source"] == source) &
        (manifest["Tissue-Raw-Code"] == tissue_raw_code)
    ]

    if match.empty:
        raise ValueError(
            f"No manifest entry found for Source='{source}' and "
            f"Tissue-Raw-Code='{tissue_raw_code}'."
        )

    if len(match) > 1:
        raise ValueError(
            f"Multiple manifest entries found for Source='{source}' and "
            f"Tissue-Raw-Code='{tissue_raw_code}'."
        )

    row = match.iloc[0]

    # Retrieve input filename from manifest
    if result_type == "peptide":
        input_filename = row["FragPipe-Output-Peptide"]
        anchor_col = "Peptide"
    else:
        input_filename = row["FragPipe-Output-Protein"]
        anchor_col = "Protein"

    input_path = fragpipe_dir / input_filename

    if not input_path.exists():
        raise FileNotFoundError(f"Input file not found: {input_path}")

    # Read FragPipe TSV file
    data = pd.read_csv(input_path, sep="\t", header=0)

    # Check required Protein column for contaminant assignment
    if "Protein" not in data.columns:
        raise KeyError(f"'Protein' column not found in {input_filename}")

    # Add traceability columns
    data["Index"] = range(1, len(data) + 1)
    data["Source"] = row["Source"]
    data["Species"] = row["Species"]
    data["Tissue"] = row["Tissue-Raw-Code"]
    data["Batch"] = row["Batch"]
    data["FragPipe_result"] = result_type
    data["Contaminant"] = data["Protein"].astype(str).str.startswith("Traes").map(
        {True: "no", False: "yes"}
    )

    new_cols = [
        "Index",
        "Source",
        "Species",
        "Tissue",
        "Batch",
        "FragPipe_result",
        "Contaminant"
    ]

    # Insert metadata columns before Peptide or Protein column when possible
    remaining_cols = [col for col in data.columns if col not in new_cols]

    if anchor_col in remaining_cols:
        anchor_pos = remaining_cols.index(anchor_col)
        ordered_cols = (
            remaining_cols[:anchor_pos] +
            new_cols +
            remaining_cols[anchor_pos:]
        )
    else:
        ordered_cols = new_cols + remaining_cols

    data = data[ordered_cols]

    # Save annotated table
    output_filename = input_filename.replace(".tsv", "_annotated.csv")
    output_path = output_dir / output_filename

    data.to_csv(output_path, index=False)

    print(f"Annotated file saved: {output_path}")
    print(f"Rows: {data.shape[0]:,}")
    print(f"Columns: {data.shape[1]:,}")

    return data


# -----------------------------
# 4. Example usage to check code
# -----------------------------

# Example peptide file
annotated_peptide = annotate_fragpipe_output(
    source="PXD050500",
    tissue_raw_code="radicle",
    result_type="peptide"
)

# Example protein file
annotated_protein = annotate_fragpipe_output(
    source="PXD050500",
    tissue_raw_code="radicle",
    result_type="protein"
)

# -----------------------------
# 5. Annotate all FragPipe TSV files
# -----------------------------

for _, row in manifest.iterrows():
    source = row["Source"]
    tissue_raw_code = row["Tissue-Raw-Code"]

    print(f"\nProcessing {source} | {tissue_raw_code}")

    annotate_fragpipe_output(
        source=source,
        tissue_raw_code=tissue_raw_code,
        result_type="peptide"
    )

    annotate_fragpipe_output(
        source=source,
        tissue_raw_code=tissue_raw_code,
        result_type="protein"
    )


# -----------------------------
# 6. Create Step 6 summary table
# -----------------------------

step6_summary_records = []

for _, row in manifest.iterrows():
    source = row["Source"]
    species = row["Species"]
    tissue_raw_code = row["Tissue-Raw-Code"]
    batch = row["Batch"]

    for result_type in ["peptide", "protein"]:

        if result_type == "peptide":
            input_filename = row["FragPipe-Output-Peptide"]
            count_label = "peptide_count"
        else:
            input_filename = row["FragPipe-Output-Protein"]
            count_label = "protein_count"

        annotated_filename = input_filename.replace(".tsv", "_annotated.csv")
        annotated_path = output_dir / annotated_filename

        if not annotated_path.exists():
            print(f"Warning: annotated file not found, skipped: {annotated_path}")
            continue

        data = pd.read_csv(annotated_path)

        total_count = len(data)
        contaminant_count = (data["Contaminant"] == "yes").sum()
        non_contaminant_count = (data["Contaminant"] == "no").sum()

        step6_summary_records.append({
            "Source": source,
            "Species": species,
            "Tissue": tissue_raw_code,
            "Batch": batch,
            "FragPipe_result": result_type,
            "Annotated_file": annotated_filename,
            count_label: total_count,
            "Contaminant_count": contaminant_count,
            "Non_contaminant_count": non_contaminant_count
        })

step6_summary = pd.DataFrame(step6_summary_records)

# Keep peptide/protein counts in one generic column as well
step6_summary["Total_rows"] = step6_summary[["peptide_count", "protein_count"]].sum(axis=1, skipna=True)

# Reorder columns
summary_cols = [
    "Source",
    "Species",
    "Tissue",
    "Batch",
    "FragPipe_result",
    "Annotated_file",
    "Total_rows",
    "peptide_count",
    "protein_count",
    "Contaminant_count",
    "Non_contaminant_count"
]

step6_summary = step6_summary[summary_cols]

# Save summary table
step6_summary_out = output_dir / "wheat_fragpipe_annotation_summary_step6.csv"
step6_summary.to_csv(step6_summary_out, index=False)

print(f"Step 6 summary saved: {step6_summary_out}")
print(f"Summary rows: {step6_summary.shape[0]:,}")

display(step6_summary)
```


<div>
<style scoped>
    .dataframe tbody tr th:only-of-type {
        vertical-align: middle;
    }

    .dataframe tbody tr th {
        vertical-align: top;
    }

    .dataframe thead th {
        text-align: right;
    }
</style>
<table border="1" class="dataframe">
  <thead>
    <tr style="text-align: right;">
      <th></th>
      <th>Order</th>
      <th>First-Author</th>
      <th>Source</th>
      <th>Species</th>
      <th>Tissue-Name</th>
      <th>Tissue-Raw-Code</th>
      <th>Tissue-Description</th>
      <th>Batch</th>
      <th>FragPipe-Output-Peptide</th>
      <th>FragPipe-Output-Protein</th>
    </tr>
  </thead>
  <tbody>
    <tr>
      <th>0</th>
      <td>1</td>
      <td>Vincent</td>
      <td>MSV000090572</td>
      <td>bread wheat</td>
      <td>stored_grain</td>
      <td>stored_grain</td>
      <td>NaN</td>
      <td>single</td>
      <td>FragPipe_Vincent_MSV000090572_stored-grain_pep...</td>
      <td>FragPipe_Vincent_MSV000090572_stored-grain_pro...</td>
    </tr>
    <tr>
      <th>1</th>
      <td>2</td>
      <td>Liu</td>
      <td>PXD050500</td>
      <td>bread wheat</td>
      <td>germ</td>
      <td>coleoptile</td>
      <td>The protective, white, tubular sheath that eme...</td>
      <td>single</td>
      <td>FragPipe_Liu_PXD050500_coleoptile_peptide.tsv</td>
      <td>FragPipe_Liu_PXD050500_coleoptile_protein.tsv</td>
    </tr>
    <tr>
      <th>2</th>
      <td>3</td>
      <td>Liu</td>
      <td>PXD050500</td>
      <td>bread wheat</td>
      <td>node</td>
      <td>node</td>
      <td>The crown area where the seminal roots, advent...</td>
      <td>single</td>
      <td>FragPipe_Liu_PXD050500_node_peptide.tsv</td>
      <td>FragPipe_Liu_PXD050500_node_protein.tsv</td>
    </tr>
    <tr>
      <th>3</th>
      <td>4</td>
      <td>Liu</td>
      <td>PXD050500</td>
      <td>bread wheat</td>
      <td>root</td>
      <td>radicle</td>
      <td>The first primary root that emerges from the s...</td>
      <td>single</td>
      <td>FragPipe_Liu_PXD050500_radicle_peptide.tsv</td>
      <td>FragPipe_Liu_PXD050500_radicle_protein.tsv</td>
    </tr>
    <tr>
      <th>4</th>
      <td>5</td>
      <td>Duncan</td>
      <td>PXD004720</td>
      <td>bread wheat</td>
      <td>anther</td>
      <td>anther</td>
      <td>NaN</td>
      <td>single</td>
      <td>FragPipe_Duncan_PXD004720_anther_peptide.tsv</td>
      <td>FragPipe_Duncan_PXD004720_anther_protein.tsv</td>
    </tr>
  </tbody>
</table>
</div>


    Annotated file saved: python_outputs\tables\FragPipe_Liu_PXD050500_radicle_peptide_annotated.csv
    Rows: 333,400
    Columns: 30
    Annotated file saved: python_outputs\tables\FragPipe_Liu_PXD050500_radicle_protein_annotated.csv
    Rows: 105,970
    Columns: 33
    
    Processing MSV000090572 | stored_grain
    Annotated file saved: python_outputs\tables\FragPipe_Vincent_MSV000090572_stored-grain_peptide_annotated.csv
    Rows: 9,343
    Columns: 30
    Annotated file saved: python_outputs\tables\FragPipe_Vincent_MSV000090572_stored-grain_protein_annotated.csv
    Rows: 7,074
    Columns: 33
    
    Processing PXD050500 | coleoptile
    Annotated file saved: python_outputs\tables\FragPipe_Liu_PXD050500_coleoptile_peptide_annotated.csv
    Rows: 583,184
    Columns: 30
    Annotated file saved: python_outputs\tables\FragPipe_Liu_PXD050500_coleoptile_protein_annotated.csv
    Rows: 137,841
    Columns: 33
    
    Processing PXD050500 | node
    Annotated file saved: python_outputs\tables\FragPipe_Liu_PXD050500_node_peptide_annotated.csv
    Rows: 596,437
    Columns: 30
    Annotated file saved: python_outputs\tables\FragPipe_Liu_PXD050500_node_protein_annotated.csv
    Rows: 141,772
    Columns: 33
    
    Processing PXD050500 | radicle
    Annotated file saved: python_outputs\tables\FragPipe_Liu_PXD050500_radicle_peptide_annotated.csv
    Rows: 333,400
    Columns: 30
    Annotated file saved: python_outputs\tables\FragPipe_Liu_PXD050500_radicle_protein_annotated.csv
    Rows: 105,970
    Columns: 33
    
    Processing PXD004720 | anther
    Annotated file saved: python_outputs\tables\FragPipe_Duncan_PXD004720_anther_peptide_annotated.csv
    Rows: 34,203
    Columns: 30
    Annotated file saved: python_outputs\tables\FragPipe_Duncan_PXD004720_anther_protein_annotated.csv
    Rows: 10,095
    Columns: 33
    
    Processing PXD004720 | boot
    Annotated file saved: python_outputs\tables\FragPipe_Duncan_PXD004720_boot_peptide_annotated.csv
    Rows: 3,644
    Columns: 30
    Annotated file saved: python_outputs\tables\FragPipe_Duncan_PXD004720_boot_protein_annotated.csv
    Rows: 2,828
    Columns: 33
    
    Processing PXD004720 | coleoptile
    Annotated file saved: python_outputs\tables\FragPipe_Duncan_PXD004720_coleoptile_peptide_annotated.csv
    Rows: 41,126
    Columns: 30
    Annotated file saved: python_outputs\tables\FragPipe_Duncan_PXD004720_coleoptile_protein_annotated.csv
    Rows: 12,216
    Columns: 33
    
    Processing PXD004720 | embryo
    Annotated file saved: python_outputs\tables\FragPipe_Duncan_PXD004720_embryo_peptide_annotated.csv
    Rows: 2,868
    Columns: 30
    Annotated file saved: python_outputs\tables\FragPipe_Duncan_PXD004720_embryo_protein_annotated.csv
    Rows: 2,403
    Columns: 33
    
    Processing PXD004720 | endosperm
    Annotated file saved: python_outputs\tables\FragPipe_Duncan_PXD004720_endosperm_peptide_annotated.csv
    Rows: 20,174
    Columns: 30
    Annotated file saved: python_outputs\tables\FragPipe_Duncan_PXD004720_endosperm_protein_annotated.csv
    Rows: 7,427
    Columns: 33
    
    Processing PXD004720 | glume
    Annotated file saved: python_outputs\tables\FragPipe_Duncan_PXD004720_glume_peptide_annotated.csv
    Rows: 28,683
    Columns: 30
    Annotated file saved: python_outputs\tables\FragPipe_Duncan_PXD004720_glume_protein_annotated.csv
    Rows: 9,612
    Columns: 33
    
    Processing PXD004720 | grain-zadoks-70
    Annotated file saved: python_outputs\tables\FragPipe_Duncan_PXD004720_grain-zadoks-70_peptide_annotated.csv
    Rows: 26,229
    Columns: 30
    Annotated file saved: python_outputs\tables\FragPipe_Duncan_PXD004720_grain-zadoks-70_protein_annotated.csv
    Rows: 9,821
    Columns: 33
    
    Processing PXD004720 | grain-zadoks-71
    Annotated file saved: python_outputs\tables\FragPipe_Duncan_PXD004720_grain-zadoks-71_peptide_annotated.csv
    Rows: 35,990
    Columns: 30
    Annotated file saved: python_outputs\tables\FragPipe_Duncan_PXD004720_grain-zadoks-71_protein_annotated.csv
    Rows: 12,336
    Columns: 33
    
    Processing PXD004720 | grain-zadoks-75
    Annotated file saved: python_outputs\tables\FragPipe_Duncan_PXD004720_grain-zadoks-75_peptide_annotated.csv
    Rows: 27,552
    Columns: 30
    Annotated file saved: python_outputs\tables\FragPipe_Duncan_PXD004720_grain-zadoks-75_protein_annotated.csv
    Rows: 11,100
    Columns: 33
    
    Processing PXD004720 | grain-zadoks-83
    Annotated file saved: python_outputs\tables\FragPipe_Duncan_PXD004720_grain-zadoks-83_peptide_annotated.csv
    Rows: 24,051
    Columns: 30
    Annotated file saved: python_outputs\tables\FragPipe_Duncan_PXD004720_grain-zadoks-83_protein_annotated.csv
    Rows: 10,194
    Columns: 33
    
    Processing PXD004720 | grain-zadoks-87
    Annotated file saved: python_outputs\tables\FragPipe_Duncan_PXD004720_grain-zadoks-87_peptide_annotated.csv
    Rows: 24,674
    Columns: 30
    Annotated file saved: python_outputs\tables\FragPipe_Duncan_PXD004720_grain-zadoks-87_protein_annotated.csv
    Rows: 10,389
    Columns: 33
    
    Processing PXD004720 | leaf-flag-mature
    Annotated file saved: python_outputs\tables\FragPipe_Duncan_PXD004720_leaf-flag-mature_peptide_annotated.csv
    Rows: 29,941
    Columns: 30
    Annotated file saved: python_outputs\tables\FragPipe_Duncan_PXD004720_leaf-flag-mature_protein_annotated.csv
    Rows: 9,639
    Columns: 33
    
    Processing PXD004720 | leaf-flag-senescing
    Annotated file saved: python_outputs\tables\FragPipe_Duncan_PXD004720_leaf-flag-senescing_peptide_annotated.csv
    Rows: 11,394
    Columns: 30
    Annotated file saved: python_outputs\tables\FragPipe_Duncan_PXD004720_leaf-flag-senescing_protein_annotated.csv
    Rows: 8,034
    Columns: 33
    
    Processing PXD004720 | leaf-flag-young
    Annotated file saved: python_outputs\tables\FragPipe_Duncan_PXD004720_leaf-flag-young_peptide_annotated.csv
    Rows: 23,441
    Columns: 30
    Annotated file saved: python_outputs\tables\FragPipe_Duncan_PXD004720_leaf-flag-young_protein_annotated.csv
    Rows: 7,542
    Columns: 33
    
    Processing PXD004720 | lemma
    Annotated file saved: python_outputs\tables\FragPipe_Duncan_PXD004720_lemma_peptide_annotated.csv
    Rows: 29,842
    Columns: 30
    Annotated file saved: python_outputs\tables\FragPipe_Duncan_PXD004720_lemma_protein_annotated.csv
    Rows: 9,956
    Columns: 33
    
    Processing PXD004720 | node
    Annotated file saved: python_outputs\tables\FragPipe_Duncan_PXD004720_node_peptide_annotated.csv
    Rows: 21,508
    Columns: 30
    Annotated file saved: python_outputs\tables\FragPipe_Duncan_PXD004720_node_protein_annotated.csv
    Rows: 10,530
    Columns: 33
    
    Processing PXD004720 | node_secretion
    Annotated file saved: python_outputs\tables\FragPipe_Duncan_PXD004720_node-secretion_peptide_annotated.csv
    Rows: 34,194
    Columns: 30
    Annotated file saved: python_outputs\tables\FragPipe_Duncan_PXD004720_node-secretion_protein_annotated.csv
    Rows: 10,572
    Columns: 33
    
    Processing PXD004720 | palea
    Annotated file saved: python_outputs\tables\FragPipe_Duncan_PXD004720_palea_peptide_annotated.csv
    Rows: 21,774
    Columns: 30
    Annotated file saved: python_outputs\tables\FragPipe_Duncan_PXD004720_palea_protein_annotated.csv
    Rows: 6,942
    Columns: 33
    
    Processing PXD004720 | pericarp
    Annotated file saved: python_outputs\tables\FragPipe_Duncan_PXD004720_pericarp_peptide_annotated.csv
    Rows: 28,448
    Columns: 30
    Annotated file saved: python_outputs\tables\FragPipe_Duncan_PXD004720_pericarp_protein_annotated.csv
    Rows: 9,466
    Columns: 33
    
    Processing PXD004720 | pollen
    Annotated file saved: python_outputs\tables\FragPipe_Duncan_PXD004720_pollen_peptide_annotated.csv
    Rows: 13,593
    Columns: 30
    Annotated file saved: python_outputs\tables\FragPipe_Duncan_PXD004720_pollen_protein_annotated.csv
    Rows: 4,676
    Columns: 33
    
    Processing PXD004720 | rachilla
    Annotated file saved: python_outputs\tables\FragPipe_Duncan_PXD004720_rachilla_peptide_annotated.csv
    Rows: 31,213
    Columns: 30
    Annotated file saved: python_outputs\tables\FragPipe_Duncan_PXD004720_rachilla_protein_annotated.csv
    Rows: 9,717
    Columns: 33
    
    Processing PXD004720 | radicle
    Annotated file saved: python_outputs\tables\FragPipe_Duncan_PXD004720_radicle_peptide_annotated.csv
    Rows: 39,704
    Columns: 30
    Annotated file saved: python_outputs\tables\FragPipe_Duncan_PXD004720_radicle_protein_annotated.csv
    Rows: 11,752
    Columns: 33
    
    Processing PXD004720 | root-mature
    Annotated file saved: python_outputs\tables\FragPipe_Duncan_PXD004720_root-mature_peptide_annotated.csv
    Rows: 21,390
    Columns: 30
    Annotated file saved: python_outputs\tables\FragPipe_Duncan_PXD004720_root-mature_protein_annotated.csv
    Rows: 12,214
    Columns: 33
    
    Processing PXD004720 | root-secretion
    Annotated file saved: python_outputs\tables\FragPipe_Duncan_PXD004720_root-secretion_peptide_annotated.csv
    Rows: 21,329
    Columns: 30
    Annotated file saved: python_outputs\tables\FragPipe_Duncan_PXD004720_root-secretion_protein_annotated.csv
    Rows: 10,309
    Columns: 33
    
    Processing PXD004720 | root-tip
    Annotated file saved: python_outputs\tables\FragPipe_Duncan_PXD004720_root-tip_peptide_annotated.csv
    Rows: 38,681
    Columns: 30
    Annotated file saved: python_outputs\tables\FragPipe_Duncan_PXD004720_root-tip_protein_annotated.csv
    Rows: 10,344
    Columns: 33
    
    Processing PXD004720 | root-vasculature
    Annotated file saved: python_outputs\tables\FragPipe_Duncan_PXD004720_root-vasculature_peptide_annotated.csv
    Rows: 20,911
    Columns: 30
    Annotated file saved: python_outputs\tables\FragPipe_Duncan_PXD004720_root-vasculature_protein_annotated.csv
    Rows: 7,577
    Columns: 33
    
    Processing PXD004720 | spike-immature
    Annotated file saved: python_outputs\tables\FragPipe_Duncan_PXD004720_spike-immature_peptide_annotated.csv
    Rows: 36,414
    Columns: 30
    Annotated file saved: python_outputs\tables\FragPipe_Duncan_PXD004720_spike-immature_protein_annotated.csv
    Rows: 10,948
    Columns: 33
    
    Processing PXD004720 | stem
    Annotated file saved: python_outputs\tables\FragPipe_Duncan_PXD004720_stem_peptide_annotated.csv
    Rows: 14,389
    Columns: 30
    Annotated file saved: python_outputs\tables\FragPipe_Duncan_PXD004720_stem_protein_annotated.csv
    Rows: 8,984
    Columns: 33
    Step 6 summary saved: python_outputs\tables\wheat_fragpipe_annotation_summary_step6.csv
    Summary rows: 64
    


<div>
<style scoped>
    .dataframe tbody tr th:only-of-type {
        vertical-align: middle;
    }

    .dataframe tbody tr th {
        vertical-align: top;
    }

    .dataframe thead th {
        text-align: right;
    }
</style>
<table border="1" class="dataframe">
  <thead>
    <tr style="text-align: right;">
      <th></th>
      <th>Source</th>
      <th>Species</th>
      <th>Tissue</th>
      <th>Batch</th>
      <th>FragPipe_result</th>
      <th>Annotated_file</th>
      <th>Total_rows</th>
      <th>peptide_count</th>
      <th>protein_count</th>
      <th>Contaminant_count</th>
      <th>Non_contaminant_count</th>
    </tr>
  </thead>
  <tbody>
    <tr>
      <th>0</th>
      <td>MSV000090572</td>
      <td>bread wheat</td>
      <td>stored_grain</td>
      <td>single</td>
      <td>peptide</td>
      <td>FragPipe_Vincent_MSV000090572_stored-grain_pep...</td>
      <td>9343.0</td>
      <td>9343.0</td>
      <td>NaN</td>
      <td>14</td>
      <td>9329</td>
    </tr>
    <tr>
      <th>1</th>
      <td>MSV000090572</td>
      <td>bread wheat</td>
      <td>stored_grain</td>
      <td>single</td>
      <td>protein</td>
      <td>FragPipe_Vincent_MSV000090572_stored-grain_pro...</td>
      <td>7074.0</td>
      <td>NaN</td>
      <td>7074.0</td>
      <td>6</td>
      <td>7068</td>
    </tr>
    <tr>
      <th>2</th>
      <td>PXD050500</td>
      <td>bread wheat</td>
      <td>coleoptile</td>
      <td>single</td>
      <td>peptide</td>
      <td>FragPipe_Liu_PXD050500_coleoptile_peptide_anno...</td>
      <td>583184.0</td>
      <td>583184.0</td>
      <td>NaN</td>
      <td>409</td>
      <td>582775</td>
    </tr>
    <tr>
      <th>3</th>
      <td>PXD050500</td>
      <td>bread wheat</td>
      <td>coleoptile</td>
      <td>single</td>
      <td>protein</td>
      <td>FragPipe_Liu_PXD050500_coleoptile_protein_anno...</td>
      <td>137841.0</td>
      <td>NaN</td>
      <td>137841.0</td>
      <td>79</td>
      <td>137762</td>
    </tr>
    <tr>
      <th>4</th>
      <td>PXD050500</td>
      <td>bread wheat</td>
      <td>node</td>
      <td>single</td>
      <td>peptide</td>
      <td>FragPipe_Liu_PXD050500_node_peptide_annotated.csv</td>
      <td>596437.0</td>
      <td>596437.0</td>
      <td>NaN</td>
      <td>483</td>
      <td>595954</td>
    </tr>
    <tr>
      <th>...</th>
      <td>...</td>
      <td>...</td>
      <td>...</td>
      <td>...</td>
      <td>...</td>
      <td>...</td>
      <td>...</td>
      <td>...</td>
      <td>...</td>
      <td>...</td>
      <td>...</td>
    </tr>
    <tr>
      <th>59</th>
      <td>PXD004720</td>
      <td>bread wheat</td>
      <td>root-vasculature</td>
      <td>single</td>
      <td>protein</td>
      <td>FragPipe_Duncan_PXD004720_root-vasculature_pro...</td>
      <td>7577.0</td>
      <td>NaN</td>
      <td>7577.0</td>
      <td>9</td>
      <td>7568</td>
    </tr>
    <tr>
      <th>60</th>
      <td>PXD004720</td>
      <td>bread wheat</td>
      <td>spike-immature</td>
      <td>single</td>
      <td>peptide</td>
      <td>FragPipe_Duncan_PXD004720_spike-immature_pepti...</td>
      <td>36414.0</td>
      <td>36414.0</td>
      <td>NaN</td>
      <td>44</td>
      <td>36370</td>
    </tr>
    <tr>
      <th>61</th>
      <td>PXD004720</td>
      <td>bread wheat</td>
      <td>spike-immature</td>
      <td>single</td>
      <td>protein</td>
      <td>FragPipe_Duncan_PXD004720_spike-immature_prote...</td>
      <td>10948.0</td>
      <td>NaN</td>
      <td>10948.0</td>
      <td>8</td>
      <td>10940</td>
    </tr>
    <tr>
      <th>62</th>
      <td>PXD004720</td>
      <td>bread wheat</td>
      <td>stem</td>
      <td>single</td>
      <td>peptide</td>
      <td>FragPipe_Duncan_PXD004720_stem_peptide_annotat...</td>
      <td>14389.0</td>
      <td>14389.0</td>
      <td>NaN</td>
      <td>51</td>
      <td>14338</td>
    </tr>
    <tr>
      <th>63</th>
      <td>PXD004720</td>
      <td>bread wheat</td>
      <td>stem</td>
      <td>single</td>
      <td>protein</td>
      <td>FragPipe_Duncan_PXD004720_stem_protein_annotat...</td>
      <td>8984.0</td>
      <td>NaN</td>
      <td>8984.0</td>
      <td>10</td>
      <td>8974</td>
    </tr>
  </tbody>
</table>
<p>64 rows × 11 columns</p>
</div>


# Step 7 — Build Non-Contaminant Peptide–Protein Evidence Tables

This step converts the annotated FragPipe peptide-level outputs into standardised peptide–protein evidence tables for each wheat tissue.

Only wheat-derived identifications were retained for downstream analysis.

### Filtering rule

```text
Contaminant == "no"
```

This removed cRAP contaminants and other non-wheat entries before protein mapping and genome projection.

---

## Purpose

The aim of this step was to create a clean intermediate table linking each identified peptide sequence to its associated wheat protein accession(s).

This table provides the bridge between:

```text
FragPipe peptide identifications
```

and:

```text
protein accession → gene model → genomic coordinates
```

---

## Protein and isoform handling

FragPipe reports a primary protein accession in the `Protein` column and, when available, additional related accessions in the `Mapped Proteins` column.

To preserve protein isoform and shared-peptide information, this step expands the peptide table into peptide–protein pairs.

### Protein mapping categories

| Category | Description |
|---|---|
| primary | Protein accession reported in the FragPipe `Protein` column |
| mapped | Additional protein accession reported in the FragPipe `Mapped Proteins` column |

This avoids prematurely collapsing peptides that may support multiple protein isoforms or closely related wheat gene models.

---

## Output files

One peptide–protein evidence table was exported per tissue.

### Example output

```text
FragPipe_Duncan_PXD004720_anther_peptide_protein_evidence.csv
```

### Output directory

```text
python_outputs/tables/
```

---

## Summary metrics

A Step 7 summary table was generated across all tissues.

### Metrics captured

| Metric | Description |
|---|---|
| Non_contaminant_peptide_protein_pairs | Total number of retained peptide–protein evidence rows |
| Unique_peptides | Number of unique peptide sequences |
| Unique_proteins | Number of unique wheat protein accessions |
| Primary_peptide_protein_pairs | Number of primary peptide–protein associations |
| Mapped_peptide_protein_pairs | Number of additional mapped protein/isoform associations |
| Peptides_mapping_to_multiple_proteins | Number of peptides associated with more than one protein accession |
| Proteins_supported_by_one_peptide | Number of proteins supported by one unique peptide |
| Proteins_supported_by_two_or_more_peptides | Number of proteins supported by at least two unique peptides |

### Output summary file

```text
wheat_fragpipe_peptide_protein_evidence_summary_step7.csv
```

This evidence table is used in the next step to link protein accessions to wheat gene models and genome annotation coordinates.


```python
# ============================================================
# Step 7 — Build non-contaminant peptide–protein evidence tables
# ============================================================

import pandas as pd
from pathlib import Path

# -----------------------------
# 1. Input / output paths
# -----------------------------
fragpipe_dir = Path("FragPipe_results")
tables_dir = Path("python_outputs/tables")
tables_dir.mkdir(parents=True, exist_ok=True)

manifest_file = fragpipe_dir / "wheat_tissues_FragPipe-result-manifest_2026-05-11.csv"
step6_summary_file = tables_dir / "wheat_fragpipe_annotation_summary_step6.csv"

step7_summary_out = tables_dir / "wheat_fragpipe_peptide_protein_evidence_summary_step7.csv"

# -----------------------------
# 2. Load manifest
# -----------------------------
manifest = pd.read_csv(manifest_file, encoding="utf-8-sig")


# -----------------------------
# 3. Helper functions
# -----------------------------
def split_mapped_proteins(value):
    """
    Split FragPipe mapped protein accessions into a clean list.

    Handles empty values and common separators used in protein tables.
    """
    if pd.isna(value):
        return []

    value = str(value).strip()

    if value == "" or value.lower() in ["nan", "na", "none"]:
        return []

    for sep in [";", ","]:
        value = value.replace(sep, "|")

    return [x.strip() for x in value.split("|") if x.strip()]


def find_optional_column(data, possible_names):
    """
    Return the first matching column name from a list of possible names.
    """
    for col in possible_names:
        if col in data.columns:
            return col
    return None


def build_peptide_protein_evidence(source, tissue_raw_code):
    """
    Build a non-contaminant peptide–protein evidence table for one tissue.

    Parameters
    ----------
    source : str
        Proteomics dataset source, e.g. "PXD004720".

    tissue_raw_code : str
        Tissue code as listed in the manifest, e.g. "anther".

    Returns
    -------
    pd.DataFrame
        Expanded peptide–protein evidence table.
    """

    # Find matching manifest row
    match = manifest[
        (manifest["Source"] == source) &
        (manifest["Tissue-Raw-Code"] == tissue_raw_code)
    ]

    if match.empty:
        raise ValueError(
            f"No manifest entry found for Source='{source}' and "
            f"Tissue-Raw-Code='{tissue_raw_code}'."
        )

    if len(match) > 1:
        raise ValueError(
            f"Multiple manifest entries found for Source='{source}' and "
            f"Tissue-Raw-Code='{tissue_raw_code}'."
        )

    row = match.iloc[0]

    # Input annotated peptide file from Step 6
    peptide_filename = row["FragPipe-Output-Peptide"]
    annotated_filename = peptide_filename.replace(".tsv", "_annotated.csv")
    annotated_path = tables_dir / annotated_filename

    if not annotated_path.exists():
        raise FileNotFoundError(f"Annotated peptide file not found: {annotated_path}")

    # Read annotated peptide table
    data = pd.read_csv(annotated_path)

    # Required columns
    required_cols = ["Peptide", "Protein", "Contaminant"]
    missing_cols = [col for col in required_cols if col not in data.columns]

    if missing_cols:
        raise KeyError(
            f"Missing required column(s) in {annotated_filename}: {missing_cols}"
        )

    # Keep wheat-derived identifications only
    data = data[data["Contaminant"] == "no"].copy()

    # Optional FragPipe evidence columns
    mapped_col = find_optional_column(
        data,
        ["Mapped Proteins", "Mapped Protein", "Mapped_Proteins", "Mapped proteins"]
    )

    probability_col = find_optional_column(
        data,
        ["Probability", "PeptideProphet Probability", "Peptide Probability"]
    )

    spectral_count_col = find_optional_column(
        data,
        ["Spectral Count", "Spectral_Count", "SpectralCount"]
    )

    charge_col = find_optional_column(
        data,
        ["Charges", "Charge"]
    )

    records = []

    for _, peptide_row in data.iterrows():

        peptide = peptide_row["Peptide"]
        primary_protein = peptide_row["Protein"]

        # Primary protein evidence
        protein_records = [{
            "ProteinID": primary_protein,
            "Protein_mapping_type": "primary"
        }]

        # Additional mapped proteins / isoforms, when present
        if mapped_col is not None:
            mapped_proteins = split_mapped_proteins(peptide_row[mapped_col])

            for mapped_protein in mapped_proteins:
                if mapped_protein != primary_protein:
                    protein_records.append({
                        "ProteinID": mapped_protein,
                        "Protein_mapping_type": "mapped"
                    })

        for protein_record in protein_records:

            record = {
                "Source": row["Source"],
                "Species": row["Species"],
                "Tissue": row["Tissue-Raw-Code"],
                "Batch": row["Batch"],
                "Peptide": peptide,
                "ProteinID": protein_record["ProteinID"],
                "Protein_mapping_type": protein_record["Protein_mapping_type"],
                "Contaminant": "no"
            }

            if probability_col is not None:
                record["Probability"] = peptide_row[probability_col]

            if spectral_count_col is not None:
                record["Spectral_Count"] = peptide_row[spectral_count_col]

            if charge_col is not None:
                record["Charges"] = peptide_row[charge_col]

            records.append(record)

    evidence = pd.DataFrame(records)

    # Add sequential index
    evidence.insert(0, "Index", range(1, len(evidence) + 1))

    # Remove accidental duplicated peptide–protein rows within a tissue
    # This should usually not be needed, but keeps the output robust.
    evidence = evidence.drop_duplicates(
        subset=["Source", "Tissue", "Peptide", "ProteinID", "Protein_mapping_type"]
    ).reset_index(drop=True)

    evidence["Index"] = range(1, len(evidence) + 1)

    # Save output
    output_filename = peptide_filename.replace(
        "_peptide.tsv",
        "_peptide_protein_evidence.csv"
    )

    output_path = tables_dir / output_filename
    evidence.to_csv(output_path, index=False)

    print(f"Saved peptide–protein evidence table: {output_path}")
    print(f"Rows: {len(evidence):,}")
    print(f"Unique peptides: {evidence['Peptide'].nunique():,}")
    print(f"Unique proteins: {evidence['ProteinID'].nunique():,}")

    return evidence


# -----------------------------
# 4. Run Step 7 for all tissues
# -----------------------------
step7_summary_records = []

for _, row in manifest.iterrows():

    source = row["Source"]
    species = row["Species"]
    tissue_raw_code = row["Tissue-Raw-Code"]
    batch = row["Batch"]

    print(f"\nProcessing {source} | {tissue_raw_code}")

    evidence = build_peptide_protein_evidence(
        source=source,
        tissue_raw_code=tissue_raw_code
    )

    output_filename = row["FragPipe-Output-Peptide"].replace(
        "_peptide.tsv",
        "_peptide_protein_evidence.csv"
    )

    primary_evidence = evidence[evidence["Protein_mapping_type"] == "primary"]
    mapped_evidence = evidence[evidence["Protein_mapping_type"] == "mapped"]

    peptides_per_protein = (
        evidence.groupby("ProteinID")["Peptide"]
        .nunique()
        .reset_index(name="unique_peptides_per_protein")
    )

    proteins_per_peptide = (
        evidence.groupby("Peptide")["ProteinID"]
        .nunique()
        .reset_index(name="protein_isoforms_per_peptide")
    )

    step7_summary_records.append({
        "Source": source,
        "Species": species,
        "Tissue": tissue_raw_code,
        "Batch": batch,
        "Evidence_file": output_filename,
        "Non_contaminant_peptide_protein_pairs": len(evidence),
        "Unique_peptides": evidence["Peptide"].nunique(),
        "Unique_proteins": evidence["ProteinID"].nunique(),
        "Primary_peptide_protein_pairs": len(primary_evidence),
        "Mapped_peptide_protein_pairs": len(mapped_evidence),
        "Peptides_mapping_to_multiple_proteins": (
            proteins_per_peptide["protein_isoforms_per_peptide"] > 1
        ).sum(),
        "Proteins_supported_by_one_peptide": (
            peptides_per_protein["unique_peptides_per_protein"] == 1
        ).sum(),
        "Proteins_supported_by_two_or_more_peptides": (
            peptides_per_protein["unique_peptides_per_protein"] >= 2
        ).sum()
    })

step7_summary = pd.DataFrame(step7_summary_records)

# Save Step 7 summary
step7_summary.to_csv(step7_summary_out, index=False)

print(f"\nStep 7 summary saved: {step7_summary_out}")
print(f"Summary rows: {step7_summary.shape[0]:,}")

display(step7_summary)
```

    
    Processing MSV000090572 | stored_grain
    Saved peptide–protein evidence table: python_outputs\tables\FragPipe_Vincent_MSV000090572_stored-grain_peptide_protein_evidence.csv
    Rows: 30,317
    Unique peptides: 9,329
    Unique proteins: 17,481
    
    Processing PXD050500 | coleoptile
    Saved peptide–protein evidence table: python_outputs\tables\FragPipe_Liu_PXD050500_coleoptile_peptide_protein_evidence.csv
    Rows: 1,850,247
    Unique peptides: 582,775
    Unique proteins: 239,514
    
    Processing PXD050500 | node
    Saved peptide–protein evidence table: python_outputs\tables\FragPipe_Liu_PXD050500_node_peptide_protein_evidence.csv
    Rows: 1,887,009
    Unique peptides: 595,954
    Unique proteins: 242,842
    
    Processing PXD050500 | radicle
    Saved peptide–protein evidence table: python_outputs\tables\FragPipe_Liu_PXD050500_radicle_peptide_protein_evidence.csv
    Rows: 1,062,587
    Unique peptides: 333,058
    Unique proteins: 206,049
    
    Processing PXD004720 | anther
    Saved peptide–protein evidence table: python_outputs\tables\FragPipe_Duncan_PXD004720_anther_peptide_protein_evidence.csv
    Rows: 164,397
    Unique peptides: 34,098
    Unique proteins: 36,839
    
    Processing PXD004720 | boot
    Saved peptide–protein evidence table: python_outputs\tables\FragPipe_Duncan_PXD004720_boot_peptide_protein_evidence.csv
    Rows: 13,230
    Unique peptides: 3,643
    Unique proteins: 9,211
    
    Processing PXD004720 | coleoptile
    Saved peptide–protein evidence table: python_outputs\tables\FragPipe_Duncan_PXD004720_coleoptile_peptide_protein_evidence.csv
    Rows: 204,518
    Unique peptides: 41,025
    Unique proteins: 46,513
    
    Processing PXD004720 | embryo
    Saved peptide–protein evidence table: python_outputs\tables\FragPipe_Duncan_PXD004720_embryo_peptide_protein_evidence.csv
    Rows: 8,852
    Unique peptides: 2,867
    Unique proteins: 7,220
    
    Processing PXD004720 | endosperm
    Saved peptide–protein evidence table: python_outputs\tables\FragPipe_Duncan_PXD004720_endosperm_peptide_protein_evidence.csv
    Rows: 102,845
    Unique peptides: 20,081
    Unique proteins: 27,461
    
    Processing PXD004720 | glume
    Saved peptide–protein evidence table: python_outputs\tables\FragPipe_Duncan_PXD004720_glume_peptide_protein_evidence.csv
    Rows: 145,154
    Unique peptides: 28,648
    Unique proteins: 35,551
    
    Processing PXD004720 | grain-zadoks-70
    Saved peptide–protein evidence table: python_outputs\tables\FragPipe_Duncan_PXD004720_grain-zadoks-70_peptide_protein_evidence.csv
    Rows: 126,227
    Unique peptides: 26,196
    Unique proteins: 35,549
    
    Processing PXD004720 | grain-zadoks-71
    Saved peptide–protein evidence table: python_outputs\tables\FragPipe_Duncan_PXD004720_grain-zadoks-71_peptide_protein_evidence.csv
    Rows: 179,751
    Unique peptides: 35,960
    Unique proteins: 47,089
    
    Processing PXD004720 | grain-zadoks-75
    Saved peptide–protein evidence table: python_outputs\tables\FragPipe_Duncan_PXD004720_grain-zadoks-75_peptide_protein_evidence.csv
    Rows: 132,719
    Unique peptides: 27,463
    Unique proteins: 41,042
    
    Processing PXD004720 | grain-zadoks-83
    Saved peptide–protein evidence table: python_outputs\tables\FragPipe_Duncan_PXD004720_grain-zadoks-83_peptide_protein_evidence.csv
    Rows: 113,236
    Unique peptides: 23,990
    Unique proteins: 36,654
    
    Processing PXD004720 | grain-zadoks-87
    Saved peptide–protein evidence table: python_outputs\tables\FragPipe_Duncan_PXD004720_grain-zadoks-87_peptide_protein_evidence.csv
    Rows: 111,819
    Unique peptides: 24,636
    Unique proteins: 34,889
    
    Processing PXD004720 | leaf-flag-mature
    Saved peptide–protein evidence table: python_outputs\tables\FragPipe_Duncan_PXD004720_leaf-flag-mature_peptide_protein_evidence.csv
    Rows: 145,650
    Unique peptides: 29,840
    Unique proteins: 37,963
    
    Processing PXD004720 | leaf-flag-senescing
    Saved peptide–protein evidence table: python_outputs\tables\FragPipe_Duncan_PXD004720_leaf-flag-senescing_peptide_protein_evidence.csv
    Rows: 46,206
    Unique peptides: 11,354
    Unique proteins: 25,701
    
    Processing PXD004720 | leaf-flag-young
    Saved peptide–protein evidence table: python_outputs\tables\FragPipe_Duncan_PXD004720_leaf-flag-young_peptide_protein_evidence.csv
    Rows: 120,833
    Unique peptides: 23,373
    Unique proteins: 32,629
    
    Processing PXD004720 | lemma
    Saved peptide–protein evidence table: python_outputs\tables\FragPipe_Duncan_PXD004720_lemma_peptide_protein_evidence.csv
    Rows: 147,650
    Unique peptides: 29,788
    Unique proteins: 37,946
    
    Processing PXD004720 | node
    Saved peptide–protein evidence table: python_outputs\tables\FragPipe_Duncan_PXD004720_node_peptide_protein_evidence.csv
    Rows: 99,844
    Unique peptides: 21,457
    Unique proteins: 36,555
    
    Processing PXD004720 | node_secretion
    Saved peptide–protein evidence table: python_outputs\tables\FragPipe_Duncan_PXD004720_node-secretion_peptide_protein_evidence.csv
    Rows: 169,705
    Unique peptides: 34,124
    Unique proteins: 42,371
    
    Processing PXD004720 | palea
    Saved peptide–protein evidence table: python_outputs\tables\FragPipe_Duncan_PXD004720_palea_peptide_protein_evidence.csv
    Rows: 116,960
    Unique peptides: 21,707
    Unique proteins: 27,591
    
    Processing PXD004720 | pericarp
    Saved peptide–protein evidence table: python_outputs\tables\FragPipe_Duncan_PXD004720_pericarp_peptide_protein_evidence.csv
    Rows: 131,245
    Unique peptides: 28,332
    Unique proteins: 32,676
    
    Processing PXD004720 | pollen
    Saved peptide–protein evidence table: python_outputs\tables\FragPipe_Duncan_PXD004720_pollen_peptide_protein_evidence.csv
    Rows: 74,722
    Unique peptides: 13,502
    Unique proteins: 19,942
    
    Processing PXD004720 | rachilla
    Saved peptide–protein evidence table: python_outputs\tables\FragPipe_Duncan_PXD004720_rachilla_peptide_protein_evidence.csv
    Rows: 160,234
    Unique peptides: 31,145
    Unique proteins: 37,219
    
    Processing PXD004720 | radicle
    Saved peptide–protein evidence table: python_outputs\tables\FragPipe_Duncan_PXD004720_radicle_peptide_protein_evidence.csv
    Rows: 194,730
    Unique peptides: 39,643
    Unique proteins: 42,976
    
    Processing PXD004720 | root-mature
    Saved peptide–protein evidence table: python_outputs\tables\FragPipe_Duncan_PXD004720_root-mature_peptide_protein_evidence.csv
    Rows: 97,211
    Unique peptides: 21,350
    Unique proteins: 39,165
    
    Processing PXD004720 | root-secretion
    Saved peptide–protein evidence table: python_outputs\tables\FragPipe_Duncan_PXD004720_root-secretion_peptide_protein_evidence.csv
    Rows: 100,654
    Unique peptides: 21,284
    Unique proteins: 32,049
    
    Processing PXD004720 | root-tip
    Saved peptide–protein evidence table: python_outputs\tables\FragPipe_Duncan_PXD004720_root-tip_peptide_protein_evidence.csv
    Rows: 191,973
    Unique peptides: 38,634
    Unique proteins: 39,785
    
    Processing PXD004720 | root-vasculature
    Saved peptide–protein evidence table: python_outputs\tables\FragPipe_Duncan_PXD004720_root-vasculature_peptide_protein_evidence.csv
    Rows: 111,846
    Unique peptides: 20,815
    Unique proteins: 30,631
    
    Processing PXD004720 | spike-immature
    Saved peptide–protein evidence table: python_outputs\tables\FragPipe_Duncan_PXD004720_spike-immature_peptide_protein_evidence.csv
    Rows: 188,868
    Unique peptides: 36,370
    Unique proteins: 40,220
    
    Processing PXD004720 | stem
    Saved peptide–protein evidence table: python_outputs\tables\FragPipe_Duncan_PXD004720_stem_peptide_protein_evidence.csv
    Rows: 60,749
    Unique peptides: 14,338
    Unique proteins: 29,417
    
    Step 7 summary saved: python_outputs\tables\wheat_fragpipe_peptide_protein_evidence_summary_step7.csv
    Summary rows: 32
    


<div>
<style scoped>
    .dataframe tbody tr th:only-of-type {
        vertical-align: middle;
    }

    .dataframe tbody tr th {
        vertical-align: top;
    }

    .dataframe thead th {
        text-align: right;
    }
</style>
<table border="1" class="dataframe">
  <thead>
    <tr style="text-align: right;">
      <th></th>
      <th>Source</th>
      <th>Species</th>
      <th>Tissue</th>
      <th>Batch</th>
      <th>Evidence_file</th>
      <th>Non_contaminant_peptide_protein_pairs</th>
      <th>Unique_peptides</th>
      <th>Unique_proteins</th>
      <th>Primary_peptide_protein_pairs</th>
      <th>Mapped_peptide_protein_pairs</th>
      <th>Peptides_mapping_to_multiple_proteins</th>
      <th>Proteins_supported_by_one_peptide</th>
      <th>Proteins_supported_by_two_or_more_peptides</th>
    </tr>
  </thead>
  <tbody>
    <tr>
      <th>0</th>
      <td>MSV000090572</td>
      <td>bread wheat</td>
      <td>stored_grain</td>
      <td>single</td>
      <td>FragPipe_Vincent_MSV000090572_stored-grain_pep...</td>
      <td>30317</td>
      <td>9329</td>
      <td>17481</td>
      <td>9329</td>
      <td>20988</td>
      <td>4290</td>
      <td>14135</td>
      <td>3346</td>
    </tr>
    <tr>
      <th>1</th>
      <td>PXD050500</td>
      <td>bread wheat</td>
      <td>coleoptile</td>
      <td>single</td>
      <td>FragPipe_Liu_PXD050500_coleoptile_peptide_prot...</td>
      <td>1850247</td>
      <td>582775</td>
      <td>239514</td>
      <td>582775</td>
      <td>1267472</td>
      <td>338464</td>
      <td>59431</td>
      <td>180083</td>
    </tr>
    <tr>
      <th>2</th>
      <td>PXD050500</td>
      <td>bread wheat</td>
      <td>node</td>
      <td>single</td>
      <td>FragPipe_Liu_PXD050500_node_peptide_protein_ev...</td>
      <td>1887009</td>
      <td>595954</td>
      <td>242842</td>
      <td>595954</td>
      <td>1291055</td>
      <td>340536</td>
      <td>57486</td>
      <td>185356</td>
    </tr>
    <tr>
      <th>3</th>
      <td>PXD050500</td>
      <td>bread wheat</td>
      <td>radicle</td>
      <td>single</td>
      <td>FragPipe_Liu_PXD050500_radicle_peptide_protein...</td>
      <td>1062587</td>
      <td>333058</td>
      <td>206049</td>
      <td>333058</td>
      <td>729529</td>
      <td>190016</td>
      <td>68213</td>
      <td>137836</td>
    </tr>
    <tr>
      <th>4</th>
      <td>PXD004720</td>
      <td>bread wheat</td>
      <td>anther</td>
      <td>single</td>
      <td>FragPipe_Duncan_PXD004720_anther_peptide_prote...</td>
      <td>164397</td>
      <td>34098</td>
      <td>36839</td>
      <td>34098</td>
      <td>130299</td>
      <td>27707</td>
      <td>19944</td>
      <td>16895</td>
    </tr>
    <tr>
      <th>5</th>
      <td>PXD004720</td>
      <td>bread wheat</td>
      <td>boot</td>
      <td>single</td>
      <td>FragPipe_Duncan_PXD004720_boot_peptide_protein...</td>
      <td>13230</td>
      <td>3643</td>
      <td>9211</td>
      <td>3643</td>
      <td>9587</td>
      <td>2302</td>
      <td>7288</td>
      <td>1923</td>
    </tr>
    <tr>
      <th>6</th>
      <td>PXD004720</td>
      <td>bread wheat</td>
      <td>coleoptile</td>
      <td>single</td>
      <td>FragPipe_Duncan_PXD004720_coleoptile_peptide_p...</td>
      <td>204518</td>
      <td>41025</td>
      <td>46513</td>
      <td>41025</td>
      <td>163493</td>
      <td>34608</td>
      <td>23582</td>
      <td>22931</td>
    </tr>
    <tr>
      <th>7</th>
      <td>PXD004720</td>
      <td>bread wheat</td>
      <td>embryo</td>
      <td>single</td>
      <td>FragPipe_Duncan_PXD004720_embryo_peptide_prote...</td>
      <td>8852</td>
      <td>2867</td>
      <td>7220</td>
      <td>2867</td>
      <td>5985</td>
      <td>1581</td>
      <td>6285</td>
      <td>935</td>
    </tr>
    <tr>
      <th>8</th>
      <td>PXD004720</td>
      <td>bread wheat</td>
      <td>endosperm</td>
      <td>single</td>
      <td>FragPipe_Duncan_PXD004720_endosperm_peptide_pr...</td>
      <td>102845</td>
      <td>20081</td>
      <td>27461</td>
      <td>20081</td>
      <td>82764</td>
      <td>15925</td>
      <td>15845</td>
      <td>11616</td>
    </tr>
    <tr>
      <th>9</th>
      <td>PXD004720</td>
      <td>bread wheat</td>
      <td>glume</td>
      <td>single</td>
      <td>FragPipe_Duncan_PXD004720_glume_peptide_protei...</td>
      <td>145154</td>
      <td>28648</td>
      <td>35551</td>
      <td>28648</td>
      <td>116506</td>
      <td>23601</td>
      <td>19974</td>
      <td>15577</td>
    </tr>
    <tr>
      <th>10</th>
      <td>PXD004720</td>
      <td>bread wheat</td>
      <td>grain-zadoks-70</td>
      <td>single</td>
      <td>FragPipe_Duncan_PXD004720_grain-zadoks-70_pept...</td>
      <td>126227</td>
      <td>26196</td>
      <td>35549</td>
      <td>26196</td>
      <td>100031</td>
      <td>20748</td>
      <td>20524</td>
      <td>15025</td>
    </tr>
    <tr>
      <th>11</th>
      <td>PXD004720</td>
      <td>bread wheat</td>
      <td>grain-zadoks-71</td>
      <td>single</td>
      <td>FragPipe_Duncan_PXD004720_grain-zadoks-71_pept...</td>
      <td>179751</td>
      <td>35960</td>
      <td>47089</td>
      <td>35960</td>
      <td>143791</td>
      <td>30079</td>
      <td>24698</td>
      <td>22391</td>
    </tr>
    <tr>
      <th>12</th>
      <td>PXD004720</td>
      <td>bread wheat</td>
      <td>grain-zadoks-75</td>
      <td>single</td>
      <td>FragPipe_Duncan_PXD004720_grain-zadoks-75_pept...</td>
      <td>132719</td>
      <td>27463</td>
      <td>41042</td>
      <td>27463</td>
      <td>105256</td>
      <td>21885</td>
      <td>23263</td>
      <td>17779</td>
    </tr>
    <tr>
      <th>13</th>
      <td>PXD004720</td>
      <td>bread wheat</td>
      <td>grain-zadoks-83</td>
      <td>single</td>
      <td>FragPipe_Duncan_PXD004720_grain-zadoks-83_pept...</td>
      <td>113236</td>
      <td>23990</td>
      <td>36654</td>
      <td>23990</td>
      <td>89246</td>
      <td>18722</td>
      <td>21688</td>
      <td>14966</td>
    </tr>
    <tr>
      <th>14</th>
      <td>PXD004720</td>
      <td>bread wheat</td>
      <td>grain-zadoks-87</td>
      <td>single</td>
      <td>FragPipe_Duncan_PXD004720_grain-zadoks-87_pept...</td>
      <td>111819</td>
      <td>24636</td>
      <td>34889</td>
      <td>24636</td>
      <td>87183</td>
      <td>18599</td>
      <td>21245</td>
      <td>13644</td>
    </tr>
    <tr>
      <th>15</th>
      <td>PXD004720</td>
      <td>bread wheat</td>
      <td>leaf-flag-mature</td>
      <td>single</td>
      <td>FragPipe_Duncan_PXD004720_leaf-flag-mature_pep...</td>
      <td>145650</td>
      <td>29840</td>
      <td>37963</td>
      <td>29840</td>
      <td>115810</td>
      <td>24930</td>
      <td>19764</td>
      <td>18199</td>
    </tr>
    <tr>
      <th>16</th>
      <td>PXD004720</td>
      <td>bread wheat</td>
      <td>leaf-flag-senescing</td>
      <td>single</td>
      <td>FragPipe_Duncan_PXD004720_leaf-flag-senescing_...</td>
      <td>46206</td>
      <td>11354</td>
      <td>25701</td>
      <td>11354</td>
      <td>34852</td>
      <td>7264</td>
      <td>18840</td>
      <td>6861</td>
    </tr>
    <tr>
      <th>17</th>
      <td>PXD004720</td>
      <td>bread wheat</td>
      <td>leaf-flag-young</td>
      <td>single</td>
      <td>FragPipe_Duncan_PXD004720_leaf-flag-young_pept...</td>
      <td>120833</td>
      <td>23373</td>
      <td>32629</td>
      <td>23373</td>
      <td>97460</td>
      <td>19862</td>
      <td>17539</td>
      <td>15090</td>
    </tr>
    <tr>
      <th>18</th>
      <td>PXD004720</td>
      <td>bread wheat</td>
      <td>lemma</td>
      <td>single</td>
      <td>FragPipe_Duncan_PXD004720_lemma_peptide_protei...</td>
      <td>147650</td>
      <td>29788</td>
      <td>37946</td>
      <td>29788</td>
      <td>117862</td>
      <td>24643</td>
      <td>20871</td>
      <td>17075</td>
    </tr>
    <tr>
      <th>19</th>
      <td>PXD004720</td>
      <td>bread wheat</td>
      <td>node</td>
      <td>single</td>
      <td>FragPipe_Duncan_PXD004720_node_peptide_protein...</td>
      <td>99844</td>
      <td>21457</td>
      <td>36555</td>
      <td>21457</td>
      <td>78387</td>
      <td>15947</td>
      <td>23379</td>
      <td>13176</td>
    </tr>
    <tr>
      <th>20</th>
      <td>PXD004720</td>
      <td>bread wheat</td>
      <td>node_secretion</td>
      <td>single</td>
      <td>FragPipe_Duncan_PXD004720_node-secretion_pepti...</td>
      <td>169705</td>
      <td>34124</td>
      <td>42371</td>
      <td>34124</td>
      <td>135581</td>
      <td>28864</td>
      <td>21386</td>
      <td>20985</td>
    </tr>
    <tr>
      <th>21</th>
      <td>PXD004720</td>
      <td>bread wheat</td>
      <td>palea</td>
      <td>single</td>
      <td>FragPipe_Duncan_PXD004720_palea_peptide_protei...</td>
      <td>116960</td>
      <td>21707</td>
      <td>27591</td>
      <td>21707</td>
      <td>95253</td>
      <td>18324</td>
      <td>14027</td>
      <td>13564</td>
    </tr>
    <tr>
      <th>22</th>
      <td>PXD004720</td>
      <td>bread wheat</td>
      <td>pericarp</td>
      <td>single</td>
      <td>FragPipe_Duncan_PXD004720_pericarp_peptide_pro...</td>
      <td>131245</td>
      <td>28332</td>
      <td>32676</td>
      <td>28332</td>
      <td>102913</td>
      <td>21946</td>
      <td>18600</td>
      <td>14076</td>
    </tr>
    <tr>
      <th>23</th>
      <td>PXD004720</td>
      <td>bread wheat</td>
      <td>pollen</td>
      <td>single</td>
      <td>FragPipe_Duncan_PXD004720_pollen_peptide_prote...</td>
      <td>74722</td>
      <td>13502</td>
      <td>19942</td>
      <td>13502</td>
      <td>61220</td>
      <td>11116</td>
      <td>10811</td>
      <td>9131</td>
    </tr>
    <tr>
      <th>24</th>
      <td>PXD004720</td>
      <td>bread wheat</td>
      <td>rachilla</td>
      <td>single</td>
      <td>FragPipe_Duncan_PXD004720_rachilla_peptide_pro...</td>
      <td>160234</td>
      <td>31145</td>
      <td>37219</td>
      <td>31145</td>
      <td>129089</td>
      <td>26163</td>
      <td>19600</td>
      <td>17619</td>
    </tr>
    <tr>
      <th>25</th>
      <td>PXD004720</td>
      <td>bread wheat</td>
      <td>radicle</td>
      <td>single</td>
      <td>FragPipe_Duncan_PXD004720_radicle_peptide_prot...</td>
      <td>194730</td>
      <td>39643</td>
      <td>42976</td>
      <td>39643</td>
      <td>155087</td>
      <td>32778</td>
      <td>22396</td>
      <td>20580</td>
    </tr>
    <tr>
      <th>26</th>
      <td>PXD004720</td>
      <td>bread wheat</td>
      <td>root-mature</td>
      <td>single</td>
      <td>FragPipe_Duncan_PXD004720_root-mature_peptide_...</td>
      <td>97211</td>
      <td>21350</td>
      <td>39165</td>
      <td>21350</td>
      <td>75861</td>
      <td>14600</td>
      <td>26303</td>
      <td>12862</td>
    </tr>
    <tr>
      <th>27</th>
      <td>PXD004720</td>
      <td>bread wheat</td>
      <td>root-secretion</td>
      <td>single</td>
      <td>FragPipe_Duncan_PXD004720_root-secretion_pepti...</td>
      <td>100654</td>
      <td>21284</td>
      <td>32049</td>
      <td>21284</td>
      <td>79370</td>
      <td>15607</td>
      <td>20041</td>
      <td>12008</td>
    </tr>
    <tr>
      <th>28</th>
      <td>PXD004720</td>
      <td>bread wheat</td>
      <td>root-tip</td>
      <td>single</td>
      <td>FragPipe_Duncan_PXD004720_root-tip_peptide_pro...</td>
      <td>191973</td>
      <td>38634</td>
      <td>39785</td>
      <td>38634</td>
      <td>153339</td>
      <td>33012</td>
      <td>18865</td>
      <td>20920</td>
    </tr>
    <tr>
      <th>29</th>
      <td>PXD004720</td>
      <td>bread wheat</td>
      <td>root-vasculature</td>
      <td>single</td>
      <td>FragPipe_Duncan_PXD004720_root-vasculature_pep...</td>
      <td>111846</td>
      <td>20815</td>
      <td>30631</td>
      <td>20815</td>
      <td>91031</td>
      <td>17251</td>
      <td>16694</td>
      <td>13937</td>
    </tr>
    <tr>
      <th>30</th>
      <td>PXD004720</td>
      <td>bread wheat</td>
      <td>spike-immature</td>
      <td>single</td>
      <td>FragPipe_Duncan_PXD004720_spike-immature_pepti...</td>
      <td>188868</td>
      <td>36370</td>
      <td>40220</td>
      <td>36370</td>
      <td>152498</td>
      <td>30739</td>
      <td>19857</td>
      <td>20363</td>
    </tr>
    <tr>
      <th>31</th>
      <td>PXD004720</td>
      <td>bread wheat</td>
      <td>stem</td>
      <td>single</td>
      <td>FragPipe_Duncan_PXD004720_stem_peptide_protein...</td>
      <td>60749</td>
      <td>14338</td>
      <td>29417</td>
      <td>14338</td>
      <td>46411</td>
      <td>9710</td>
      <td>20198</td>
      <td>9219</td>
    </tr>
  </tbody>
</table>
</div>


# Step 8 — Link Peptide–Protein Evidence to Wheat Gene Models

This step links the non-contaminant peptide–protein evidence tables generated in Step 7 to wheat gene model annotations derived from the IWGSC RefSeq v2.1 GFF3 files.

The purpose of this step is to connect FragPipe protein accessions to their corresponding wheat gene models, transcripts, and genomic annotation features.

---

## Input files

### Peptide–protein evidence tables from Step 7

One file per tissue:

```text
FragPipe_FirstAuthor_Source_Tissue-Raw-Code_peptide_protein_evidence.csv
```

### Protein-to-gene mapping table from Step 5

```text
wheat_protein_gene_mapping_HC_LC.csv
```

This table was generated by parsing the high-confidence and low-confidence wheat GFF3 annotation files.

---

## Mapping strategy

Each peptide–protein evidence row was merged with the protein-to-gene mapping table using:

```text
ProteinID
```

This links each identified protein accession to available gene model annotation fields, including transcript and gene identifiers.

---

## Mapping status

A mapping status column was added to retain unmatched entries for audit purposes.

| Status | Description |
|---|---|
| mapped | ProteinID was found in the GFF3-derived protein-to-gene mapping table |
| unmapped | ProteinID was not found in the protein-to-gene mapping table |

Unmapped proteins were retained rather than discarded so that mapping completeness could be evaluated across tissues.

---

## Output files

One peptide–protein–gene mapping table was exported per tissue.

### Example output

```text
FragPipe_Duncan_PXD004720_anther_peptide_protein_gene_mapping.csv
```

### Output directory

```text
python_outputs/tables/
```

---

## Summary metrics

A Step 8 summary table was generated across all tissues.

### Metrics captured

| Metric | Description |
|---|---|
| Peptide_protein_pairs | Total number of peptide–protein evidence rows |
| Mapped_peptide_protein_pairs | Number of peptide–protein rows successfully linked to gene models |
| Unmapped_peptide_protein_pairs | Number of peptide–protein rows not linked to gene models |
| Mapping_rate_percent | Percentage of peptide–protein rows successfully mapped |
| Unique_peptides | Number of unique peptide sequences |
| Unique_proteins | Number of unique protein accessions |
| Mapped_unique_proteins | Number of unique protein accessions successfully mapped |
| Unmapped_unique_proteins | Number of unique protein accessions not mapped |
| Unique_gene_models | Number of unique gene models identified |
| Unique_transcripts | Number of unique transcripts identified |

### Output summary file

```text
wheat_peptide_protein_gene_mapping_summary_step8.csv
```

This mapped table is used in the next step to project peptide positions from protein coordinates onto wheat genomic coordinates.


```python
# ============================================================
# Step 8 — Link peptide–protein evidence to wheat gene models
# ============================================================

import pandas as pd
from pathlib import Path

# -----------------------------
# 1. Input / output paths
# -----------------------------
fragpipe_dir = Path("FragPipe_results")
tables_dir = Path("python_outputs/tables")
tables_dir.mkdir(parents=True, exist_ok=True)

manifest_file = fragpipe_dir / "wheat_tissues_FragPipe-result-manifest_2026-05-11.csv"

protein_gene_mapping_file = tables_dir / "wheat_protein_gene_mapping_HC_LC.csv"

step8_summary_out = tables_dir / "wheat_peptide_protein_gene_mapping_summary_step8.csv"

# -----------------------------
# 2. Load manifest and protein-to-gene mapping table
# -----------------------------
manifest = pd.read_csv(manifest_file, encoding="utf-8-sig")
protein_gene_mapping = pd.read_csv(protein_gene_mapping_file)

print("Protein-to-gene mapping table loaded:")
print(f"Rows: {protein_gene_mapping.shape[0]:,}")
print(f"Columns: {protein_gene_mapping.shape[1]:,}")

display(protein_gene_mapping.head())


# -----------------------------
# 3. Helper function
# -----------------------------
def map_peptide_protein_to_gene_models(source, tissue_raw_code):
    """
    Merge one tissue-level peptide–protein evidence table with the
    GFF3-derived wheat protein-to-gene mapping table.

    Parameters
    ----------
    source : str
        Proteomics dataset source, e.g. "PXD004720".

    tissue_raw_code : str
        Tissue code as listed in the manifest, e.g. "anther".

    Returns
    -------
    pd.DataFrame
        Peptide–protein–gene evidence table.
    """

    # Find matching manifest row
    match = manifest[
        (manifest["Source"] == source) &
        (manifest["Tissue-Raw-Code"] == tissue_raw_code)
    ]

    if match.empty:
        raise ValueError(
            f"No manifest entry found for Source='{source}' and "
            f"Tissue-Raw-Code='{tissue_raw_code}'."
        )

    if len(match) > 1:
        raise ValueError(
            f"Multiple manifest entries found for Source='{source}' and "
            f"Tissue-Raw-Code='{tissue_raw_code}'."
        )

    row = match.iloc[0]

    # Input Step 7 evidence file
    evidence_filename = row["FragPipe-Output-Peptide"].replace(
        "_peptide.tsv",
        "_peptide_protein_evidence.csv"
    )

    evidence_path = tables_dir / evidence_filename

    if not evidence_path.exists():
        raise FileNotFoundError(f"Step 7 evidence file not found: {evidence_path}")

    evidence = pd.read_csv(evidence_path)

    if "ProteinID" not in evidence.columns:
        raise KeyError(f"'ProteinID' column not found in {evidence_filename}")

    if "ProteinID" not in protein_gene_mapping.columns:
        raise KeyError("'ProteinID' column not found in protein-to-gene mapping table.")

    # Merge peptide–protein evidence with GFF3-derived mapping
    mapped = evidence.merge(
        protein_gene_mapping,
        on="ProteinID",
        how="left",
        indicator=True
    )

    mapped["Gene_model_mapping_status"] = mapped["_merge"].map({
        "both": "mapped",
        "left_only": "unmapped",
        "right_only": "mapping_only"
    })

    mapped = mapped.drop(columns=["_merge"])

    # Add sequential index
    if "Index" in mapped.columns:
        mapped = mapped.drop(columns=["Index"])

    mapped.insert(0, "Index", range(1, len(mapped) + 1))

    # Save output
    output_filename = evidence_filename.replace(
        "_peptide_protein_evidence.csv",
        "_peptide_protein_gene_mapping.csv"
    )

    output_path = tables_dir / output_filename
    mapped.to_csv(output_path, index=False)

    # Mapping summary
    total_pairs = len(mapped)
    mapped_pairs = (mapped["Gene_model_mapping_status"] == "mapped").sum()
    unmapped_pairs = (mapped["Gene_model_mapping_status"] == "unmapped").sum()

    print(f"\nSaved peptide–protein–gene mapping table: {output_path}")
    print(f"Rows: {total_pairs:,}")
    print(f"Mapped pairs: {mapped_pairs:,}")
    print(f"Unmapped pairs: {unmapped_pairs:,}")

    return mapped


# -----------------------------
# 4. Run Step 8 for all tissues
# -----------------------------
step8_summary_records = []

for _, row in manifest.iterrows():

    source = row["Source"]
    species = row["Species"]
    tissue_raw_code = row["Tissue-Raw-Code"]
    batch = row["Batch"]

    print(f"\nProcessing {source} | {tissue_raw_code}")

    mapped = map_peptide_protein_to_gene_models(
        source=source,
        tissue_raw_code=tissue_raw_code
    )

    output_filename = row["FragPipe-Output-Peptide"].replace(
        "_peptide.tsv",
        "_peptide_protein_gene_mapping.csv"
    )

    total_pairs = len(mapped)
    mapped_pairs = (mapped["Gene_model_mapping_status"] == "mapped").sum()
    unmapped_pairs = (mapped["Gene_model_mapping_status"] == "unmapped").sum()

    step8_summary_records.append({
        "Source": source,
        "Species": species,
        "Tissue": tissue_raw_code,
        "Batch": batch,
        "Gene_mapping_file": output_filename,
        "Peptide_protein_pairs": total_pairs,
        "Mapped_peptide_protein_pairs": mapped_pairs,
        "Unmapped_peptide_protein_pairs": unmapped_pairs,
        "Mapping_rate_percent": round((mapped_pairs / total_pairs) * 100, 2) if total_pairs > 0 else 0,
        "Unique_peptides": mapped["Peptide"].nunique() if "Peptide" in mapped.columns else pd.NA,
        "Unique_proteins": mapped["ProteinID"].nunique() if "ProteinID" in mapped.columns else pd.NA,
        "Mapped_unique_proteins": mapped.loc[
            mapped["Gene_model_mapping_status"] == "mapped", "ProteinID"
        ].nunique(),
        "Unmapped_unique_proteins": mapped.loc[
            mapped["Gene_model_mapping_status"] == "unmapped", "ProteinID"
        ].nunique(),
        "Unique_gene_models": mapped["GeneID"].nunique() if "GeneID" in mapped.columns else pd.NA,
        "Unique_transcripts": mapped["TranscriptID"].nunique() if "TranscriptID" in mapped.columns else pd.NA
    })

step8_summary = pd.DataFrame(step8_summary_records)

# Save Step 8 summary
step8_summary.to_csv(step8_summary_out, index=False)

print(f"\nStep 8 summary saved: {step8_summary_out}")
print(f"Summary rows: {step8_summary.shape[0]:,}")

display(step8_summary)
```

    C:\Users\Owner\AppData\Local\Temp\ipykernel_10260\2744825148.py:25: DtypeWarning: Columns (17,19) have mixed types. Specify dtype option on import or set low_memory=False.
      protein_gene_mapping = pd.read_csv(protein_gene_mapping_file)
    

    Protein-to-gene mapping table loaded:
    Rows: 295,914
    Columns: 24
    


<div>
<style scoped>
    .dataframe tbody tr th:only-of-type {
        vertical-align: middle;
    }

    .dataframe tbody tr th {
        vertical-align: top;
    }

    .dataframe thead th {
        text-align: right;
    }
</style>
<table border="1" class="dataframe">
  <thead>
    <tr style="text-align: right;">
      <th></th>
      <th>ProteinID</th>
      <th>GeneModel</th>
      <th>TranscriptID</th>
      <th>Chromosome</th>
      <th>Strand</th>
      <th>Gene_start</th>
      <th>Gene_end</th>
      <th>Transcript_start</th>
      <th>Transcript_end</th>
      <th>CDS_start</th>
      <th>...</th>
      <th>CDS_phase_values</th>
      <th>Annotation_confidence</th>
      <th>Gene_Name</th>
      <th>Gene_biotype</th>
      <th>Transcript_Name</th>
      <th>Transcript_biotype</th>
      <th>ProteinID_found_in_GFF3</th>
      <th>GeneModel_found_in_GFF3</th>
      <th>Has_gene_coordinates</th>
      <th>Has_transcript_coordinates</th>
    </tr>
  </thead>
  <tbody>
    <tr>
      <th>0</th>
      <td>TraesCS1A03G0000200.1</td>
      <td>TraesCS1A03G0000200</td>
      <td>TraesCS1A03G0000200.1</td>
      <td>Chr1A</td>
      <td>-</td>
      <td>40098</td>
      <td>70338</td>
      <td>40098</td>
      <td>70338</td>
      <td>58508</td>
      <td>...</td>
      <td>0</td>
      <td>HC</td>
      <td>TraesCS1A03G0000200</td>
      <td>NaN</td>
      <td>TraesCS1A03G0000200.1</td>
      <td>NaN</td>
      <td>True</td>
      <td>True</td>
      <td>True</td>
      <td>True</td>
    </tr>
    <tr>
      <th>1</th>
      <td>TraesCS1A03G0000400.1</td>
      <td>TraesCS1A03G0000400</td>
      <td>TraesCS1A03G0000400.1</td>
      <td>Chr1A</td>
      <td>+</td>
      <td>70239</td>
      <td>89245</td>
      <td>70239</td>
      <td>89245</td>
      <td>70239</td>
      <td>...</td>
      <td>0</td>
      <td>HC</td>
      <td>TraesCS1A03G0000400</td>
      <td>NaN</td>
      <td>TraesCS1A03G0000400.1</td>
      <td>NaN</td>
      <td>True</td>
      <td>True</td>
      <td>True</td>
      <td>True</td>
    </tr>
    <tr>
      <th>2</th>
      <td>TraesCS1A03G0000600.1</td>
      <td>TraesCS1A03G0000600</td>
      <td>TraesCS1A03G0000600.1</td>
      <td>Chr1A</td>
      <td>+</td>
      <td>95906</td>
      <td>104903</td>
      <td>95906</td>
      <td>104903</td>
      <td>104607</td>
      <td>...</td>
      <td>0</td>
      <td>HC</td>
      <td>TraesCS1A03G0000600</td>
      <td>NaN</td>
      <td>TraesCS1A03G0000600.1</td>
      <td>NaN</td>
      <td>True</td>
      <td>True</td>
      <td>True</td>
      <td>True</td>
    </tr>
    <tr>
      <th>3</th>
      <td>TraesCS1A03G0000800.1</td>
      <td>TraesCS1A03G0000800</td>
      <td>TraesCS1A03G0000800.1</td>
      <td>Chr1A</td>
      <td>+</td>
      <td>102794</td>
      <td>122504</td>
      <td>102794</td>
      <td>122504</td>
      <td>121263</td>
      <td>...</td>
      <td>0</td>
      <td>HC</td>
      <td>TraesCS1A03G0000800</td>
      <td>NaN</td>
      <td>TraesCS1A03G0000800.1</td>
      <td>NaN</td>
      <td>True</td>
      <td>True</td>
      <td>True</td>
      <td>True</td>
    </tr>
    <tr>
      <th>4</th>
      <td>TraesCS1A03G0001000.1</td>
      <td>TraesCS1A03G0001000</td>
      <td>TraesCS1A03G0001000.1</td>
      <td>Chr1A</td>
      <td>-</td>
      <td>149490</td>
      <td>154559</td>
      <td>149490</td>
      <td>154559</td>
      <td>149490</td>
      <td>...</td>
      <td>0; 2</td>
      <td>HC</td>
      <td>TraesCS1A03G0001000</td>
      <td>NaN</td>
      <td>TraesCS1A03G0001000.1</td>
      <td>NaN</td>
      <td>True</td>
      <td>True</td>
      <td>True</td>
      <td>True</td>
    </tr>
  </tbody>
</table>
<p>5 rows × 24 columns</p>
</div>


    
    Processing MSV000090572 | stored_grain
    
    Saved peptide–protein–gene mapping table: python_outputs\tables\FragPipe_Vincent_MSV000090572_stored-grain_peptide_protein_gene_mapping.csv
    Rows: 30,317
    Mapped pairs: 30,314
    Unmapped pairs: 3
    
    Processing PXD050500 | coleoptile
    
    Saved peptide–protein–gene mapping table: python_outputs\tables\FragPipe_Liu_PXD050500_coleoptile_peptide_protein_gene_mapping.csv
    Rows: 1,850,247
    Mapped pairs: 1,850,136
    Unmapped pairs: 111
    
    Processing PXD050500 | node
    
    Saved peptide–protein–gene mapping table: python_outputs\tables\FragPipe_Liu_PXD050500_node_peptide_protein_gene_mapping.csv
    Rows: 1,887,009
    Mapped pairs: 1,886,922
    Unmapped pairs: 87
    
    Processing PXD050500 | radicle
    
    Saved peptide–protein–gene mapping table: python_outputs\tables\FragPipe_Liu_PXD050500_radicle_peptide_protein_gene_mapping.csv
    Rows: 1,062,587
    Mapped pairs: 1,062,508
    Unmapped pairs: 79
    
    Processing PXD004720 | anther
    
    Saved peptide–protein–gene mapping table: python_outputs\tables\FragPipe_Duncan_PXD004720_anther_peptide_protein_gene_mapping.csv
    Rows: 164,397
    Mapped pairs: 164,365
    Unmapped pairs: 32
    
    Processing PXD004720 | boot
    
    Saved peptide–protein–gene mapping table: python_outputs\tables\FragPipe_Duncan_PXD004720_boot_peptide_protein_gene_mapping.csv
    Rows: 13,230
    Mapped pairs: 13,227
    Unmapped pairs: 3
    
    Processing PXD004720 | coleoptile
    
    Saved peptide–protein–gene mapping table: python_outputs\tables\FragPipe_Duncan_PXD004720_coleoptile_peptide_protein_gene_mapping.csv
    Rows: 204,518
    Mapped pairs: 204,476
    Unmapped pairs: 42
    
    Processing PXD004720 | embryo
    
    Saved peptide–protein–gene mapping table: python_outputs\tables\FragPipe_Duncan_PXD004720_embryo_peptide_protein_gene_mapping.csv
    Rows: 8,852
    Mapped pairs: 8,852
    Unmapped pairs: 0
    
    Processing PXD004720 | endosperm
    
    Saved peptide–protein–gene mapping table: python_outputs\tables\FragPipe_Duncan_PXD004720_endosperm_peptide_protein_gene_mapping.csv
    Rows: 102,845
    Mapped pairs: 102,830
    Unmapped pairs: 15
    
    Processing PXD004720 | glume
    
    Saved peptide–protein–gene mapping table: python_outputs\tables\FragPipe_Duncan_PXD004720_glume_peptide_protein_gene_mapping.csv
    Rows: 145,154
    Mapped pairs: 145,133
    Unmapped pairs: 21
    
    Processing PXD004720 | grain-zadoks-70
    
    Saved peptide–protein–gene mapping table: python_outputs\tables\FragPipe_Duncan_PXD004720_grain-zadoks-70_peptide_protein_gene_mapping.csv
    Rows: 126,227
    Mapped pairs: 126,195
    Unmapped pairs: 32
    
    Processing PXD004720 | grain-zadoks-71
    
    Saved peptide–protein–gene mapping table: python_outputs\tables\FragPipe_Duncan_PXD004720_grain-zadoks-71_peptide_protein_gene_mapping.csv
    Rows: 179,751
    Mapped pairs: 179,725
    Unmapped pairs: 26
    
    Processing PXD004720 | grain-zadoks-75
    
    Saved peptide–protein–gene mapping table: python_outputs\tables\FragPipe_Duncan_PXD004720_grain-zadoks-75_peptide_protein_gene_mapping.csv
    Rows: 132,719
    Mapped pairs: 132,701
    Unmapped pairs: 18
    
    Processing PXD004720 | grain-zadoks-83
    
    Saved peptide–protein–gene mapping table: python_outputs\tables\FragPipe_Duncan_PXD004720_grain-zadoks-83_peptide_protein_gene_mapping.csv
    Rows: 113,236
    Mapped pairs: 113,203
    Unmapped pairs: 33
    
    Processing PXD004720 | grain-zadoks-87
    
    Saved peptide–protein–gene mapping table: python_outputs\tables\FragPipe_Duncan_PXD004720_grain-zadoks-87_peptide_protein_gene_mapping.csv
    Rows: 111,819
    Mapped pairs: 111,792
    Unmapped pairs: 27
    
    Processing PXD004720 | leaf-flag-mature
    
    Saved peptide–protein–gene mapping table: python_outputs\tables\FragPipe_Duncan_PXD004720_leaf-flag-mature_peptide_protein_gene_mapping.csv
    Rows: 145,650
    Mapped pairs: 145,632
    Unmapped pairs: 18
    
    Processing PXD004720 | leaf-flag-senescing
    
    Saved peptide–protein–gene mapping table: python_outputs\tables\FragPipe_Duncan_PXD004720_leaf-flag-senescing_peptide_protein_gene_mapping.csv
    Rows: 46,206
    Mapped pairs: 46,200
    Unmapped pairs: 6
    
    Processing PXD004720 | leaf-flag-young
    
    Saved peptide–protein–gene mapping table: python_outputs\tables\FragPipe_Duncan_PXD004720_leaf-flag-young_peptide_protein_gene_mapping.csv
    Rows: 120,833
    Mapped pairs: 120,821
    Unmapped pairs: 12
    
    Processing PXD004720 | lemma
    
    Saved peptide–protein–gene mapping table: python_outputs\tables\FragPipe_Duncan_PXD004720_lemma_peptide_protein_gene_mapping.csv
    Rows: 147,650
    Mapped pairs: 147,618
    Unmapped pairs: 32
    
    Processing PXD004720 | node
    
    Saved peptide–protein–gene mapping table: python_outputs\tables\FragPipe_Duncan_PXD004720_node_peptide_protein_gene_mapping.csv
    Rows: 99,844
    Mapped pairs: 99,823
    Unmapped pairs: 21
    
    Processing PXD004720 | node_secretion
    
    Saved peptide–protein–gene mapping table: python_outputs\tables\FragPipe_Duncan_PXD004720_node-secretion_peptide_protein_gene_mapping.csv
    Rows: 169,705
    Mapped pairs: 169,675
    Unmapped pairs: 30
    
    Processing PXD004720 | palea
    
    Saved peptide–protein–gene mapping table: python_outputs\tables\FragPipe_Duncan_PXD004720_palea_peptide_protein_gene_mapping.csv
    Rows: 116,960
    Mapped pairs: 116,936
    Unmapped pairs: 24
    
    Processing PXD004720 | pericarp
    
    Saved peptide–protein–gene mapping table: python_outputs\tables\FragPipe_Duncan_PXD004720_pericarp_peptide_protein_gene_mapping.csv
    Rows: 131,245
    Mapped pairs: 131,215
    Unmapped pairs: 30
    
    Processing PXD004720 | pollen
    
    Saved peptide–protein–gene mapping table: python_outputs\tables\FragPipe_Duncan_PXD004720_pollen_peptide_protein_gene_mapping.csv
    Rows: 74,722
    Mapped pairs: 74,698
    Unmapped pairs: 24
    
    Processing PXD004720 | rachilla
    
    Saved peptide–protein–gene mapping table: python_outputs\tables\FragPipe_Duncan_PXD004720_rachilla_peptide_protein_gene_mapping.csv
    Rows: 160,234
    Mapped pairs: 160,204
    Unmapped pairs: 30
    
    Processing PXD004720 | radicle
    
    Saved peptide–protein–gene mapping table: python_outputs\tables\FragPipe_Duncan_PXD004720_radicle_peptide_protein_gene_mapping.csv
    Rows: 194,730
    Mapped pairs: 194,694
    Unmapped pairs: 36
    
    Processing PXD004720 | root-mature
    
    Saved peptide–protein–gene mapping table: python_outputs\tables\FragPipe_Duncan_PXD004720_root-mature_peptide_protein_gene_mapping.csv
    Rows: 97,211
    Mapped pairs: 97,188
    Unmapped pairs: 23
    
    Processing PXD004720 | root-secretion
    
    Saved peptide–protein–gene mapping table: python_outputs\tables\FragPipe_Duncan_PXD004720_root-secretion_peptide_protein_gene_mapping.csv
    Rows: 100,654
    Mapped pairs: 100,639
    Unmapped pairs: 15
    
    Processing PXD004720 | root-tip
    
    Saved peptide–protein–gene mapping table: python_outputs\tables\FragPipe_Duncan_PXD004720_root-tip_peptide_protein_gene_mapping.csv
    Rows: 191,973
    Mapped pairs: 191,943
    Unmapped pairs: 30
    
    Processing PXD004720 | root-vasculature
    
    Saved peptide–protein–gene mapping table: python_outputs\tables\FragPipe_Duncan_PXD004720_root-vasculature_peptide_protein_gene_mapping.csv
    Rows: 111,846
    Mapped pairs: 111,831
    Unmapped pairs: 15
    
    Processing PXD004720 | spike-immature
    
    Saved peptide–protein–gene mapping table: python_outputs\tables\FragPipe_Duncan_PXD004720_spike-immature_peptide_protein_gene_mapping.csv
    Rows: 188,868
    Mapped pairs: 188,826
    Unmapped pairs: 42
    
    Processing PXD004720 | stem
    
    Saved peptide–protein–gene mapping table: python_outputs\tables\FragPipe_Duncan_PXD004720_stem_peptide_protein_gene_mapping.csv
    Rows: 60,749
    Mapped pairs: 60,734
    Unmapped pairs: 15
    
    Step 8 summary saved: python_outputs\tables\wheat_peptide_protein_gene_mapping_summary_step8.csv
    Summary rows: 32
    


<div>
<style scoped>
    .dataframe tbody tr th:only-of-type {
        vertical-align: middle;
    }

    .dataframe tbody tr th {
        vertical-align: top;
    }

    .dataframe thead th {
        text-align: right;
    }
</style>
<table border="1" class="dataframe">
  <thead>
    <tr style="text-align: right;">
      <th></th>
      <th>Source</th>
      <th>Species</th>
      <th>Tissue</th>
      <th>Batch</th>
      <th>Gene_mapping_file</th>
      <th>Peptide_protein_pairs</th>
      <th>Mapped_peptide_protein_pairs</th>
      <th>Unmapped_peptide_protein_pairs</th>
      <th>Mapping_rate_percent</th>
      <th>Unique_peptides</th>
      <th>Unique_proteins</th>
      <th>Mapped_unique_proteins</th>
      <th>Unmapped_unique_proteins</th>
      <th>Unique_gene_models</th>
      <th>Unique_transcripts</th>
    </tr>
  </thead>
  <tbody>
    <tr>
      <th>0</th>
      <td>MSV000090572</td>
      <td>bread wheat</td>
      <td>stored_grain</td>
      <td>single</td>
      <td>FragPipe_Vincent_MSV000090572_stored-grain_pep...</td>
      <td>30317</td>
      <td>30314</td>
      <td>3</td>
      <td>99.99</td>
      <td>9329</td>
      <td>17481</td>
      <td>17478</td>
      <td>3</td>
      <td>&lt;NA&gt;</td>
      <td>17478</td>
    </tr>
    <tr>
      <th>1</th>
      <td>PXD050500</td>
      <td>bread wheat</td>
      <td>coleoptile</td>
      <td>single</td>
      <td>FragPipe_Liu_PXD050500_coleoptile_peptide_prot...</td>
      <td>1850247</td>
      <td>1850136</td>
      <td>111</td>
      <td>99.99</td>
      <td>582775</td>
      <td>239514</td>
      <td>239461</td>
      <td>53</td>
      <td>&lt;NA&gt;</td>
      <td>239461</td>
    </tr>
    <tr>
      <th>2</th>
      <td>PXD050500</td>
      <td>bread wheat</td>
      <td>node</td>
      <td>single</td>
      <td>FragPipe_Liu_PXD050500_node_peptide_protein_ge...</td>
      <td>1887009</td>
      <td>1886922</td>
      <td>87</td>
      <td>100.00</td>
      <td>595954</td>
      <td>242842</td>
      <td>242813</td>
      <td>29</td>
      <td>&lt;NA&gt;</td>
      <td>242813</td>
    </tr>
    <tr>
      <th>3</th>
      <td>PXD050500</td>
      <td>bread wheat</td>
      <td>radicle</td>
      <td>single</td>
      <td>FragPipe_Liu_PXD050500_radicle_peptide_protein...</td>
      <td>1062587</td>
      <td>1062508</td>
      <td>79</td>
      <td>99.99</td>
      <td>333058</td>
      <td>206049</td>
      <td>206016</td>
      <td>33</td>
      <td>&lt;NA&gt;</td>
      <td>206016</td>
    </tr>
    <tr>
      <th>4</th>
      <td>PXD004720</td>
      <td>bread wheat</td>
      <td>anther</td>
      <td>single</td>
      <td>FragPipe_Duncan_PXD004720_anther_peptide_prote...</td>
      <td>164397</td>
      <td>164365</td>
      <td>32</td>
      <td>99.98</td>
      <td>34098</td>
      <td>36839</td>
      <td>36826</td>
      <td>13</td>
      <td>&lt;NA&gt;</td>
      <td>36826</td>
    </tr>
    <tr>
      <th>5</th>
      <td>PXD004720</td>
      <td>bread wheat</td>
      <td>boot</td>
      <td>single</td>
      <td>FragPipe_Duncan_PXD004720_boot_peptide_protein...</td>
      <td>13230</td>
      <td>13227</td>
      <td>3</td>
      <td>99.98</td>
      <td>3643</td>
      <td>9211</td>
      <td>9208</td>
      <td>3</td>
      <td>&lt;NA&gt;</td>
      <td>9208</td>
    </tr>
    <tr>
      <th>6</th>
      <td>PXD004720</td>
      <td>bread wheat</td>
      <td>coleoptile</td>
      <td>single</td>
      <td>FragPipe_Duncan_PXD004720_coleoptile_peptide_p...</td>
      <td>204518</td>
      <td>204476</td>
      <td>42</td>
      <td>99.98</td>
      <td>41025</td>
      <td>46513</td>
      <td>46496</td>
      <td>17</td>
      <td>&lt;NA&gt;</td>
      <td>46496</td>
    </tr>
    <tr>
      <th>7</th>
      <td>PXD004720</td>
      <td>bread wheat</td>
      <td>embryo</td>
      <td>single</td>
      <td>FragPipe_Duncan_PXD004720_embryo_peptide_prote...</td>
      <td>8852</td>
      <td>8852</td>
      <td>0</td>
      <td>100.00</td>
      <td>2867</td>
      <td>7220</td>
      <td>7220</td>
      <td>0</td>
      <td>&lt;NA&gt;</td>
      <td>7220</td>
    </tr>
    <tr>
      <th>8</th>
      <td>PXD004720</td>
      <td>bread wheat</td>
      <td>endosperm</td>
      <td>single</td>
      <td>FragPipe_Duncan_PXD004720_endosperm_peptide_pr...</td>
      <td>102845</td>
      <td>102830</td>
      <td>15</td>
      <td>99.99</td>
      <td>20081</td>
      <td>27461</td>
      <td>27454</td>
      <td>7</td>
      <td>&lt;NA&gt;</td>
      <td>27454</td>
    </tr>
    <tr>
      <th>9</th>
      <td>PXD004720</td>
      <td>bread wheat</td>
      <td>glume</td>
      <td>single</td>
      <td>FragPipe_Duncan_PXD004720_glume_peptide_protei...</td>
      <td>145154</td>
      <td>145133</td>
      <td>21</td>
      <td>99.99</td>
      <td>28648</td>
      <td>35551</td>
      <td>35540</td>
      <td>11</td>
      <td>&lt;NA&gt;</td>
      <td>35540</td>
    </tr>
    <tr>
      <th>10</th>
      <td>PXD004720</td>
      <td>bread wheat</td>
      <td>grain-zadoks-70</td>
      <td>single</td>
      <td>FragPipe_Duncan_PXD004720_grain-zadoks-70_pept...</td>
      <td>126227</td>
      <td>126195</td>
      <td>32</td>
      <td>99.97</td>
      <td>26196</td>
      <td>35549</td>
      <td>35532</td>
      <td>17</td>
      <td>&lt;NA&gt;</td>
      <td>35532</td>
    </tr>
    <tr>
      <th>11</th>
      <td>PXD004720</td>
      <td>bread wheat</td>
      <td>grain-zadoks-71</td>
      <td>single</td>
      <td>FragPipe_Duncan_PXD004720_grain-zadoks-71_pept...</td>
      <td>179751</td>
      <td>179725</td>
      <td>26</td>
      <td>99.99</td>
      <td>35960</td>
      <td>47089</td>
      <td>47076</td>
      <td>13</td>
      <td>&lt;NA&gt;</td>
      <td>47076</td>
    </tr>
    <tr>
      <th>12</th>
      <td>PXD004720</td>
      <td>bread wheat</td>
      <td>grain-zadoks-75</td>
      <td>single</td>
      <td>FragPipe_Duncan_PXD004720_grain-zadoks-75_pept...</td>
      <td>132719</td>
      <td>132701</td>
      <td>18</td>
      <td>99.99</td>
      <td>27463</td>
      <td>41042</td>
      <td>41029</td>
      <td>13</td>
      <td>&lt;NA&gt;</td>
      <td>41029</td>
    </tr>
    <tr>
      <th>13</th>
      <td>PXD004720</td>
      <td>bread wheat</td>
      <td>grain-zadoks-83</td>
      <td>single</td>
      <td>FragPipe_Duncan_PXD004720_grain-zadoks-83_pept...</td>
      <td>113236</td>
      <td>113203</td>
      <td>33</td>
      <td>99.97</td>
      <td>23990</td>
      <td>36654</td>
      <td>36635</td>
      <td>19</td>
      <td>&lt;NA&gt;</td>
      <td>36635</td>
    </tr>
    <tr>
      <th>14</th>
      <td>PXD004720</td>
      <td>bread wheat</td>
      <td>grain-zadoks-87</td>
      <td>single</td>
      <td>FragPipe_Duncan_PXD004720_grain-zadoks-87_pept...</td>
      <td>111819</td>
      <td>111792</td>
      <td>27</td>
      <td>99.98</td>
      <td>24636</td>
      <td>34889</td>
      <td>34874</td>
      <td>15</td>
      <td>&lt;NA&gt;</td>
      <td>34874</td>
    </tr>
    <tr>
      <th>15</th>
      <td>PXD004720</td>
      <td>bread wheat</td>
      <td>leaf-flag-mature</td>
      <td>single</td>
      <td>FragPipe_Duncan_PXD004720_leaf-flag-mature_pep...</td>
      <td>145650</td>
      <td>145632</td>
      <td>18</td>
      <td>99.99</td>
      <td>29840</td>
      <td>37963</td>
      <td>37954</td>
      <td>9</td>
      <td>&lt;NA&gt;</td>
      <td>37954</td>
    </tr>
    <tr>
      <th>16</th>
      <td>PXD004720</td>
      <td>bread wheat</td>
      <td>leaf-flag-senescing</td>
      <td>single</td>
      <td>FragPipe_Duncan_PXD004720_leaf-flag-senescing_...</td>
      <td>46206</td>
      <td>46200</td>
      <td>6</td>
      <td>99.99</td>
      <td>11354</td>
      <td>25701</td>
      <td>25696</td>
      <td>5</td>
      <td>&lt;NA&gt;</td>
      <td>25696</td>
    </tr>
    <tr>
      <th>17</th>
      <td>PXD004720</td>
      <td>bread wheat</td>
      <td>leaf-flag-young</td>
      <td>single</td>
      <td>FragPipe_Duncan_PXD004720_leaf-flag-young_pept...</td>
      <td>120833</td>
      <td>120821</td>
      <td>12</td>
      <td>99.99</td>
      <td>23373</td>
      <td>32629</td>
      <td>32620</td>
      <td>9</td>
      <td>&lt;NA&gt;</td>
      <td>32620</td>
    </tr>
    <tr>
      <th>18</th>
      <td>PXD004720</td>
      <td>bread wheat</td>
      <td>lemma</td>
      <td>single</td>
      <td>FragPipe_Duncan_PXD004720_lemma_peptide_protei...</td>
      <td>147650</td>
      <td>147618</td>
      <td>32</td>
      <td>99.98</td>
      <td>29788</td>
      <td>37946</td>
      <td>37927</td>
      <td>19</td>
      <td>&lt;NA&gt;</td>
      <td>37927</td>
    </tr>
    <tr>
      <th>19</th>
      <td>PXD004720</td>
      <td>bread wheat</td>
      <td>node</td>
      <td>single</td>
      <td>FragPipe_Duncan_PXD004720_node_peptide_protein...</td>
      <td>99844</td>
      <td>99823</td>
      <td>21</td>
      <td>99.98</td>
      <td>21457</td>
      <td>36555</td>
      <td>36542</td>
      <td>13</td>
      <td>&lt;NA&gt;</td>
      <td>36542</td>
    </tr>
    <tr>
      <th>20</th>
      <td>PXD004720</td>
      <td>bread wheat</td>
      <td>node_secretion</td>
      <td>single</td>
      <td>FragPipe_Duncan_PXD004720_node-secretion_pepti...</td>
      <td>169705</td>
      <td>169675</td>
      <td>30</td>
      <td>99.98</td>
      <td>34124</td>
      <td>42371</td>
      <td>42356</td>
      <td>15</td>
      <td>&lt;NA&gt;</td>
      <td>42356</td>
    </tr>
    <tr>
      <th>21</th>
      <td>PXD004720</td>
      <td>bread wheat</td>
      <td>palea</td>
      <td>single</td>
      <td>FragPipe_Duncan_PXD004720_palea_peptide_protei...</td>
      <td>116960</td>
      <td>116936</td>
      <td>24</td>
      <td>99.98</td>
      <td>21707</td>
      <td>27591</td>
      <td>27580</td>
      <td>11</td>
      <td>&lt;NA&gt;</td>
      <td>27580</td>
    </tr>
    <tr>
      <th>22</th>
      <td>PXD004720</td>
      <td>bread wheat</td>
      <td>pericarp</td>
      <td>single</td>
      <td>FragPipe_Duncan_PXD004720_pericarp_peptide_pro...</td>
      <td>131245</td>
      <td>131215</td>
      <td>30</td>
      <td>99.98</td>
      <td>28332</td>
      <td>32676</td>
      <td>32667</td>
      <td>9</td>
      <td>&lt;NA&gt;</td>
      <td>32667</td>
    </tr>
    <tr>
      <th>23</th>
      <td>PXD004720</td>
      <td>bread wheat</td>
      <td>pollen</td>
      <td>single</td>
      <td>FragPipe_Duncan_PXD004720_pollen_peptide_prote...</td>
      <td>74722</td>
      <td>74698</td>
      <td>24</td>
      <td>99.97</td>
      <td>13502</td>
      <td>19942</td>
      <td>19931</td>
      <td>11</td>
      <td>&lt;NA&gt;</td>
      <td>19931</td>
    </tr>
    <tr>
      <th>24</th>
      <td>PXD004720</td>
      <td>bread wheat</td>
      <td>rachilla</td>
      <td>single</td>
      <td>FragPipe_Duncan_PXD004720_rachilla_peptide_pro...</td>
      <td>160234</td>
      <td>160204</td>
      <td>30</td>
      <td>99.98</td>
      <td>31145</td>
      <td>37219</td>
      <td>37204</td>
      <td>15</td>
      <td>&lt;NA&gt;</td>
      <td>37204</td>
    </tr>
    <tr>
      <th>25</th>
      <td>PXD004720</td>
      <td>bread wheat</td>
      <td>radicle</td>
      <td>single</td>
      <td>FragPipe_Duncan_PXD004720_radicle_peptide_prot...</td>
      <td>194730</td>
      <td>194694</td>
      <td>36</td>
      <td>99.98</td>
      <td>39643</td>
      <td>42976</td>
      <td>42965</td>
      <td>11</td>
      <td>&lt;NA&gt;</td>
      <td>42965</td>
    </tr>
    <tr>
      <th>26</th>
      <td>PXD004720</td>
      <td>bread wheat</td>
      <td>root-mature</td>
      <td>single</td>
      <td>FragPipe_Duncan_PXD004720_root-mature_peptide_...</td>
      <td>97211</td>
      <td>97188</td>
      <td>23</td>
      <td>99.98</td>
      <td>21350</td>
      <td>39165</td>
      <td>39156</td>
      <td>9</td>
      <td>&lt;NA&gt;</td>
      <td>39156</td>
    </tr>
    <tr>
      <th>27</th>
      <td>PXD004720</td>
      <td>bread wheat</td>
      <td>root-secretion</td>
      <td>single</td>
      <td>FragPipe_Duncan_PXD004720_root-secretion_pepti...</td>
      <td>100654</td>
      <td>100639</td>
      <td>15</td>
      <td>99.99</td>
      <td>21284</td>
      <td>32049</td>
      <td>32046</td>
      <td>3</td>
      <td>&lt;NA&gt;</td>
      <td>32046</td>
    </tr>
    <tr>
      <th>28</th>
      <td>PXD004720</td>
      <td>bread wheat</td>
      <td>root-tip</td>
      <td>single</td>
      <td>FragPipe_Duncan_PXD004720_root-tip_peptide_pro...</td>
      <td>191973</td>
      <td>191943</td>
      <td>30</td>
      <td>99.98</td>
      <td>38634</td>
      <td>39785</td>
      <td>39774</td>
      <td>11</td>
      <td>&lt;NA&gt;</td>
      <td>39774</td>
    </tr>
    <tr>
      <th>29</th>
      <td>PXD004720</td>
      <td>bread wheat</td>
      <td>root-vasculature</td>
      <td>single</td>
      <td>FragPipe_Duncan_PXD004720_root-vasculature_pep...</td>
      <td>111846</td>
      <td>111831</td>
      <td>15</td>
      <td>99.99</td>
      <td>20815</td>
      <td>30631</td>
      <td>30626</td>
      <td>5</td>
      <td>&lt;NA&gt;</td>
      <td>30626</td>
    </tr>
    <tr>
      <th>30</th>
      <td>PXD004720</td>
      <td>bread wheat</td>
      <td>spike-immature</td>
      <td>single</td>
      <td>FragPipe_Duncan_PXD004720_spike-immature_pepti...</td>
      <td>188868</td>
      <td>188826</td>
      <td>42</td>
      <td>99.98</td>
      <td>36370</td>
      <td>40220</td>
      <td>40207</td>
      <td>13</td>
      <td>&lt;NA&gt;</td>
      <td>40207</td>
    </tr>
    <tr>
      <th>31</th>
      <td>PXD004720</td>
      <td>bread wheat</td>
      <td>stem</td>
      <td>single</td>
      <td>FragPipe_Duncan_PXD004720_stem_peptide_protein...</td>
      <td>60749</td>
      <td>60734</td>
      <td>15</td>
      <td>99.98</td>
      <td>14338</td>
      <td>29417</td>
      <td>29410</td>
      <td>7</td>
      <td>&lt;NA&gt;</td>
      <td>29410</td>
    </tr>
  </tbody>
</table>
</div>


# Step 9 — Project Identified Peptides onto Wheat Genomic Coordinates

This step projects identified non-contaminant peptides from protein coordinates onto wheat genomic coordinates using the IWGSC RefSeq v2.1 genome annotation.

The aim is to convert peptide-level proteomics evidence into genome browser-compatible coordinate information for downstream BED file generation and JBrowse visualisation.

To improve computational efficiency on standard desktop hardware, CDS annotation features were pre-indexed into transcript-level dictionaries prior to peptide projection. This substantially reduced repeated scanning of the large wheat GFF3 annotation table (>2.8 million rows).

---

## Input files

### Peptide–protein–gene mapping tables from Step 8

One file per tissue:

`FragPipe_FirstAuthor_Source_Tissue-Raw-Code_peptide_protein_gene_mapping.csv`

### Protein FASTA database

`iwgsc_refseqv2.1_annotation_200916_HC_LC_pep_cRAP_with_DECOY.fasta`

### Parsed GFF3 feature table from Step 5

`wheat_gff3_parsed_features_HC_LC.csv`

---

## Projection strategy

Genomic coding coordinate vectors were reconstructed directly from ordered CDS intervals extracted from the GFF3 annotation. CDS phase values were retained from the annotation for reference but were not used to trim CDS nucleotide coordinates during peptide projection, because CDS features already define coding sequence boundaries. This ensured continuous exon-aware reconstruction of transcript coding coordinates prior to peptide genomic projection.

For each peptide–protein–transcript entry, the workflow:

1. Locates the peptide amino acid sequence within the corresponding protein sequence.
2. Converts the peptide amino acid interval into coding nucleotide positions.
3. Retrieves the CDS blocks associated with the corresponding transcript.
4. Projects coding nucleotide positions onto genomic coordinates.
5. Collapses projected nucleotide positions into genomic blocks.
6. Generates an intron-aware peptide display label in which exon-spanning junctions are explicitly shown using dash characters.

This allows peptides spanning CDS junctions to be represented as multi-block genomic features and labelled in JBrowse with an explicit visual indication of intron-spanning structure.

---

## Intron-aware peptide labels

For peptides projected across more than one CDS block, an additional display field is generated:

`Peptide_intron_gapped`

This field preserves the original peptide sequence but inserts dash characters at positions where the peptide crosses an intron.

Example:

`AFVVPGLTDADGVGYVAQ--------------------------------------GEGVLTVIENGEKR`

This makes exon-spanning peptides easier to interpret in Apollo/JBrowse, particularly for visualising peptide-supported splice structures across CDS boundaries.

For peptides contained within a single CDS block, `Peptide_intron_gapped` is identical to the original peptide sequence.

---

## Coordinate systems

Both biological and BED-compatible coordinate systems were retained.

| Coordinate field | Description |
|---|---|
| `Genomic_start_1based` | Genomic start coordinate using 1-based inclusive coordinates |
| `Genomic_end_1based` | Genomic end coordinate using 1-based inclusive coordinates |
| `BED_start_0based` | BED-compatible 0-based start coordinate |
| `BED_end_0based_exclusive` | BED-compatible end coordinate |
| `BED_block_count` | Number of genomic blocks covered by the peptide |
| `BED_block_sizes` | BED12-compatible block sizes |
| `BED_block_starts` | BED12-compatible block starts relative to feature start |
| `Peptide_intron_gapped` | Peptide label with dash characters inserted across intron-spanning junctions |

---

## Projection status

A projection status column was added to support quality control and computational validation of peptide genomic projections.

| Status | Description |
|---|---|
| `projected` | Peptide was successfully projected onto genomic coordinates |
| `protein_sequence_not_found` | Protein accession was not found in the FASTA database |
| `peptide_not_found_in_protein` | Peptide sequence could not be located in the protein sequence |
| `cds_blocks_not_found` | CDS features could not be found for the transcript |
| `incomplete_coding_projection` | Peptide coding interval could not be fully projected |

Only successfully projected rows are used for BED file generation in the next step.

---

## Output files

One peptide genome projection table was exported per tissue.

Example output:

`FragPipe_Duncan_PXD004720_anther_peptide_genome_projection.csv`

Output directory:

`python_outputs/tables/`

---

## Summary metrics

A Step 9 summary table was generated across all tissues.

| Metric | Description |
|---|---|
| `Peptide_protein_gene_rows` | Number of mapped peptide–protein–gene rows considered |
| `Projected_rows` | Number of rows successfully projected onto genomic coordinates |
| `Unprojected_rows` | Number of rows not successfully projected |
| `Projection_rate_percent` | Percentage of rows successfully projected |
| `Unique_projected_peptides` | Number of unique projected peptide sequences |
| `Unique_projected_proteins` | Number of unique projected proteins |
| `Unique_projected_gene_models` | Number of unique projected gene models |
| `Peptides_crossing_CDS_blocks` | Number of projected peptide rows spanning more than one CDS block |
| `Peptides_with_intron_gapped_label` | Number of projected peptide rows with dash-containing intron-aware labels |

Output summary file:

`wheat_peptide_genome_projection_summary_step9.csv`

Projection accuracy was subsequently validated computationally (Step 9B) by reconstructing genomic coding sequences from projected peptide coordinates, translating the reconstructed sequences, and comparing them against the original peptide sequences. Validation of 10,000 randomly sampled projected peptides demonstrated >98% peptide reconstruction concordance after I/L normalisation, supporting the accuracy of exon-aware coordinate reconstruction, strand handling, and genomic peptide projection.

The projected genomic coordinate tables are used in the next step to generate BED6 and BED12 files for visualisation in Apollo/JBrowse.


```python
# ============================================================
# Step 9 — Peptide-to-genome projection with intron-gapped labels (takes 60 min)
# ============================================================

import pandas as pd
from pathlib import Path

# -----------------------------
# 1. Input / output paths
# -----------------------------
fragpipe_dir = Path("FragPipe_results")
tables_dir = Path("python_outputs/tables")

manifest_file = fragpipe_dir / "wheat_tissues_FragPipe-result-manifest_2026-05-11.csv"
fasta_file = Path("protein_database/iwgsc_refseqv2.1_annotation_200916_HC_LC_pep_cRAP_with_DECOY.fasta")
gff3_features_file = tables_dir / "wheat_gff3_parsed_features_HC_LC.csv"

step9_summary_out = tables_dir / "wheat_peptide_genome_projection_summary_step9.csv"

manifest = pd.read_csv(manifest_file, encoding="utf-8-sig")


# -----------------------------
# 2. FASTA reader
# -----------------------------
def read_fasta_sequences(fasta_path):
    sequences = {}
    current_id = None
    current_seq = []

    with open(fasta_path, "r", encoding="utf-8", errors="ignore") as handle:
        for line in handle:
            line = line.strip()

            if not line:
                continue

            if line.startswith(">"):
                if current_id is not None:
                    sequences[current_id] = "".join(current_seq)

                current_id = line[1:].split()[0]
                current_seq = []
            else:
                current_seq.append(line)

        if current_id is not None:
            sequences[current_id] = "".join(current_seq)

    return sequences


# -----------------------------
# 3. Build CDS dictionary once
# -----------------------------
def build_cds_dictionary(gff3_features_file):
    usecols = [
        "SeqID",
        "FeatureType",
        "Start",
        "End",
        "Strand",
        "Phase",
        "Parent"
    ]

    gff = pd.read_csv(
        gff3_features_file,
        usecols=usecols,
        low_memory=False
    )

    cds = gff[gff["FeatureType"].astype(str).str.lower() == "cds"].copy()

    cds["Start"] = cds["Start"].astype(int)
    cds["End"] = cds["End"].astype(int)
    cds["Phase"] = pd.to_numeric(cds["Phase"], errors="coerce").fillna(0).astype(int)

    cds_dict = {}

    for transcript_id, group in cds.groupby("Parent", sort=False):

        strand = group["Strand"].iloc[0]

        if strand == "+":
            group = group.sort_values(["Start", "End"])
        else:
            group = group.sort_values(["End", "Start"], ascending=[False, False])

        cds_dict[transcript_id] = group[[
            "SeqID",
            "Start",
            "End",
            "Strand",
            "Phase"
        ]].to_dict("records")

    return cds_dict


# -----------------------------
# 4. Projection helpers
# -----------------------------
def locate_peptide_in_protein(peptide, protein_sequence):
    peptide_clean = str(peptide).replace("I", "L")
    protein_clean = str(protein_sequence).replace("I", "L")

    start_0based = protein_clean.find(peptide_clean)

    if start_0based == -1:
        return None, None, 0

    occurrence_count = protein_clean.count(peptide_clean)

    aa_start = start_0based + 1
    aa_end = aa_start + len(peptide_clean) - 1

    return aa_start, aa_end, occurrence_count


def build_coding_position_vector_from_blocks(cds_blocks):
    """
    Build transcript-ordered genomic coding positions from CDS blocks.

    Important:
    The GFF3 Phase field is NOT used to trim CDS coordinates here.
    CDS features already define the coding sequence. Phase describes
    codon continuity across CDS features, but bases should not be removed
    when reconstructing peptide genomic positions.
    """
    coding_positions = []

    for block in cds_blocks:
        start = int(block["Start"])
        end = int(block["End"])
        strand = block["Strand"]

        if strand == "+":
            coding_positions.extend(range(start, end + 1))
        else:
            coding_positions.extend(range(end, start - 1, -1))

    return coding_positions


def collapse_positions_to_blocks(genomic_positions):
    if len(genomic_positions) == 0:
        return []

    sorted_positions = sorted(genomic_positions)

    blocks = []
    block_start = sorted_positions[0]
    previous = sorted_positions[0]

    for pos in sorted_positions[1:]:
        if pos == previous + 1:
            previous = pos
        else:
            blocks.append((block_start, previous))
            block_start = pos
            previous = pos

    blocks.append((block_start, previous))

    return blocks


def build_intron_gapped_peptide_from_nt_positions(
    peptide,
    peptide_nt_positions,
    exact_dash_count=True,
    max_dashes=80
):
    """
    Builds a display peptide sequence with dashes inserted where the peptide
    crosses introns.

    The peptide split is based on the actual nucleotide positions contributing
    to each amino acid codon, so the exon boundary placement is exact at
    amino-acid resolution.

    If exact_dash_count=True:
        number of dashes = intron length in nucleotides.
    If exact_dash_count=False:
        dashes are capped to max_dashes for easier JBrowse display.
    """

    peptide = str(peptide)

    if len(peptide_nt_positions) != len(peptide) * 3:
        return peptide

    parts = []

    for aa_index, aa in enumerate(peptide):
        codon_positions = peptide_nt_positions[aa_index * 3:(aa_index + 1) * 3]
        parts.append(aa)

        if aa_index == len(peptide) - 1:
            continue

        next_codon_positions = peptide_nt_positions[(aa_index + 1) * 3:(aa_index + 2) * 3]

        current_genomic_max = max(codon_positions)
        current_genomic_min = min(codon_positions)
        next_genomic_max = max(next_codon_positions)
        next_genomic_min = min(next_codon_positions)

        # Consecutive codons in the same exon are adjacent in genomic space.
        same_exon_forward = next_genomic_min == current_genomic_max + 1
        same_exon_reverse = current_genomic_min == next_genomic_max + 1

        if not (same_exon_forward or same_exon_reverse):

            if next_genomic_min > current_genomic_max:
                intron_nt_len = next_genomic_min - current_genomic_max - 1
            else:
                intron_nt_len = current_genomic_min - next_genomic_max - 1

            if exact_dash_count:
                dash_n = max(1, int(intron_nt_len))
            else:
                dash_n = min(max(3, int(intron_nt_len)), max_dashes)

            parts.append("-" * dash_n)

    return "".join(parts)


# -----------------------------
# 5. Project one tissue
# -----------------------------
def project_one_tissue_peptides_to_genome_fast(source, tissue_raw_code, protein_sequences, cds_dict):

    match = manifest[
        (manifest["Source"] == source) &
        (manifest["Tissue-Raw-Code"] == tissue_raw_code)
    ]

    if match.empty:
        raise ValueError(f"No manifest entry found for {source} | {tissue_raw_code}")

    row = match.iloc[0]

    step8_filename = row["FragPipe-Output-Peptide"].replace(
        "_peptide.tsv",
        "_peptide_protein_gene_mapping.csv"
    )

    step8_path = tables_dir / step8_filename

    mapped = pd.read_csv(step8_path, low_memory=False)

    mapped = mapped[
        mapped["Gene_model_mapping_status"] == "mapped"
    ].copy()

    records = []

    for peptide_row in mapped.itertuples(index=False):

        row_dict = peptide_row._asdict()

        peptide = row_dict["Peptide"]
        protein_id = row_dict["ProteinID"]
        transcript_id = row_dict["TranscriptID"]

        projection_status = "projected"
        peptide_intron_gapped = str(peptide)
        peptide_intron_gapped_compact = str(peptide)

        protein_sequence = protein_sequences.get(protein_id)

        if protein_sequence is None:
            projection_status = "protein_sequence_not_found"
            aa_start = pd.NA
            aa_end = pd.NA
            occurrence_count = pd.NA
            chromosome = pd.NA
            strand = pd.NA
            peptide_blocks = []

        else:
            aa_start, aa_end, occurrence_count = locate_peptide_in_protein(
                peptide,
                protein_sequence
            )

            if aa_start is None:
                projection_status = "peptide_not_found_in_protein"
                aa_start = pd.NA
                aa_end = pd.NA
                chromosome = pd.NA
                strand = pd.NA
                peptide_blocks = []

            else:
                cds_blocks = cds_dict.get(transcript_id)

                if cds_blocks is None:
                    projection_status = "cds_blocks_not_found"
                    chromosome = pd.NA
                    strand = pd.NA
                    peptide_blocks = []

                else:
                    coding_positions = build_coding_position_vector_from_blocks(cds_blocks)

                    nt_start = (int(aa_start) - 1) * 3
                    nt_end = int(aa_end) * 3

                    peptide_nt_positions = coding_positions[nt_start:nt_end]

                    chromosome = cds_blocks[0]["SeqID"]
                    strand = cds_blocks[0]["Strand"]

                    if len(peptide_nt_positions) != len(str(peptide)) * 3:
                        projection_status = "incomplete_coding_projection"
                        peptide_blocks = []
                    else:
                        peptide_blocks = collapse_positions_to_blocks(peptide_nt_positions)

                        # Full-resolution label for BED/JBrowse
                        peptide_intron_gapped = build_intron_gapped_peptide_from_nt_positions(
                            peptide=peptide,
                            peptide_nt_positions=peptide_nt_positions,
                            exact_dash_count=True
                        )
                        
                        # Compact label for CSV storage / downstream loading
                        peptide_intron_gapped_compact = build_intron_gapped_peptide_from_nt_positions(
                            peptide=peptide,
                            peptide_nt_positions=peptide_nt_positions,
                            exact_dash_count=False,
                            max_dashes=10
                        )

        if peptide_blocks:
            genomic_start_1based = min(start for start, end in peptide_blocks)
            genomic_end_1based = max(end for start, end in peptide_blocks)

            bed_start = genomic_start_1based - 1
            bed_end = genomic_end_1based

            block_count = len(peptide_blocks)
            block_sizes = ",".join(str(end - start + 1) for start, end in peptide_blocks)
            block_starts = ",".join(str(start - genomic_start_1based) for start, end in peptide_blocks)

        else:
            genomic_start_1based = pd.NA
            genomic_end_1based = pd.NA
            bed_start = pd.NA
            bed_end = pd.NA
            block_count = 0
            block_sizes = ""
            block_starts = ""

        row_dict.update({
            "AA_start": aa_start,
            "AA_end": aa_end,
            "Peptide_length_AA": len(str(peptide)),
            "Peptide_occurrences_in_protein": occurrence_count,
            "Peptide_intron_gapped": peptide_intron_gapped,
            "Peptide_intron_gapped_compact": peptide_intron_gapped_compact,
            "Chromosome": chromosome,
            "Strand": strand,
            "Genomic_start_1based": genomic_start_1based,
            "Genomic_end_1based": genomic_end_1based,
            "BED_start_0based": bed_start,
            "BED_end_0based_exclusive": bed_end,
            "BED_block_count": block_count,
            "BED_block_sizes": block_sizes,
            "BED_block_starts": block_starts,
            "Projection_status": projection_status
        })

        records.append(row_dict)

    projected = pd.DataFrame(records)

    if "Index" in projected.columns:
        projected = projected.drop(columns=["Index"])

    projected.insert(0, "Index", range(1, len(projected) + 1))

    output_filename = step8_filename.replace(
        "_peptide_protein_gene_mapping.csv",
        "_peptide_genome_projection.csv"
    )

    output_path = tables_dir / output_filename
    projected.to_csv(output_path, index=False)

    print(f"\nSaved: {output_path}")
    print(projected["Projection_status"].value_counts(dropna=False))

    unprojected = projected[
        projected["Projection_status"] != "projected"
    ].copy()

    unprojected_filename = output_filename.replace(
        "_peptide_genome_projection.csv",
        "_unprojected_peptides_for_tblastn.csv"
    )

    unprojected_path = tables_dir / unprojected_filename
    unprojected.to_csv(unprojected_path, index=False)

    print(f"Unprojected peptide table saved: {unprojected_path}")
    print(f"Unprojected rows: {len(unprojected):,}")

    return projected


# -----------------------------
# 6. Pre-load large resources once
# -----------------------------
print("Loading protein FASTA...")
protein_sequences = read_fasta_sequences(fasta_file)
print(f"Protein sequences loaded: {len(protein_sequences):,}")

print("Building CDS dictionary...")
cds_dict = build_cds_dictionary(gff3_features_file)
print(f"Transcript CDS entries loaded: {len(cds_dict):,}")

# # -----------------------------
# # 7-test. Run Step 9 on one tissue only
# # -----------------------------

# test_source = "PXD004720"
# test_tissue = "embryo"

# print(f"\nTEST RUN — Step 9 projection for one tissue only:")
# print(f"Source: {test_source}")
# print(f"Tissue: {test_tissue}")

# test_projection = project_one_tissue_peptides_to_genome_fast(
#     source=test_source,
#     tissue_raw_code=test_tissue,
#     protein_sequences=protein_sequences,
#     cds_dict=cds_dict
# )

# print("\nTest projection completed.")
# print(f"Rows: {len(test_projection):,}")

# display(test_projection.head())
# display(test_projection["Projection_status"].value_counts(dropna=False))

# -----------------------------
# 7. Run all tissues
# -----------------------------
step9_summary_records = []

for _, row in manifest.iterrows():

    source = row["Source"]
    species = row["Species"]
    tissue_raw_code = row["Tissue-Raw-Code"]
    batch = row["Batch"]

    print(f"\nProcessing {source} | {tissue_raw_code}")

    projected = project_one_tissue_peptides_to_genome_fast(
        source=source,
        tissue_raw_code=tissue_raw_code,
        protein_sequences=protein_sequences,
        cds_dict=cds_dict
    )

    output_filename = row["FragPipe-Output-Peptide"].replace(
        "_peptide.tsv",
        "_peptide_genome_projection.csv"
    )

    projected_count = (projected["Projection_status"] == "projected").sum()
    unprojected_count = len(projected) - projected_count

    step9_summary_records.append({
        "Source": source,
        "Species": species,
        "Tissue": tissue_raw_code,
        "Batch": batch,
        "Genome_projection_file": output_filename,
        "Peptide_protein_gene_rows": len(projected),
        "Projected_rows": projected_count,
        "Unprojected_rows": unprojected_count,
        "Projection_rate_percent": round((projected_count / len(projected)) * 100, 2) if len(projected) > 0 else 0,
        "Unique_projected_peptides": projected.loc[
            projected["Projection_status"] == "projected", "Peptide"
        ].nunique(),
        "Unique_projected_proteins": projected.loc[
            projected["Projection_status"] == "projected", "ProteinID"
        ].nunique(),
        "Unique_projected_gene_models": projected.loc[
            projected["Projection_status"] == "projected", "GeneID"
        ].nunique() if "GeneID" in projected.columns else pd.NA,
        "Peptides_crossing_CDS_blocks": (
            projected.loc[
                projected["Projection_status"] == "projected", "BED_block_count"
            ] > 1
        ).sum(),
        "Peptides_with_intron_gapped_label": (
            projected.loc[
                projected["Projection_status"] == "projected", "Peptide_intron_gapped"
            ].astype(str).str.contains("-", regex=False)
        ).sum()
    })

step9_summary = pd.DataFrame(step9_summary_records)

step9_summary.to_csv(step9_summary_out, index=False)

print(f"\nStep 9 summary saved: {step9_summary_out}")
display(step9_summary)

```

    Loading protein FASTA...
    Protein sequences loaded: 592,060
    Building CDS dictionary...
    Transcript CDS entries loaded: 295,914
    
    Processing MSV000090572 | stored_grain
    
    Saved: python_outputs\tables\FragPipe_Vincent_MSV000090572_stored-grain_peptide_genome_projection.csv
    Projection_status
    projected    30314
    Name: count, dtype: int64
    Unprojected peptide table saved: python_outputs\tables\FragPipe_Vincent_MSV000090572_stored-grain_unprojected_peptides_for_tblastn.csv
    Unprojected rows: 0
    
    Processing PXD050500 | coleoptile
    
    Saved: python_outputs\tables\FragPipe_Liu_PXD050500_coleoptile_peptide_genome_projection.csv
    Projection_status
    projected    1850136
    Name: count, dtype: int64
    Unprojected peptide table saved: python_outputs\tables\FragPipe_Liu_PXD050500_coleoptile_unprojected_peptides_for_tblastn.csv
    Unprojected rows: 0
    
    Processing PXD050500 | node
    
    Saved: python_outputs\tables\FragPipe_Liu_PXD050500_node_peptide_genome_projection.csv
    Projection_status
    projected    1886922
    Name: count, dtype: int64
    Unprojected peptide table saved: python_outputs\tables\FragPipe_Liu_PXD050500_node_unprojected_peptides_for_tblastn.csv
    Unprojected rows: 0
    
    Processing PXD050500 | radicle
    
    Saved: python_outputs\tables\FragPipe_Liu_PXD050500_radicle_peptide_genome_projection.csv
    Projection_status
    projected    1062508
    Name: count, dtype: int64
    Unprojected peptide table saved: python_outputs\tables\FragPipe_Liu_PXD050500_radicle_unprojected_peptides_for_tblastn.csv
    Unprojected rows: 0
    
    Processing PXD004720 | anther
    
    Saved: python_outputs\tables\FragPipe_Duncan_PXD004720_anther_peptide_genome_projection.csv
    Projection_status
    projected    164365
    Name: count, dtype: int64
    Unprojected peptide table saved: python_outputs\tables\FragPipe_Duncan_PXD004720_anther_unprojected_peptides_for_tblastn.csv
    Unprojected rows: 0
    
    Processing PXD004720 | boot
    
    Saved: python_outputs\tables\FragPipe_Duncan_PXD004720_boot_peptide_genome_projection.csv
    Projection_status
    projected    13227
    Name: count, dtype: int64
    Unprojected peptide table saved: python_outputs\tables\FragPipe_Duncan_PXD004720_boot_unprojected_peptides_for_tblastn.csv
    Unprojected rows: 0
    
    Processing PXD004720 | coleoptile
    
    Saved: python_outputs\tables\FragPipe_Duncan_PXD004720_coleoptile_peptide_genome_projection.csv
    Projection_status
    projected    204476
    Name: count, dtype: int64
    Unprojected peptide table saved: python_outputs\tables\FragPipe_Duncan_PXD004720_coleoptile_unprojected_peptides_for_tblastn.csv
    Unprojected rows: 0
    
    Processing PXD004720 | embryo
    
    Saved: python_outputs\tables\FragPipe_Duncan_PXD004720_embryo_peptide_genome_projection.csv
    Projection_status
    projected    8852
    Name: count, dtype: int64
    Unprojected peptide table saved: python_outputs\tables\FragPipe_Duncan_PXD004720_embryo_unprojected_peptides_for_tblastn.csv
    Unprojected rows: 0
    
    Processing PXD004720 | endosperm
    
    Saved: python_outputs\tables\FragPipe_Duncan_PXD004720_endosperm_peptide_genome_projection.csv
    Projection_status
    projected    102830
    Name: count, dtype: int64
    Unprojected peptide table saved: python_outputs\tables\FragPipe_Duncan_PXD004720_endosperm_unprojected_peptides_for_tblastn.csv
    Unprojected rows: 0
    
    Processing PXD004720 | glume
    
    Saved: python_outputs\tables\FragPipe_Duncan_PXD004720_glume_peptide_genome_projection.csv
    Projection_status
    projected    145133
    Name: count, dtype: int64
    Unprojected peptide table saved: python_outputs\tables\FragPipe_Duncan_PXD004720_glume_unprojected_peptides_for_tblastn.csv
    Unprojected rows: 0
    
    Processing PXD004720 | grain-zadoks-70
    
    Saved: python_outputs\tables\FragPipe_Duncan_PXD004720_grain-zadoks-70_peptide_genome_projection.csv
    Projection_status
    projected    126195
    Name: count, dtype: int64
    Unprojected peptide table saved: python_outputs\tables\FragPipe_Duncan_PXD004720_grain-zadoks-70_unprojected_peptides_for_tblastn.csv
    Unprojected rows: 0
    
    Processing PXD004720 | grain-zadoks-71
    
    Saved: python_outputs\tables\FragPipe_Duncan_PXD004720_grain-zadoks-71_peptide_genome_projection.csv
    Projection_status
    projected    179725
    Name: count, dtype: int64
    Unprojected peptide table saved: python_outputs\tables\FragPipe_Duncan_PXD004720_grain-zadoks-71_unprojected_peptides_for_tblastn.csv
    Unprojected rows: 0
    
    Processing PXD004720 | grain-zadoks-75
    
    Saved: python_outputs\tables\FragPipe_Duncan_PXD004720_grain-zadoks-75_peptide_genome_projection.csv
    Projection_status
    projected    132701
    Name: count, dtype: int64
    Unprojected peptide table saved: python_outputs\tables\FragPipe_Duncan_PXD004720_grain-zadoks-75_unprojected_peptides_for_tblastn.csv
    Unprojected rows: 0
    
    Processing PXD004720 | grain-zadoks-83
    
    Saved: python_outputs\tables\FragPipe_Duncan_PXD004720_grain-zadoks-83_peptide_genome_projection.csv
    Projection_status
    projected    113203
    Name: count, dtype: int64
    Unprojected peptide table saved: python_outputs\tables\FragPipe_Duncan_PXD004720_grain-zadoks-83_unprojected_peptides_for_tblastn.csv
    Unprojected rows: 0
    
    Processing PXD004720 | grain-zadoks-87
    
    Saved: python_outputs\tables\FragPipe_Duncan_PXD004720_grain-zadoks-87_peptide_genome_projection.csv
    Projection_status
    projected    111792
    Name: count, dtype: int64
    Unprojected peptide table saved: python_outputs\tables\FragPipe_Duncan_PXD004720_grain-zadoks-87_unprojected_peptides_for_tblastn.csv
    Unprojected rows: 0
    
    Processing PXD004720 | leaf-flag-mature
    
    Saved: python_outputs\tables\FragPipe_Duncan_PXD004720_leaf-flag-mature_peptide_genome_projection.csv
    Projection_status
    projected    145632
    Name: count, dtype: int64
    Unprojected peptide table saved: python_outputs\tables\FragPipe_Duncan_PXD004720_leaf-flag-mature_unprojected_peptides_for_tblastn.csv
    Unprojected rows: 0
    
    Processing PXD004720 | leaf-flag-senescing
    
    Saved: python_outputs\tables\FragPipe_Duncan_PXD004720_leaf-flag-senescing_peptide_genome_projection.csv
    Projection_status
    projected    46200
    Name: count, dtype: int64
    Unprojected peptide table saved: python_outputs\tables\FragPipe_Duncan_PXD004720_leaf-flag-senescing_unprojected_peptides_for_tblastn.csv
    Unprojected rows: 0
    
    Processing PXD004720 | leaf-flag-young
    
    Saved: python_outputs\tables\FragPipe_Duncan_PXD004720_leaf-flag-young_peptide_genome_projection.csv
    Projection_status
    projected    120821
    Name: count, dtype: int64
    Unprojected peptide table saved: python_outputs\tables\FragPipe_Duncan_PXD004720_leaf-flag-young_unprojected_peptides_for_tblastn.csv
    Unprojected rows: 0
    
    Processing PXD004720 | lemma
    
    Saved: python_outputs\tables\FragPipe_Duncan_PXD004720_lemma_peptide_genome_projection.csv
    Projection_status
    projected    147618
    Name: count, dtype: int64
    Unprojected peptide table saved: python_outputs\tables\FragPipe_Duncan_PXD004720_lemma_unprojected_peptides_for_tblastn.csv
    Unprojected rows: 0
    
    Processing PXD004720 | node
    
    Saved: python_outputs\tables\FragPipe_Duncan_PXD004720_node_peptide_genome_projection.csv
    Projection_status
    projected    99823
    Name: count, dtype: int64
    Unprojected peptide table saved: python_outputs\tables\FragPipe_Duncan_PXD004720_node_unprojected_peptides_for_tblastn.csv
    Unprojected rows: 0
    
    Processing PXD004720 | node_secretion
    
    Saved: python_outputs\tables\FragPipe_Duncan_PXD004720_node-secretion_peptide_genome_projection.csv
    Projection_status
    projected    169675
    Name: count, dtype: int64
    Unprojected peptide table saved: python_outputs\tables\FragPipe_Duncan_PXD004720_node-secretion_unprojected_peptides_for_tblastn.csv
    Unprojected rows: 0
    
    Processing PXD004720 | palea
    
    Saved: python_outputs\tables\FragPipe_Duncan_PXD004720_palea_peptide_genome_projection.csv
    Projection_status
    projected    116936
    Name: count, dtype: int64
    Unprojected peptide table saved: python_outputs\tables\FragPipe_Duncan_PXD004720_palea_unprojected_peptides_for_tblastn.csv
    Unprojected rows: 0
    
    Processing PXD004720 | pericarp
    
    Saved: python_outputs\tables\FragPipe_Duncan_PXD004720_pericarp_peptide_genome_projection.csv
    Projection_status
    projected    131215
    Name: count, dtype: int64
    Unprojected peptide table saved: python_outputs\tables\FragPipe_Duncan_PXD004720_pericarp_unprojected_peptides_for_tblastn.csv
    Unprojected rows: 0
    
    Processing PXD004720 | pollen
    
    Saved: python_outputs\tables\FragPipe_Duncan_PXD004720_pollen_peptide_genome_projection.csv
    Projection_status
    projected    74698
    Name: count, dtype: int64
    Unprojected peptide table saved: python_outputs\tables\FragPipe_Duncan_PXD004720_pollen_unprojected_peptides_for_tblastn.csv
    Unprojected rows: 0
    
    Processing PXD004720 | rachilla
    
    Saved: python_outputs\tables\FragPipe_Duncan_PXD004720_rachilla_peptide_genome_projection.csv
    Projection_status
    projected    160204
    Name: count, dtype: int64
    Unprojected peptide table saved: python_outputs\tables\FragPipe_Duncan_PXD004720_rachilla_unprojected_peptides_for_tblastn.csv
    Unprojected rows: 0
    
    Processing PXD004720 | radicle
    
    Saved: python_outputs\tables\FragPipe_Duncan_PXD004720_radicle_peptide_genome_projection.csv
    Projection_status
    projected    194694
    Name: count, dtype: int64
    Unprojected peptide table saved: python_outputs\tables\FragPipe_Duncan_PXD004720_radicle_unprojected_peptides_for_tblastn.csv
    Unprojected rows: 0
    
    Processing PXD004720 | root-mature
    
    Saved: python_outputs\tables\FragPipe_Duncan_PXD004720_root-mature_peptide_genome_projection.csv
    Projection_status
    projected    97188
    Name: count, dtype: int64
    Unprojected peptide table saved: python_outputs\tables\FragPipe_Duncan_PXD004720_root-mature_unprojected_peptides_for_tblastn.csv
    Unprojected rows: 0
    
    Processing PXD004720 | root-secretion
    
    Saved: python_outputs\tables\FragPipe_Duncan_PXD004720_root-secretion_peptide_genome_projection.csv
    Projection_status
    projected    100639
    Name: count, dtype: int64
    Unprojected peptide table saved: python_outputs\tables\FragPipe_Duncan_PXD004720_root-secretion_unprojected_peptides_for_tblastn.csv
    Unprojected rows: 0
    
    Processing PXD004720 | root-tip
    
    Saved: python_outputs\tables\FragPipe_Duncan_PXD004720_root-tip_peptide_genome_projection.csv
    Projection_status
    projected    191943
    Name: count, dtype: int64
    Unprojected peptide table saved: python_outputs\tables\FragPipe_Duncan_PXD004720_root-tip_unprojected_peptides_for_tblastn.csv
    Unprojected rows: 0
    
    Processing PXD004720 | root-vasculature
    
    Saved: python_outputs\tables\FragPipe_Duncan_PXD004720_root-vasculature_peptide_genome_projection.csv
    Projection_status
    projected    111831
    Name: count, dtype: int64
    Unprojected peptide table saved: python_outputs\tables\FragPipe_Duncan_PXD004720_root-vasculature_unprojected_peptides_for_tblastn.csv
    Unprojected rows: 0
    
    Processing PXD004720 | spike-immature
    
    Saved: python_outputs\tables\FragPipe_Duncan_PXD004720_spike-immature_peptide_genome_projection.csv
    Projection_status
    projected    188826
    Name: count, dtype: int64
    Unprojected peptide table saved: python_outputs\tables\FragPipe_Duncan_PXD004720_spike-immature_unprojected_peptides_for_tblastn.csv
    Unprojected rows: 0
    
    Processing PXD004720 | stem
    
    Saved: python_outputs\tables\FragPipe_Duncan_PXD004720_stem_peptide_genome_projection.csv
    Projection_status
    projected    60734
    Name: count, dtype: int64
    Unprojected peptide table saved: python_outputs\tables\FragPipe_Duncan_PXD004720_stem_unprojected_peptides_for_tblastn.csv
    Unprojected rows: 0
    
    Step 9 summary saved: python_outputs\tables\wheat_peptide_genome_projection_summary_step9.csv
    


<div>
<style scoped>
    .dataframe tbody tr th:only-of-type {
        vertical-align: middle;
    }

    .dataframe tbody tr th {
        vertical-align: top;
    }

    .dataframe thead th {
        text-align: right;
    }
</style>
<table border="1" class="dataframe">
  <thead>
    <tr style="text-align: right;">
      <th></th>
      <th>Source</th>
      <th>Species</th>
      <th>Tissue</th>
      <th>Batch</th>
      <th>Genome_projection_file</th>
      <th>Peptide_protein_gene_rows</th>
      <th>Projected_rows</th>
      <th>Unprojected_rows</th>
      <th>Projection_rate_percent</th>
      <th>Unique_projected_peptides</th>
      <th>Unique_projected_proteins</th>
      <th>Unique_projected_gene_models</th>
      <th>Peptides_crossing_CDS_blocks</th>
      <th>Peptides_with_intron_gapped_label</th>
    </tr>
  </thead>
  <tbody>
    <tr>
      <th>0</th>
      <td>MSV000090572</td>
      <td>bread wheat</td>
      <td>stored_grain</td>
      <td>single</td>
      <td>FragPipe_Vincent_MSV000090572_stored-grain_pep...</td>
      <td>30314</td>
      <td>30314</td>
      <td>0</td>
      <td>100.0</td>
      <td>9329</td>
      <td>17478</td>
      <td>&lt;NA&gt;</td>
      <td>3291</td>
      <td>1828</td>
    </tr>
    <tr>
      <th>1</th>
      <td>PXD050500</td>
      <td>bread wheat</td>
      <td>coleoptile</td>
      <td>single</td>
      <td>FragPipe_Liu_PXD050500_coleoptile_peptide_geno...</td>
      <td>1850136</td>
      <td>1850136</td>
      <td>0</td>
      <td>100.0</td>
      <td>582775</td>
      <td>239461</td>
      <td>&lt;NA&gt;</td>
      <td>228372</td>
      <td>122385</td>
    </tr>
    <tr>
      <th>2</th>
      <td>PXD050500</td>
      <td>bread wheat</td>
      <td>node</td>
      <td>single</td>
      <td>FragPipe_Liu_PXD050500_node_peptide_genome_pro...</td>
      <td>1886922</td>
      <td>1886922</td>
      <td>0</td>
      <td>100.0</td>
      <td>595954</td>
      <td>242813</td>
      <td>&lt;NA&gt;</td>
      <td>235642</td>
      <td>127047</td>
    </tr>
    <tr>
      <th>3</th>
      <td>PXD050500</td>
      <td>bread wheat</td>
      <td>radicle</td>
      <td>single</td>
      <td>FragPipe_Liu_PXD050500_radicle_peptide_genome_...</td>
      <td>1062508</td>
      <td>1062508</td>
      <td>0</td>
      <td>100.0</td>
      <td>333058</td>
      <td>206016</td>
      <td>&lt;NA&gt;</td>
      <td>118195</td>
      <td>62337</td>
    </tr>
    <tr>
      <th>4</th>
      <td>PXD004720</td>
      <td>bread wheat</td>
      <td>anther</td>
      <td>single</td>
      <td>FragPipe_Duncan_PXD004720_anther_peptide_genom...</td>
      <td>164365</td>
      <td>164365</td>
      <td>0</td>
      <td>100.0</td>
      <td>34098</td>
      <td>36826</td>
      <td>&lt;NA&gt;</td>
      <td>27776</td>
      <td>16150</td>
    </tr>
    <tr>
      <th>5</th>
      <td>PXD004720</td>
      <td>bread wheat</td>
      <td>boot</td>
      <td>single</td>
      <td>FragPipe_Duncan_PXD004720_boot_peptide_genome_...</td>
      <td>13227</td>
      <td>13227</td>
      <td>0</td>
      <td>100.0</td>
      <td>3643</td>
      <td>9208</td>
      <td>&lt;NA&gt;</td>
      <td>2728</td>
      <td>1537</td>
    </tr>
    <tr>
      <th>6</th>
      <td>PXD004720</td>
      <td>bread wheat</td>
      <td>coleoptile</td>
      <td>single</td>
      <td>FragPipe_Duncan_PXD004720_coleoptile_peptide_g...</td>
      <td>204476</td>
      <td>204476</td>
      <td>0</td>
      <td>100.0</td>
      <td>41025</td>
      <td>46496</td>
      <td>&lt;NA&gt;</td>
      <td>31344</td>
      <td>18679</td>
    </tr>
    <tr>
      <th>7</th>
      <td>PXD004720</td>
      <td>bread wheat</td>
      <td>embryo</td>
      <td>single</td>
      <td>FragPipe_Duncan_PXD004720_embryo_peptide_genom...</td>
      <td>8852</td>
      <td>8852</td>
      <td>0</td>
      <td>100.0</td>
      <td>2867</td>
      <td>7220</td>
      <td>&lt;NA&gt;</td>
      <td>1411</td>
      <td>760</td>
    </tr>
    <tr>
      <th>8</th>
      <td>PXD004720</td>
      <td>bread wheat</td>
      <td>endosperm</td>
      <td>single</td>
      <td>FragPipe_Duncan_PXD004720_endosperm_peptide_ge...</td>
      <td>102830</td>
      <td>102830</td>
      <td>0</td>
      <td>100.0</td>
      <td>20081</td>
      <td>27454</td>
      <td>&lt;NA&gt;</td>
      <td>14269</td>
      <td>7971</td>
    </tr>
    <tr>
      <th>9</th>
      <td>PXD004720</td>
      <td>bread wheat</td>
      <td>glume</td>
      <td>single</td>
      <td>FragPipe_Duncan_PXD004720_glume_peptide_genome...</td>
      <td>145133</td>
      <td>145133</td>
      <td>0</td>
      <td>100.0</td>
      <td>28648</td>
      <td>35540</td>
      <td>&lt;NA&gt;</td>
      <td>22131</td>
      <td>12759</td>
    </tr>
    <tr>
      <th>10</th>
      <td>PXD004720</td>
      <td>bread wheat</td>
      <td>grain-zadoks-70</td>
      <td>single</td>
      <td>FragPipe_Duncan_PXD004720_grain-zadoks-70_pept...</td>
      <td>126195</td>
      <td>126195</td>
      <td>0</td>
      <td>100.0</td>
      <td>26196</td>
      <td>35532</td>
      <td>&lt;NA&gt;</td>
      <td>17478</td>
      <td>9892</td>
    </tr>
    <tr>
      <th>11</th>
      <td>PXD004720</td>
      <td>bread wheat</td>
      <td>grain-zadoks-71</td>
      <td>single</td>
      <td>FragPipe_Duncan_PXD004720_grain-zadoks-71_pept...</td>
      <td>179725</td>
      <td>179725</td>
      <td>0</td>
      <td>100.0</td>
      <td>35960</td>
      <td>47076</td>
      <td>&lt;NA&gt;</td>
      <td>27658</td>
      <td>15967</td>
    </tr>
    <tr>
      <th>12</th>
      <td>PXD004720</td>
      <td>bread wheat</td>
      <td>grain-zadoks-75</td>
      <td>single</td>
      <td>FragPipe_Duncan_PXD004720_grain-zadoks-75_pept...</td>
      <td>132701</td>
      <td>132701</td>
      <td>0</td>
      <td>100.0</td>
      <td>27463</td>
      <td>41029</td>
      <td>&lt;NA&gt;</td>
      <td>18650</td>
      <td>10504</td>
    </tr>
    <tr>
      <th>13</th>
      <td>PXD004720</td>
      <td>bread wheat</td>
      <td>grain-zadoks-83</td>
      <td>single</td>
      <td>FragPipe_Duncan_PXD004720_grain-zadoks-83_pept...</td>
      <td>113203</td>
      <td>113203</td>
      <td>0</td>
      <td>100.0</td>
      <td>23990</td>
      <td>36635</td>
      <td>&lt;NA&gt;</td>
      <td>14734</td>
      <td>8452</td>
    </tr>
    <tr>
      <th>14</th>
      <td>PXD004720</td>
      <td>bread wheat</td>
      <td>grain-zadoks-87</td>
      <td>single</td>
      <td>FragPipe_Duncan_PXD004720_grain-zadoks-87_pept...</td>
      <td>111792</td>
      <td>111792</td>
      <td>0</td>
      <td>100.0</td>
      <td>24636</td>
      <td>34874</td>
      <td>&lt;NA&gt;</td>
      <td>14886</td>
      <td>8402</td>
    </tr>
    <tr>
      <th>15</th>
      <td>PXD004720</td>
      <td>bread wheat</td>
      <td>leaf-flag-mature</td>
      <td>single</td>
      <td>FragPipe_Duncan_PXD004720_leaf-flag-mature_pep...</td>
      <td>145632</td>
      <td>145632</td>
      <td>0</td>
      <td>100.0</td>
      <td>29840</td>
      <td>37954</td>
      <td>&lt;NA&gt;</td>
      <td>19448</td>
      <td>11285</td>
    </tr>
    <tr>
      <th>16</th>
      <td>PXD004720</td>
      <td>bread wheat</td>
      <td>leaf-flag-senescing</td>
      <td>single</td>
      <td>FragPipe_Duncan_PXD004720_leaf-flag-senescing_...</td>
      <td>46200</td>
      <td>46200</td>
      <td>0</td>
      <td>100.0</td>
      <td>11354</td>
      <td>25696</td>
      <td>&lt;NA&gt;</td>
      <td>4572</td>
      <td>2415</td>
    </tr>
    <tr>
      <th>17</th>
      <td>PXD004720</td>
      <td>bread wheat</td>
      <td>leaf-flag-young</td>
      <td>single</td>
      <td>FragPipe_Duncan_PXD004720_leaf-flag-young_pept...</td>
      <td>120821</td>
      <td>120821</td>
      <td>0</td>
      <td>100.0</td>
      <td>23373</td>
      <td>32620</td>
      <td>&lt;NA&gt;</td>
      <td>15227</td>
      <td>8981</td>
    </tr>
    <tr>
      <th>18</th>
      <td>PXD004720</td>
      <td>bread wheat</td>
      <td>lemma</td>
      <td>single</td>
      <td>FragPipe_Duncan_PXD004720_lemma_peptide_genome...</td>
      <td>147618</td>
      <td>147618</td>
      <td>0</td>
      <td>100.0</td>
      <td>29788</td>
      <td>37927</td>
      <td>&lt;NA&gt;</td>
      <td>21753</td>
      <td>12440</td>
    </tr>
    <tr>
      <th>19</th>
      <td>PXD004720</td>
      <td>bread wheat</td>
      <td>node</td>
      <td>single</td>
      <td>FragPipe_Duncan_PXD004720_node_peptide_genome_...</td>
      <td>99823</td>
      <td>99823</td>
      <td>0</td>
      <td>100.0</td>
      <td>21457</td>
      <td>36542</td>
      <td>&lt;NA&gt;</td>
      <td>12765</td>
      <td>7108</td>
    </tr>
    <tr>
      <th>20</th>
      <td>PXD004720</td>
      <td>bread wheat</td>
      <td>node_secretion</td>
      <td>single</td>
      <td>FragPipe_Duncan_PXD004720_node-secretion_pepti...</td>
      <td>169675</td>
      <td>169675</td>
      <td>0</td>
      <td>100.0</td>
      <td>34124</td>
      <td>42356</td>
      <td>&lt;NA&gt;</td>
      <td>22888</td>
      <td>13292</td>
    </tr>
    <tr>
      <th>21</th>
      <td>PXD004720</td>
      <td>bread wheat</td>
      <td>palea</td>
      <td>single</td>
      <td>FragPipe_Duncan_PXD004720_palea_peptide_genome...</td>
      <td>116936</td>
      <td>116936</td>
      <td>0</td>
      <td>100.0</td>
      <td>21707</td>
      <td>27580</td>
      <td>&lt;NA&gt;</td>
      <td>20681</td>
      <td>12105</td>
    </tr>
    <tr>
      <th>22</th>
      <td>PXD004720</td>
      <td>bread wheat</td>
      <td>pericarp</td>
      <td>single</td>
      <td>FragPipe_Duncan_PXD004720_pericarp_peptide_gen...</td>
      <td>131215</td>
      <td>131215</td>
      <td>0</td>
      <td>100.0</td>
      <td>28332</td>
      <td>32667</td>
      <td>&lt;NA&gt;</td>
      <td>20964</td>
      <td>11953</td>
    </tr>
    <tr>
      <th>23</th>
      <td>PXD004720</td>
      <td>bread wheat</td>
      <td>pollen</td>
      <td>single</td>
      <td>FragPipe_Duncan_PXD004720_pollen_peptide_genom...</td>
      <td>74698</td>
      <td>74698</td>
      <td>0</td>
      <td>100.0</td>
      <td>13502</td>
      <td>19931</td>
      <td>&lt;NA&gt;</td>
      <td>9490</td>
      <td>5338</td>
    </tr>
    <tr>
      <th>24</th>
      <td>PXD004720</td>
      <td>bread wheat</td>
      <td>rachilla</td>
      <td>single</td>
      <td>FragPipe_Duncan_PXD004720_rachilla_peptide_gen...</td>
      <td>160204</td>
      <td>160204</td>
      <td>0</td>
      <td>100.0</td>
      <td>31145</td>
      <td>37204</td>
      <td>&lt;NA&gt;</td>
      <td>24784</td>
      <td>14463</td>
    </tr>
    <tr>
      <th>25</th>
      <td>PXD004720</td>
      <td>bread wheat</td>
      <td>radicle</td>
      <td>single</td>
      <td>FragPipe_Duncan_PXD004720_radicle_peptide_geno...</td>
      <td>194694</td>
      <td>194694</td>
      <td>0</td>
      <td>100.0</td>
      <td>39643</td>
      <td>42965</td>
      <td>&lt;NA&gt;</td>
      <td>32499</td>
      <td>19061</td>
    </tr>
    <tr>
      <th>26</th>
      <td>PXD004720</td>
      <td>bread wheat</td>
      <td>root-mature</td>
      <td>single</td>
      <td>FragPipe_Duncan_PXD004720_root-mature_peptide_...</td>
      <td>97188</td>
      <td>97188</td>
      <td>0</td>
      <td>100.0</td>
      <td>21350</td>
      <td>39156</td>
      <td>&lt;NA&gt;</td>
      <td>10898</td>
      <td>6135</td>
    </tr>
    <tr>
      <th>27</th>
      <td>PXD004720</td>
      <td>bread wheat</td>
      <td>root-secretion</td>
      <td>single</td>
      <td>FragPipe_Duncan_PXD004720_root-secretion_pepti...</td>
      <td>100639</td>
      <td>100639</td>
      <td>0</td>
      <td>100.0</td>
      <td>21284</td>
      <td>32046</td>
      <td>&lt;NA&gt;</td>
      <td>15547</td>
      <td>8991</td>
    </tr>
    <tr>
      <th>28</th>
      <td>PXD004720</td>
      <td>bread wheat</td>
      <td>root-tip</td>
      <td>single</td>
      <td>FragPipe_Duncan_PXD004720_root-tip_peptide_gen...</td>
      <td>191943</td>
      <td>191943</td>
      <td>0</td>
      <td>100.0</td>
      <td>38634</td>
      <td>39774</td>
      <td>&lt;NA&gt;</td>
      <td>35031</td>
      <td>21057</td>
    </tr>
    <tr>
      <th>29</th>
      <td>PXD004720</td>
      <td>bread wheat</td>
      <td>root-vasculature</td>
      <td>single</td>
      <td>FragPipe_Duncan_PXD004720_root-vasculature_pep...</td>
      <td>111831</td>
      <td>111831</td>
      <td>0</td>
      <td>100.0</td>
      <td>20815</td>
      <td>30626</td>
      <td>&lt;NA&gt;</td>
      <td>14833</td>
      <td>8183</td>
    </tr>
    <tr>
      <th>30</th>
      <td>PXD004720</td>
      <td>bread wheat</td>
      <td>spike-immature</td>
      <td>single</td>
      <td>FragPipe_Duncan_PXD004720_spike-immature_pepti...</td>
      <td>188826</td>
      <td>188826</td>
      <td>0</td>
      <td>100.0</td>
      <td>36370</td>
      <td>40207</td>
      <td>&lt;NA&gt;</td>
      <td>33193</td>
      <td>19376</td>
    </tr>
    <tr>
      <th>31</th>
      <td>PXD004720</td>
      <td>bread wheat</td>
      <td>stem</td>
      <td>single</td>
      <td>FragPipe_Duncan_PXD004720_stem_peptide_genome_...</td>
      <td>60734</td>
      <td>60734</td>
      <td>0</td>
      <td>100.0</td>
      <td>14338</td>
      <td>29410</td>
      <td>&lt;NA&gt;</td>
      <td>6949</td>
      <td>3980</td>
    </tr>
  </tbody>
</table>
</div>


```python
# Testing intron annotation at the peptide level
test_gapped = projected[
    projected["Peptide_intron_gapped"].astype(str).str.contains("-", regex=False)
][["Peptide", "Peptide_intron_gapped", "BED_block_count", "BED_block_sizes", "BED_block_starts"]]

display(test_gapped.head(20))
```


<div>
<style scoped>
    .dataframe tbody tr th:only-of-type {
        vertical-align: middle;
    }

    .dataframe tbody tr th {
        vertical-align: top;
    }

    .dataframe thead th {
        text-align: right;
    }
</style>
<table border="1" class="dataframe">
  <thead>
    <tr style="text-align: right;">
      <th></th>
      <th>Peptide</th>
      <th>Peptide_intron_gapped</th>
      <th>BED_block_count</th>
      <th>BED_block_sizes</th>
      <th>BED_block_starts</th>
    </tr>
  </thead>
  <tbody>
    <tr>
      <th>202</th>
      <td>AAAAAGEPQNGKR</td>
      <td>AAAAAGE------------------PQNGKR</td>
      <td>2</td>
      <td>18,21</td>
      <td>0,36</td>
    </tr>
    <tr>
      <th>309</th>
      <td>AAAAEGDLDLQAK</td>
      <td>AAAAE-----------------------------------------...</td>
      <td>2</td>
      <td>15,24</td>
      <td>0,1904</td>
    </tr>
    <tr>
      <th>310</th>
      <td>AAAAEGDLDLQAK</td>
      <td>AAAAE-----------------------------------------...</td>
      <td>2</td>
      <td>15,24</td>
      <td>0,1120</td>
    </tr>
    <tr>
      <th>311</th>
      <td>AAAAEGDLDLQAK</td>
      <td>AAAAE-----------------------------------------...</td>
      <td>2</td>
      <td>15,24</td>
      <td>0,2111</td>
    </tr>
    <tr>
      <th>314</th>
      <td>AAAAGAAPSGGPVFKSGPLFISSK</td>
      <td>AAAAGAAPSGGP----------------------------------...</td>
      <td>2</td>
      <td>36,36</td>
      <td>0,964</td>
    </tr>
    <tr>
      <th>315</th>
      <td>AAAAGAAPSGGPVFKSGPLFISSK</td>
      <td>AAAAGAAPSGGP----------------------------------...</td>
      <td>2</td>
      <td>36,36</td>
      <td>0,964</td>
    </tr>
    <tr>
      <th>316</th>
      <td>AAAAGAAPSGGPVFKSGPLFISSK</td>
      <td>AAAAGAAPSGGP----------------------------------...</td>
      <td>2</td>
      <td>36,36</td>
      <td>0,964</td>
    </tr>
    <tr>
      <th>317</th>
      <td>AAAAGAAPSGGPVFKSGPLFISSK</td>
      <td>AAAAGAAPSGGP----------------------------------...</td>
      <td>2</td>
      <td>36,36</td>
      <td>0,964</td>
    </tr>
    <tr>
      <th>318</th>
      <td>AAAAGAAPSGGPVFKSGPLFISSK</td>
      <td>AAAAGAAPSGGP----------------------------------...</td>
      <td>2</td>
      <td>36,36</td>
      <td>0,1019</td>
    </tr>
    <tr>
      <th>319</th>
      <td>AAAAGAAPSGGPVFKSGPLFISSK</td>
      <td>AAAAGAAPSGGP----------------------------------...</td>
      <td>2</td>
      <td>36,36</td>
      <td>0,1019</td>
    </tr>
    <tr>
      <th>320</th>
      <td>AAAAGAAPSGGPVFKSGPLFISSK</td>
      <td>AAAAGAAPSGGP----------------------------------...</td>
      <td>2</td>
      <td>36,36</td>
      <td>0,1019</td>
    </tr>
    <tr>
      <th>321</th>
      <td>AAAAGAAPSGGPVFKSGPLFISSK</td>
      <td>AAAAGAAPSGGP----------------------------------...</td>
      <td>2</td>
      <td>36,36</td>
      <td>0,1019</td>
    </tr>
    <tr>
      <th>322</th>
      <td>AAAAGAAPSGGPVFKSGPLFISSK</td>
      <td>AAAAGAAPSGGP----------------------------------...</td>
      <td>2</td>
      <td>36,36</td>
      <td>0,1019</td>
    </tr>
    <tr>
      <th>323</th>
      <td>AAAAGAAPSGGPVFKSGPLFISSK</td>
      <td>AAAAGAAPSGGP----------------------------------...</td>
      <td>2</td>
      <td>36,36</td>
      <td>0,1019</td>
    </tr>
    <tr>
      <th>324</th>
      <td>AAAAGAAPSGGPVFKSGPLFISSK</td>
      <td>AAAAGAAPSGGP----------------------------------...</td>
      <td>2</td>
      <td>36,36</td>
      <td>0,1018</td>
    </tr>
    <tr>
      <th>325</th>
      <td>AAAAGAAPSGGPVFKSGPLFISSK</td>
      <td>AAAAGAAPSGGP----------------------------------...</td>
      <td>2</td>
      <td>36,36</td>
      <td>0,1018</td>
    </tr>
    <tr>
      <th>326</th>
      <td>AAAAGAAPSGGPVFKSGPLFISSK</td>
      <td>AAAAGAAPSGGP----------------------------------...</td>
      <td>2</td>
      <td>36,36</td>
      <td>0,1018</td>
    </tr>
    <tr>
      <th>327</th>
      <td>AAAAGAAPSGGPVFKSGPLFISSK</td>
      <td>AAAAGAAPSGGP----------------------------------...</td>
      <td>2</td>
      <td>36,36</td>
      <td>0,1018</td>
    </tr>
    <tr>
      <th>328</th>
      <td>AAAAGAAPSGGPVFKSGPLFISSK</td>
      <td>AAAAGAAPSGGP----------------------------------...</td>
      <td>2</td>
      <td>36,36</td>
      <td>0,1018</td>
    </tr>
    <tr>
      <th>329</th>
      <td>AAAAGAAPSGGPVFKSGPLFISSK</td>
      <td>AAAAGAAPSGGP----------------------------------...</td>
      <td>2</td>
      <td>36,36</td>
      <td>0,1018</td>
    </tr>
  </tbody>
</table>
</div>


# Step 10 — Translation validation of annotation-guided peptide genome projections

This step validates a stratified random subset of peptide genome projections generated in Step 9.

---

The validation tests whether projected genomic coordinates can reconstruct the original peptide sequence. For each sampled projected peptide, the workflow:

1. Reads the peptide genomic coordinates from the Step 9 projection tables.
2. Extracts the corresponding genomic DNA sequence from the wheat reference genome.
3. Reconstructs the spliced coding sequence using the BED block structure.
4. Reverse-complements the sequence when the peptide maps to the negative strand.
5. Translates the reconstructed coding DNA sequence.
6. Compares the translated amino acid sequence with the original FragPipe peptide sequence.

This provides a direct validation of the annotation-guided projection logic, including:

- exon-aware block reconstruction,
- splice-junction handling,
- strand-aware sequence recovery,
- peptide-level coordinate accuracy.

Unlike heuristic alignment approaches, this strategy computationally confirms that projected genomic coordinates can regenerate the experimentally observed peptide sequence.

---

Two matching criteria are reported:

- `Exact_match`  
  The translated peptide is identical to the original peptide sequence.

- `IL_normalised_match`  
  The translated peptide matches after converting both I and L to L, because tandem mass spectrometry usually cannot distinguish isoleucine from leucine.

The I/L-normalised comparison is considered the biologically relevant validation criterion.

---

The workflow also records:

- reconstructed CDS nucleotide length,
- validation status categories,
- multi-block (exon-spanning) peptide projections,
- negative-strand peptide projections.

This allows independent assessment of complex projection scenarios such as splice-junction peptides and reverse-strand gene models.

---

The output files are:

- `wheat_projection_validation_stratified100percent_step10.csv`
- `wheat_projection_validation_100%_summary_step10.csv`
- `wheat_projection_validation_100%_tissue_summary_step10.csv`


```python
# # install python library
# !pip install pyfaidx
```


```python
# ============================================================
# Step 10 — Validate peptide genome projections by translation (takes 30 min)
# ============================================================

import pandas as pd
import numpy as np
from pathlib import Path
import re

# -----------------------------
# 1. User settings
# -----------------------------
tables_dir = Path("python_outputs/tables")
genome_fasta = Path("genome_annotation/iwgsc_refseqv2.1_assembly.fa")

sample_fraction = 1 # to test on a portion of the data, lower this number (e.g. 0.1 for 10% of the data)
chunk_size = 30_000 # reduce this for low performance computer
random_seed = 42

validation_out = tables_dir / "wheat_projection_validation_stratified100percent_step10.csv"
summary_out = tables_dir / "wheat_projection_validation_100%_summary_step10.csv"
tissue_summary_out = tables_dir / "wheat_projection_validation_100%_tissue_summary_step10.csv"

# Clear previous output if rerunning
if validation_out.exists():
    validation_out.unlink()

# -----------------------------
# 2. Check pyfaidx availability
# -----------------------------
try:
    from pyfaidx import Fasta
except ImportError:
    raise ImportError(
        "Install pyfaidx first:\n\npip install pyfaidx"
    )

# -----------------------------
# 3. Find projection files
# -----------------------------
projection_files = sorted(tables_dir.glob("*_peptide_genome_projection.csv"))

if len(projection_files) == 0:
    raise FileNotFoundError(
        f"No Step 9 projection files found in: {tables_dir}"
    )

print(f"Projection files found: {len(projection_files)}")

if not genome_fasta.exists():
    raise FileNotFoundError(f"Genome FASTA not found:\n{genome_fasta}")

# -----------------------------
# 4. Helper functions
# -----------------------------
def clean_peptide_sequence(seq):
    seq = str(seq).upper()
    seq = re.sub(r"[^ACDEFGHIKLMNPQRSTVWY]", "", seq)
    return seq


def il_normalise(seq):
    return str(seq).upper().replace("I", "L")


def reverse_complement(seq):
    complement = str.maketrans("ACGTNacgtn", "TGCANtgcan")
    return str(seq).translate(complement)[::-1].upper()


genetic_code = {
    "TTT": "F", "TTC": "F", "TTA": "L", "TTG": "L",
    "TCT": "S", "TCC": "S", "TCA": "S", "TCG": "S",
    "TAT": "Y", "TAC": "Y", "TAA": "*", "TAG": "*",
    "TGT": "C", "TGC": "C", "TGA": "*", "TGG": "W",
    "CTT": "L", "CTC": "L", "CTA": "L", "CTG": "L",
    "CCT": "P", "CCC": "P", "CCA": "P", "CCG": "P",
    "CAT": "H", "CAC": "H", "CAA": "Q", "CAG": "Q",
    "CGT": "R", "CGC": "R", "CGA": "R", "CGG": "R",
    "ATT": "I", "ATC": "I", "ATA": "I", "ATG": "M",
    "ACT": "T", "ACC": "T", "ACA": "T", "ACG": "T",
    "AAT": "N", "AAC": "N", "AAA": "K", "AAG": "K",
    "AGT": "S", "AGC": "S", "AGA": "R", "AGG": "R",
    "GTT": "V", "GTC": "V", "GTA": "V", "GTG": "V",
    "GCT": "A", "GCC": "A", "GCA": "A", "GCG": "A",
    "GAT": "D", "GAC": "D", "GAA": "E", "GAG": "E",
    "GGT": "G", "GGC": "G", "GGA": "G", "GGG": "G"
}


def translate_dna(seq):
    seq = str(seq).upper()
    return "".join(
        genetic_code.get(seq[i:i + 3], "X")
        for i in range(0, len(seq) - 2, 3)
    )


def parse_bed_blocks(row):
    feature_start_1based = int(row["BED_start_0based"]) + 1

    sizes = [
        int(x) for x in str(row["BED_block_sizes"]).strip(",").split(",")
        if str(x).strip() != ""
    ]

    starts = [
        int(x) for x in str(row["BED_block_starts"]).strip(",").split(",")
        if str(x).strip() != ""
    ]

    if len(sizes) != len(starts):
        return []

    return [
        (feature_start_1based + rel_start,
         feature_start_1based + rel_start + size - 1)
        for size, rel_start in zip(sizes, starts)
    ]


def extract_projected_cds(row, genome):
    chrom = str(row["Chromosome"])
    strand = str(row["Strand"])

    blocks = parse_bed_blocks(row)

    if len(blocks) == 0:
        return ""

    seq_parts = []

    for start, end in blocks:
        seq_parts.append(genome[chrom][start - 1:end].seq)

    cds_seq = "".join(seq_parts).upper()

    if strand == "-":
        cds_seq = reverse_complement(cds_seq)

    return cds_seq


def validate_projection_row(row_dict, genome):
    peptide = clean_peptide_sequence(row_dict["Peptide"])
    validation_status = "validated"

    try:
        cds_seq = extract_projected_cds(row_dict, genome)
        translated = translate_dna(cds_seq)

        exact_match = translated == peptide
        il_match = il_normalise(translated) == il_normalise(peptide)

        expected_nt_length = len(peptide) * 3
        observed_nt_length = len(cds_seq)

        if observed_nt_length != expected_nt_length:
            validation_status = "length_mismatch"
        elif "N" in cds_seq:
            validation_status = "contains_N"
        elif not il_match:
            validation_status = "translation_mismatch"

    except KeyError:
        cds_seq = ""
        translated = ""
        exact_match = False
        il_match = False
        expected_nt_length = len(peptide) * 3
        observed_nt_length = 0
        validation_status = "chromosome_not_found_in_genome_fasta"

    except Exception as e:
        cds_seq = ""
        translated = ""
        exact_match = False
        il_match = False
        expected_nt_length = len(peptide) * 3
        observed_nt_length = 0
        validation_status = f"error: {type(e).__name__}"

    row_dict.update({
        "Original_peptide_clean": peptide,
        "Reconstructed_CDS_length_nt": observed_nt_length,
        "Expected_CDS_length_nt": expected_nt_length,
        "Translated_projected_peptide": translated,
        "Exact_match": exact_match,
        "IL_normalised_match": il_match,
        "Validation_status": validation_status
    })

    return row_dict


# -----------------------------
# 5. Required columns
# -----------------------------
required_cols = [
    "Source",
    "Species",
    "Tissue",
    "Batch",
    "Peptide",
    "ProteinID",
    "TranscriptID",
    "GeneModel",
    "GeneID",
    "Annotation_confidence",
    "Chromosome",
    "Strand",
    "AA_start",
    "AA_end",
    "BED_start_0based",
    "BED_end_0based_exclusive",
    "BED_block_count",
    "BED_block_sizes",
    "BED_block_starts",
    "Projection_status",
    "Probability",
    "Peptide_intron_gapped",
    "Peptide_intron_gapped_compact",
    "Tissues_count"
]

# -----------------------------
# 6. Open genome FASTA
# -----------------------------
print("\nOpening indexed genome FASTA...")

# Use rebuild=True only the first time. Afterwards, rebuild=False is faster.
genome = Fasta(str(genome_fasta), rebuild=False)

print(f"Genome sequences available: {len(genome.keys()):,}")

# -----------------------------
# 7. Validate stratified 100% per projection file
# -----------------------------
rng = np.random.default_rng(random_seed)

overall_records = []
tissue_summary_records = []
header_written = False

print("\nValidating stratified 100% sample per tissue/source...")

for file_i, projection_file in enumerate(projection_files, start=1):

    print(f"\n[{file_i}/{len(projection_files)}] {projection_file.name}")

    header = pd.read_csv(projection_file, nrows=0)
    available_cols = [c for c in required_cols if c in header.columns]

    file_total_projected = 0
    file_sampled = 0
    file_exact = 0
    file_il = 0
    file_multiblock = 0
    file_negative = 0
    file_status_counts = {}

    for chunk_i, chunk in enumerate(
        pd.read_csv(
            projection_file,
            usecols=available_cols,
            chunksize=chunk_size,
            low_memory=True
        ),
        start=1
    ):

        if "Projection_status" not in chunk.columns:
            continue

        chunk = chunk[chunk["Projection_status"] == "projected"].copy()

        if chunk.empty:
            continue

        file_total_projected += len(chunk)

        # Stratified 100% sample within this file/chunk
        sampled = chunk.sample(
            frac=sample_fraction,
            random_state=random_seed + file_i + chunk_i
        ).copy()

        if sampled.empty:
            continue

        sampled["_source_file"] = projection_file.name

        validation_records = []

        for row in sampled.itertuples(index=False):
            row_dict = row._asdict()
            validation_records.append(validate_projection_row(row_dict, genome))

        validation_chunk = pd.DataFrame(validation_records)

        # Update counters
        file_sampled += len(validation_chunk)
        file_exact += int(validation_chunk["Exact_match"].sum())
        file_il += int(validation_chunk["IL_normalised_match"].sum())
        file_multiblock += int(
            (pd.to_numeric(validation_chunk["BED_block_count"], errors="coerce") > 1).sum()
        )
        file_negative += int((validation_chunk["Strand"].astype(str) == "-").sum())

        for status, count in validation_chunk["Validation_status"].value_counts().items():
            file_status_counts[status] = file_status_counts.get(status, 0) + int(count)

        # Append to CSV immediately
        validation_chunk.to_csv(
            validation_out,
            index=False,
            mode="a",
            header=not header_written
        )

        header_written = True

        print(
            f"  Chunk {chunk_i}: sampled {len(validation_chunk):,} "
            f"| cumulative sampled {file_sampled:,}"
        )

    tissue_summary_records.append({
        "Projection_file": projection_file.name,
        "Projected_rows_available": file_total_projected,
        "Sample_fraction": sample_fraction,
        "Rows_validated": file_sampled,
        "Exact_translation_matches": file_exact,
        "Exact_translation_match_rate_percent": round((file_exact / file_sampled) * 100, 2) if file_sampled > 0 else pd.NA,
        "IL_normalised_translation_matches": file_il,
        "IL_normalised_translation_match_rate_percent": round((file_il / file_sampled) * 100, 2) if file_sampled > 0 else pd.NA,
        "Multi_block_peptide_projections_tested": file_multiblock,
        "Negative_strand_peptide_projections_tested": file_negative,
        **{f"Validation_status_{k}": v for k, v in file_status_counts.items()}
    })

# -----------------------------
# 8. Build final summaries
# -----------------------------
tissue_summary = pd.DataFrame(tissue_summary_records)
tissue_summary.to_csv(tissue_summary_out, index=False)

total_validated = int(tissue_summary["Rows_validated"].sum())
total_exact = int(tissue_summary["Exact_translation_matches"].sum())
total_il = int(tissue_summary["IL_normalised_translation_matches"].sum())
total_multiblock = int(tissue_summary["Multi_block_peptide_projections_tested"].sum())
total_negative = int(tissue_summary["Negative_strand_peptide_projections_tested"].sum())

summary_records = [
    {"Metric": "Sampling strategy", "Value": "100% stratified per projection file/source-tissue"},
    {"Metric": "Sample fraction", "Value": sample_fraction},
    {"Metric": "Projection files validated", "Value": len(tissue_summary)},
    {"Metric": "Projected rows available across files", "Value": int(tissue_summary["Projected_rows_available"].sum())},
    {"Metric": "Projected peptide rows validated", "Value": total_validated},
    {"Metric": "Exact translation matches", "Value": total_exact},
    {"Metric": "Exact translation match rate (%)", "Value": round((total_exact / total_validated) * 100, 2) if total_validated > 0 else pd.NA},
    {"Metric": "I/L-normalised translation matches", "Value": total_il},
    {"Metric": "I/L-normalised translation match rate (%)", "Value": round((total_il / total_validated) * 100, 2) if total_validated > 0 else pd.NA},
    {"Metric": "Multi-block peptide projections tested", "Value": total_multiblock},
    {"Metric": "Negative-strand peptide projections tested", "Value": total_negative}
]

status_cols = [c for c in tissue_summary.columns if c.startswith("Validation_status_")]

for col in status_cols:
    summary_records.append({
        "Metric": col.replace("Validation_status_", "Validation status: "),
        "Value": int(tissue_summary[col].fillna(0).sum())
    })

summary_df = pd.DataFrame(summary_records)
summary_df.to_csv(summary_out, index=False)

print("\n===== STEP 10 STRATIFIED VALIDATION SUMMARY =====")
display(summary_df)

print(f"\nValidation table saved: {validation_out}")
print(f"Tissue-level validation summary saved: {tissue_summary_out}")
print(f"Overall validation summary saved: {summary_out}")
```

    Projection files found: 32
    
    Opening indexed genome FASTA...
    Genome sequences available: 22
    
    Validating stratified 100% sample per tissue/source...
    
    [1/32] FragPipe_Duncan_PXD004720_anther_peptide_genome_projection.csv
      Chunk 1: sampled 30,000 | cumulative sampled 30,000
      Chunk 2: sampled 30,000 | cumulative sampled 60,000
      Chunk 3: sampled 30,000 | cumulative sampled 90,000
      Chunk 4: sampled 30,000 | cumulative sampled 120,000
      Chunk 5: sampled 30,000 | cumulative sampled 150,000
      Chunk 6: sampled 14,365 | cumulative sampled 164,365
    
    [2/32] FragPipe_Duncan_PXD004720_boot_peptide_genome_projection.csv
      Chunk 1: sampled 13,227 | cumulative sampled 13,227
    
    [3/32] FragPipe_Duncan_PXD004720_coleoptile_peptide_genome_projection.csv
      Chunk 1: sampled 30,000 | cumulative sampled 30,000
      Chunk 2: sampled 30,000 | cumulative sampled 60,000
      Chunk 3: sampled 30,000 | cumulative sampled 90,000
      Chunk 4: sampled 30,000 | cumulative sampled 120,000
      Chunk 5: sampled 30,000 | cumulative sampled 150,000
      Chunk 6: sampled 30,000 | cumulative sampled 180,000
      Chunk 7: sampled 24,476 | cumulative sampled 204,476
    
    [4/32] FragPipe_Duncan_PXD004720_embryo_peptide_genome_projection.csv
      Chunk 1: sampled 8,852 | cumulative sampled 8,852
    
    [5/32] FragPipe_Duncan_PXD004720_endosperm_peptide_genome_projection.csv
      Chunk 1: sampled 30,000 | cumulative sampled 30,000
      Chunk 2: sampled 30,000 | cumulative sampled 60,000
      Chunk 3: sampled 30,000 | cumulative sampled 90,000
      Chunk 4: sampled 12,830 | cumulative sampled 102,830
    
    [6/32] FragPipe_Duncan_PXD004720_glume_peptide_genome_projection.csv
      Chunk 1: sampled 30,000 | cumulative sampled 30,000
      Chunk 2: sampled 30,000 | cumulative sampled 60,000
      Chunk 3: sampled 30,000 | cumulative sampled 90,000
      Chunk 4: sampled 30,000 | cumulative sampled 120,000
      Chunk 5: sampled 25,133 | cumulative sampled 145,133
    
    [7/32] FragPipe_Duncan_PXD004720_grain-zadoks-70_peptide_genome_projection.csv
      Chunk 1: sampled 30,000 | cumulative sampled 30,000
      Chunk 2: sampled 30,000 | cumulative sampled 60,000
      Chunk 3: sampled 30,000 | cumulative sampled 90,000
      Chunk 4: sampled 30,000 | cumulative sampled 120,000
      Chunk 5: sampled 6,195 | cumulative sampled 126,195
    
    [8/32] FragPipe_Duncan_PXD004720_grain-zadoks-71_peptide_genome_projection.csv
      Chunk 1: sampled 30,000 | cumulative sampled 30,000
      Chunk 2: sampled 30,000 | cumulative sampled 60,000
      Chunk 3: sampled 30,000 | cumulative sampled 90,000
      Chunk 4: sampled 30,000 | cumulative sampled 120,000
      Chunk 5: sampled 30,000 | cumulative sampled 150,000
      Chunk 6: sampled 29,725 | cumulative sampled 179,725
    
    [9/32] FragPipe_Duncan_PXD004720_grain-zadoks-75_peptide_genome_projection.csv
      Chunk 1: sampled 30,000 | cumulative sampled 30,000
      Chunk 2: sampled 30,000 | cumulative sampled 60,000
      Chunk 3: sampled 30,000 | cumulative sampled 90,000
      Chunk 4: sampled 30,000 | cumulative sampled 120,000
      Chunk 5: sampled 12,701 | cumulative sampled 132,701
    
    [10/32] FragPipe_Duncan_PXD004720_grain-zadoks-83_peptide_genome_projection.csv
      Chunk 1: sampled 30,000 | cumulative sampled 30,000
      Chunk 2: sampled 30,000 | cumulative sampled 60,000
      Chunk 3: sampled 30,000 | cumulative sampled 90,000
      Chunk 4: sampled 23,203 | cumulative sampled 113,203
    
    [11/32] FragPipe_Duncan_PXD004720_grain-zadoks-87_peptide_genome_projection.csv
      Chunk 1: sampled 30,000 | cumulative sampled 30,000
      Chunk 2: sampled 30,000 | cumulative sampled 60,000
      Chunk 3: sampled 30,000 | cumulative sampled 90,000
      Chunk 4: sampled 21,792 | cumulative sampled 111,792
    
    [12/32] FragPipe_Duncan_PXD004720_leaf-flag-mature_peptide_genome_projection.csv
      Chunk 1: sampled 30,000 | cumulative sampled 30,000
      Chunk 2: sampled 30,000 | cumulative sampled 60,000
      Chunk 3: sampled 30,000 | cumulative sampled 90,000
      Chunk 4: sampled 30,000 | cumulative sampled 120,000
      Chunk 5: sampled 25,632 | cumulative sampled 145,632
    
    [13/32] FragPipe_Duncan_PXD004720_leaf-flag-senescing_peptide_genome_projection.csv
      Chunk 1: sampled 30,000 | cumulative sampled 30,000
      Chunk 2: sampled 16,200 | cumulative sampled 46,200
    
    [14/32] FragPipe_Duncan_PXD004720_leaf-flag-young_peptide_genome_projection.csv
      Chunk 1: sampled 30,000 | cumulative sampled 30,000
      Chunk 2: sampled 30,000 | cumulative sampled 60,000
      Chunk 3: sampled 30,000 | cumulative sampled 90,000
      Chunk 4: sampled 30,000 | cumulative sampled 120,000
      Chunk 5: sampled 821 | cumulative sampled 120,821
    
    [15/32] FragPipe_Duncan_PXD004720_lemma_peptide_genome_projection.csv
      Chunk 1: sampled 30,000 | cumulative sampled 30,000
      Chunk 2: sampled 30,000 | cumulative sampled 60,000
      Chunk 3: sampled 30,000 | cumulative sampled 90,000
      Chunk 4: sampled 30,000 | cumulative sampled 120,000
      Chunk 5: sampled 27,618 | cumulative sampled 147,618
    
    [16/32] FragPipe_Duncan_PXD004720_node-secretion_peptide_genome_projection.csv
      Chunk 1: sampled 30,000 | cumulative sampled 30,000
      Chunk 2: sampled 30,000 | cumulative sampled 60,000
      Chunk 3: sampled 30,000 | cumulative sampled 90,000
      Chunk 4: sampled 30,000 | cumulative sampled 120,000
      Chunk 5: sampled 30,000 | cumulative sampled 150,000
      Chunk 6: sampled 19,675 | cumulative sampled 169,675
    
    [17/32] FragPipe_Duncan_PXD004720_node_peptide_genome_projection.csv
      Chunk 1: sampled 30,000 | cumulative sampled 30,000
      Chunk 2: sampled 30,000 | cumulative sampled 60,000
      Chunk 3: sampled 30,000 | cumulative sampled 90,000
      Chunk 4: sampled 9,823 | cumulative sampled 99,823
    
    [18/32] FragPipe_Duncan_PXD004720_palea_peptide_genome_projection.csv
      Chunk 1: sampled 30,000 | cumulative sampled 30,000
      Chunk 2: sampled 30,000 | cumulative sampled 60,000
      Chunk 3: sampled 30,000 | cumulative sampled 90,000
      Chunk 4: sampled 26,936 | cumulative sampled 116,936
    
    [19/32] FragPipe_Duncan_PXD004720_pericarp_peptide_genome_projection.csv
      Chunk 1: sampled 30,000 | cumulative sampled 30,000
      Chunk 2: sampled 30,000 | cumulative sampled 60,000
      Chunk 3: sampled 30,000 | cumulative sampled 90,000
      Chunk 4: sampled 30,000 | cumulative sampled 120,000
      Chunk 5: sampled 11,215 | cumulative sampled 131,215
    
    [20/32] FragPipe_Duncan_PXD004720_pollen_peptide_genome_projection.csv
      Chunk 1: sampled 30,000 | cumulative sampled 30,000
      Chunk 2: sampled 30,000 | cumulative sampled 60,000
      Chunk 3: sampled 14,698 | cumulative sampled 74,698
    
    [21/32] FragPipe_Duncan_PXD004720_rachilla_peptide_genome_projection.csv
      Chunk 1: sampled 30,000 | cumulative sampled 30,000
      Chunk 2: sampled 30,000 | cumulative sampled 60,000
      Chunk 3: sampled 30,000 | cumulative sampled 90,000
      Chunk 4: sampled 30,000 | cumulative sampled 120,000
      Chunk 5: sampled 30,000 | cumulative sampled 150,000
      Chunk 6: sampled 10,204 | cumulative sampled 160,204
    
    [22/32] FragPipe_Duncan_PXD004720_radicle_peptide_genome_projection.csv
      Chunk 1: sampled 30,000 | cumulative sampled 30,000
      Chunk 2: sampled 30,000 | cumulative sampled 60,000
      Chunk 3: sampled 30,000 | cumulative sampled 90,000
      Chunk 4: sampled 30,000 | cumulative sampled 120,000
      Chunk 5: sampled 30,000 | cumulative sampled 150,000
      Chunk 6: sampled 30,000 | cumulative sampled 180,000
      Chunk 7: sampled 14,694 | cumulative sampled 194,694
    
    [23/32] FragPipe_Duncan_PXD004720_root-mature_peptide_genome_projection.csv
      Chunk 1: sampled 30,000 | cumulative sampled 30,000
      Chunk 2: sampled 30,000 | cumulative sampled 60,000
      Chunk 3: sampled 30,000 | cumulative sampled 90,000
      Chunk 4: sampled 7,188 | cumulative sampled 97,188
    
    [24/32] FragPipe_Duncan_PXD004720_root-secretion_peptide_genome_projection.csv
      Chunk 1: sampled 30,000 | cumulative sampled 30,000
      Chunk 2: sampled 30,000 | cumulative sampled 60,000
      Chunk 3: sampled 30,000 | cumulative sampled 90,000
      Chunk 4: sampled 10,639 | cumulative sampled 100,639
    
    [25/32] FragPipe_Duncan_PXD004720_root-tip_peptide_genome_projection.csv
      Chunk 1: sampled 30,000 | cumulative sampled 30,000
      Chunk 2: sampled 30,000 | cumulative sampled 60,000
      Chunk 3: sampled 30,000 | cumulative sampled 90,000
      Chunk 4: sampled 30,000 | cumulative sampled 120,000
      Chunk 5: sampled 30,000 | cumulative sampled 150,000
      Chunk 6: sampled 30,000 | cumulative sampled 180,000
      Chunk 7: sampled 11,943 | cumulative sampled 191,943
    
    [26/32] FragPipe_Duncan_PXD004720_root-vasculature_peptide_genome_projection.csv
      Chunk 1: sampled 30,000 | cumulative sampled 30,000
      Chunk 2: sampled 30,000 | cumulative sampled 60,000
      Chunk 3: sampled 30,000 | cumulative sampled 90,000
      Chunk 4: sampled 21,831 | cumulative sampled 111,831
    
    [27/32] FragPipe_Duncan_PXD004720_spike-immature_peptide_genome_projection.csv
      Chunk 1: sampled 30,000 | cumulative sampled 30,000
      Chunk 2: sampled 30,000 | cumulative sampled 60,000
      Chunk 3: sampled 30,000 | cumulative sampled 90,000
      Chunk 4: sampled 30,000 | cumulative sampled 120,000
      Chunk 5: sampled 30,000 | cumulative sampled 150,000
      Chunk 6: sampled 30,000 | cumulative sampled 180,000
      Chunk 7: sampled 8,826 | cumulative sampled 188,826
    
    [28/32] FragPipe_Duncan_PXD004720_stem_peptide_genome_projection.csv
      Chunk 1: sampled 30,000 | cumulative sampled 30,000
      Chunk 2: sampled 30,000 | cumulative sampled 60,000
      Chunk 3: sampled 734 | cumulative sampled 60,734
    
    [29/32] FragPipe_Liu_PXD050500_coleoptile_peptide_genome_projection.csv
      Chunk 1: sampled 30,000 | cumulative sampled 30,000
      Chunk 2: sampled 30,000 | cumulative sampled 60,000
      Chunk 3: sampled 30,000 | cumulative sampled 90,000
      Chunk 4: sampled 30,000 | cumulative sampled 120,000
      Chunk 5: sampled 30,000 | cumulative sampled 150,000
      Chunk 6: sampled 30,000 | cumulative sampled 180,000
      Chunk 7: sampled 30,000 | cumulative sampled 210,000
      Chunk 8: sampled 30,000 | cumulative sampled 240,000
      Chunk 9: sampled 30,000 | cumulative sampled 270,000
      Chunk 10: sampled 30,000 | cumulative sampled 300,000
      Chunk 11: sampled 30,000 | cumulative sampled 330,000
      Chunk 12: sampled 30,000 | cumulative sampled 360,000
      Chunk 13: sampled 30,000 | cumulative sampled 390,000
      Chunk 14: sampled 30,000 | cumulative sampled 420,000
      Chunk 15: sampled 30,000 | cumulative sampled 450,000
      Chunk 16: sampled 30,000 | cumulative sampled 480,000
      Chunk 17: sampled 30,000 | cumulative sampled 510,000
      Chunk 18: sampled 30,000 | cumulative sampled 540,000
      Chunk 19: sampled 30,000 | cumulative sampled 570,000
      Chunk 20: sampled 30,000 | cumulative sampled 600,000
      Chunk 21: sampled 30,000 | cumulative sampled 630,000
      Chunk 22: sampled 30,000 | cumulative sampled 660,000
      Chunk 23: sampled 30,000 | cumulative sampled 690,000
      Chunk 24: sampled 30,000 | cumulative sampled 720,000
      Chunk 25: sampled 30,000 | cumulative sampled 750,000
      Chunk 26: sampled 30,000 | cumulative sampled 780,000
      Chunk 27: sampled 30,000 | cumulative sampled 810,000
      Chunk 28: sampled 30,000 | cumulative sampled 840,000
      Chunk 29: sampled 30,000 | cumulative sampled 870,000
      Chunk 30: sampled 30,000 | cumulative sampled 900,000
      Chunk 31: sampled 30,000 | cumulative sampled 930,000
      Chunk 32: sampled 30,000 | cumulative sampled 960,000
      Chunk 33: sampled 30,000 | cumulative sampled 990,000
      Chunk 34: sampled 30,000 | cumulative sampled 1,020,000
      Chunk 35: sampled 30,000 | cumulative sampled 1,050,000
      Chunk 36: sampled 30,000 | cumulative sampled 1,080,000
      Chunk 37: sampled 30,000 | cumulative sampled 1,110,000
      Chunk 38: sampled 30,000 | cumulative sampled 1,140,000
      Chunk 39: sampled 30,000 | cumulative sampled 1,170,000
      Chunk 40: sampled 30,000 | cumulative sampled 1,200,000
      Chunk 41: sampled 30,000 | cumulative sampled 1,230,000
      Chunk 42: sampled 30,000 | cumulative sampled 1,260,000
      Chunk 43: sampled 30,000 | cumulative sampled 1,290,000
      Chunk 44: sampled 30,000 | cumulative sampled 1,320,000
      Chunk 45: sampled 30,000 | cumulative sampled 1,350,000
      Chunk 46: sampled 30,000 | cumulative sampled 1,380,000
      Chunk 47: sampled 30,000 | cumulative sampled 1,410,000
      Chunk 48: sampled 30,000 | cumulative sampled 1,440,000
      Chunk 49: sampled 30,000 | cumulative sampled 1,470,000
      Chunk 50: sampled 30,000 | cumulative sampled 1,500,000
      Chunk 51: sampled 30,000 | cumulative sampled 1,530,000
      Chunk 52: sampled 30,000 | cumulative sampled 1,560,000
      Chunk 53: sampled 30,000 | cumulative sampled 1,590,000
      Chunk 54: sampled 30,000 | cumulative sampled 1,620,000
      Chunk 55: sampled 30,000 | cumulative sampled 1,650,000
      Chunk 56: sampled 30,000 | cumulative sampled 1,680,000
      Chunk 57: sampled 30,000 | cumulative sampled 1,710,000
      Chunk 58: sampled 30,000 | cumulative sampled 1,740,000
      Chunk 59: sampled 30,000 | cumulative sampled 1,770,000
      Chunk 60: sampled 30,000 | cumulative sampled 1,800,000
      Chunk 61: sampled 30,000 | cumulative sampled 1,830,000
      Chunk 62: sampled 20,136 | cumulative sampled 1,850,136
    
    [30/32] FragPipe_Liu_PXD050500_node_peptide_genome_projection.csv
      Chunk 1: sampled 30,000 | cumulative sampled 30,000
      Chunk 2: sampled 30,000 | cumulative sampled 60,000
      Chunk 3: sampled 30,000 | cumulative sampled 90,000
      Chunk 4: sampled 30,000 | cumulative sampled 120,000
      Chunk 5: sampled 30,000 | cumulative sampled 150,000
      Chunk 6: sampled 30,000 | cumulative sampled 180,000
      Chunk 7: sampled 30,000 | cumulative sampled 210,000
      Chunk 8: sampled 30,000 | cumulative sampled 240,000
      Chunk 9: sampled 30,000 | cumulative sampled 270,000
      Chunk 10: sampled 30,000 | cumulative sampled 300,000
      Chunk 11: sampled 30,000 | cumulative sampled 330,000
      Chunk 12: sampled 30,000 | cumulative sampled 360,000
      Chunk 13: sampled 30,000 | cumulative sampled 390,000
      Chunk 14: sampled 30,000 | cumulative sampled 420,000
      Chunk 15: sampled 30,000 | cumulative sampled 450,000
      Chunk 16: sampled 30,000 | cumulative sampled 480,000
      Chunk 17: sampled 30,000 | cumulative sampled 510,000
      Chunk 18: sampled 30,000 | cumulative sampled 540,000
      Chunk 19: sampled 30,000 | cumulative sampled 570,000
      Chunk 20: sampled 30,000 | cumulative sampled 600,000
      Chunk 21: sampled 30,000 | cumulative sampled 630,000
      Chunk 22: sampled 30,000 | cumulative sampled 660,000
      Chunk 23: sampled 30,000 | cumulative sampled 690,000
      Chunk 24: sampled 30,000 | cumulative sampled 720,000
      Chunk 25: sampled 30,000 | cumulative sampled 750,000
      Chunk 26: sampled 30,000 | cumulative sampled 780,000
      Chunk 27: sampled 30,000 | cumulative sampled 810,000
      Chunk 28: sampled 30,000 | cumulative sampled 840,000
      Chunk 29: sampled 30,000 | cumulative sampled 870,000
      Chunk 30: sampled 30,000 | cumulative sampled 900,000
      Chunk 31: sampled 30,000 | cumulative sampled 930,000
      Chunk 32: sampled 30,000 | cumulative sampled 960,000
      Chunk 33: sampled 30,000 | cumulative sampled 990,000
      Chunk 34: sampled 30,000 | cumulative sampled 1,020,000
      Chunk 35: sampled 30,000 | cumulative sampled 1,050,000
      Chunk 36: sampled 30,000 | cumulative sampled 1,080,000
      Chunk 37: sampled 30,000 | cumulative sampled 1,110,000
      Chunk 38: sampled 30,000 | cumulative sampled 1,140,000
      Chunk 39: sampled 30,000 | cumulative sampled 1,170,000
      Chunk 40: sampled 30,000 | cumulative sampled 1,200,000
      Chunk 41: sampled 30,000 | cumulative sampled 1,230,000
      Chunk 42: sampled 30,000 | cumulative sampled 1,260,000
      Chunk 43: sampled 30,000 | cumulative sampled 1,290,000
      Chunk 44: sampled 30,000 | cumulative sampled 1,320,000
      Chunk 45: sampled 30,000 | cumulative sampled 1,350,000
      Chunk 46: sampled 30,000 | cumulative sampled 1,380,000
      Chunk 47: sampled 30,000 | cumulative sampled 1,410,000
      Chunk 48: sampled 30,000 | cumulative sampled 1,440,000
      Chunk 49: sampled 30,000 | cumulative sampled 1,470,000
      Chunk 50: sampled 30,000 | cumulative sampled 1,500,000
      Chunk 51: sampled 30,000 | cumulative sampled 1,530,000
      Chunk 52: sampled 30,000 | cumulative sampled 1,560,000
      Chunk 53: sampled 30,000 | cumulative sampled 1,590,000
      Chunk 54: sampled 30,000 | cumulative sampled 1,620,000
      Chunk 55: sampled 30,000 | cumulative sampled 1,650,000
      Chunk 56: sampled 30,000 | cumulative sampled 1,680,000
      Chunk 57: sampled 30,000 | cumulative sampled 1,710,000
      Chunk 58: sampled 30,000 | cumulative sampled 1,740,000
      Chunk 59: sampled 30,000 | cumulative sampled 1,770,000
      Chunk 60: sampled 30,000 | cumulative sampled 1,800,000
      Chunk 61: sampled 30,000 | cumulative sampled 1,830,000
      Chunk 62: sampled 30,000 | cumulative sampled 1,860,000
      Chunk 63: sampled 26,922 | cumulative sampled 1,886,922
    
    [31/32] FragPipe_Liu_PXD050500_radicle_peptide_genome_projection.csv
      Chunk 1: sampled 30,000 | cumulative sampled 30,000
      Chunk 2: sampled 30,000 | cumulative sampled 60,000
      Chunk 3: sampled 30,000 | cumulative sampled 90,000
      Chunk 4: sampled 30,000 | cumulative sampled 120,000
      Chunk 5: sampled 30,000 | cumulative sampled 150,000
      Chunk 6: sampled 30,000 | cumulative sampled 180,000
      Chunk 7: sampled 30,000 | cumulative sampled 210,000
      Chunk 8: sampled 30,000 | cumulative sampled 240,000
      Chunk 9: sampled 30,000 | cumulative sampled 270,000
      Chunk 10: sampled 30,000 | cumulative sampled 300,000
      Chunk 11: sampled 30,000 | cumulative sampled 330,000
      Chunk 12: sampled 30,000 | cumulative sampled 360,000
      Chunk 13: sampled 30,000 | cumulative sampled 390,000
      Chunk 14: sampled 30,000 | cumulative sampled 420,000
      Chunk 15: sampled 30,000 | cumulative sampled 450,000
      Chunk 16: sampled 30,000 | cumulative sampled 480,000
      Chunk 17: sampled 30,000 | cumulative sampled 510,000
      Chunk 18: sampled 30,000 | cumulative sampled 540,000
      Chunk 19: sampled 30,000 | cumulative sampled 570,000
      Chunk 20: sampled 30,000 | cumulative sampled 600,000
      Chunk 21: sampled 30,000 | cumulative sampled 630,000
      Chunk 22: sampled 30,000 | cumulative sampled 660,000
      Chunk 23: sampled 30,000 | cumulative sampled 690,000
      Chunk 24: sampled 30,000 | cumulative sampled 720,000
      Chunk 25: sampled 30,000 | cumulative sampled 750,000
      Chunk 26: sampled 30,000 | cumulative sampled 780,000
      Chunk 27: sampled 30,000 | cumulative sampled 810,000
      Chunk 28: sampled 30,000 | cumulative sampled 840,000
      Chunk 29: sampled 30,000 | cumulative sampled 870,000
      Chunk 30: sampled 30,000 | cumulative sampled 900,000
      Chunk 31: sampled 30,000 | cumulative sampled 930,000
      Chunk 32: sampled 30,000 | cumulative sampled 960,000
      Chunk 33: sampled 30,000 | cumulative sampled 990,000
      Chunk 34: sampled 30,000 | cumulative sampled 1,020,000
      Chunk 35: sampled 30,000 | cumulative sampled 1,050,000
      Chunk 36: sampled 12,508 | cumulative sampled 1,062,508
    
    [32/32] FragPipe_Vincent_MSV000090572_stored-grain_peptide_genome_projection.csv
      Chunk 1: sampled 30,000 | cumulative sampled 30,000
      Chunk 2: sampled 314 | cumulative sampled 30,314
    
    ===== STEP 10 STRATIFIED VALIDATION SUMMARY =====
    


<div>
<style scoped>
    .dataframe tbody tr th:only-of-type {
        vertical-align: middle;
    }

    .dataframe tbody tr th {
        vertical-align: top;
    }

    .dataframe thead th {
        text-align: right;
    }
</style>
<table border="1" class="dataframe">
  <thead>
    <tr style="text-align: right;">
      <th></th>
      <th>Metric</th>
      <th>Value</th>
    </tr>
  </thead>
  <tbody>
    <tr>
      <th>0</th>
      <td>Sampling strategy</td>
      <td>100% stratified per projection file/source-tissue</td>
    </tr>
    <tr>
      <th>1</th>
      <td>Sample fraction</td>
      <td>1</td>
    </tr>
    <tr>
      <th>2</th>
      <td>Projection files validated</td>
      <td>32</td>
    </tr>
    <tr>
      <th>3</th>
      <td>Projected rows available across files</td>
      <td>8291056</td>
    </tr>
    <tr>
      <th>4</th>
      <td>Projected peptide rows validated</td>
      <td>8291056</td>
    </tr>
    <tr>
      <th>5</th>
      <td>Exact translation matches</td>
      <td>8078586</td>
    </tr>
    <tr>
      <th>6</th>
      <td>Exact translation match rate (%)</td>
      <td>97.44</td>
    </tr>
    <tr>
      <th>7</th>
      <td>I/L-normalised translation matches</td>
      <td>8214230</td>
    </tr>
    <tr>
      <th>8</th>
      <td>I/L-normalised translation match rate (%)</td>
      <td>99.07</td>
    </tr>
    <tr>
      <th>9</th>
      <td>Multi-block peptide projections tested</td>
      <td>1100087</td>
    </tr>
    <tr>
      <th>10</th>
      <td>Negative-strand peptide projections tested</td>
      <td>4236479</td>
    </tr>
    <tr>
      <th>11</th>
      <td>Validation status: validated</td>
      <td>8214230</td>
    </tr>
    <tr>
      <th>12</th>
      <td>Validation status: translation_mismatch</td>
      <td>76811</td>
    </tr>
    <tr>
      <th>13</th>
      <td>Validation status: contains_N</td>
      <td>15</td>
    </tr>
  </tbody>
</table>
</div>


    
    Validation table saved: python_outputs\tables\wheat_projection_validation_stratified100percent_step10.csv
    Tissue-level validation summary saved: python_outputs\tables\wheat_projection_validation_100%_tissue_summary_step10.csv
    Overall validation summary saved: python_outputs\tables\wheat_projection_validation_100%_summary_step10.csv
    

# Step 11 — Sanity Checks for Translation-Validated Peptide Genome Projections

This step performs final quality-control checks on the projected peptide genome coordinates before public upload to Apollo/JBrowse.

The aim is to verify that the annotation-guided peptide projections generated in Step 10 and validated by translation are internally consistent and suitable for genome-browser visualisation.

---

Three independent sanity checks are performed:

1. **BED geometry check**  
   Confirms that BED coordinates are valid, that start coordinates are smaller than end coordinates, and that BED block sizes and block starts are consistent with each peptide genomic interval.

2. **Chromosome and strand check**  
   Confirms that projected peptides are assigned only to expected wheat chromosomes and valid strand values (`+` or `-`).

3. **Protein-coordinate consistency check**  
   Confirms that each peptide projection has coherent amino-acid coordinates, including matching peptide length and valid protein coordinate intervals.

For each projected peptide row, the workflow records pass/fail flags for each check and assigns an overall sanity-check status.

---

The outputs include:

- a full sanity-check table for all projected peptide rows,
- a failed-row diagnostic table,
- a tissue-level sanity-check summary table.

---

These checks provide an additional safeguard before sharing the BED tracks as a public genome-browser resource.


```python
# ============================================================
# Step 11 — Sanity checks for translation-validated peptide genome projections (takes 30 min)
# ============================================================

import pandas as pd
from pathlib import Path

# -----------------------------
# 1. Input / output paths
# -----------------------------
fragpipe_dir = Path("FragPipe_results")
tables_dir = Path("python_outputs/tables")
tables_dir.mkdir(parents=True, exist_ok=True)

manifest_file = fragpipe_dir / "wheat_tissues_FragPipe-result-manifest_2026-05-11.csv"

sanity_full_out = tables_dir / "wheat_projection_translation_validated_sanity_checks_full_step11.csv"
sanity_failed_out = tables_dir / "wheat_projection_translation_validated_sanity_checks_failed_rows_step11.csv"
sanity_summary_out = tables_dir / "wheat_projection_translation_validated_sanity_checks_summary_step11.csv"

manifest = pd.read_csv(manifest_file, encoding="utf-8-sig")

# Step 10 output: full translation-validation table
validation_file = tables_dir / "wheat_projection_validation_stratified100percent_step10.csv"

if not validation_file.exists():
    raise FileNotFoundError(
        f"Step 10 translation-validation file not found:\n{validation_file}\n\n"
        "Please run Step 10 first."
    )

# -----------------------------
# 2. Expected wheat chromosomes and valid strands
# -----------------------------
valid_chromosomes = {
    "Chr1A", "Chr1B", "Chr1D",
    "Chr2A", "Chr2B", "Chr2D",
    "Chr3A", "Chr3B", "Chr3D",
    "Chr4A", "Chr4B", "Chr4D",
    "Chr5A", "Chr5B", "Chr5D",
    "Chr6A", "Chr6B", "Chr6D",
    "Chr7A", "Chr7B", "Chr7D"
}

valid_strands = {"+", "-"}

# -----------------------------
# 3. Helper functions
# -----------------------------
def parse_bed_list(value):
    """
    Convert comma-separated BED block size/start strings into integer lists.
    Handles empty or missing values.
    """
    if pd.isna(value):
        return []

    value = str(value).strip().strip(",")

    if value == "":
        return []

    try:
        return [int(x) for x in value.split(",") if str(x).strip() != ""]
    except ValueError:
        return []


def check_bed_geometry(row):
    """
    Check BED interval and BED12-style block geometry.
    """
    try:
        bed_start = int(row["BED_start_0based"])
        bed_end = int(row["BED_end_0based_exclusive"])
        block_count = int(row["BED_block_count"])
    except Exception:
        return False

    if bed_start < 0:
        return False

    if bed_end <= bed_start:
        return False

    block_sizes = parse_bed_list(row["BED_block_sizes"])
    block_starts = parse_bed_list(row["BED_block_starts"])

    if block_count <= 0:
        return False

    if len(block_sizes) != block_count:
        return False

    if len(block_starts) != block_count:
        return False

    if any(size <= 0 for size in block_sizes):
        return False

    if any(start < 0 for start in block_starts):
        return False

    interval_length = bed_end - bed_start

    # Every block must fit inside the BED interval
    for block_start, block_size in zip(block_starts, block_sizes):
        if block_start + block_size > interval_length:
            return False

    return True


def check_block_nt_length(row):
    """
    Check that the sum of BED block nucleotide lengths matches peptide length × 3.
    """
    try:
        peptide_len = int(row["Peptide_length_AA"])
    except Exception:
        return False

    block_sizes = parse_bed_list(row["BED_block_sizes"])

    if len(block_sizes) == 0:
        return False

    return sum(block_sizes) == peptide_len * 3


def check_chromosome_and_strand(row):
    """
    Check valid chromosome and strand assignment.
    """
    chrom = str(row["Chromosome"])
    strand = str(row["Strand"])

    return chrom in valid_chromosomes and strand in valid_strands


def check_protein_coordinates(row):
    """
    Check that amino-acid coordinates are internally consistent with peptide length.
    """
    try:
        aa_start = int(row["AA_start"])
        aa_end = int(row["AA_end"])
        peptide_len = int(row["Peptide_length_AA"])
    except Exception:
        return False

    if aa_start <= 0:
        return False

    if aa_end < aa_start:
        return False

    if (aa_end - aa_start + 1) != peptide_len:
        return False

    peptide = str(row["Peptide"])

    if len(peptide) != peptide_len:
        return False

    return True

# -----------------------------
# 4. Run sanity checks across translation-validated Step 10 output
# -----------------------------

print("\nRunning sanity checks on Step 10 translation-validated projections...")

chunk_size = 100_000

# Clear previous outputs if rerunning
for out_file in [sanity_full_out, sanity_failed_out, sanity_summary_out]:
    if out_file.exists():
        out_file.unlink()

# Build lookup from Step 9 projection filename to manifest metadata
manifest_lookup = {}

for _, manifest_row in manifest.iterrows():

    projection_filename = manifest_row["FragPipe-Output-Peptide"].replace(
        "_peptide.tsv",
        "_peptide_genome_projection.csv"
    )

    manifest_lookup[projection_filename] = {
        "Source": manifest_row["Source"],
        "Species": manifest_row["Species"],
        "Tissue": manifest_row["Tissue-Raw-Code"],
        "Batch": manifest_row["Batch"]
    }


def detect_source_file_column(chunk):
    """
    Detect the column containing the original Step 9 projection filename.

    Step 10 intended to write '_source_file', but pandas itertuples()
    may rename columns beginning with '_' into positional names.
    This helper first checks for '_source_file', then searches for a
    column containing values ending in '_peptide_genome_projection.csv'.
    """

    if "_source_file" in chunk.columns:
        return "_source_file"

    for col in chunk.columns:
        sample_values = (
            chunk[col]
            .dropna()
            .astype(str)
            .head(20)
        )

        if sample_values.str.contains(
            "_peptide_genome_projection.csv",
            regex=False
        ).any():
            return col

    return None


summary_dict = {}
header_written_full = False
header_written_failed = False
first_failed_examples = []
source_file_col = None

total_rows_read = 0
total_translation_validated = 0
total_translation_excluded = 0

for chunk_i, chunk in enumerate(
    pd.read_csv(
        validation_file,
        chunksize=chunk_size,
        low_memory=False
    ),
    start=1
):

    total_rows_read += len(chunk)

    # Detect source-file column once, then reuse it
    if source_file_col is None:
        source_file_col = detect_source_file_column(chunk)

        if source_file_col is None:
            raise KeyError(
                "Could not identify the Step 10 source-file column. "
                "Expected '_source_file' or a column containing values ending with "
                "'_peptide_genome_projection.csv'."
            )

        print(f"Detected Step 10 source-file column: {source_file_col}")

    if "Validation_status" not in chunk.columns:
        raise KeyError(
            "Column 'Validation_status' was not found in the Step 10 validation table."
        )

    # Count all Step 10 rows by source file and translation status
    for projection_filename, file_group in chunk.groupby(source_file_col):

        projection_filename = str(projection_filename)

        if projection_filename not in summary_dict:
            meta = manifest_lookup.get(
                projection_filename,
                {
                    "Source": pd.NA,
                    "Species": pd.NA,
                    "Tissue": pd.NA,
                    "Batch": pd.NA
                }
            )

            summary_dict[projection_filename] = {
                "Source": meta["Source"],
                "Species": meta["Species"],
                "Tissue": meta["Tissue"],
                "Batch": meta["Batch"],
                "Projection_file": projection_filename,

                "Rows_from_step10_validation_table": 0,
                "Rows_translation_validated": 0,
                "Rows_excluded_by_translation_validation": 0,

                "Translation_validated_rows_checked": 0,
                "Rows_passing_all_sanity_checks": 0,
                "Rows_failing_any_sanity_check": 0,

                "BED_geometry_failures": 0,
                "Block_nt_length_failures": 0,
                "Chromosome_strand_failures": 0,
                "Protein_coordinate_failures": 0
            }

        summary_dict[projection_filename]["Rows_from_step10_validation_table"] += len(file_group)

        n_validated = int((file_group["Validation_status"].astype(str) == "validated").sum())
        n_excluded = len(file_group) - n_validated

        summary_dict[projection_filename]["Rows_translation_validated"] += n_validated
        summary_dict[projection_filename]["Rows_excluded_by_translation_validation"] += n_excluded

    # Keep only translation-validated rows for sanity checks
    projected = chunk[
        chunk["Validation_status"].astype(str) == "validated"
    ].copy()

    total_translation_validated += len(projected)
    total_translation_excluded += len(chunk) - len(projected)

    if projected.empty:
        print(
            f"Chunk {chunk_i}: read {len(chunk):,} rows | "
            f"no translation-validated rows"
        )
        continue

    # Derive peptide length if not already present
    if "Peptide_length_AA" not in projected.columns:
        if "Original_peptide_clean" in projected.columns:
            projected["Peptide_length_AA"] = (
                projected["Original_peptide_clean"]
                .astype(str)
                .str.len()
            )
        else:
            projected["Peptide_length_AA"] = (
                projected["Peptide"]
                .astype(str)
                .str.len()
            )

    # Add/standardise projection file metadata
    projected["Projection_file"] = projected[source_file_col].astype(str)

    projected["Source"] = projected["Projection_file"].map(
        lambda x: manifest_lookup.get(x, {}).get("Source", pd.NA)
    )
    projected["Species"] = projected["Projection_file"].map(
        lambda x: manifest_lookup.get(x, {}).get("Species", pd.NA)
    )
    projected["Tissue"] = projected["Projection_file"].map(
        lambda x: manifest_lookup.get(x, {}).get("Tissue", pd.NA)
    )
    projected["Batch"] = projected["Projection_file"].map(
        lambda x: manifest_lookup.get(x, {}).get("Batch", pd.NA)
    )

    # Sanity checks
    projected["Check_BED_geometry"] = projected.apply(check_bed_geometry, axis=1)
    projected["Check_block_nt_length"] = projected.apply(check_block_nt_length, axis=1)
    projected["Check_chromosome_and_strand"] = projected.apply(check_chromosome_and_strand, axis=1)
    projected["Check_protein_coordinates"] = projected.apply(check_protein_coordinates, axis=1)

    check_cols = [
        "Check_BED_geometry",
        "Check_block_nt_length",
        "Check_chromosome_and_strand",
        "Check_protein_coordinates"
    ]

    projected["All_sanity_checks_passed"] = projected[check_cols].all(axis=1)

    projected["Sanity_check_status"] = projected["All_sanity_checks_passed"].map({
        True: "passed",
        False: "failed"
    })

    # Update summary counts by source projection file
    for projection_filename, file_group in projected.groupby("Projection_file"):

        projection_filename = str(projection_filename)

        if projection_filename not in summary_dict:
            meta = manifest_lookup.get(
                projection_filename,
                {
                    "Source": pd.NA,
                    "Species": pd.NA,
                    "Tissue": pd.NA,
                    "Batch": pd.NA
                }
            )

            summary_dict[projection_filename] = {
                "Source": meta["Source"],
                "Species": meta["Species"],
                "Tissue": meta["Tissue"],
                "Batch": meta["Batch"],
                "Projection_file": projection_filename,

                "Rows_from_step10_validation_table": 0,
                "Rows_translation_validated": 0,
                "Rows_excluded_by_translation_validation": 0,

                "Translation_validated_rows_checked": 0,
                "Rows_passing_all_sanity_checks": 0,
                "Rows_failing_any_sanity_check": 0,

                "BED_geometry_failures": 0,
                "Block_nt_length_failures": 0,
                "Chromosome_strand_failures": 0,
                "Protein_coordinate_failures": 0
            }

        summary_dict[projection_filename]["Translation_validated_rows_checked"] += len(file_group)
        summary_dict[projection_filename]["Rows_passing_all_sanity_checks"] += int(
            file_group["All_sanity_checks_passed"].sum()
        )
        summary_dict[projection_filename]["Rows_failing_any_sanity_check"] += int(
            (~file_group["All_sanity_checks_passed"]).sum()
        )

        summary_dict[projection_filename]["BED_geometry_failures"] += int(
            (~file_group["Check_BED_geometry"]).sum()
        )
        summary_dict[projection_filename]["Block_nt_length_failures"] += int(
            (~file_group["Check_block_nt_length"]).sum()
        )
        summary_dict[projection_filename]["Chromosome_strand_failures"] += int(
            (~file_group["Check_chromosome_and_strand"]).sum()
        )
        summary_dict[projection_filename]["Protein_coordinate_failures"] += int(
            (~file_group["Check_protein_coordinates"]).sum()
        )

    # Write full sanity-check output incrementally
    projected.to_csv(
        sanity_full_out,
        index=False,
        mode="a",
        header=not header_written_full
    )

    header_written_full = True

    # Write failed rows incrementally
    failed = projected[
        projected["Sanity_check_status"] == "failed"
    ].copy()

    if not failed.empty:

        failed.to_csv(
            sanity_failed_out,
            index=False,
            mode="a",
            header=not header_written_failed
        )

        header_written_failed = True

        if len(first_failed_examples) < 20:
            remaining = 20 - len(first_failed_examples)
            first_failed_examples.append(failed.head(remaining))

    print(
        f"Chunk {chunk_i}: read {len(chunk):,} rows | "
        f"translation-validated {len(projected):,} | "
        f"sanity failed {len(failed):,}"
    )


# -----------------------------
# 5. Build and export summary table
# -----------------------------
if not header_written_full:
    raise ValueError(
        "No translation-validated projected peptide rows were available for sanity checking."
    )

sanity_summary = pd.DataFrame(summary_dict.values())

# Add percentages
sanity_summary["Percent_translation_validated"] = (
    sanity_summary["Rows_translation_validated"] /
    sanity_summary["Rows_from_step10_validation_table"] *
    100
).round(4)

sanity_summary["Percent_passing_all_sanity_checks"] = (
    sanity_summary["Rows_passing_all_sanity_checks"] /
    sanity_summary["Translation_validated_rows_checked"] *
    100
).round(4)

sanity_summary.to_csv(sanity_summary_out, index=False)

# Build a small failed-row preview for display
if len(first_failed_examples) > 0:
    sanity_failed_preview = pd.concat(first_failed_examples, ignore_index=True).head(20)
else:
    sanity_failed_preview = pd.DataFrame()


# -----------------------------
# 6. Overall summary
# -----------------------------
overall_checked = int(sanity_summary["Translation_validated_rows_checked"].sum())
overall_passed = int(sanity_summary["Rows_passing_all_sanity_checks"].sum())
overall_failed = int(sanity_summary["Rows_failing_any_sanity_check"].sum())

overall_pass_percent = round(
    (overall_passed / overall_checked) * 100,
    4
) if overall_checked > 0 else pd.NA

print("\n===== STEP 11 SANITY CHECK SUMMARY =====")
print(f"Rows read from Step 10 validation table: {total_rows_read:,}")
print(f"Rows excluded by translation validation: {total_translation_excluded:,}")
print(f"Translation-validated rows checked: {overall_checked:,}")
print(f"Rows passing all sanity checks: {overall_passed:,}")
print(f"Rows failing at least one sanity check: {overall_failed:,}")
print(f"Overall sanity-check pass rate among translation-validated rows: {overall_pass_percent}%")

print(f"\nFull sanity-check table saved: {sanity_full_out}")
print(f"Failed-row diagnostic table saved: {sanity_failed_out}")
print(f"Tissue-level sanity summary saved: {sanity_summary_out}")

display(sanity_summary)

if not sanity_failed_preview.empty:
    display(sanity_failed_preview)
else:
    print("\nNo failed sanity-check rows to display.")
```

    
    Running sanity checks on Step 10 translation-validated projections...
    Detected Step 10 source-file column: _22
    Chunk 1: read 100,000 rows | translation-validated 99,476 | sanity failed 890
    Chunk 2: read 100,000 rows | translation-validated 99,426 | sanity failed 953
    Chunk 3: read 100,000 rows | translation-validated 99,342 | sanity failed 1,023
    Chunk 4: read 100,000 rows | translation-validated 99,392 | sanity failed 933
    Chunk 5: read 100,000 rows | translation-validated 99,450 | sanity failed 840
    Chunk 6: read 100,000 rows | translation-validated 99,285 | sanity failed 1,169
    Chunk 7: read 100,000 rows | translation-validated 99,365 | sanity failed 954
    Chunk 8: read 100,000 rows | translation-validated 99,354 | sanity failed 875
    Chunk 9: read 100,000 rows | translation-validated 99,505 | sanity failed 764
    Chunk 10: read 100,000 rows | translation-validated 99,501 | sanity failed 763
    Chunk 11: read 100,000 rows | translation-validated 99,434 | sanity failed 809
    Chunk 12: read 100,000 rows | translation-validated 99,319 | sanity failed 923
    Chunk 13: read 100,000 rows | translation-validated 99,437 | sanity failed 793
    Chunk 14: read 100,000 rows | translation-validated 99,306 | sanity failed 1,059
    Chunk 15: read 100,000 rows | translation-validated 99,132 | sanity failed 1,178
    Chunk 16: read 100,000 rows | translation-validated 99,263 | sanity failed 1,095
    Chunk 17: read 100,000 rows | translation-validated 99,256 | sanity failed 997
    Chunk 18: read 100,000 rows | translation-validated 99,369 | sanity failed 942
    Chunk 19: read 100,000 rows | translation-validated 99,286 | sanity failed 1,045
    Chunk 20: read 100,000 rows | translation-validated 98,989 | sanity failed 1,118
    Chunk 21: read 100,000 rows | translation-validated 99,334 | sanity failed 1,102
    Chunk 22: read 100,000 rows | translation-validated 99,505 | sanity failed 1,012
    Chunk 23: read 100,000 rows | translation-validated 99,561 | sanity failed 915
    Chunk 24: read 100,000 rows | translation-validated 99,464 | sanity failed 951
    Chunk 25: read 100,000 rows | translation-validated 99,373 | sanity failed 1,025
    Chunk 26: read 100,000 rows | translation-validated 99,536 | sanity failed 828
    Chunk 27: read 100,000 rows | translation-validated 99,638 | sanity failed 745
    Chunk 28: read 100,000 rows | translation-validated 99,215 | sanity failed 1,062
    Chunk 29: read 100,000 rows | translation-validated 99,444 | sanity failed 969
    Chunk 30: read 100,000 rows | translation-validated 99,562 | sanity failed 720
    Chunk 31: read 100,000 rows | translation-validated 99,724 | sanity failed 633
    Chunk 32: read 100,000 rows | translation-validated 99,533 | sanity failed 896
    Chunk 33: read 100,000 rows | translation-validated 99,557 | sanity failed 900
    Chunk 34: read 100,000 rows | translation-validated 99,593 | sanity failed 850
    Chunk 35: read 100,000 rows | translation-validated 99,263 | sanity failed 857
    Chunk 36: read 100,000 rows | translation-validated 99,161 | sanity failed 839
    Chunk 37: read 100,000 rows | translation-validated 98,841 | sanity failed 901
    Chunk 38: read 100,000 rows | translation-validated 98,936 | sanity failed 804
    Chunk 39: read 100,000 rows | translation-validated 98,967 | sanity failed 874
    Chunk 40: read 100,000 rows | translation-validated 98,988 | sanity failed 973
    Chunk 41: read 100,000 rows | translation-validated 98,863 | sanity failed 1,010
    Chunk 42: read 100,000 rows | translation-validated 98,549 | sanity failed 993
    Chunk 43: read 100,000 rows | translation-validated 98,710 | sanity failed 905
    Chunk 44: read 100,000 rows | translation-validated 98,681 | sanity failed 844
    Chunk 45: read 100,000 rows | translation-validated 99,065 | sanity failed 1,055
    Chunk 46: read 100,000 rows | translation-validated 98,648 | sanity failed 1,214
    Chunk 47: read 100,000 rows | translation-validated 98,674 | sanity failed 1,020
    Chunk 48: read 100,000 rows | translation-validated 98,781 | sanity failed 949
    Chunk 49: read 100,000 rows | translation-validated 98,855 | sanity failed 975
    Chunk 50: read 100,000 rows | translation-validated 98,971 | sanity failed 841
    Chunk 51: read 100,000 rows | translation-validated 99,035 | sanity failed 855
    Chunk 52: read 100,000 rows | translation-validated 99,018 | sanity failed 839
    Chunk 53: read 100,000 rows | translation-validated 98,620 | sanity failed 860
    Chunk 54: read 100,000 rows | translation-validated 99,201 | sanity failed 858
    Chunk 55: read 100,000 rows | translation-validated 98,586 | sanity failed 984
    Chunk 56: read 100,000 rows | translation-validated 98,751 | sanity failed 870
    Chunk 57: read 100,000 rows | translation-validated 98,933 | sanity failed 858
    Chunk 58: read 100,000 rows | translation-validated 99,051 | sanity failed 899
    Chunk 59: read 100,000 rows | translation-validated 98,832 | sanity failed 942
    Chunk 60: read 100,000 rows | translation-validated 98,643 | sanity failed 1,067
    Chunk 61: read 100,000 rows | translation-validated 98,680 | sanity failed 936
    Chunk 62: read 100,000 rows | translation-validated 98,679 | sanity failed 983
    Chunk 63: read 100,000 rows | translation-validated 98,817 | sanity failed 960
    Chunk 64: read 100,000 rows | translation-validated 98,927 | sanity failed 1,156
    Chunk 65: read 100,000 rows | translation-validated 98,459 | sanity failed 1,135
    Chunk 66: read 100,000 rows | translation-validated 98,605 | sanity failed 1,063
    Chunk 67: read 100,000 rows | translation-validated 98,867 | sanity failed 906
    Chunk 68: read 100,000 rows | translation-validated 98,829 | sanity failed 932
    Chunk 69: read 100,000 rows | translation-validated 98,929 | sanity failed 828
    Chunk 70: read 100,000 rows | translation-validated 98,927 | sanity failed 757
    Chunk 71: read 100,000 rows | translation-validated 98,990 | sanity failed 874
    Chunk 72: read 100,000 rows | translation-validated 98,661 | sanity failed 928
    Chunk 73: read 100,000 rows | translation-validated 99,181 | sanity failed 848
    Chunk 74: read 100,000 rows | translation-validated 98,771 | sanity failed 876
    Chunk 75: read 100,000 rows | translation-validated 98,701 | sanity failed 905
    Chunk 76: read 100,000 rows | translation-validated 98,660 | sanity failed 960
    Chunk 77: read 100,000 rows | translation-validated 98,836 | sanity failed 944
    Chunk 78: read 100,000 rows | translation-validated 98,863 | sanity failed 939
    Chunk 79: read 100,000 rows | translation-validated 98,473 | sanity failed 1,148
    Chunk 80: read 100,000 rows | translation-validated 98,816 | sanity failed 1,001
    Chunk 81: read 100,000 rows | translation-validated 98,978 | sanity failed 788
    Chunk 82: read 100,000 rows | translation-validated 99,176 | sanity failed 846
    Chunk 83: read 91,056 rows | translation-validated 89,856 | sanity failed 1,013
    
    ===== STEP 11 SANITY CHECK SUMMARY =====
    Rows read from Step 10 validation table: 8,291,056
    Rows excluded by translation validation: 76,826
    Translation-validated rows checked: 8,214,230
    Rows passing all sanity checks: 8,136,687
    Rows failing at least one sanity check: 77,543
    Overall sanity-check pass rate among translation-validated rows: 99.056%
    
    Full sanity-check table saved: python_outputs\tables\wheat_projection_translation_validated_sanity_checks_full_step11.csv
    Failed-row diagnostic table saved: python_outputs\tables\wheat_projection_translation_validated_sanity_checks_failed_rows_step11.csv
    Tissue-level sanity summary saved: python_outputs\tables\wheat_projection_translation_validated_sanity_checks_summary_step11.csv
    


<div>
<style scoped>
    .dataframe tbody tr th:only-of-type {
        vertical-align: middle;
    }

    .dataframe tbody tr th {
        vertical-align: top;
    }

    .dataframe thead th {
        text-align: right;
    }
</style>
<table border="1" class="dataframe">
  <thead>
    <tr style="text-align: right;">
      <th></th>
      <th>Source</th>
      <th>Species</th>
      <th>Tissue</th>
      <th>Batch</th>
      <th>Projection_file</th>
      <th>Rows_from_step10_validation_table</th>
      <th>Rows_translation_validated</th>
      <th>Rows_excluded_by_translation_validation</th>
      <th>Translation_validated_rows_checked</th>
      <th>Rows_passing_all_sanity_checks</th>
      <th>Rows_failing_any_sanity_check</th>
      <th>BED_geometry_failures</th>
      <th>Block_nt_length_failures</th>
      <th>Chromosome_strand_failures</th>
      <th>Protein_coordinate_failures</th>
      <th>Percent_translation_validated</th>
      <th>Percent_passing_all_sanity_checks</th>
    </tr>
  </thead>
  <tbody>
    <tr>
      <th>0</th>
      <td>PXD004720</td>
      <td>bread wheat</td>
      <td>anther</td>
      <td>single</td>
      <td>FragPipe_Duncan_PXD004720_anther_peptide_genom...</td>
      <td>164365</td>
      <td>163467</td>
      <td>898</td>
      <td>163467</td>
      <td>161955</td>
      <td>1512</td>
      <td>0</td>
      <td>0</td>
      <td>1512</td>
      <td>0</td>
      <td>99.4537</td>
      <td>99.0750</td>
    </tr>
    <tr>
      <th>1</th>
      <td>PXD004720</td>
      <td>bread wheat</td>
      <td>boot</td>
      <td>single</td>
      <td>FragPipe_Duncan_PXD004720_boot_peptide_genome_...</td>
      <td>13227</td>
      <td>13137</td>
      <td>90</td>
      <td>13137</td>
      <td>13024</td>
      <td>113</td>
      <td>0</td>
      <td>0</td>
      <td>113</td>
      <td>0</td>
      <td>99.3196</td>
      <td>99.1398</td>
    </tr>
    <tr>
      <th>2</th>
      <td>PXD004720</td>
      <td>bread wheat</td>
      <td>coleoptile</td>
      <td>single</td>
      <td>FragPipe_Duncan_PXD004720_coleoptile_peptide_g...</td>
      <td>204476</td>
      <td>203264</td>
      <td>1212</td>
      <td>203264</td>
      <td>201259</td>
      <td>2005</td>
      <td>0</td>
      <td>0</td>
      <td>2005</td>
      <td>0</td>
      <td>99.4073</td>
      <td>99.0136</td>
    </tr>
    <tr>
      <th>3</th>
      <td>PXD004720</td>
      <td>bread wheat</td>
      <td>embryo</td>
      <td>single</td>
      <td>FragPipe_Duncan_PXD004720_embryo_peptide_genom...</td>
      <td>8852</td>
      <td>8742</td>
      <td>110</td>
      <td>8742</td>
      <td>8662</td>
      <td>80</td>
      <td>0</td>
      <td>0</td>
      <td>80</td>
      <td>0</td>
      <td>98.7573</td>
      <td>99.0849</td>
    </tr>
    <tr>
      <th>4</th>
      <td>PXD004720</td>
      <td>bread wheat</td>
      <td>endosperm</td>
      <td>single</td>
      <td>FragPipe_Duncan_PXD004720_endosperm_peptide_ge...</td>
      <td>102830</td>
      <td>102289</td>
      <td>541</td>
      <td>102289</td>
      <td>101413</td>
      <td>876</td>
      <td>0</td>
      <td>0</td>
      <td>876</td>
      <td>0</td>
      <td>99.4739</td>
      <td>99.1436</td>
    </tr>
    <tr>
      <th>5</th>
      <td>PXD004720</td>
      <td>bread wheat</td>
      <td>glume</td>
      <td>single</td>
      <td>FragPipe_Duncan_PXD004720_glume_peptide_genome...</td>
      <td>145133</td>
      <td>144079</td>
      <td>1054</td>
      <td>144079</td>
      <td>142449</td>
      <td>1630</td>
      <td>0</td>
      <td>0</td>
      <td>1630</td>
      <td>0</td>
      <td>99.2738</td>
      <td>98.8687</td>
    </tr>
    <tr>
      <th>6</th>
      <td>PXD004720</td>
      <td>bread wheat</td>
      <td>grain-zadoks-70</td>
      <td>single</td>
      <td>FragPipe_Duncan_PXD004720_grain-zadoks-70_pept...</td>
      <td>126195</td>
      <td>125355</td>
      <td>840</td>
      <td>125355</td>
      <td>124251</td>
      <td>1104</td>
      <td>0</td>
      <td>0</td>
      <td>1104</td>
      <td>0</td>
      <td>99.3344</td>
      <td>99.1193</td>
    </tr>
    <tr>
      <th>7</th>
      <td>PXD004720</td>
      <td>bread wheat</td>
      <td>grain-zadoks-71</td>
      <td>single</td>
      <td>FragPipe_Duncan_PXD004720_grain-zadoks-71_pept...</td>
      <td>179725</td>
      <td>178814</td>
      <td>911</td>
      <td>178814</td>
      <td>177392</td>
      <td>1422</td>
      <td>0</td>
      <td>0</td>
      <td>1422</td>
      <td>0</td>
      <td>99.4931</td>
      <td>99.2048</td>
    </tr>
    <tr>
      <th>8</th>
      <td>PXD004720</td>
      <td>bread wheat</td>
      <td>grain-zadoks-75</td>
      <td>single</td>
      <td>FragPipe_Duncan_PXD004720_grain-zadoks-75_pept...</td>
      <td>132701</td>
      <td>132081</td>
      <td>620</td>
      <td>132081</td>
      <td>131066</td>
      <td>1015</td>
      <td>0</td>
      <td>0</td>
      <td>1015</td>
      <td>0</td>
      <td>99.5328</td>
      <td>99.2315</td>
    </tr>
    <tr>
      <th>9</th>
      <td>PXD004720</td>
      <td>bread wheat</td>
      <td>grain-zadoks-83</td>
      <td>single</td>
      <td>FragPipe_Duncan_PXD004720_grain-zadoks-83_pept...</td>
      <td>113203</td>
      <td>112389</td>
      <td>814</td>
      <td>112389</td>
      <td>111335</td>
      <td>1054</td>
      <td>0</td>
      <td>0</td>
      <td>1054</td>
      <td>0</td>
      <td>99.2809</td>
      <td>99.0622</td>
    </tr>
    <tr>
      <th>10</th>
      <td>PXD004720</td>
      <td>bread wheat</td>
      <td>grain-zadoks-87</td>
      <td>single</td>
      <td>FragPipe_Duncan_PXD004720_grain-zadoks-87_pept...</td>
      <td>111792</td>
      <td>111156</td>
      <td>636</td>
      <td>111156</td>
      <td>110256</td>
      <td>900</td>
      <td>0</td>
      <td>0</td>
      <td>900</td>
      <td>0</td>
      <td>99.4311</td>
      <td>99.1903</td>
    </tr>
    <tr>
      <th>11</th>
      <td>PXD004720</td>
      <td>bread wheat</td>
      <td>leaf-flag-mature</td>
      <td>single</td>
      <td>FragPipe_Duncan_PXD004720_leaf-flag-mature_pep...</td>
      <td>145632</td>
      <td>144641</td>
      <td>991</td>
      <td>144641</td>
      <td>143118</td>
      <td>1523</td>
      <td>0</td>
      <td>0</td>
      <td>1523</td>
      <td>0</td>
      <td>99.3195</td>
      <td>98.9470</td>
    </tr>
    <tr>
      <th>12</th>
      <td>PXD004720</td>
      <td>bread wheat</td>
      <td>leaf-flag-senescing</td>
      <td>single</td>
      <td>FragPipe_Duncan_PXD004720_leaf-flag-senescing_...</td>
      <td>46200</td>
      <td>45680</td>
      <td>520</td>
      <td>45680</td>
      <td>45054</td>
      <td>626</td>
      <td>0</td>
      <td>0</td>
      <td>626</td>
      <td>0</td>
      <td>98.8745</td>
      <td>98.6296</td>
    </tr>
    <tr>
      <th>13</th>
      <td>PXD004720</td>
      <td>bread wheat</td>
      <td>leaf-flag-young</td>
      <td>single</td>
      <td>FragPipe_Duncan_PXD004720_leaf-flag-young_pept...</td>
      <td>120821</td>
      <td>119922</td>
      <td>899</td>
      <td>119922</td>
      <td>118601</td>
      <td>1321</td>
      <td>0</td>
      <td>0</td>
      <td>1321</td>
      <td>0</td>
      <td>99.2559</td>
      <td>98.8985</td>
    </tr>
    <tr>
      <th>14</th>
      <td>PXD004720</td>
      <td>bread wheat</td>
      <td>lemma</td>
      <td>single</td>
      <td>FragPipe_Duncan_PXD004720_lemma_peptide_genome...</td>
      <td>147618</td>
      <td>146562</td>
      <td>1056</td>
      <td>146562</td>
      <td>145101</td>
      <td>1461</td>
      <td>0</td>
      <td>0</td>
      <td>1461</td>
      <td>0</td>
      <td>99.2846</td>
      <td>99.0032</td>
    </tr>
    <tr>
      <th>15</th>
      <td>PXD004720</td>
      <td>bread wheat</td>
      <td>node_secretion</td>
      <td>single</td>
      <td>FragPipe_Duncan_PXD004720_node-secretion_pepti...</td>
      <td>169675</td>
      <td>168564</td>
      <td>1111</td>
      <td>168564</td>
      <td>166896</td>
      <td>1668</td>
      <td>0</td>
      <td>0</td>
      <td>1668</td>
      <td>0</td>
      <td>99.3452</td>
      <td>99.0105</td>
    </tr>
    <tr>
      <th>16</th>
      <td>PXD004720</td>
      <td>bread wheat</td>
      <td>node</td>
      <td>single</td>
      <td>FragPipe_Duncan_PXD004720_node_peptide_genome_...</td>
      <td>99823</td>
      <td>98731</td>
      <td>1092</td>
      <td>98731</td>
      <td>97553</td>
      <td>1178</td>
      <td>0</td>
      <td>0</td>
      <td>1178</td>
      <td>0</td>
      <td>98.9061</td>
      <td>98.8069</td>
    </tr>
    <tr>
      <th>17</th>
      <td>PXD004720</td>
      <td>bread wheat</td>
      <td>palea</td>
      <td>single</td>
      <td>FragPipe_Duncan_PXD004720_palea_peptide_genome...</td>
      <td>116936</td>
      <td>116285</td>
      <td>651</td>
      <td>116285</td>
      <td>115047</td>
      <td>1238</td>
      <td>0</td>
      <td>0</td>
      <td>1238</td>
      <td>0</td>
      <td>99.4433</td>
      <td>98.9354</td>
    </tr>
    <tr>
      <th>18</th>
      <td>PXD004720</td>
      <td>bread wheat</td>
      <td>pericarp</td>
      <td>single</td>
      <td>FragPipe_Duncan_PXD004720_pericarp_peptide_gen...</td>
      <td>131215</td>
      <td>130645</td>
      <td>570</td>
      <td>130645</td>
      <td>129415</td>
      <td>1230</td>
      <td>0</td>
      <td>0</td>
      <td>1230</td>
      <td>0</td>
      <td>99.5656</td>
      <td>99.0585</td>
    </tr>
    <tr>
      <th>19</th>
      <td>PXD004720</td>
      <td>bread wheat</td>
      <td>pollen</td>
      <td>single</td>
      <td>FragPipe_Duncan_PXD004720_pollen_peptide_genom...</td>
      <td>74698</td>
      <td>74334</td>
      <td>364</td>
      <td>74334</td>
      <td>73586</td>
      <td>748</td>
      <td>0</td>
      <td>0</td>
      <td>748</td>
      <td>0</td>
      <td>99.5127</td>
      <td>98.9937</td>
    </tr>
    <tr>
      <th>20</th>
      <td>PXD004720</td>
      <td>bread wheat</td>
      <td>rachilla</td>
      <td>single</td>
      <td>FragPipe_Duncan_PXD004720_rachilla_peptide_gen...</td>
      <td>160204</td>
      <td>159219</td>
      <td>985</td>
      <td>159219</td>
      <td>157648</td>
      <td>1571</td>
      <td>0</td>
      <td>0</td>
      <td>1571</td>
      <td>0</td>
      <td>99.3852</td>
      <td>99.0133</td>
    </tr>
    <tr>
      <th>21</th>
      <td>PXD004720</td>
      <td>bread wheat</td>
      <td>radicle</td>
      <td>single</td>
      <td>FragPipe_Duncan_PXD004720_radicle_peptide_geno...</td>
      <td>194694</td>
      <td>193938</td>
      <td>756</td>
      <td>193938</td>
      <td>192432</td>
      <td>1506</td>
      <td>0</td>
      <td>0</td>
      <td>1506</td>
      <td>0</td>
      <td>99.6117</td>
      <td>99.2235</td>
    </tr>
    <tr>
      <th>22</th>
      <td>PXD004720</td>
      <td>bread wheat</td>
      <td>root-mature</td>
      <td>single</td>
      <td>FragPipe_Duncan_PXD004720_root-mature_peptide_...</td>
      <td>97188</td>
      <td>96387</td>
      <td>801</td>
      <td>96387</td>
      <td>95329</td>
      <td>1058</td>
      <td>0</td>
      <td>0</td>
      <td>1058</td>
      <td>0</td>
      <td>99.1758</td>
      <td>98.9023</td>
    </tr>
    <tr>
      <th>23</th>
      <td>PXD004720</td>
      <td>bread wheat</td>
      <td>root-secretion</td>
      <td>single</td>
      <td>FragPipe_Duncan_PXD004720_root-secretion_pepti...</td>
      <td>100639</td>
      <td>100085</td>
      <td>554</td>
      <td>100085</td>
      <td>99127</td>
      <td>958</td>
      <td>0</td>
      <td>0</td>
      <td>958</td>
      <td>0</td>
      <td>99.4495</td>
      <td>99.0428</td>
    </tr>
    <tr>
      <th>24</th>
      <td>PXD004720</td>
      <td>bread wheat</td>
      <td>root-tip</td>
      <td>single</td>
      <td>FragPipe_Duncan_PXD004720_root-tip_peptide_gen...</td>
      <td>191943</td>
      <td>191262</td>
      <td>681</td>
      <td>191262</td>
      <td>189975</td>
      <td>1287</td>
      <td>0</td>
      <td>0</td>
      <td>1287</td>
      <td>0</td>
      <td>99.6452</td>
      <td>99.3271</td>
    </tr>
    <tr>
      <th>25</th>
      <td>PXD004720</td>
      <td>bread wheat</td>
      <td>root-vasculature</td>
      <td>single</td>
      <td>FragPipe_Duncan_PXD004720_root-vasculature_pep...</td>
      <td>111831</td>
      <td>111315</td>
      <td>516</td>
      <td>111315</td>
      <td>110307</td>
      <td>1008</td>
      <td>0</td>
      <td>0</td>
      <td>1008</td>
      <td>0</td>
      <td>99.5386</td>
      <td>99.0945</td>
    </tr>
    <tr>
      <th>26</th>
      <td>PXD004720</td>
      <td>bread wheat</td>
      <td>spike-immature</td>
      <td>single</td>
      <td>FragPipe_Duncan_PXD004720_spike-immature_pepti...</td>
      <td>188826</td>
      <td>188025</td>
      <td>801</td>
      <td>188025</td>
      <td>186379</td>
      <td>1646</td>
      <td>0</td>
      <td>0</td>
      <td>1646</td>
      <td>0</td>
      <td>99.5758</td>
      <td>99.1246</td>
    </tr>
    <tr>
      <th>27</th>
      <td>PXD004720</td>
      <td>bread wheat</td>
      <td>stem</td>
      <td>single</td>
      <td>FragPipe_Duncan_PXD004720_stem_peptide_genome_...</td>
      <td>60734</td>
      <td>60181</td>
      <td>553</td>
      <td>60181</td>
      <td>59623</td>
      <td>558</td>
      <td>0</td>
      <td>0</td>
      <td>558</td>
      <td>0</td>
      <td>99.0895</td>
      <td>99.0728</td>
    </tr>
    <tr>
      <th>28</th>
      <td>PXD050500</td>
      <td>bread wheat</td>
      <td>coleoptile</td>
      <td>single</td>
      <td>FragPipe_Liu_PXD050500_coleoptile_peptide_geno...</td>
      <td>1850136</td>
      <td>1829196</td>
      <td>20940</td>
      <td>1829196</td>
      <td>1812027</td>
      <td>17169</td>
      <td>0</td>
      <td>0</td>
      <td>17169</td>
      <td>0</td>
      <td>98.8682</td>
      <td>99.0614</td>
    </tr>
    <tr>
      <th>29</th>
      <td>PXD050500</td>
      <td>bread wheat</td>
      <td>node</td>
      <td>single</td>
      <td>FragPipe_Liu_PXD050500_node_peptide_genome_pro...</td>
      <td>1886922</td>
      <td>1864415</td>
      <td>22507</td>
      <td>1864415</td>
      <td>1846620</td>
      <td>17795</td>
      <td>0</td>
      <td>0</td>
      <td>17795</td>
      <td>0</td>
      <td>98.8072</td>
      <td>99.0455</td>
    </tr>
    <tr>
      <th>30</th>
      <td>PXD050500</td>
      <td>bread wheat</td>
      <td>radicle</td>
      <td>single</td>
      <td>FragPipe_Liu_PXD050500_radicle_peptide_genome_...</td>
      <td>1062508</td>
      <td>1050178</td>
      <td>12330</td>
      <td>1050178</td>
      <td>1040357</td>
      <td>9821</td>
      <td>0</td>
      <td>0</td>
      <td>9821</td>
      <td>0</td>
      <td>98.8395</td>
      <td>99.0648</td>
    </tr>
    <tr>
      <th>31</th>
      <td>MSV000090572</td>
      <td>bread wheat</td>
      <td>stored_grain</td>
      <td>single</td>
      <td>FragPipe_Vincent_MSV000090572_stored-grain_pep...</td>
      <td>30314</td>
      <td>29892</td>
      <td>422</td>
      <td>29892</td>
      <td>29430</td>
      <td>462</td>
      <td>0</td>
      <td>0</td>
      <td>462</td>
      <td>0</td>
      <td>98.6079</td>
      <td>98.4544</td>
    </tr>
  </tbody>
</table>
</div>


<div>
<style scoped>
    .dataframe tbody tr th:only-of-type {
        vertical-align: middle;
    }

    .dataframe tbody tr th {
        vertical-align: top;
    }

    .dataframe thead th {
        text-align: right;
    }
</style>
<table border="1" class="dataframe">
  <thead>
    <tr style="text-align: right;">
      <th></th>
      <th>Source</th>
      <th>Species</th>
      <th>Tissue</th>
      <th>Batch</th>
      <th>Peptide</th>
      <th>ProteinID</th>
      <th>Probability</th>
      <th>GeneModel</th>
      <th>TranscriptID</th>
      <th>Chromosome</th>
      <th>...</th>
      <th>IL_normalised_match</th>
      <th>Validation_status</th>
      <th>Peptide_length_AA</th>
      <th>Projection_file</th>
      <th>Check_BED_geometry</th>
      <th>Check_block_nt_length</th>
      <th>Check_chromosome_and_strand</th>
      <th>Check_protein_coordinates</th>
      <th>All_sanity_checks_passed</th>
      <th>Sanity_check_status</th>
    </tr>
  </thead>
  <tbody>
    <tr>
      <th>0</th>
      <td>PXD004720</td>
      <td>bread wheat</td>
      <td>anther</td>
      <td>single</td>
      <td>ADLHHLAMFLAGRQPDAPQEALQVLDIVLR</td>
      <td>TraesCSU03G0061400.2</td>
      <td>1.0000</td>
      <td>TraesCSU03G0061400</td>
      <td>TraesCSU03G0061400.2</td>
      <td>ChrUnknown</td>
      <td>...</td>
      <td>True</td>
      <td>validated</td>
      <td>30</td>
      <td>FragPipe_Duncan_PXD004720_anther_peptide_genom...</td>
      <td>True</td>
      <td>True</td>
      <td>False</td>
      <td>True</td>
      <td>False</td>
      <td>failed</td>
    </tr>
    <tr>
      <th>1</th>
      <td>PXD004720</td>
      <td>bread wheat</td>
      <td>anther</td>
      <td>single</td>
      <td>AKQMAESFNMSTFANREDLLGQSK</td>
      <td>TraesCSU03G0111000.1</td>
      <td>0.4495</td>
      <td>TraesCSU03G0111000</td>
      <td>TraesCSU03G0111000.1</td>
      <td>ChrUnknown</td>
      <td>...</td>
      <td>True</td>
      <td>validated</td>
      <td>24</td>
      <td>FragPipe_Duncan_PXD004720_anther_peptide_genom...</td>
      <td>True</td>
      <td>True</td>
      <td>False</td>
      <td>True</td>
      <td>False</td>
      <td>failed</td>
    </tr>
    <tr>
      <th>2</th>
      <td>PXD004720</td>
      <td>bread wheat</td>
      <td>anther</td>
      <td>single</td>
      <td>AHEELVK</td>
      <td>TraesCSU03G0430400LC.1</td>
      <td>0.9305</td>
      <td>TraesCSU03G0430400LC</td>
      <td>TraesCSU03G0430400LC.1</td>
      <td>ChrUnknown</td>
      <td>...</td>
      <td>True</td>
      <td>validated</td>
      <td>7</td>
      <td>FragPipe_Duncan_PXD004720_anther_peptide_genom...</td>
      <td>True</td>
      <td>True</td>
      <td>False</td>
      <td>True</td>
      <td>False</td>
      <td>failed</td>
    </tr>
    <tr>
      <th>3</th>
      <td>PXD004720</td>
      <td>bread wheat</td>
      <td>anther</td>
      <td>single</td>
      <td>AAAMGVEEAVR</td>
      <td>TraesCSU03G0402100.1</td>
      <td>0.3517</td>
      <td>TraesCSU03G0402100</td>
      <td>TraesCSU03G0402100.1</td>
      <td>ChrUnknown</td>
      <td>...</td>
      <td>True</td>
      <td>validated</td>
      <td>11</td>
      <td>FragPipe_Duncan_PXD004720_anther_peptide_genom...</td>
      <td>True</td>
      <td>True</td>
      <td>False</td>
      <td>True</td>
      <td>False</td>
      <td>failed</td>
    </tr>
    <tr>
      <th>4</th>
      <td>PXD004720</td>
      <td>bread wheat</td>
      <td>anther</td>
      <td>single</td>
      <td>APGIIER</td>
      <td>TraesCSU03G0138700LC.1</td>
      <td>0.7045</td>
      <td>TraesCSU03G0138700LC</td>
      <td>TraesCSU03G0138700LC.1</td>
      <td>ChrUnknown</td>
      <td>...</td>
      <td>True</td>
      <td>validated</td>
      <td>7</td>
      <td>FragPipe_Duncan_PXD004720_anther_peptide_genom...</td>
      <td>True</td>
      <td>True</td>
      <td>False</td>
      <td>True</td>
      <td>False</td>
      <td>failed</td>
    </tr>
    <tr>
      <th>5</th>
      <td>PXD004720</td>
      <td>bread wheat</td>
      <td>anther</td>
      <td>single</td>
      <td>DSDMQSADGDGR</td>
      <td>TraesCSU03G0144900LC.1</td>
      <td>0.5074</td>
      <td>TraesCSU03G0144900LC</td>
      <td>TraesCSU03G0144900LC.1</td>
      <td>ChrUnknown</td>
      <td>...</td>
      <td>True</td>
      <td>validated</td>
      <td>12</td>
      <td>FragPipe_Duncan_PXD004720_anther_peptide_genom...</td>
      <td>True</td>
      <td>True</td>
      <td>False</td>
      <td>True</td>
      <td>False</td>
      <td>failed</td>
    </tr>
    <tr>
      <th>6</th>
      <td>PXD004720</td>
      <td>bread wheat</td>
      <td>anther</td>
      <td>single</td>
      <td>AVGVSTEK</td>
      <td>TraesCSU03G0112900LC.1</td>
      <td>0.1800</td>
      <td>TraesCSU03G0112900LC</td>
      <td>TraesCSU03G0112900LC.1</td>
      <td>ChrUnknown</td>
      <td>...</td>
      <td>True</td>
      <td>validated</td>
      <td>8</td>
      <td>FragPipe_Duncan_PXD004720_anther_peptide_genom...</td>
      <td>True</td>
      <td>True</td>
      <td>False</td>
      <td>True</td>
      <td>False</td>
      <td>failed</td>
    </tr>
    <tr>
      <th>7</th>
      <td>PXD004720</td>
      <td>bread wheat</td>
      <td>anther</td>
      <td>single</td>
      <td>APGIIER</td>
      <td>TraesCSU03G0351700LC.1</td>
      <td>0.7045</td>
      <td>TraesCSU03G0351700LC</td>
      <td>TraesCSU03G0351700LC.1</td>
      <td>ChrUnknown</td>
      <td>...</td>
      <td>True</td>
      <td>validated</td>
      <td>7</td>
      <td>FragPipe_Duncan_PXD004720_anther_peptide_genom...</td>
      <td>True</td>
      <td>True</td>
      <td>False</td>
      <td>True</td>
      <td>False</td>
      <td>failed</td>
    </tr>
    <tr>
      <th>8</th>
      <td>PXD004720</td>
      <td>bread wheat</td>
      <td>anther</td>
      <td>single</td>
      <td>AEHLASIFGTEKDR</td>
      <td>TraesCSU03G0553300LC.1</td>
      <td>1.0000</td>
      <td>TraesCSU03G0553300LC</td>
      <td>TraesCSU03G0553300LC.1</td>
      <td>ChrUnknown</td>
      <td>...</td>
      <td>True</td>
      <td>validated</td>
      <td>14</td>
      <td>FragPipe_Duncan_PXD004720_anther_peptide_genom...</td>
      <td>True</td>
      <td>True</td>
      <td>False</td>
      <td>True</td>
      <td>False</td>
      <td>failed</td>
    </tr>
    <tr>
      <th>9</th>
      <td>PXD004720</td>
      <td>bread wheat</td>
      <td>anther</td>
      <td>single</td>
      <td>AVANETGAFFFLINGPEIMSK</td>
      <td>TraesCSU03G0001700.1</td>
      <td>1.0000</td>
      <td>TraesCSU03G0001700</td>
      <td>TraesCSU03G0001700.1</td>
      <td>ChrUnknown</td>
      <td>...</td>
      <td>True</td>
      <td>validated</td>
      <td>21</td>
      <td>FragPipe_Duncan_PXD004720_anther_peptide_genom...</td>
      <td>True</td>
      <td>True</td>
      <td>False</td>
      <td>True</td>
      <td>False</td>
      <td>failed</td>
    </tr>
    <tr>
      <th>10</th>
      <td>PXD004720</td>
      <td>bread wheat</td>
      <td>anther</td>
      <td>single</td>
      <td>DNGMHALIIYDDLSK</td>
      <td>TraesCSU03G0519600LC.1</td>
      <td>1.0000</td>
      <td>TraesCSU03G0519600LC</td>
      <td>TraesCSU03G0519600LC.1</td>
      <td>ChrUnknown</td>
      <td>...</td>
      <td>True</td>
      <td>validated</td>
      <td>15</td>
      <td>FragPipe_Duncan_PXD004720_anther_peptide_genom...</td>
      <td>True</td>
      <td>True</td>
      <td>False</td>
      <td>True</td>
      <td>False</td>
      <td>failed</td>
    </tr>
    <tr>
      <th>11</th>
      <td>PXD004720</td>
      <td>bread wheat</td>
      <td>anther</td>
      <td>single</td>
      <td>ASVAPPTVGKER</td>
      <td>TraesCSU03G0433300LC.1</td>
      <td>0.6832</td>
      <td>TraesCSU03G0433300LC</td>
      <td>TraesCSU03G0433300LC.1</td>
      <td>ChrUnknown</td>
      <td>...</td>
      <td>True</td>
      <td>validated</td>
      <td>12</td>
      <td>FragPipe_Duncan_PXD004720_anther_peptide_genom...</td>
      <td>True</td>
      <td>True</td>
      <td>False</td>
      <td>True</td>
      <td>False</td>
      <td>failed</td>
    </tr>
    <tr>
      <th>12</th>
      <td>PXD004720</td>
      <td>bread wheat</td>
      <td>anther</td>
      <td>single</td>
      <td>AFTELLK</td>
      <td>TraesCSU03G0362800LC.1</td>
      <td>0.9993</td>
      <td>TraesCSU03G0362800LC</td>
      <td>TraesCSU03G0362800LC.1</td>
      <td>ChrUnknown</td>
      <td>...</td>
      <td>True</td>
      <td>validated</td>
      <td>7</td>
      <td>FragPipe_Duncan_PXD004720_anther_peptide_genom...</td>
      <td>True</td>
      <td>True</td>
      <td>False</td>
      <td>True</td>
      <td>False</td>
      <td>failed</td>
    </tr>
    <tr>
      <th>13</th>
      <td>PXD004720</td>
      <td>bread wheat</td>
      <td>anther</td>
      <td>single</td>
      <td>AVTSPSAR</td>
      <td>TraesCSU03G0279300.1</td>
      <td>0.0573</td>
      <td>TraesCSU03G0279300</td>
      <td>TraesCSU03G0279300.1</td>
      <td>ChrUnknown</td>
      <td>...</td>
      <td>True</td>
      <td>validated</td>
      <td>8</td>
      <td>FragPipe_Duncan_PXD004720_anther_peptide_genom...</td>
      <td>True</td>
      <td>True</td>
      <td>False</td>
      <td>True</td>
      <td>False</td>
      <td>failed</td>
    </tr>
    <tr>
      <th>14</th>
      <td>PXD004720</td>
      <td>bread wheat</td>
      <td>anther</td>
      <td>single</td>
      <td>ADGFAGVFPEHKYEIVK</td>
      <td>TraesCSU03G0076400.4</td>
      <td>0.5887</td>
      <td>TraesCSU03G0076400</td>
      <td>TraesCSU03G0076400.4</td>
      <td>ChrUnknown</td>
      <td>...</td>
      <td>True</td>
      <td>validated</td>
      <td>17</td>
      <td>FragPipe_Duncan_PXD004720_anther_peptide_genom...</td>
      <td>True</td>
      <td>True</td>
      <td>False</td>
      <td>True</td>
      <td>False</td>
      <td>failed</td>
    </tr>
    <tr>
      <th>15</th>
      <td>PXD004720</td>
      <td>bread wheat</td>
      <td>anther</td>
      <td>single</td>
      <td>AHVIVMGATNRPNSIDPALR</td>
      <td>TraesCSU03G0001700.1</td>
      <td>1.0000</td>
      <td>TraesCSU03G0001700</td>
      <td>TraesCSU03G0001700.1</td>
      <td>ChrUnknown</td>
      <td>...</td>
      <td>True</td>
      <td>validated</td>
      <td>20</td>
      <td>FragPipe_Duncan_PXD004720_anther_peptide_genom...</td>
      <td>True</td>
      <td>True</td>
      <td>False</td>
      <td>True</td>
      <td>False</td>
      <td>failed</td>
    </tr>
    <tr>
      <th>16</th>
      <td>PXD004720</td>
      <td>bread wheat</td>
      <td>anther</td>
      <td>single</td>
      <td>AAGIPSK</td>
      <td>TraesCSU03G0054300.1</td>
      <td>0.9320</td>
      <td>TraesCSU03G0054300</td>
      <td>TraesCSU03G0054300.1</td>
      <td>ChrUnknown</td>
      <td>...</td>
      <td>True</td>
      <td>validated</td>
      <td>7</td>
      <td>FragPipe_Duncan_PXD004720_anther_peptide_genom...</td>
      <td>True</td>
      <td>True</td>
      <td>False</td>
      <td>True</td>
      <td>False</td>
      <td>failed</td>
    </tr>
    <tr>
      <th>17</th>
      <td>PXD004720</td>
      <td>bread wheat</td>
      <td>anther</td>
      <td>single</td>
      <td>AGVYHPWGPGNGRR</td>
      <td>TraesCSU03G0292000LC.1</td>
      <td>0.2867</td>
      <td>TraesCSU03G0292000LC</td>
      <td>TraesCSU03G0292000LC.1</td>
      <td>ChrUnknown</td>
      <td>...</td>
      <td>True</td>
      <td>validated</td>
      <td>14</td>
      <td>FragPipe_Duncan_PXD004720_anther_peptide_genom...</td>
      <td>True</td>
      <td>True</td>
      <td>False</td>
      <td>True</td>
      <td>False</td>
      <td>failed</td>
    </tr>
    <tr>
      <th>18</th>
      <td>PXD004720</td>
      <td>bread wheat</td>
      <td>anther</td>
      <td>single</td>
      <td>AAQLSGR</td>
      <td>TraesCSU03G0023500LC.1</td>
      <td>0.6112</td>
      <td>TraesCSU03G0023500LC</td>
      <td>TraesCSU03G0023500LC.1</td>
      <td>ChrUnknown</td>
      <td>...</td>
      <td>True</td>
      <td>validated</td>
      <td>7</td>
      <td>FragPipe_Duncan_PXD004720_anther_peptide_genom...</td>
      <td>True</td>
      <td>True</td>
      <td>False</td>
      <td>True</td>
      <td>False</td>
      <td>failed</td>
    </tr>
    <tr>
      <th>19</th>
      <td>PXD004720</td>
      <td>bread wheat</td>
      <td>anther</td>
      <td>single</td>
      <td>AAVFQMEHNKAP</td>
      <td>TraesCSU03G0002700LC.1</td>
      <td>0.0546</td>
      <td>TraesCSU03G0002700LC</td>
      <td>TraesCSU03G0002700LC.1</td>
      <td>ChrUnknown</td>
      <td>...</td>
      <td>True</td>
      <td>validated</td>
      <td>12</td>
      <td>FragPipe_Duncan_PXD004720_anther_peptide_genom...</td>
      <td>True</td>
      <td>True</td>
      <td>False</td>
      <td>True</td>
      <td>False</td>
      <td>failed</td>
    </tr>
  </tbody>
</table>
<p>20 rows × 38 columns</p>
</div>


# Step 11B — Diagnostic Audit: Fate of ChrUnknown Peptide Projections

This diagnostic step tracks peptide projections assigned to `ChrUnknown` across the validation workflow.

The purpose is to determine whether `ChrUnknown` peptide projections were:

1. present after initial annotation-guided peptide genome projection,
2. retained after translation validation,
3. excluded during sanity checking,
4. absent from the final non-redundant validated peptide set.

This audit was added because final validated BED and EDA outputs no longer contained peptide projections assigned to `ChrUnknown`.

---

## Workflow stages audited

The diagnostic compares `ChrUnknown` counts across:

| Stage | File type | Interpretation |
|---|---|---|
| Step 9 | Tissue-level peptide genome projection files | Initial annotation-guided projected peptide coordinates |
| Step 10 | Full translation-validation table | Rows tested by translation reconstruction |
| Step 11 | Full sanity-check table | Translation-validated rows after BED/chromosome/protein-coordinate checks |
| Step 13 | All-tissue non-redundant validated table | Final non-redundant validated peptide projection set |

---

## Expected outcome

If `ChrUnknown` rows are present in Step 9 and Step 10 but fail Step 11, this indicates that they were removed by the sanity-check layer rather than by peptide-to-genome projection or translation validation.

In particular, rows assigned to `ChrUnknown` are expected to fail the chromosome/strand sanity check when `ChrUnknown` is not included in the accepted chromosome list used for final chromosome-scale BED export.

---

## Output files

### Summary table

```text
wheat_ChrUnknown_validation_fate_summary_step11B.csv


```python
# ============================================================
# Step 11B — Diagnostic audit: fate of ChrUnknown peptide projections
# Memory-light version for large CSV files
# ============================================================

import pandas as pd
from pathlib import Path
from collections import defaultdict

# -----------------------------
# 1. Paths
# -----------------------------
fragpipe_dir = Path("FragPipe_results")
tables_dir = Path("python_outputs/tables")
tables_dir.mkdir(parents=True, exist_ok=True)

manifest_file = fragpipe_dir / "wheat_tissues_FragPipe-result-manifest_2026-05-11.csv"

# Existing large workflow files
step10_validation_file = tables_dir / "wheat_projection_validation_stratified100percent_step10.csv"
step11_sanity_file = tables_dir / "wheat_projection_translation_validated_sanity_checks_full_step11.csv"
step13_combined_file = tables_dir / "wheat_all_tissues_nonredundant_validated_peptides_step13.csv"

summary_out = tables_dir / "wheat_ChrUnknown_validation_fate_summary_step11B.csv"
failed_examples_out = tables_dir / "wheat_ChrUnknown_failed_sanity_examples_step11B.csv"

chunksize = 25_000
max_failed_examples = 100

manifest = pd.read_csv(manifest_file, encoding="utf-8-sig")

# -----------------------------
# 2. Helper functions
# -----------------------------
def normalise_chr(value):
    """
    Normalise chromosome labels for ChrUnknown audit.
    """
    if pd.isna(value):
        return "NA"

    value = str(value).strip()

    if value.lower() in ["chrunknown", "unknown"]:
        return "ChrUnknown"

    if value.lower() in ["nan", "none", "", "<na>"]:
        return "NA"

    return value


def build_manifest_lookup(manifest):
    """
    Build lookup from Step 9 projection filename to source/tissue metadata.
    """
    lookup = {}

    for _, row in manifest.iterrows():

        projection_filename = row["FragPipe-Output-Peptide"].replace(
            "_peptide.tsv",
            "_peptide_genome_projection.csv"
        )

        lookup[projection_filename] = {
            "Source": row["Source"],
            "Species": row["Species"],
            "Tissue": row["Tissue-Raw-Code"],
            "Batch": row["Batch"]
        }

    return lookup


manifest_lookup = build_manifest_lookup(manifest)


def get_meta_from_projection_file(projection_file):
    """
    Recover Source/Species/Tissue/Batch from projection filename.
    """
    projection_file = Path(str(projection_file)).name

    return manifest_lookup.get(
        projection_file,
        {
            "Source": "UNKNOWN",
            "Species": "bread wheat",
            "Tissue": "UNKNOWN",
            "Batch": "UNKNOWN"
        }
    )


def get_gene_col(columns):
    """
    Return the first available gene-model column.
    """
    for col in ["GeneModel", "GeneID", "Gene_label", "Gene_model"]:
        if col in columns:
            return col
    return None


def init_record():
    """
    Initialise a compact audit record.
    """
    return {
        "Total_rows": 0,
        "ChrUnknown_rows": 0,
        "ChrUnknown_peptides": set(),
        "ChrUnknown_proteins": set(),
        "ChrUnknown_genes": set()
    }


def update_record(records, key, chunk, chr_mask):
    """
    Update compact counts for one audit group.
    Only unique values from ChrUnknown rows are retained.
    """
    rec = records[key]

    rec["Total_rows"] += len(chunk)
    rec["ChrUnknown_rows"] += int(chr_mask.sum())

    chr_chunk = chunk.loc[chr_mask]

    if chr_chunk.empty:
        return

    if "Peptide" in chr_chunk.columns:
        rec["ChrUnknown_peptides"].update(
            chr_chunk["Peptide"].dropna().astype(str).unique()
        )

    if "ProteinID" in chr_chunk.columns:
        rec["ChrUnknown_proteins"].update(
            chr_chunk["ProteinID"].dropna().astype(str).unique()
        )

    gene_col = get_gene_col(chr_chunk.columns)

    if gene_col is not None:
        rec["ChrUnknown_genes"].update(
            chr_chunk[gene_col].dropna().astype(str).unique()
        )


def records_to_dataframe(records):
    """
    Convert compact audit dictionary to a DataFrame.
    """
    rows = []

    for key, rec in records.items():

        stage, source, species, tissue, batch, projection_file, status_detail = key

        total_rows = rec["Total_rows"]
        chr_rows = rec["ChrUnknown_rows"]

        rows.append({
            "Stage": stage,
            "Source": source,
            "Species": species,
            "Tissue": tissue,
            "Batch": batch,
            "Projection_file": projection_file,
            "Status_detail": status_detail,
            "Total_rows": total_rows,
            "ChrUnknown_rows": chr_rows,
            "ChrUnknown_percent_rows": round((chr_rows / total_rows) * 100, 4) if total_rows > 0 else 0,
            "ChrUnknown_unique_peptides": len(rec["ChrUnknown_peptides"]),
            "ChrUnknown_unique_proteins": len(rec["ChrUnknown_proteins"]),
            "ChrUnknown_unique_gene_models": len(rec["ChrUnknown_genes"])
        })

    return pd.DataFrame(rows)


# -----------------------------
# 3. Stage A — Step 9 initial projected rows
# -----------------------------
print("Auditing Step 9 projected rows...")

step9_records = defaultdict(init_record)

for _, row in manifest.iterrows():

    projection_filename = row["FragPipe-Output-Peptide"].replace(
        "_peptide.tsv",
        "_peptide_genome_projection.csv"
    )

    projection_path = tables_dir / projection_filename

    if not projection_path.exists():
        print(f"Skipped missing Step 9 file: {projection_path}")
        continue

    header = pd.read_csv(projection_path, nrows=0)
    header_cols = list(header.columns)

    required = ["Chromosome", "Projection_status"]
    missing = [c for c in required if c not in header_cols]

    if missing:
        raise KeyError(
            f"Missing required column(s) in {projection_filename}: {missing}"
        )

    usecols = [
        col for col in [
            "Chromosome",
            "Projection_status",
            "Peptide",
            "ProteinID",
            "GeneModel",
            "GeneID",
            "Gene_label"
        ]
        if col in header_cols
    ]

    key = (
        "Step 9 projected rows",
        row["Source"],
        row["Species"],
        row["Tissue-Raw-Code"],
        row["Batch"],
        projection_filename,
        "Projection_status == projected"
    )

    for chunk in pd.read_csv(
        projection_path,
        usecols=usecols,
        chunksize=chunksize,
        low_memory=False
    ):

        chunk = chunk[
            chunk["Projection_status"].astype(str) == "projected"
        ].copy()

        if chunk.empty:
            continue

        chr_norm = chunk["Chromosome"].apply(normalise_chr)
        chr_mask = chr_norm == "ChrUnknown"

        update_record(step9_records, key, chunk, chr_mask)

step9_summary = records_to_dataframe(step9_records)

# -----------------------------
# 4. Stage B — Step 10 translation validation
# -----------------------------
print("Auditing Step 10 translation-validation rows...")

if not step10_validation_file.exists():
    raise FileNotFoundError(
        f"Step 10 file not found:\n{step10_validation_file}"
    )

step10_header = pd.read_csv(step10_validation_file, nrows=0)
step10_cols = list(step10_header.columns)

# Step 10 may or may not contain a source/projection filename column.
# If absent, we still audit ChrUnknown globally by Validation_status.
source_file_col = None

for candidate in ["_source_file", "Projection_file", "Genome_projection_file"]:
    if candidate in step10_cols:
        source_file_col = candidate
        break

required = ["Chromosome", "Validation_status"]
missing = [c for c in required if c not in step10_cols]

if missing:
    raise KeyError(
        f"Missing required Step 10 column(s): {missing}"
    )

step10_usecols = [
    col for col in [
        source_file_col,
        "Source",
        "Species",
        "Tissue",
        "Batch",
        "Chromosome",
        "Validation_status",
        "Peptide",
        "ProteinID",
        "GeneModel",
        "GeneID",
        "Gene_label"
    ]
    if col is not None and col in step10_cols
]

print(f"Step 10 columns used: {step10_usecols}")

if source_file_col is None:
    print(
        "No Step 10 source/projection filename column found. "
        "Step 10 ChrUnknown audit will be summarised globally by Validation_status."
    )
else:
    print(f"Detected Step 10 source-file column: {source_file_col}")

step10_records = defaultdict(init_record)

for chunk_i, chunk in enumerate(
    pd.read_csv(
        step10_validation_file,
        usecols=step10_usecols,
        chunksize=chunksize,
        low_memory=False
    ),
    start=1
):

    chunk["Chromosome_norm"] = chunk["Chromosome"].apply(normalise_chr)

    # Case 1: Step 10 has explicit source/tissue metadata
    if all(col in chunk.columns for col in ["Source", "Species", "Tissue", "Batch"]):

        for (source, species, tissue, batch, status), group in chunk.groupby(
            ["Source", "Species", "Tissue", "Batch", "Validation_status"],
            dropna=False
        ):

            key = (
                "Step 10 translation validation",
                source,
                species,
                tissue,
                batch,
                "Step10_global_table",
                f"Validation_status == {status}"
            )

            chr_mask = group["Chromosome_norm"] == "ChrUnknown"

            update_record(step10_records, key, group, chr_mask)

    # Case 2: Step 10 has projection filename metadata but not Source/Tissue
    elif source_file_col is not None:

        for (projection_file, status), group in chunk.groupby(
            [source_file_col, "Validation_status"],
            dropna=False
        ):

            projection_file_name = Path(str(projection_file)).name
            meta = get_meta_from_projection_file(projection_file_name)

            key = (
                "Step 10 translation validation",
                meta["Source"],
                meta["Species"],
                meta["Tissue"],
                meta["Batch"],
                projection_file_name,
                f"Validation_status == {status}"
            )

            chr_mask = group["Chromosome_norm"] == "ChrUnknown"

            update_record(step10_records, key, group, chr_mask)

    # Case 3: Step 10 has no source/projection metadata
    else:

        for status, group in chunk.groupby("Validation_status", dropna=False):

            key = (
                "Step 10 translation validation",
                "ALL",
                "bread wheat",
                "ALL",
                "ALL",
                step10_validation_file.name,
                f"Validation_status == {status}"
            )

            chr_mask = group["Chromosome_norm"] == "ChrUnknown"

            update_record(step10_records, key, group, chr_mask)

    if chunk_i % 20 == 0:
        print(f"  Step 10 chunks processed: {chunk_i}")

step10_summary = records_to_dataframe(step10_records)

# -----------------------------
# 5. Stage C — Step 11 sanity checks
# -----------------------------
print("Auditing Step 11 sanity-check rows...")

if not step11_sanity_file.exists():
    raise FileNotFoundError(
        f"Step 11 file not found:\n{step11_sanity_file}"
    )

step11_header = pd.read_csv(step11_sanity_file, nrows=0)
step11_cols = list(step11_header.columns)

required = [
    "Source",
    "Species",
    "Tissue",
    "Batch",
    "Projection_file",
    "Chromosome",
    "Sanity_check_status"
]

missing = [c for c in required if c not in step11_cols]

if missing:
    raise KeyError(
        f"Missing required Step 11 column(s): {missing}"
    )

step11_usecols = [
    col for col in [
        "Source",
        "Species",
        "Tissue",
        "Batch",
        "Projection_file",
        "Chromosome",
        "Sanity_check_status",
        "Validation_status",
        "Peptide",
        "ProteinID",
        "GeneModel",
        "GeneID",
        "Gene_label",
        "Check_BED_geometry",
        "Check_block_nt_length",
        "Check_chromosome_and_strand",
        "Check_protein_coordinates",
        "All_sanity_checks_passed"
    ]
    if col in step11_cols
]

step11_records = defaultdict(init_record)
failed_examples = []

for chunk_i, chunk in enumerate(
    pd.read_csv(
        step11_sanity_file,
        usecols=step11_usecols,
        chunksize=chunksize,
        low_memory=False
    ),
    start=1
):

    chunk["Chromosome_norm"] = chunk["Chromosome"].apply(normalise_chr)

    for (source, species, tissue, batch, projection_file, status), group in chunk.groupby(
        ["Source", "Species", "Tissue", "Batch", "Projection_file", "Sanity_check_status"],
        dropna=False
    ):

        projection_file_name = Path(str(projection_file)).name

        key = (
            "Step 11 sanity checks",
            source,
            species,
            tissue,
            batch,
            projection_file_name,
            f"Sanity_check_status == {status}"
        )

        chr_mask = group["Chromosome_norm"] == "ChrUnknown"

        update_record(step11_records, key, group, chr_mask)

    # Keep only a small number of failed ChrUnknown examples
    if len(failed_examples) < max_failed_examples:

        chr_failed = chunk[
            (chunk["Chromosome_norm"] == "ChrUnknown") &
            (chunk["Sanity_check_status"].astype(str) == "failed")
        ].copy()

        if not chr_failed.empty:

            slots_remaining = max_failed_examples - len(failed_examples)
            failed_examples.extend(
                chr_failed.head(slots_remaining).to_dict("records")
            )

    if chunk_i % 20 == 0:
        print(f"  Step 11 chunks processed: {chunk_i}")

step11_summary = records_to_dataframe(step11_records)

failed_examples_df = pd.DataFrame(failed_examples)
failed_examples_df.to_csv(failed_examples_out, index=False)

# -----------------------------
# 6. Stage D — Step 13 final non-redundant validated table
# -----------------------------
print("Auditing Step 13 final non-redundant validated table...")

step13_records = defaultdict(init_record)

if step13_combined_file.exists():

    step13_header = pd.read_csv(step13_combined_file, nrows=0)
    step13_cols = list(step13_header.columns)

    if "Chromosome" not in step13_cols:
        raise KeyError("Step 13 combined table lacks 'Chromosome' column.")

    step13_usecols = [
        col for col in [
            "Chromosome",
            "Peptide",
            "ProteinID",
            "GeneModel",
            "GeneID",
            "Gene_label"
        ]
        if col in step13_cols
    ]

    key = (
        "Step 13 final non-redundant validated table",
        "ALL",
        "bread wheat",
        "ALL",
        "ALL",
        step13_combined_file.name,
        "Final non-redundant validated rows"
    )

    for chunk_i, chunk in enumerate(
        pd.read_csv(
            step13_combined_file,
            usecols=step13_usecols,
            chunksize=chunksize,
            low_memory=False
        ),
        start=1
    ):

        chr_norm = chunk["Chromosome"].apply(normalise_chr)
        chr_mask = chr_norm == "ChrUnknown"

        update_record(step13_records, key, chunk, chr_mask)

        if chunk_i % 20 == 0:
            print(f"  Step 13 chunks processed: {chunk_i}")

else:
    print(f"Step 13 file not found, skipped: {step13_combined_file}")

step13_summary = records_to_dataframe(step13_records)

# -----------------------------
# 7. Combine, filter, export
# -----------------------------
audit_summary = pd.concat(
    [
        step9_summary,
        step10_summary,
        step11_summary,
        step13_summary
    ],
    ignore_index=True
)

# Keep only informative rows:
# - any row with ChrUnknown
# - plus the final Step 13 row, even if ChrUnknown count is zero
audit_summary = audit_summary[
    (audit_summary["ChrUnknown_rows"] > 0) |
    (audit_summary["Stage"].str.contains("Step 13", regex=False))
].copy()

audit_summary = audit_summary.sort_values(
    [
        "Stage",
        "Source",
        "Tissue",
        "Projection_file",
        "Status_detail"
    ],
    na_position="last"
).reset_index(drop=True)

audit_summary.to_csv(summary_out, index=False)

print("\n===== ChrUnknown VALIDATION FATE SUMMARY =====")
print(f"Summary saved: {summary_out}")
print(f"Failed examples saved: {failed_examples_out}")
print(f"Summary rows: {len(audit_summary):,}")

display(audit_summary)

if not failed_examples_df.empty:
    print("\nExample ChrUnknown rows failing Step 11 sanity checks:")
    display(failed_examples_df.head(20))
else:
    print("\nNo failed ChrUnknown examples were found in Step 11.")
```

    Auditing Step 9 projected rows...
    Auditing Step 10 translation-validation rows...
    Step 10 columns used: ['Source', 'Species', 'Tissue', 'Batch', 'Chromosome', 'Validation_status', 'Peptide', 'ProteinID', 'GeneModel']
    No Step 10 source/projection filename column found. Step 10 ChrUnknown audit will be summarised globally by Validation_status.
      Step 10 chunks processed: 20
      Step 10 chunks processed: 40
      Step 10 chunks processed: 60
      Step 10 chunks processed: 80
      Step 10 chunks processed: 100
      Step 10 chunks processed: 120
      Step 10 chunks processed: 140
      Step 10 chunks processed: 160
      Step 10 chunks processed: 180
      Step 10 chunks processed: 200
      Step 10 chunks processed: 220
      Step 10 chunks processed: 240
      Step 10 chunks processed: 260
      Step 10 chunks processed: 280
      Step 10 chunks processed: 300
      Step 10 chunks processed: 320
    Auditing Step 11 sanity-check rows...
      Step 11 chunks processed: 20
      Step 11 chunks processed: 40
      Step 11 chunks processed: 60
      Step 11 chunks processed: 80
      Step 11 chunks processed: 100
      Step 11 chunks processed: 120
      Step 11 chunks processed: 140
      Step 11 chunks processed: 160
      Step 11 chunks processed: 180
      Step 11 chunks processed: 200
      Step 11 chunks processed: 220
      Step 11 chunks processed: 240
      Step 11 chunks processed: 260
      Step 11 chunks processed: 280
      Step 11 chunks processed: 300
      Step 11 chunks processed: 320
    Auditing Step 13 final non-redundant validated table...
      Step 13 chunks processed: 20
      Step 13 chunks processed: 40
      Step 13 chunks processed: 60
      Step 13 chunks processed: 80
      Step 13 chunks processed: 100
      Step 13 chunks processed: 120
    
    ===== ChrUnknown VALIDATION FATE SUMMARY =====
    Summary saved: python_outputs\tables\wheat_ChrUnknown_validation_fate_summary_step11B.csv
    Failed examples saved: python_outputs\tables\wheat_ChrUnknown_failed_sanity_examples_step11B.csv
    Summary rows: 128
    


<div>
<style scoped>
    .dataframe tbody tr th:only-of-type {
        vertical-align: middle;
    }

    .dataframe tbody tr th {
        vertical-align: top;
    }

    .dataframe thead th {
        text-align: right;
    }
</style>
<table border="1" class="dataframe">
  <thead>
    <tr style="text-align: right;">
      <th></th>
      <th>Stage</th>
      <th>Source</th>
      <th>Species</th>
      <th>Tissue</th>
      <th>Batch</th>
      <th>Projection_file</th>
      <th>Status_detail</th>
      <th>Total_rows</th>
      <th>ChrUnknown_rows</th>
      <th>ChrUnknown_percent_rows</th>
      <th>ChrUnknown_unique_peptides</th>
      <th>ChrUnknown_unique_proteins</th>
      <th>ChrUnknown_unique_gene_models</th>
    </tr>
  </thead>
  <tbody>
    <tr>
      <th>0</th>
      <td>Step 10 translation validation</td>
      <td>MSV000090572</td>
      <td>bread wheat</td>
      <td>stored_grain</td>
      <td>single</td>
      <td>Step10_global_table</td>
      <td>Validation_status == translation_mismatch</td>
      <td>422</td>
      <td>16</td>
      <td>3.7915</td>
      <td>13</td>
      <td>11</td>
      <td>11</td>
    </tr>
    <tr>
      <th>1</th>
      <td>Step 10 translation validation</td>
      <td>MSV000090572</td>
      <td>bread wheat</td>
      <td>stored_grain</td>
      <td>single</td>
      <td>Step10_global_table</td>
      <td>Validation_status == validated</td>
      <td>29892</td>
      <td>462</td>
      <td>1.5456</td>
      <td>209</td>
      <td>291</td>
      <td>273</td>
    </tr>
    <tr>
      <th>2</th>
      <td>Step 10 translation validation</td>
      <td>PXD004720</td>
      <td>bread wheat</td>
      <td>anther</td>
      <td>single</td>
      <td>Step10_global_table</td>
      <td>Validation_status == translation_mismatch</td>
      <td>898</td>
      <td>19</td>
      <td>2.1158</td>
      <td>18</td>
      <td>13</td>
      <td>13</td>
    </tr>
    <tr>
      <th>3</th>
      <td>Step 10 translation validation</td>
      <td>PXD004720</td>
      <td>bread wheat</td>
      <td>anther</td>
      <td>single</td>
      <td>Step10_global_table</td>
      <td>Validation_status == validated</td>
      <td>163467</td>
      <td>1512</td>
      <td>0.9250</td>
      <td>696</td>
      <td>467</td>
      <td>437</td>
    </tr>
    <tr>
      <th>4</th>
      <td>Step 10 translation validation</td>
      <td>PXD004720</td>
      <td>bread wheat</td>
      <td>boot</td>
      <td>single</td>
      <td>Step10_global_table</td>
      <td>Validation_status == validated</td>
      <td>13137</td>
      <td>113</td>
      <td>0.8602</td>
      <td>69</td>
      <td>98</td>
      <td>88</td>
    </tr>
    <tr>
      <th>...</th>
      <td>...</td>
      <td>...</td>
      <td>...</td>
      <td>...</td>
      <td>...</td>
      <td>...</td>
      <td>...</td>
      <td>...</td>
      <td>...</td>
      <td>...</td>
      <td>...</td>
      <td>...</td>
      <td>...</td>
    </tr>
    <tr>
      <th>123</th>
      <td>Step 9 projected rows</td>
      <td>PXD004720</td>
      <td>bread wheat</td>
      <td>spike-immature</td>
      <td>single</td>
      <td>FragPipe_Duncan_PXD004720_spike-immature_pepti...</td>
      <td>Projection_status == projected</td>
      <td>188826</td>
      <td>1666</td>
      <td>0.8823</td>
      <td>749</td>
      <td>511</td>
      <td>469</td>
    </tr>
    <tr>
      <th>124</th>
      <td>Step 9 projected rows</td>
      <td>PXD004720</td>
      <td>bread wheat</td>
      <td>stem</td>
      <td>single</td>
      <td>FragPipe_Duncan_PXD004720_stem_peptide_genome_...</td>
      <td>Projection_status == projected</td>
      <td>60734</td>
      <td>572</td>
      <td>0.9418</td>
      <td>304</td>
      <td>349</td>
      <td>329</td>
    </tr>
    <tr>
      <th>125</th>
      <td>Step 9 projected rows</td>
      <td>PXD050500</td>
      <td>bread wheat</td>
      <td>coleoptile</td>
      <td>single</td>
      <td>FragPipe_Liu_PXD050500_coleoptile_peptide_geno...</td>
      <td>Projection_status == projected</td>
      <td>1850136</td>
      <td>17731</td>
      <td>0.9584</td>
      <td>10538</td>
      <td>4324</td>
      <td>4180</td>
    </tr>
    <tr>
      <th>126</th>
      <td>Step 9 projected rows</td>
      <td>PXD050500</td>
      <td>bread wheat</td>
      <td>node</td>
      <td>single</td>
      <td>FragPipe_Liu_PXD050500_node_peptide_genome_pro...</td>
      <td>Projection_status == projected</td>
      <td>1886922</td>
      <td>18421</td>
      <td>0.9762</td>
      <td>10806</td>
      <td>4452</td>
      <td>4309</td>
    </tr>
    <tr>
      <th>127</th>
      <td>Step 9 projected rows</td>
      <td>PXD050500</td>
      <td>bread wheat</td>
      <td>radicle</td>
      <td>single</td>
      <td>FragPipe_Liu_PXD050500_radicle_peptide_genome_...</td>
      <td>Projection_status == projected</td>
      <td>1062508</td>
      <td>10116</td>
      <td>0.9521</td>
      <td>5891</td>
      <td>3509</td>
      <td>3379</td>
    </tr>
  </tbody>
</table>
<p>128 rows × 13 columns</p>
</div>


    
    Example ChrUnknown rows failing Step 11 sanity checks:
    


<div>
<style scoped>
    .dataframe tbody tr th:only-of-type {
        vertical-align: middle;
    }

    .dataframe tbody tr th {
        vertical-align: top;
    }

    .dataframe thead th {
        text-align: right;
    }
</style>
<table border="1" class="dataframe">
  <thead>
    <tr style="text-align: right;">
      <th></th>
      <th>Source</th>
      <th>Species</th>
      <th>Tissue</th>
      <th>Batch</th>
      <th>Peptide</th>
      <th>ProteinID</th>
      <th>GeneModel</th>
      <th>Chromosome</th>
      <th>Validation_status</th>
      <th>Projection_file</th>
      <th>Check_BED_geometry</th>
      <th>Check_block_nt_length</th>
      <th>Check_chromosome_and_strand</th>
      <th>Check_protein_coordinates</th>
      <th>All_sanity_checks_passed</th>
      <th>Sanity_check_status</th>
      <th>Chromosome_norm</th>
    </tr>
  </thead>
  <tbody>
    <tr>
      <th>0</th>
      <td>PXD004720</td>
      <td>bread wheat</td>
      <td>anther</td>
      <td>single</td>
      <td>ADLHHLAMFLAGRQPDAPQEALQVLDIVLR</td>
      <td>TraesCSU03G0061400.2</td>
      <td>TraesCSU03G0061400</td>
      <td>ChrUnknown</td>
      <td>validated</td>
      <td>FragPipe_Duncan_PXD004720_anther_peptide_genom...</td>
      <td>True</td>
      <td>True</td>
      <td>False</td>
      <td>True</td>
      <td>False</td>
      <td>failed</td>
      <td>ChrUnknown</td>
    </tr>
    <tr>
      <th>1</th>
      <td>PXD004720</td>
      <td>bread wheat</td>
      <td>anther</td>
      <td>single</td>
      <td>AKQMAESFNMSTFANREDLLGQSK</td>
      <td>TraesCSU03G0111000.1</td>
      <td>TraesCSU03G0111000</td>
      <td>ChrUnknown</td>
      <td>validated</td>
      <td>FragPipe_Duncan_PXD004720_anther_peptide_genom...</td>
      <td>True</td>
      <td>True</td>
      <td>False</td>
      <td>True</td>
      <td>False</td>
      <td>failed</td>
      <td>ChrUnknown</td>
    </tr>
    <tr>
      <th>2</th>
      <td>PXD004720</td>
      <td>bread wheat</td>
      <td>anther</td>
      <td>single</td>
      <td>AHEELVK</td>
      <td>TraesCSU03G0430400LC.1</td>
      <td>TraesCSU03G0430400LC</td>
      <td>ChrUnknown</td>
      <td>validated</td>
      <td>FragPipe_Duncan_PXD004720_anther_peptide_genom...</td>
      <td>True</td>
      <td>True</td>
      <td>False</td>
      <td>True</td>
      <td>False</td>
      <td>failed</td>
      <td>ChrUnknown</td>
    </tr>
    <tr>
      <th>3</th>
      <td>PXD004720</td>
      <td>bread wheat</td>
      <td>anther</td>
      <td>single</td>
      <td>AAAMGVEEAVR</td>
      <td>TraesCSU03G0402100.1</td>
      <td>TraesCSU03G0402100</td>
      <td>ChrUnknown</td>
      <td>validated</td>
      <td>FragPipe_Duncan_PXD004720_anther_peptide_genom...</td>
      <td>True</td>
      <td>True</td>
      <td>False</td>
      <td>True</td>
      <td>False</td>
      <td>failed</td>
      <td>ChrUnknown</td>
    </tr>
    <tr>
      <th>4</th>
      <td>PXD004720</td>
      <td>bread wheat</td>
      <td>anther</td>
      <td>single</td>
      <td>APGIIER</td>
      <td>TraesCSU03G0138700LC.1</td>
      <td>TraesCSU03G0138700LC</td>
      <td>ChrUnknown</td>
      <td>validated</td>
      <td>FragPipe_Duncan_PXD004720_anther_peptide_genom...</td>
      <td>True</td>
      <td>True</td>
      <td>False</td>
      <td>True</td>
      <td>False</td>
      <td>failed</td>
      <td>ChrUnknown</td>
    </tr>
    <tr>
      <th>5</th>
      <td>PXD004720</td>
      <td>bread wheat</td>
      <td>anther</td>
      <td>single</td>
      <td>DSDMQSADGDGR</td>
      <td>TraesCSU03G0144900LC.1</td>
      <td>TraesCSU03G0144900LC</td>
      <td>ChrUnknown</td>
      <td>validated</td>
      <td>FragPipe_Duncan_PXD004720_anther_peptide_genom...</td>
      <td>True</td>
      <td>True</td>
      <td>False</td>
      <td>True</td>
      <td>False</td>
      <td>failed</td>
      <td>ChrUnknown</td>
    </tr>
    <tr>
      <th>6</th>
      <td>PXD004720</td>
      <td>bread wheat</td>
      <td>anther</td>
      <td>single</td>
      <td>AVGVSTEK</td>
      <td>TraesCSU03G0112900LC.1</td>
      <td>TraesCSU03G0112900LC</td>
      <td>ChrUnknown</td>
      <td>validated</td>
      <td>FragPipe_Duncan_PXD004720_anther_peptide_genom...</td>
      <td>True</td>
      <td>True</td>
      <td>False</td>
      <td>True</td>
      <td>False</td>
      <td>failed</td>
      <td>ChrUnknown</td>
    </tr>
    <tr>
      <th>7</th>
      <td>PXD004720</td>
      <td>bread wheat</td>
      <td>anther</td>
      <td>single</td>
      <td>APGIIER</td>
      <td>TraesCSU03G0351700LC.1</td>
      <td>TraesCSU03G0351700LC</td>
      <td>ChrUnknown</td>
      <td>validated</td>
      <td>FragPipe_Duncan_PXD004720_anther_peptide_genom...</td>
      <td>True</td>
      <td>True</td>
      <td>False</td>
      <td>True</td>
      <td>False</td>
      <td>failed</td>
      <td>ChrUnknown</td>
    </tr>
    <tr>
      <th>8</th>
      <td>PXD004720</td>
      <td>bread wheat</td>
      <td>anther</td>
      <td>single</td>
      <td>AEHLASIFGTEKDR</td>
      <td>TraesCSU03G0553300LC.1</td>
      <td>TraesCSU03G0553300LC</td>
      <td>ChrUnknown</td>
      <td>validated</td>
      <td>FragPipe_Duncan_PXD004720_anther_peptide_genom...</td>
      <td>True</td>
      <td>True</td>
      <td>False</td>
      <td>True</td>
      <td>False</td>
      <td>failed</td>
      <td>ChrUnknown</td>
    </tr>
    <tr>
      <th>9</th>
      <td>PXD004720</td>
      <td>bread wheat</td>
      <td>anther</td>
      <td>single</td>
      <td>AVANETGAFFFLINGPEIMSK</td>
      <td>TraesCSU03G0001700.1</td>
      <td>TraesCSU03G0001700</td>
      <td>ChrUnknown</td>
      <td>validated</td>
      <td>FragPipe_Duncan_PXD004720_anther_peptide_genom...</td>
      <td>True</td>
      <td>True</td>
      <td>False</td>
      <td>True</td>
      <td>False</td>
      <td>failed</td>
      <td>ChrUnknown</td>
    </tr>
    <tr>
      <th>10</th>
      <td>PXD004720</td>
      <td>bread wheat</td>
      <td>anther</td>
      <td>single</td>
      <td>DNGMHALIIYDDLSK</td>
      <td>TraesCSU03G0519600LC.1</td>
      <td>TraesCSU03G0519600LC</td>
      <td>ChrUnknown</td>
      <td>validated</td>
      <td>FragPipe_Duncan_PXD004720_anther_peptide_genom...</td>
      <td>True</td>
      <td>True</td>
      <td>False</td>
      <td>True</td>
      <td>False</td>
      <td>failed</td>
      <td>ChrUnknown</td>
    </tr>
    <tr>
      <th>11</th>
      <td>PXD004720</td>
      <td>bread wheat</td>
      <td>anther</td>
      <td>single</td>
      <td>ASVAPPTVGKER</td>
      <td>TraesCSU03G0433300LC.1</td>
      <td>TraesCSU03G0433300LC</td>
      <td>ChrUnknown</td>
      <td>validated</td>
      <td>FragPipe_Duncan_PXD004720_anther_peptide_genom...</td>
      <td>True</td>
      <td>True</td>
      <td>False</td>
      <td>True</td>
      <td>False</td>
      <td>failed</td>
      <td>ChrUnknown</td>
    </tr>
    <tr>
      <th>12</th>
      <td>PXD004720</td>
      <td>bread wheat</td>
      <td>anther</td>
      <td>single</td>
      <td>AFTELLK</td>
      <td>TraesCSU03G0362800LC.1</td>
      <td>TraesCSU03G0362800LC</td>
      <td>ChrUnknown</td>
      <td>validated</td>
      <td>FragPipe_Duncan_PXD004720_anther_peptide_genom...</td>
      <td>True</td>
      <td>True</td>
      <td>False</td>
      <td>True</td>
      <td>False</td>
      <td>failed</td>
      <td>ChrUnknown</td>
    </tr>
    <tr>
      <th>13</th>
      <td>PXD004720</td>
      <td>bread wheat</td>
      <td>anther</td>
      <td>single</td>
      <td>AVTSPSAR</td>
      <td>TraesCSU03G0279300.1</td>
      <td>TraesCSU03G0279300</td>
      <td>ChrUnknown</td>
      <td>validated</td>
      <td>FragPipe_Duncan_PXD004720_anther_peptide_genom...</td>
      <td>True</td>
      <td>True</td>
      <td>False</td>
      <td>True</td>
      <td>False</td>
      <td>failed</td>
      <td>ChrUnknown</td>
    </tr>
    <tr>
      <th>14</th>
      <td>PXD004720</td>
      <td>bread wheat</td>
      <td>anther</td>
      <td>single</td>
      <td>ADGFAGVFPEHKYEIVK</td>
      <td>TraesCSU03G0076400.4</td>
      <td>TraesCSU03G0076400</td>
      <td>ChrUnknown</td>
      <td>validated</td>
      <td>FragPipe_Duncan_PXD004720_anther_peptide_genom...</td>
      <td>True</td>
      <td>True</td>
      <td>False</td>
      <td>True</td>
      <td>False</td>
      <td>failed</td>
      <td>ChrUnknown</td>
    </tr>
    <tr>
      <th>15</th>
      <td>PXD004720</td>
      <td>bread wheat</td>
      <td>anther</td>
      <td>single</td>
      <td>AHVIVMGATNRPNSIDPALR</td>
      <td>TraesCSU03G0001700.1</td>
      <td>TraesCSU03G0001700</td>
      <td>ChrUnknown</td>
      <td>validated</td>
      <td>FragPipe_Duncan_PXD004720_anther_peptide_genom...</td>
      <td>True</td>
      <td>True</td>
      <td>False</td>
      <td>True</td>
      <td>False</td>
      <td>failed</td>
      <td>ChrUnknown</td>
    </tr>
    <tr>
      <th>16</th>
      <td>PXD004720</td>
      <td>bread wheat</td>
      <td>anther</td>
      <td>single</td>
      <td>AAGIPSK</td>
      <td>TraesCSU03G0054300.1</td>
      <td>TraesCSU03G0054300</td>
      <td>ChrUnknown</td>
      <td>validated</td>
      <td>FragPipe_Duncan_PXD004720_anther_peptide_genom...</td>
      <td>True</td>
      <td>True</td>
      <td>False</td>
      <td>True</td>
      <td>False</td>
      <td>failed</td>
      <td>ChrUnknown</td>
    </tr>
    <tr>
      <th>17</th>
      <td>PXD004720</td>
      <td>bread wheat</td>
      <td>anther</td>
      <td>single</td>
      <td>AGVYHPWGPGNGRR</td>
      <td>TraesCSU03G0292000LC.1</td>
      <td>TraesCSU03G0292000LC</td>
      <td>ChrUnknown</td>
      <td>validated</td>
      <td>FragPipe_Duncan_PXD004720_anther_peptide_genom...</td>
      <td>True</td>
      <td>True</td>
      <td>False</td>
      <td>True</td>
      <td>False</td>
      <td>failed</td>
      <td>ChrUnknown</td>
    </tr>
    <tr>
      <th>18</th>
      <td>PXD004720</td>
      <td>bread wheat</td>
      <td>anther</td>
      <td>single</td>
      <td>AAQLSGR</td>
      <td>TraesCSU03G0023500LC.1</td>
      <td>TraesCSU03G0023500LC</td>
      <td>ChrUnknown</td>
      <td>validated</td>
      <td>FragPipe_Duncan_PXD004720_anther_peptide_genom...</td>
      <td>True</td>
      <td>True</td>
      <td>False</td>
      <td>True</td>
      <td>False</td>
      <td>failed</td>
      <td>ChrUnknown</td>
    </tr>
    <tr>
      <th>19</th>
      <td>PXD004720</td>
      <td>bread wheat</td>
      <td>anther</td>
      <td>single</td>
      <td>AAVFQMEHNKAP</td>
      <td>TraesCSU03G0002700LC.1</td>
      <td>TraesCSU03G0002700LC</td>
      <td>ChrUnknown</td>
      <td>validated</td>
      <td>FragPipe_Duncan_PXD004720_anther_peptide_genom...</td>
      <td>True</td>
      <td>True</td>
      <td>False</td>
      <td>True</td>
      <td>False</td>
      <td>failed</td>
      <td>ChrUnknown</td>
    </tr>
  </tbody>
</table>
</div>


# Step 12 — Export BED6 and BED12 Files for JBrowse

This step converts successfully projected peptide genomic coordinates from Step 10 into BED files for genome browser visualisation.

Both BED6 and BED12 formats were generated for each wheat tissue.

---

## Input files

### Peptide genome projection table from Step 11


```text
wheat_projection_validation_stratified100percent_step10.csv
```

Only rows with translation-validated genomic projection were exported.

### Filtering rule

```text
Projection_status == "projected"
```

---

## BED output formats

### BED6

BED6 provides a compact representation of each peptide genomic interval.

| BED6 field | Description |
|---|---|
| chrom | Chromosome or genome sequence ID |
| chromStart | 0-based genomic start coordinate |
| chromEnd | 0-based exclusive genomic end coordinate |
| name | Peptide, protein, and gene identifier |
| score | BED score scaled from peptide probability where available |
| strand | Genomic strand |

---

### BED12

BED12 provides a block-aware representation of peptide genomic coordinates.

This is particularly important for peptides spanning CDS junctions, because such peptides need to be displayed as multi-block genomic features.

| BED12 field | Description |
|---|---|
| chrom | Chromosome or genome sequence ID |
| chromStart | 0-based genomic start coordinate |
| chromEnd | 0-based exclusive genomic end coordinate |
| name | Peptide, protein, and gene identifier |
| score | BED score scaled from peptide probability where available |
| strand | Genomic strand |
| thickStart | Start of displayed feature |
| thickEnd | End of displayed feature |
| itemRgb | RGB colour field |
| blockCount | Number of genomic blocks |
| blockSizes | Comma-separated block sizes |
| blockStarts | Comma-separated block starts relative to chromStart |

---

## BED score

The BED score field was generated as follows:

```text
Probability × 1000
```

where peptide probability was available.

If no probability column was present, a default score of `1000` was assigned.

---

## Output files

BED files were exported into:

```text
python_outputs/bed/
```

### Example BED6 output

```text
FragPipe_Duncan_PXD004720_anther_peptides.bed6
```

### Example BED12 output

```text
FragPipe_Duncan_PXD004720_anther_peptides.bed12
```

---

## Summary metrics

A Step 10 summary table was generated across all tissues.

### Metrics captured

| Metric | Description |
|---|---|
| BED_rows | Number of peptide genomic features exported |
| Unique_BED_peptides | Number of unique peptide sequences exported |
| Unique_BED_proteins | Number of unique protein accessions represented |
| Unique_BED_gene_models | Number of unique gene models represented |
| Multi_block_peptides | Number of peptide features spanning more than one genomic block |

### Output summary file

```text
wheat_bed_export_summary_step12.csv
```

The resulting BED6 and BED12 files can be loaded into JBrowse to visualise peptide evidence aligned to the wheat genome.


```python
# ============================================================
# Step 12 — Export fully validated BED6 and BED12 files for JBrowse (takes 30 min)
# Translation-validated + sanity-check-passed projections only
# ============================================================

import pandas as pd
from pathlib import Path

# -----------------------------
# 1. Input / output paths
# -----------------------------
fragpipe_dir = Path("FragPipe_results")
tables_dir = Path("python_outputs/tables")

# Final BED outputs after both validation layers:
# Step 10 translation validation + Step 11 sanity checks
bed_dir = Path("python_outputs/bed_validated")
bed_dir.mkdir(parents=True, exist_ok=True)

manifest_file = fragpipe_dir / "wheat_tissues_FragPipe-result-manifest_2026-05-11.csv"

# Step 11 output: full sanity-check table built from Step 10 translation-validated rows
sanity_file = tables_dir / "wheat_projection_translation_validated_sanity_checks_full_step11.csv"

# Step 12 output summary
step12_summary_out = tables_dir / "wheat_bed_export_validated_summary_step12.csv"

chunk_size = 100_000

manifest = pd.read_csv(manifest_file, encoding="utf-8-sig")

if not sanity_file.exists():
    raise FileNotFoundError(
        f"Step 11 sanity-check file not found:\n{sanity_file}\n\n"
        "Please run Step 11 first."
    )


# -----------------------------
# 2. Helper functions
# -----------------------------
def make_bed_score(data):
    """
    Create BED score between 0 and 1000.

    Priority:
    1. Probability column scaled to 0–1000, if available
    2. Default score = 1000
    """

    if "Probability" in data.columns:
        score = pd.to_numeric(
            data["Probability"],
            errors="coerce"
        ) * 1000

        score = (
            score
            .fillna(1000)
            .clip(0, 1000)
            .round()
            .astype(int)
        )

    else:
        score = pd.Series([1000] * len(data), index=data.index)

    return score


def clean_bed_name(value):
    """
    Clean BED name field for BED/JBrowse compatibility.
    """

    return (
        str(value)
        .replace(" ", "_")
        .replace(";", "|")
        .replace(",", "|")
        .replace("\t", "_")
        .replace("\n", "_")
    )


def build_bed_name(projected):
    """
    Build informative BED label.

    Priority:
    1. Peptide_intron_gapped, if present
    2. Peptide_intron_gapped_compact, if present
    3. Peptide

    Final structure:
    peptide|protein|gene|validated=translation+sanity|tissues=X
    """

    if "Peptide_intron_gapped" in projected.columns:
        peptide_label = projected["Peptide_intron_gapped"].astype(str)
    elif "Peptide_intron_gapped_compact" in projected.columns:
        peptide_label = projected["Peptide_intron_gapped_compact"].astype(str)
    else:
        peptide_label = projected["Peptide"].astype(str)

    if "GeneID" in projected.columns:
        bed_name = (
            peptide_label + "|" +
            projected["ProteinID"].astype(str) + "|" +
            projected["GeneID"].astype(str)
        )
    else:
        bed_name = (
            peptide_label + "|" +
            projected["ProteinID"].astype(str)
        )

    # Make final BED tracks self-documenting
    bed_name = bed_name + "|validated=translation+sanity"

    # Add tissue count if present
    if "Tissues_count" in projected.columns:
        bed_name = (
            bed_name +
            "|tissues=" +
            projected["Tissues_count"].astype(str)
        )

    return bed_name.apply(clean_bed_name)


def prepare_bed_rows(projected):
    """
    Prepare a sanity-passed DataFrame for BED6/BED12 export.
    """

    required_cols = [
        "Chromosome",
        "BED_start_0based",
        "BED_end_0based_exclusive",
        "ProteinID",
        "Strand",
        "BED_block_count",
        "BED_block_sizes",
        "BED_block_starts",
        "Peptide",
        "Sanity_check_status",
        "All_sanity_checks_passed"
    ]

    missing_cols = [
        col for col in required_cols
        if col not in projected.columns
    ]

    if missing_cols:
        raise KeyError(
            f"Missing required BED column(s): {missing_cols}"
        )

    projected = projected.copy()

    # Create BED score and label
    projected["BED_score"] = make_bed_score(projected)
    projected["BED_name"] = build_bed_name(projected)

    # Force integer coordinate types
    projected["BED_start_0based"] = (
        pd.to_numeric(projected["BED_start_0based"], errors="coerce")
        .astype("Int64")
    )

    projected["BED_end_0based_exclusive"] = (
        pd.to_numeric(projected["BED_end_0based_exclusive"], errors="coerce")
        .astype("Int64")
    )

    projected["BED_block_count"] = (
        pd.to_numeric(projected["BED_block_count"], errors="coerce")
        .astype("Int64")
    )

    # Drop rows with missing essential BED fields, just in case
    before_drop = len(projected)

    projected = projected.dropna(
        subset=[
            "Chromosome",
            "BED_start_0based",
            "BED_end_0based_exclusive",
            "Strand",
            "BED_block_count",
            "BED_block_sizes",
            "BED_block_starts"
        ]
    ).copy()

    dropped_missing_bed_fields = before_drop - len(projected)

    projected["BED_start_0based"] = projected["BED_start_0based"].astype(int)
    projected["BED_end_0based_exclusive"] = projected["BED_end_0based_exclusive"].astype(int)
    projected["BED_block_count"] = projected["BED_block_count"].astype(int)

    return projected, dropped_missing_bed_fields


def make_bed6(projected):
    """
    Build BED6 DataFrame.
    """

    return projected[[
        "Chromosome",
        "BED_start_0based",
        "BED_end_0based_exclusive",
        "BED_name",
        "BED_score",
        "Strand"
    ]].copy()


def make_bed12(projected):
    """
    Build BED12 DataFrame.
    """

    return pd.DataFrame({

        "chrom":
            projected["Chromosome"],

        "chromStart":
            projected["BED_start_0based"],

        "chromEnd":
            projected["BED_end_0based_exclusive"],

        "name":
            projected["BED_name"],

        "score":
            projected["BED_score"],

        "strand":
            projected["Strand"],

        "thickStart":
            projected["BED_start_0based"],

        "thickEnd":
            projected["BED_end_0based_exclusive"],

        "itemRgb":
            "0",

        "blockCount":
            projected["BED_block_count"],

        "blockSizes":
            projected["BED_block_sizes"],

        "blockStarts":
            projected["BED_block_starts"]
    })


# -----------------------------
# 3. Build manifest lookup and output file names
# -----------------------------
manifest_lookup = {}

for _, row in manifest.iterrows():

    projection_filename = row["FragPipe-Output-Peptide"].replace(
        "_peptide.tsv",
        "_peptide_genome_projection.csv"
    )

    bed6_filename = projection_filename.replace(
        "_peptide_genome_projection.csv",
        "_validated_peptides.bed6"
    )

    bed12_filename = projection_filename.replace(
        "_peptide_genome_projection.csv",
        "_validated_peptides.bed12"
    )

    manifest_lookup[projection_filename] = {
        "Source": row["Source"],
        "Species": row["Species"],
        "Tissue": row["Tissue-Raw-Code"],
        "Batch": row["Batch"],
        "BED6_file": bed6_filename,
        "BED12_file": bed12_filename,
        "BED6_path": bed_dir / bed6_filename,
        "BED12_path": bed_dir / bed12_filename
    }


# Clear previous BED outputs if rerunning this step
for info in manifest_lookup.values():
    for path_key in ["BED6_path", "BED12_path"]:
        path = info[path_key]
        if path.exists():
            path.unlink()

if step12_summary_out.exists():
    step12_summary_out.unlink()


# -----------------------------
# 4. Initialise summary dictionary
# -----------------------------
summary_dict = {}

for projection_filename, info in manifest_lookup.items():

    summary_dict[projection_filename] = {
        "Source": info["Source"],
        "Species": info["Species"],
        "Tissue": info["Tissue"],
        "Batch": info["Batch"],
        "Projection_file": projection_filename,
        "BED6_file": info["BED6_file"],
        "BED12_file": info["BED12_file"],

        "Rows_in_step11_sanity_file": 0,
        "Rows_passing_all_sanity_checks": 0,
        "Rows_excluded_by_sanity_checks": 0,
        "Rows_dropped_missing_BED_fields": 0,

        "BED_rows": 0,
        "Unique_BED_peptides": 0,
        "Unique_BED_proteins": 0,
        "Unique_BED_gene_models": 0,
        "Multi_block_peptides": 0,
        "Within_exon_peptides": 0,
        "Intron_spanning_BED_rows": 0,
        "Within_exon_BED_rows": 0,
        "Unique_intron_spanning_peptides": 0,
        "Unique_within_exon_peptides": 0,
        "BED_labels_with_introns": 0
    }


# For unique counts across chunks
unique_peptides = {k: set() for k in manifest_lookup}
unique_proteins = {k: set() for k in manifest_lookup}
unique_genes = {k: set() for k in manifest_lookup}
unique_intron_spanning_peptides = {k: set() for k in manifest_lookup}
unique_within_exon_peptides = {k: set() for k in manifest_lookup}

# Track whether each BED file has already been written to
bed_file_written = {
    projection_filename: {
        "bed6": False,
        "bed12": False
    }
    for projection_filename in manifest_lookup
}


# -----------------------------
# 5. Read Step 11 sanity-check output in chunks and export BED files
# -----------------------------
print("\nExporting BED6/BED12 files from Step 11 sanity-passed rows...")

header = pd.read_csv(sanity_file, nrows=0)

required_step11_cols = [
    "Projection_file",
    "Sanity_check_status",
    "All_sanity_checks_passed"
]

missing_step11_cols = [
    col for col in required_step11_cols
    if col not in header.columns
]

if missing_step11_cols:
    raise KeyError(
        f"Missing required Step 11 column(s): {missing_step11_cols}"
    )

total_rows_read = 0
total_sanity_passed = 0
total_sanity_failed = 0
total_bed_rows = 0

for chunk_i, chunk in enumerate(
    pd.read_csv(
        sanity_file,
        chunksize=chunk_size,
        low_memory=False
    ),
    start=1
):

    total_rows_read += len(chunk)

    # Count sanity status per projection file
    for projection_filename, file_group in chunk.groupby("Projection_file"):

        projection_filename = str(projection_filename)

        if projection_filename not in summary_dict:
            continue

        n_rows = len(file_group)
        n_passed = int((file_group["Sanity_check_status"].astype(str) == "passed").sum())
        n_failed = n_rows - n_passed

        summary_dict[projection_filename]["Rows_in_step11_sanity_file"] += n_rows
        summary_dict[projection_filename]["Rows_passing_all_sanity_checks"] += n_passed
        summary_dict[projection_filename]["Rows_excluded_by_sanity_checks"] += n_failed

    # Keep only rows passing all sanity checks
    passed = chunk[
        chunk["Sanity_check_status"].astype(str) == "passed"
    ].copy()

    total_sanity_passed += len(passed)
    total_sanity_failed += len(chunk) - len(passed)

    if passed.empty:
        print(
            f"Chunk {chunk_i}: read {len(chunk):,} rows | "
            f"no sanity-passed rows"
        )
        continue

    # Export per projection/source-tissue file
    for projection_filename, projected in passed.groupby("Projection_file"):

        projection_filename = str(projection_filename)

        if projection_filename not in manifest_lookup:
            print(f"  Warning: projection file not found in manifest, skipped: {projection_filename}")
            continue

        info = manifest_lookup[projection_filename]

        projected_prepared, dropped_missing_bed_fields = prepare_bed_rows(projected)

        if projected_prepared.empty:
            summary_dict[projection_filename]["Rows_dropped_missing_BED_fields"] += dropped_missing_bed_fields
            continue

        bed6 = make_bed6(projected_prepared)
        bed12 = make_bed12(projected_prepared)

        # Append to BED6
        bed6.to_csv(
            info["BED6_path"],
            sep="\t",
            header=False,
            index=False,
            mode="a"
        )

        # Append to BED12
        bed12.to_csv(
            info["BED12_path"],
            sep="\t",
            header=False,
            index=False,
            mode="a"
        )

        # Update summary
        n_bed = len(projected_prepared)

        summary_dict[projection_filename]["BED_rows"] += n_bed
        summary_dict[projection_filename]["Rows_dropped_missing_BED_fields"] += dropped_missing_bed_fields

        if "Peptide" in projected_prepared.columns:
            unique_peptides[projection_filename].update(
                projected_prepared["Peptide"].dropna().astype(str).unique()
            )

        if "ProteinID" in projected_prepared.columns:
            unique_proteins[projection_filename].update(
                projected_prepared["ProteinID"].dropna().astype(str).unique()
            )

        if "GeneID" in projected_prepared.columns:
            unique_genes[projection_filename].update(
                projected_prepared["GeneID"].dropna().astype(str).unique()
            )

        # Count exon structure of peptide projections
        block_count = pd.to_numeric(
            projected_prepared["BED_block_count"],
            errors="coerce"
        )
        
        intron_spanning_mask = block_count > 1
        within_exon_mask = block_count == 1
        
        intron_spanning_rows = int(intron_spanning_mask.sum())
        within_exon_rows = int(within_exon_mask.sum())
        
        # Existing legacy-style count retained for continuity
        summary_dict[projection_filename]["Multi_block_peptides"] += intron_spanning_rows
        
        # New clearer paired counts
        summary_dict[projection_filename]["Within_exon_peptides"] += within_exon_rows
        summary_dict[projection_filename]["Intron_spanning_BED_rows"] += intron_spanning_rows
        summary_dict[projection_filename]["Within_exon_BED_rows"] += within_exon_rows
        
        # Unique peptide-sequence counts by exon structure
        if "Peptide" in projected_prepared.columns:
        
            unique_intron_spanning_peptides[projection_filename].update(
                projected_prepared.loc[
                    intron_spanning_mask,
                    "Peptide"
                ].dropna().astype(str).unique()
            )
        
            unique_within_exon_peptides[projection_filename].update(
                projected_prepared.loc[
                    within_exon_mask,
                    "Peptide"
                ].dropna().astype(str).unique()
            )
        
        # BED labels containing dashes remain useful as a visual proxy,
        # but BED_block_count is the authoritative exon-structure count.
        summary_dict[projection_filename]["BED_labels_with_introns"] += int(
            projected_prepared["BED_name"]
            .astype(str)
            .str.contains("-", regex=False)
            .sum()
        )

        total_bed_rows += n_bed

    print(
        f"Chunk {chunk_i}: read {len(chunk):,} rows | "
        f"sanity-passed {len(passed):,} | "
        f"cumulative BED rows {total_bed_rows:,}"
    )


# -----------------------------
# 6. Finalise summary table
# -----------------------------
for projection_filename in summary_dict:

    summary_dict[projection_filename]["Unique_BED_peptides"] = len(
        unique_peptides[projection_filename]
    )

    summary_dict[projection_filename]["Unique_BED_proteins"] = len(
        unique_proteins[projection_filename]
    )

    summary_dict[projection_filename]["Unique_BED_gene_models"] = len(
        unique_genes[projection_filename]
    )

    summary_dict[projection_filename]["Unique_intron_spanning_peptides"] = len(
        unique_intron_spanning_peptides[projection_filename]
    )

    summary_dict[projection_filename]["Unique_within_exon_peptides"] = len(
        unique_within_exon_peptides[projection_filename]
    )

step12_summary = pd.DataFrame(summary_dict.values())

step12_summary["Percent_sanity_passed"] = (
    step12_summary["Rows_passing_all_sanity_checks"] /
    step12_summary["Rows_in_step11_sanity_file"] *
    100
).round(4)

step12_summary["Percent_exported_to_BED"] = (
    step12_summary["BED_rows"] /
    step12_summary["Rows_in_step11_sanity_file"] *
    100
).round(4)

step12_summary.to_csv(
    step12_summary_out,
    index=False
)


# -----------------------------
# 7. Overall summary
# -----------------------------
overall_bed_files_bed6 = len(list(bed_dir.glob("*_validated_peptides.bed6")))
overall_bed_files_bed12 = len(list(bed_dir.glob("*_validated_peptides.bed12")))

print("\n===== STEP 12 VALIDATED BED EXPORT SUMMARY =====")
print(f"Rows read from Step 11 sanity-check table: {total_rows_read:,}")
print(f"Rows passing all sanity checks: {total_sanity_passed:,}")
print(f"Rows excluded by sanity checks: {total_sanity_failed:,}")
print(f"Rows exported to BED: {total_bed_rows:,}")
print(f"BED6 files created: {overall_bed_files_bed6:,}")
print(f"BED12 files created: {overall_bed_files_bed12:,}")

print(f"\nBED files saved in: {bed_dir}")
print(f"Step 12 summary saved: {step12_summary_out}")

display(step12_summary)
```

    
    Exporting BED6/BED12 files from Step 11 sanity-passed rows...
    Chunk 1: read 100,000 rows | sanity-passed 99,103 | cumulative BED rows 99,103
    Chunk 2: read 100,000 rows | sanity-passed 99,042 | cumulative BED rows 198,145
    Chunk 3: read 100,000 rows | sanity-passed 98,965 | cumulative BED rows 297,110
    Chunk 4: read 100,000 rows | sanity-passed 99,072 | cumulative BED rows 396,182
    Chunk 5: read 100,000 rows | sanity-passed 99,141 | cumulative BED rows 495,323
    Chunk 6: read 100,000 rows | sanity-passed 98,838 | cumulative BED rows 594,161
    Chunk 7: read 100,000 rows | sanity-passed 99,027 | cumulative BED rows 693,188
    Chunk 8: read 100,000 rows | sanity-passed 99,136 | cumulative BED rows 792,324
    Chunk 9: read 100,000 rows | sanity-passed 99,224 | cumulative BED rows 891,548
    Chunk 10: read 100,000 rows | sanity-passed 99,254 | cumulative BED rows 990,802
    Chunk 11: read 100,000 rows | sanity-passed 99,160 | cumulative BED rows 1,089,962
    Chunk 12: read 100,000 rows | sanity-passed 99,088 | cumulative BED rows 1,189,050
    Chunk 13: read 100,000 rows | sanity-passed 99,193 | cumulative BED rows 1,288,243
    Chunk 14: read 100,000 rows | sanity-passed 98,905 | cumulative BED rows 1,387,148
    Chunk 15: read 100,000 rows | sanity-passed 98,837 | cumulative BED rows 1,485,985
    Chunk 16: read 100,000 rows | sanity-passed 98,897 | cumulative BED rows 1,584,882
    Chunk 17: read 100,000 rows | sanity-passed 98,979 | cumulative BED rows 1,683,861
    Chunk 18: read 100,000 rows | sanity-passed 99,059 | cumulative BED rows 1,782,920
    Chunk 19: read 100,000 rows | sanity-passed 98,967 | cumulative BED rows 1,881,887
    Chunk 20: read 100,000 rows | sanity-passed 98,810 | cumulative BED rows 1,980,697
    Chunk 21: read 100,000 rows | sanity-passed 98,925 | cumulative BED rows 2,079,622
    Chunk 22: read 100,000 rows | sanity-passed 99,018 | cumulative BED rows 2,178,640
    Chunk 23: read 100,000 rows | sanity-passed 99,038 | cumulative BED rows 2,277,678
    Chunk 24: read 100,000 rows | sanity-passed 99,055 | cumulative BED rows 2,376,733
    Chunk 25: read 100,000 rows | sanity-passed 98,990 | cumulative BED rows 2,475,723
    Chunk 26: read 100,000 rows | sanity-passed 99,206 | cumulative BED rows 2,574,929
    Chunk 27: read 100,000 rows | sanity-passed 99,222 | cumulative BED rows 2,674,151
    Chunk 28: read 100,000 rows | sanity-passed 98,908 | cumulative BED rows 2,773,059
    Chunk 29: read 100,000 rows | sanity-passed 99,076 | cumulative BED rows 2,872,135
    Chunk 30: read 100,000 rows | sanity-passed 99,293 | cumulative BED rows 2,971,428
    Chunk 31: read 100,000 rows | sanity-passed 99,320 | cumulative BED rows 3,070,748
    Chunk 32: read 100,000 rows | sanity-passed 99,085 | cumulative BED rows 3,169,833
    Chunk 33: read 100,000 rows | sanity-passed 99,136 | cumulative BED rows 3,268,969
    Chunk 34: read 100,000 rows | sanity-passed 99,097 | cumulative BED rows 3,368,066
    Chunk 35: read 100,000 rows | sanity-passed 99,148 | cumulative BED rows 3,467,214
    Chunk 36: read 100,000 rows | sanity-passed 99,169 | cumulative BED rows 3,566,383
    Chunk 37: read 100,000 rows | sanity-passed 99,114 | cumulative BED rows 3,665,497
    Chunk 38: read 100,000 rows | sanity-passed 99,182 | cumulative BED rows 3,764,679
    Chunk 39: read 100,000 rows | sanity-passed 99,057 | cumulative BED rows 3,863,736
    Chunk 40: read 100,000 rows | sanity-passed 98,957 | cumulative BED rows 3,962,693
    Chunk 41: read 100,000 rows | sanity-passed 98,997 | cumulative BED rows 4,061,690
    Chunk 42: read 100,000 rows | sanity-passed 99,107 | cumulative BED rows 4,160,797
    Chunk 43: read 100,000 rows | sanity-passed 99,042 | cumulative BED rows 4,259,839
    Chunk 44: read 100,000 rows | sanity-passed 99,128 | cumulative BED rows 4,358,967
    Chunk 45: read 100,000 rows | sanity-passed 98,819 | cumulative BED rows 4,457,786
    Chunk 46: read 100,000 rows | sanity-passed 98,863 | cumulative BED rows 4,556,649
    Chunk 47: read 100,000 rows | sanity-passed 98,948 | cumulative BED rows 4,655,597
    Chunk 48: read 100,000 rows | sanity-passed 99,086 | cumulative BED rows 4,754,683
    Chunk 49: read 100,000 rows | sanity-passed 99,008 | cumulative BED rows 4,853,691
    Chunk 50: read 100,000 rows | sanity-passed 99,199 | cumulative BED rows 4,952,890
    Chunk 51: read 100,000 rows | sanity-passed 99,139 | cumulative BED rows 5,052,029
    Chunk 52: read 100,000 rows | sanity-passed 99,127 | cumulative BED rows 5,151,156
    Chunk 53: read 100,000 rows | sanity-passed 99,101 | cumulative BED rows 5,250,257
    Chunk 54: read 100,000 rows | sanity-passed 99,098 | cumulative BED rows 5,349,355
    Chunk 55: read 100,000 rows | sanity-passed 99,066 | cumulative BED rows 5,448,421
    Chunk 56: read 100,000 rows | sanity-passed 99,182 | cumulative BED rows 5,547,603
    Chunk 57: read 100,000 rows | sanity-passed 99,056 | cumulative BED rows 5,646,659
    Chunk 58: read 100,000 rows | sanity-passed 99,148 | cumulative BED rows 5,745,807
    Chunk 59: read 100,000 rows | sanity-passed 98,940 | cumulative BED rows 5,844,747
    Chunk 60: read 100,000 rows | sanity-passed 98,960 | cumulative BED rows 5,943,707
    Chunk 61: read 100,000 rows | sanity-passed 98,997 | cumulative BED rows 6,042,704
    Chunk 62: read 100,000 rows | sanity-passed 99,076 | cumulative BED rows 6,141,780
    Chunk 63: read 100,000 rows | sanity-passed 98,946 | cumulative BED rows 6,240,726
    Chunk 64: read 100,000 rows | sanity-passed 98,710 | cumulative BED rows 6,339,436
    Chunk 65: read 100,000 rows | sanity-passed 98,980 | cumulative BED rows 6,438,416
    Chunk 66: read 100,000 rows | sanity-passed 98,997 | cumulative BED rows 6,537,413
    Chunk 67: read 100,000 rows | sanity-passed 99,038 | cumulative BED rows 6,636,451
    Chunk 68: read 100,000 rows | sanity-passed 99,159 | cumulative BED rows 6,735,610
    Chunk 69: read 100,000 rows | sanity-passed 99,199 | cumulative BED rows 6,834,809
    Chunk 70: read 100,000 rows | sanity-passed 99,153 | cumulative BED rows 6,933,962
    Chunk 71: read 100,000 rows | sanity-passed 99,141 | cumulative BED rows 7,033,103
    Chunk 72: read 100,000 rows | sanity-passed 99,059 | cumulative BED rows 7,132,162
    Chunk 73: read 100,000 rows | sanity-passed 99,140 | cumulative BED rows 7,231,302
    Chunk 74: read 100,000 rows | sanity-passed 99,153 | cumulative BED rows 7,330,455
    Chunk 75: read 100,000 rows | sanity-passed 98,987 | cumulative BED rows 7,429,442
    Chunk 76: read 100,000 rows | sanity-passed 99,048 | cumulative BED rows 7,528,490
    Chunk 77: read 100,000 rows | sanity-passed 99,101 | cumulative BED rows 7,627,591
    Chunk 78: read 100,000 rows | sanity-passed 98,819 | cumulative BED rows 7,726,410
    Chunk 79: read 100,000 rows | sanity-passed 98,937 | cumulative BED rows 7,825,347
    Chunk 80: read 100,000 rows | sanity-passed 99,217 | cumulative BED rows 7,924,564
    Chunk 81: read 100,000 rows | sanity-passed 99,111 | cumulative BED rows 8,023,675
    Chunk 82: read 100,000 rows | sanity-passed 98,995 | cumulative BED rows 8,122,670
    Chunk 83: read 14,230 rows | sanity-passed 14,017 | cumulative BED rows 8,136,687
    
    ===== STEP 12 VALIDATED BED EXPORT SUMMARY =====
    Rows read from Step 11 sanity-check table: 8,214,230
    Rows passing all sanity checks: 8,136,687
    Rows excluded by sanity checks: 77,543
    Rows exported to BED: 8,136,687
    BED6 files created: 32
    BED12 files created: 32
    
    BED files saved in: python_outputs\bed_validated
    Step 12 summary saved: python_outputs\tables\wheat_bed_export_validated_summary_step12.csv
    


<div>
<style scoped>
    .dataframe tbody tr th:only-of-type {
        vertical-align: middle;
    }

    .dataframe tbody tr th {
        vertical-align: top;
    }

    .dataframe thead th {
        text-align: right;
    }
</style>
<table border="1" class="dataframe">
  <thead>
    <tr style="text-align: right;">
      <th></th>
      <th>Source</th>
      <th>Species</th>
      <th>Tissue</th>
      <th>Batch</th>
      <th>Projection_file</th>
      <th>BED6_file</th>
      <th>BED12_file</th>
      <th>Rows_in_step11_sanity_file</th>
      <th>Rows_passing_all_sanity_checks</th>
      <th>Rows_excluded_by_sanity_checks</th>
      <th>...</th>
      <th>Unique_BED_gene_models</th>
      <th>Multi_block_peptides</th>
      <th>Within_exon_peptides</th>
      <th>Intron_spanning_BED_rows</th>
      <th>Within_exon_BED_rows</th>
      <th>Unique_intron_spanning_peptides</th>
      <th>Unique_within_exon_peptides</th>
      <th>BED_labels_with_introns</th>
      <th>Percent_sanity_passed</th>
      <th>Percent_exported_to_BED</th>
    </tr>
  </thead>
  <tbody>
    <tr>
      <th>0</th>
      <td>MSV000090572</td>
      <td>bread wheat</td>
      <td>stored_grain</td>
      <td>single</td>
      <td>FragPipe_Vincent_MSV000090572_stored-grain_pep...</td>
      <td>FragPipe_Vincent_MSV000090572_stored-grain_val...</td>
      <td>FragPipe_Vincent_MSV000090572_stored-grain_val...</td>
      <td>29892</td>
      <td>29430</td>
      <td>462</td>
      <td>...</td>
      <td>0</td>
      <td>3248</td>
      <td>26182</td>
      <td>3248</td>
      <td>26182</td>
      <td>1077</td>
      <td>8024</td>
      <td>1795</td>
      <td>98.4544</td>
      <td>98.4544</td>
    </tr>
    <tr>
      <th>1</th>
      <td>PXD050500</td>
      <td>bread wheat</td>
      <td>coleoptile</td>
      <td>single</td>
      <td>FragPipe_Liu_PXD050500_coleoptile_peptide_geno...</td>
      <td>FragPipe_Liu_PXD050500_coleoptile_validated_pe...</td>
      <td>FragPipe_Liu_PXD050500_coleoptile_validated_pe...</td>
      <td>1829196</td>
      <td>1812027</td>
      <td>17169</td>
      <td>...</td>
      <td>0</td>
      <td>226291</td>
      <td>1585736</td>
      <td>226291</td>
      <td>1585736</td>
      <td>64943</td>
      <td>509164</td>
      <td>121256</td>
      <td>99.0614</td>
      <td>99.0614</td>
    </tr>
    <tr>
      <th>2</th>
      <td>PXD050500</td>
      <td>bread wheat</td>
      <td>node</td>
      <td>single</td>
      <td>FragPipe_Liu_PXD050500_node_peptide_genome_pro...</td>
      <td>FragPipe_Liu_PXD050500_node_validated_peptides...</td>
      <td>FragPipe_Liu_PXD050500_node_validated_peptides...</td>
      <td>1864415</td>
      <td>1846620</td>
      <td>17795</td>
      <td>...</td>
      <td>0</td>
      <td>233388</td>
      <td>1613232</td>
      <td>233388</td>
      <td>1613232</td>
      <td>66647</td>
      <td>520105</td>
      <td>125778</td>
      <td>99.0455</td>
      <td>99.0455</td>
    </tr>
    <tr>
      <th>3</th>
      <td>PXD050500</td>
      <td>bread wheat</td>
      <td>radicle</td>
      <td>single</td>
      <td>FragPipe_Liu_PXD050500_radicle_peptide_genome_...</td>
      <td>FragPipe_Liu_PXD050500_radicle_validated_pepti...</td>
      <td>FragPipe_Liu_PXD050500_radicle_validated_pepti...</td>
      <td>1050178</td>
      <td>1040357</td>
      <td>9821</td>
      <td>...</td>
      <td>0</td>
      <td>117056</td>
      <td>923301</td>
      <td>117056</td>
      <td>923301</td>
      <td>33950</td>
      <td>294305</td>
      <td>61713</td>
      <td>99.0648</td>
      <td>99.0648</td>
    </tr>
    <tr>
      <th>4</th>
      <td>PXD004720</td>
      <td>bread wheat</td>
      <td>anther</td>
      <td>single</td>
      <td>FragPipe_Duncan_PXD004720_anther_peptide_genom...</td>
      <td>FragPipe_Duncan_PXD004720_anther_validated_pep...</td>
      <td>FragPipe_Duncan_PXD004720_anther_validated_pep...</td>
      <td>163467</td>
      <td>161955</td>
      <td>1512</td>
      <td>...</td>
      <td>0</td>
      <td>27584</td>
      <td>134371</td>
      <td>27584</td>
      <td>134371</td>
      <td>6315</td>
      <td>27842</td>
      <td>16044</td>
      <td>99.0750</td>
      <td>99.0750</td>
    </tr>
    <tr>
      <th>5</th>
      <td>PXD004720</td>
      <td>bread wheat</td>
      <td>boot</td>
      <td>single</td>
      <td>FragPipe_Duncan_PXD004720_boot_peptide_genome_...</td>
      <td>FragPipe_Duncan_PXD004720_boot_validated_pepti...</td>
      <td>FragPipe_Duncan_PXD004720_boot_validated_pepti...</td>
      <td>13137</td>
      <td>13024</td>
      <td>113</td>
      <td>...</td>
      <td>0</td>
      <td>2718</td>
      <td>10306</td>
      <td>2718</td>
      <td>10306</td>
      <td>712</td>
      <td>2885</td>
      <td>1534</td>
      <td>99.1398</td>
      <td>99.1398</td>
    </tr>
    <tr>
      <th>6</th>
      <td>PXD004720</td>
      <td>bread wheat</td>
      <td>coleoptile</td>
      <td>single</td>
      <td>FragPipe_Duncan_PXD004720_coleoptile_peptide_g...</td>
      <td>FragPipe_Duncan_PXD004720_coleoptile_validated...</td>
      <td>FragPipe_Duncan_PXD004720_coleoptile_validated...</td>
      <td>203264</td>
      <td>201259</td>
      <td>2005</td>
      <td>...</td>
      <td>0</td>
      <td>31158</td>
      <td>170101</td>
      <td>31158</td>
      <td>170101</td>
      <td>6787</td>
      <td>34329</td>
      <td>18577</td>
      <td>99.0136</td>
      <td>99.0136</td>
    </tr>
    <tr>
      <th>7</th>
      <td>PXD004720</td>
      <td>bread wheat</td>
      <td>embryo</td>
      <td>single</td>
      <td>FragPipe_Duncan_PXD004720_embryo_peptide_genom...</td>
      <td>FragPipe_Duncan_PXD004720_embryo_validated_pep...</td>
      <td>FragPipe_Duncan_PXD004720_embryo_validated_pep...</td>
      <td>8742</td>
      <td>8662</td>
      <td>80</td>
      <td>...</td>
      <td>0</td>
      <td>1394</td>
      <td>7268</td>
      <td>1394</td>
      <td>7268</td>
      <td>383</td>
      <td>2438</td>
      <td>751</td>
      <td>99.0849</td>
      <td>99.0849</td>
    </tr>
    <tr>
      <th>8</th>
      <td>PXD004720</td>
      <td>bread wheat</td>
      <td>endosperm</td>
      <td>single</td>
      <td>FragPipe_Duncan_PXD004720_endosperm_peptide_ge...</td>
      <td>FragPipe_Duncan_PXD004720_endosperm_validated_...</td>
      <td>FragPipe_Duncan_PXD004720_endosperm_validated_...</td>
      <td>102289</td>
      <td>101413</td>
      <td>876</td>
      <td>...</td>
      <td>0</td>
      <td>14186</td>
      <td>87227</td>
      <td>14186</td>
      <td>87227</td>
      <td>3066</td>
      <td>17009</td>
      <td>7927</td>
      <td>99.1436</td>
      <td>99.1436</td>
    </tr>
    <tr>
      <th>9</th>
      <td>PXD004720</td>
      <td>bread wheat</td>
      <td>glume</td>
      <td>single</td>
      <td>FragPipe_Duncan_PXD004720_glume_peptide_genome...</td>
      <td>FragPipe_Duncan_PXD004720_glume_validated_pept...</td>
      <td>FragPipe_Duncan_PXD004720_glume_validated_pept...</td>
      <td>144079</td>
      <td>142449</td>
      <td>1630</td>
      <td>...</td>
      <td>0</td>
      <td>22004</td>
      <td>120445</td>
      <td>22004</td>
      <td>120445</td>
      <td>5084</td>
      <td>23614</td>
      <td>12693</td>
      <td>98.8687</td>
      <td>98.8687</td>
    </tr>
    <tr>
      <th>10</th>
      <td>PXD004720</td>
      <td>bread wheat</td>
      <td>grain-zadoks-70</td>
      <td>single</td>
      <td>FragPipe_Duncan_PXD004720_grain-zadoks-70_pept...</td>
      <td>FragPipe_Duncan_PXD004720_grain-zadoks-70_vali...</td>
      <td>FragPipe_Duncan_PXD004720_grain-zadoks-70_vali...</td>
      <td>125355</td>
      <td>124251</td>
      <td>1104</td>
      <td>...</td>
      <td>0</td>
      <td>17383</td>
      <td>106868</td>
      <td>17383</td>
      <td>106868</td>
      <td>3871</td>
      <td>22304</td>
      <td>9839</td>
      <td>99.1193</td>
      <td>99.1193</td>
    </tr>
    <tr>
      <th>11</th>
      <td>PXD004720</td>
      <td>bread wheat</td>
      <td>grain-zadoks-71</td>
      <td>single</td>
      <td>FragPipe_Duncan_PXD004720_grain-zadoks-71_pept...</td>
      <td>FragPipe_Duncan_PXD004720_grain-zadoks-71_vali...</td>
      <td>FragPipe_Duncan_PXD004720_grain-zadoks-71_vali...</td>
      <td>178814</td>
      <td>177392</td>
      <td>1422</td>
      <td>...</td>
      <td>0</td>
      <td>27486</td>
      <td>149906</td>
      <td>27486</td>
      <td>149906</td>
      <td>5774</td>
      <td>30241</td>
      <td>15867</td>
      <td>99.2048</td>
      <td>99.2048</td>
    </tr>
    <tr>
      <th>12</th>
      <td>PXD004720</td>
      <td>bread wheat</td>
      <td>grain-zadoks-75</td>
      <td>single</td>
      <td>FragPipe_Duncan_PXD004720_grain-zadoks-75_pept...</td>
      <td>FragPipe_Duncan_PXD004720_grain-zadoks-75_vali...</td>
      <td>FragPipe_Duncan_PXD004720_grain-zadoks-75_vali...</td>
      <td>132081</td>
      <td>131066</td>
      <td>1015</td>
      <td>...</td>
      <td>0</td>
      <td>18519</td>
      <td>112547</td>
      <td>18519</td>
      <td>112547</td>
      <td>3890</td>
      <td>23567</td>
      <td>10434</td>
      <td>99.2315</td>
      <td>99.2315</td>
    </tr>
    <tr>
      <th>13</th>
      <td>PXD004720</td>
      <td>bread wheat</td>
      <td>grain-zadoks-83</td>
      <td>single</td>
      <td>FragPipe_Duncan_PXD004720_grain-zadoks-83_pept...</td>
      <td>FragPipe_Duncan_PXD004720_grain-zadoks-83_vali...</td>
      <td>FragPipe_Duncan_PXD004720_grain-zadoks-83_vali...</td>
      <td>112389</td>
      <td>111335</td>
      <td>1054</td>
      <td>...</td>
      <td>0</td>
      <td>14632</td>
      <td>96703</td>
      <td>14632</td>
      <td>96703</td>
      <td>3348</td>
      <td>20636</td>
      <td>8388</td>
      <td>99.0622</td>
      <td>99.0622</td>
    </tr>
    <tr>
      <th>14</th>
      <td>PXD004720</td>
      <td>bread wheat</td>
      <td>grain-zadoks-87</td>
      <td>single</td>
      <td>FragPipe_Duncan_PXD004720_grain-zadoks-87_pept...</td>
      <td>FragPipe_Duncan_PXD004720_grain-zadoks-87_vali...</td>
      <td>FragPipe_Duncan_PXD004720_grain-zadoks-87_vali...</td>
      <td>111156</td>
      <td>110256</td>
      <td>900</td>
      <td>...</td>
      <td>0</td>
      <td>14790</td>
      <td>95466</td>
      <td>14790</td>
      <td>95466</td>
      <td>3393</td>
      <td>21207</td>
      <td>8348</td>
      <td>99.1903</td>
      <td>99.1903</td>
    </tr>
    <tr>
      <th>15</th>
      <td>PXD004720</td>
      <td>bread wheat</td>
      <td>leaf-flag-mature</td>
      <td>single</td>
      <td>FragPipe_Duncan_PXD004720_leaf-flag-mature_pep...</td>
      <td>FragPipe_Duncan_PXD004720_leaf-flag-mature_val...</td>
      <td>FragPipe_Duncan_PXD004720_leaf-flag-mature_val...</td>
      <td>144641</td>
      <td>143118</td>
      <td>1523</td>
      <td>...</td>
      <td>0</td>
      <td>19328</td>
      <td>123790</td>
      <td>19328</td>
      <td>123790</td>
      <td>4496</td>
      <td>25420</td>
      <td>11221</td>
      <td>98.9470</td>
      <td>98.9470</td>
    </tr>
    <tr>
      <th>16</th>
      <td>PXD004720</td>
      <td>bread wheat</td>
      <td>leaf-flag-senescing</td>
      <td>single</td>
      <td>FragPipe_Duncan_PXD004720_leaf-flag-senescing_...</td>
      <td>FragPipe_Duncan_PXD004720_leaf-flag-senescing_...</td>
      <td>FragPipe_Duncan_PXD004720_leaf-flag-senescing_...</td>
      <td>45680</td>
      <td>45054</td>
      <td>626</td>
      <td>...</td>
      <td>0</td>
      <td>4522</td>
      <td>40532</td>
      <td>4522</td>
      <td>40532</td>
      <td>1203</td>
      <td>10035</td>
      <td>2386</td>
      <td>98.6296</td>
      <td>98.6296</td>
    </tr>
    <tr>
      <th>17</th>
      <td>PXD004720</td>
      <td>bread wheat</td>
      <td>leaf-flag-young</td>
      <td>single</td>
      <td>FragPipe_Duncan_PXD004720_leaf-flag-young_pept...</td>
      <td>FragPipe_Duncan_PXD004720_leaf-flag-young_vali...</td>
      <td>FragPipe_Duncan_PXD004720_leaf-flag-young_vali...</td>
      <td>119922</td>
      <td>118601</td>
      <td>1321</td>
      <td>...</td>
      <td>0</td>
      <td>15118</td>
      <td>103483</td>
      <td>15118</td>
      <td>103483</td>
      <td>3479</td>
      <td>19981</td>
      <td>8922</td>
      <td>98.8985</td>
      <td>98.8985</td>
    </tr>
    <tr>
      <th>18</th>
      <td>PXD004720</td>
      <td>bread wheat</td>
      <td>lemma</td>
      <td>single</td>
      <td>FragPipe_Duncan_PXD004720_lemma_peptide_genome...</td>
      <td>FragPipe_Duncan_PXD004720_lemma_validated_pept...</td>
      <td>FragPipe_Duncan_PXD004720_lemma_validated_pept...</td>
      <td>146562</td>
      <td>145101</td>
      <td>1461</td>
      <td>...</td>
      <td>0</td>
      <td>21630</td>
      <td>123471</td>
      <td>21630</td>
      <td>123471</td>
      <td>4944</td>
      <td>24886</td>
      <td>12377</td>
      <td>99.0032</td>
      <td>99.0032</td>
    </tr>
    <tr>
      <th>19</th>
      <td>PXD004720</td>
      <td>bread wheat</td>
      <td>node</td>
      <td>single</td>
      <td>FragPipe_Duncan_PXD004720_node_peptide_genome_...</td>
      <td>FragPipe_Duncan_PXD004720_node_validated_pepti...</td>
      <td>FragPipe_Duncan_PXD004720_node_validated_pepti...</td>
      <td>98731</td>
      <td>97553</td>
      <td>1178</td>
      <td>...</td>
      <td>0</td>
      <td>12673</td>
      <td>84880</td>
      <td>12673</td>
      <td>84880</td>
      <td>3044</td>
      <td>18365</td>
      <td>7066</td>
      <td>98.8069</td>
      <td>98.8069</td>
    </tr>
    <tr>
      <th>20</th>
      <td>PXD004720</td>
      <td>bread wheat</td>
      <td>node_secretion</td>
      <td>single</td>
      <td>FragPipe_Duncan_PXD004720_node-secretion_pepti...</td>
      <td>FragPipe_Duncan_PXD004720_node-secretion_valid...</td>
      <td>FragPipe_Duncan_PXD004720_node-secretion_valid...</td>
      <td>168564</td>
      <td>166896</td>
      <td>1668</td>
      <td>...</td>
      <td>0</td>
      <td>22746</td>
      <td>144150</td>
      <td>22746</td>
      <td>144150</td>
      <td>5101</td>
      <td>29152</td>
      <td>13209</td>
      <td>99.0105</td>
      <td>99.0105</td>
    </tr>
    <tr>
      <th>21</th>
      <td>PXD004720</td>
      <td>bread wheat</td>
      <td>palea</td>
      <td>single</td>
      <td>FragPipe_Duncan_PXD004720_palea_peptide_genome...</td>
      <td>FragPipe_Duncan_PXD004720_palea_validated_pept...</td>
      <td>FragPipe_Duncan_PXD004720_palea_validated_pept...</td>
      <td>116285</td>
      <td>115047</td>
      <td>1238</td>
      <td>...</td>
      <td>0</td>
      <td>20580</td>
      <td>94467</td>
      <td>20580</td>
      <td>94467</td>
      <td>4570</td>
      <td>17192</td>
      <td>12047</td>
      <td>98.9354</td>
      <td>98.9354</td>
    </tr>
    <tr>
      <th>22</th>
      <td>PXD004720</td>
      <td>bread wheat</td>
      <td>pericarp</td>
      <td>single</td>
      <td>FragPipe_Duncan_PXD004720_pericarp_peptide_gen...</td>
      <td>FragPipe_Duncan_PXD004720_pericarp_validated_p...</td>
      <td>FragPipe_Duncan_PXD004720_pericarp_validated_p...</td>
      <td>130645</td>
      <td>129415</td>
      <td>1230</td>
      <td>...</td>
      <td>0</td>
      <td>20838</td>
      <td>108577</td>
      <td>20838</td>
      <td>108577</td>
      <td>4773</td>
      <td>23564</td>
      <td>11878</td>
      <td>99.0585</td>
      <td>99.0585</td>
    </tr>
    <tr>
      <th>23</th>
      <td>PXD004720</td>
      <td>bread wheat</td>
      <td>pollen</td>
      <td>single</td>
      <td>FragPipe_Duncan_PXD004720_pollen_peptide_genom...</td>
      <td>FragPipe_Duncan_PXD004720_pollen_validated_pep...</td>
      <td>FragPipe_Duncan_PXD004720_pollen_validated_pep...</td>
      <td>74334</td>
      <td>73586</td>
      <td>748</td>
      <td>...</td>
      <td>0</td>
      <td>9418</td>
      <td>64168</td>
      <td>9418</td>
      <td>64168</td>
      <td>2045</td>
      <td>11486</td>
      <td>5307</td>
      <td>98.9937</td>
      <td>98.9937</td>
    </tr>
    <tr>
      <th>24</th>
      <td>PXD004720</td>
      <td>bread wheat</td>
      <td>rachilla</td>
      <td>single</td>
      <td>FragPipe_Duncan_PXD004720_rachilla_peptide_gen...</td>
      <td>FragPipe_Duncan_PXD004720_rachilla_validated_p...</td>
      <td>FragPipe_Duncan_PXD004720_rachilla_validated_p...</td>
      <td>159219</td>
      <td>157648</td>
      <td>1571</td>
      <td>...</td>
      <td>0</td>
      <td>24647</td>
      <td>133001</td>
      <td>24647</td>
      <td>133001</td>
      <td>5544</td>
      <td>25666</td>
      <td>14382</td>
      <td>99.0133</td>
      <td>99.0133</td>
    </tr>
    <tr>
      <th>25</th>
      <td>PXD004720</td>
      <td>bread wheat</td>
      <td>radicle</td>
      <td>single</td>
      <td>FragPipe_Duncan_PXD004720_radicle_peptide_geno...</td>
      <td>FragPipe_Duncan_PXD004720_radicle_validated_pe...</td>
      <td>FragPipe_Duncan_PXD004720_radicle_validated_pe...</td>
      <td>193938</td>
      <td>192432</td>
      <td>1506</td>
      <td>...</td>
      <td>0</td>
      <td>32321</td>
      <td>160111</td>
      <td>32321</td>
      <td>160111</td>
      <td>7017</td>
      <td>32674</td>
      <td>18982</td>
      <td>99.2235</td>
      <td>99.2235</td>
    </tr>
    <tr>
      <th>26</th>
      <td>PXD004720</td>
      <td>bread wheat</td>
      <td>root-mature</td>
      <td>single</td>
      <td>FragPipe_Duncan_PXD004720_root-mature_peptide_...</td>
      <td>FragPipe_Duncan_PXD004720_root-mature_validate...</td>
      <td>FragPipe_Duncan_PXD004720_root-mature_validate...</td>
      <td>96387</td>
      <td>95329</td>
      <td>1058</td>
      <td>...</td>
      <td>0</td>
      <td>10790</td>
      <td>84539</td>
      <td>10790</td>
      <td>84539</td>
      <td>2558</td>
      <td>18650</td>
      <td>6077</td>
      <td>98.9023</td>
      <td>98.9023</td>
    </tr>
    <tr>
      <th>27</th>
      <td>PXD004720</td>
      <td>bread wheat</td>
      <td>root-secretion</td>
      <td>single</td>
      <td>FragPipe_Duncan_PXD004720_root-secretion_pepti...</td>
      <td>FragPipe_Duncan_PXD004720_root-secretion_valid...</td>
      <td>FragPipe_Duncan_PXD004720_root-secretion_valid...</td>
      <td>100085</td>
      <td>99127</td>
      <td>958</td>
      <td>...</td>
      <td>0</td>
      <td>15421</td>
      <td>83706</td>
      <td>15421</td>
      <td>83706</td>
      <td>3460</td>
      <td>17736</td>
      <td>8923</td>
      <td>99.0428</td>
      <td>99.0428</td>
    </tr>
    <tr>
      <th>28</th>
      <td>PXD004720</td>
      <td>bread wheat</td>
      <td>root-tip</td>
      <td>single</td>
      <td>FragPipe_Duncan_PXD004720_root-tip_peptide_gen...</td>
      <td>FragPipe_Duncan_PXD004720_root-tip_validated_p...</td>
      <td>FragPipe_Duncan_PXD004720_root-tip_validated_p...</td>
      <td>191262</td>
      <td>189975</td>
      <td>1287</td>
      <td>...</td>
      <td>0</td>
      <td>34893</td>
      <td>155082</td>
      <td>34893</td>
      <td>155082</td>
      <td>7493</td>
      <td>31196</td>
      <td>20981</td>
      <td>99.3271</td>
      <td>99.3271</td>
    </tr>
    <tr>
      <th>29</th>
      <td>PXD004720</td>
      <td>bread wheat</td>
      <td>root-vasculature</td>
      <td>single</td>
      <td>FragPipe_Duncan_PXD004720_root-vasculature_pep...</td>
      <td>FragPipe_Duncan_PXD004720_root-vasculature_val...</td>
      <td>FragPipe_Duncan_PXD004720_root-vasculature_val...</td>
      <td>111315</td>
      <td>110307</td>
      <td>1008</td>
      <td>...</td>
      <td>0</td>
      <td>14735</td>
      <td>95572</td>
      <td>14735</td>
      <td>95572</td>
      <td>3148</td>
      <td>17733</td>
      <td>8124</td>
      <td>99.0945</td>
      <td>99.0945</td>
    </tr>
    <tr>
      <th>30</th>
      <td>PXD004720</td>
      <td>bread wheat</td>
      <td>spike-immature</td>
      <td>single</td>
      <td>FragPipe_Duncan_PXD004720_spike-immature_pepti...</td>
      <td>FragPipe_Duncan_PXD004720_spike-immature_valid...</td>
      <td>FragPipe_Duncan_PXD004720_spike-immature_valid...</td>
      <td>188025</td>
      <td>186379</td>
      <td>1646</td>
      <td>...</td>
      <td>0</td>
      <td>33010</td>
      <td>153369</td>
      <td>33010</td>
      <td>153369</td>
      <td>7001</td>
      <td>29426</td>
      <td>19253</td>
      <td>99.1246</td>
      <td>99.1246</td>
    </tr>
    <tr>
      <th>31</th>
      <td>PXD004720</td>
      <td>bread wheat</td>
      <td>stem</td>
      <td>single</td>
      <td>FragPipe_Duncan_PXD004720_stem_peptide_genome_...</td>
      <td>FragPipe_Duncan_PXD004720_stem_validated_pepti...</td>
      <td>FragPipe_Duncan_PXD004720_stem_validated_pepti...</td>
      <td>60181</td>
      <td>59623</td>
      <td>558</td>
      <td>...</td>
      <td>0</td>
      <td>6875</td>
      <td>52748</td>
      <td>6875</td>
      <td>52748</td>
      <td>1665</td>
      <td>12579</td>
      <td>3946</td>
      <td>99.0728</td>
      <td>99.0728</td>
    </tr>
  </tbody>
</table>
<p>32 rows × 24 columns</p>
</div>


# Step 13 — Create a Non-Redundant Combined Peptide BED Track

This step generates a single non-redundant genome browser track containing all successfully projected wheat peptides across all tissues.

The purpose of this combined track is to improve the user experience in Apollo/JBrowse by allowing users to display all peptide evidence at once, without needing to manually activate each individual tissue track.

---

## Input files

### Validated peptide genome projection table from Step 11

```text
wheat_projection_translation_validated_sanity_checks_full_step11.csv
```

Only fully validated projected rows were retained.

### Filtering rule

```text
Projection_status == "projected"
```

---

## Redundancy removal strategy

This step does not simply concatenate all tissue-level BED files.

Instead, redundancy was removed at the peptide–protein–genome feature level.

A feature was considered redundant when it had the same:

```text
Chromosome
BED_start_0based
BED_end_0based_exclusive
Strand
Peptide
ProteinID
GeneID
BED_block_count
BED_block_sizes
BED_block_starts
```

This means that the same peptide/protein/genomic feature detected in multiple tissues is represented only once in the combined BED track.

---

## Tissue support retention

Although redundant features were collapsed, tissue-level evidence was retained in the combined output table.

The following support metrics were added:

| Metric | Description |
|---|---|
| Sources | Proteomics source datasets in which the feature was observed |
| Tissues | Wheat tissues in which the feature was observed |
| Tissue_count | Number of tissues supporting the feature |
| Observation_count | Number of tissue-level observations collapsed into the feature |
| Max_BED_score | Maximum BED score observed for the feature |

---

## Output files

### Combined non-redundant evidence table

```text
wheat_all_tissues_nonredundant_projected_peptides.csv
```

### Combined BED6 file

```text
FragPipe_allauthors_allsources_alltissues_nonredundant_validated_peptides.bed6
```

### Combined BED12 file

```text
FragPipe_allauthors_allsources_alltissues_nonredundant_validated_peptides.bed12
```

### Output directories

```text
python_outputs/tables/
python_outputs/bed/
```

---

## Summary metrics

A Step 13 summary table was generated.

### Metrics captured

| Metric | Description |
|---|---|
| Total_projected_rows_before_deduplication | Number of projected rows across all tissue-level files |
| Nonredundant_projected_rows | Number of unique peptide/protein/genome features retained |
| Redundant_rows_removed | Number of redundant tissue-level observations collapsed |
| Unique_peptides | Number of unique peptide sequences |
| Unique_proteins | Number of unique protein accessions |
| Unique_gene_models | Number of unique gene models |
| Unique_chromosomes | Number of chromosomes or genomic sequences represented |
| Multi_block_peptides | Number of features spanning more than one genomic block |

### Output summary file

```text
wheat_all_tissues_nonredundant_validated_bed_summary_step13.csv
```

The resulting combined BED files provide a user-friendly genome-wide peptide evidence track for Apollo/JBrowse visualisation.


```python
# ============================================================
# Step 13 — Create non-redundant combined validated BED tracks (takes 30 min)
# Translation-validated + sanity-check-passed projections only
# Uses SQLite for memory-safe aggregation
# ============================================================

import pandas as pd
import sqlite3
from pathlib import Path

# -----------------------------
# 1. Input / output paths
# -----------------------------
tables_dir = Path("python_outputs/tables")
bed_dir = Path("python_outputs/bed_validated")

bed_dir.mkdir(parents=True, exist_ok=True)
tables_dir.mkdir(parents=True, exist_ok=True)

# Step 11 output: translation-validated rows with sanity-check results
sanity_file = tables_dir / "wheat_projection_translation_validated_sanity_checks_full_step11.csv"

combined_table_out = tables_dir / "wheat_all_tissues_nonredundant_validated_peptides_step13.csv"
combined_bed6_out = bed_dir / "FragPipe_allauthors_allsources_alltissues_nonredundant_validated_peptides.bed6"
combined_bed12_out = bed_dir / "FragPipe_allauthors_allsources_alltissues_nonredundant_validated_peptides.bed12"
step13_summary_out = tables_dir / "wheat_all_tissues_nonredundant_validated_bed_summary_step13.csv"

sqlite_db = tables_dir / "wheat_validated_nonredundant_step13.sqlite"

chunk_size = 100_000

if not sanity_file.exists():
    raise FileNotFoundError(
        f"Step 11 sanity-check file not found:\n{sanity_file}\n\n"
        "Please run Step 11 first."
    )

# Clear previous outputs if rerunning
for path in [
    combined_table_out,
    combined_bed6_out,
    combined_bed12_out,
    step13_summary_out,
    sqlite_db
]:
    if path.exists():
        path.unlink()


# -----------------------------
# 2. Helper functions
# -----------------------------
def clean_bed_name(value):
    return (
        str(value)
        .replace(" ", "_")
        .replace(";", "|")
        .replace(",", "|")
        .replace("\t", "_")
        .replace("\n", "_")
    )


def make_bed_score(data):
    """
    Create BED score between 0 and 1000.
    """
    if "Probability" in data.columns:
        score = pd.to_numeric(data["Probability"], errors="coerce") * 1000
        score = score.fillna(1000).clip(0, 1000).round().astype(int)
    else:
        score = pd.Series([1000] * len(data), index=data.index)

    return score


# -----------------------------
# 3. Inspect Step 11 columns
# -----------------------------
header = pd.read_csv(sanity_file, nrows=0)

required_cols = [
    "Chromosome",
    "BED_start_0based",
    "BED_end_0based_exclusive",
    "Strand",
    "Peptide",
    "ProteinID",
    "BED_block_count",
    "BED_block_sizes",
    "BED_block_starts",
    "Source",
    "Tissue",
    "Sanity_check_status"
]

missing_required = [
    col for col in required_cols
    if col not in header.columns
]

if missing_required:
    raise KeyError(
        f"Missing required Step 11 column(s): {missing_required}"
    )

# Use best available gene/model label
if "GeneModel" in header.columns:
    gene_label_col = "GeneModel"
elif "GeneID" in header.columns:
    gene_label_col = "GeneID"
else:
    gene_label_col = None

# Use best available peptide display label
if "Peptide_intron_gapped" in header.columns:
    peptide_label_col = "Peptide_intron_gapped"
elif "Peptide_intron_gapped_compact" in header.columns:
    peptide_label_col = "Peptide_intron_gapped_compact"
else:
    peptide_label_col = "Peptide"

optional_cols = []

if "Probability" in header.columns:
    optional_cols.append("Probability")

if gene_label_col is not None:
    optional_cols.append(gene_label_col)

if peptide_label_col not in required_cols:
    optional_cols.append(peptide_label_col)

usecols = list(dict.fromkeys(required_cols + optional_cols))

print("Step 13 input file:")
print(sanity_file)
print(f"Using gene/model label column: {gene_label_col}")
print(f"Using peptide label column: {peptide_label_col}")


# -----------------------------
# 4. Load sanity-passed rows into SQLite
# -----------------------------
conn = sqlite3.connect(sqlite_db)

total_rows_read = 0
total_rows_passed = 0

print("\nLoading sanity-passed rows into SQLite...")

for chunk_i, chunk in enumerate(
    pd.read_csv(
        sanity_file,
        usecols=usecols,
        chunksize=chunk_size,
        low_memory=False
    ),
    start=1
):

    total_rows_read += len(chunk)

    # Keep only rows passing both validation layers:
    # Step 10 translation validation + Step 11 sanity checks
    chunk = chunk[
        chunk["Sanity_check_status"].astype(str) == "passed"
    ].copy()

    if chunk.empty:
        print(f"Chunk {chunk_i}: read {chunk_size:,} rows | no passed rows")
        continue

    total_rows_passed += len(chunk)

    # Standardise fields used downstream
    if gene_label_col is None:
        chunk["Gene_label"] = "NA"
    else:
        chunk["Gene_label"] = chunk[gene_label_col].astype(str)

    chunk["Peptide_label"] = chunk[peptide_label_col].astype(str)

    chunk["BED_score"] = make_bed_score(chunk)

    # Keep only columns needed for non-redundant aggregation
    insert_cols = [
        "Chromosome",
        "BED_start_0based",
        "BED_end_0based_exclusive",
        "Strand",
        "Peptide",
        "Peptide_label",
        "ProteinID",
        "Gene_label",
        "BED_block_count",
        "BED_block_sizes",
        "BED_block_starts",
        "Source",
        "Tissue",
        "BED_score"
    ]

    chunk = chunk[insert_cols].copy()

    # SQLite is happier with simple string/int/float fields
    for col in [
        "Chromosome",
        "Strand",
        "Peptide",
        "Peptide_label",
        "ProteinID",
        "Gene_label",
        "BED_block_sizes",
        "BED_block_starts",
        "Source",
        "Tissue"
    ]:
        chunk[col] = chunk[col].astype(str)

    numeric_cols = [
        "BED_start_0based",
        "BED_end_0based_exclusive",
        "BED_block_count",
        "BED_score"
    ]

    for col in numeric_cols:
        chunk[col] = pd.to_numeric(chunk[col], errors="coerce")

    chunk = chunk.dropna(
        subset=[
            "Chromosome",
            "BED_start_0based",
            "BED_end_0based_exclusive",
            "Strand",
            "BED_block_count",
            "BED_block_sizes",
            "BED_block_starts"
        ]
    ).copy()

    chunk.to_sql(
        "validated_projection_rows",
        conn,
        if_exists="append",
        index=False
    )

    print(
        f"Chunk {chunk_i}: cumulative rows read {total_rows_read:,} | "
        f"cumulative sanity-passed rows loaded {total_rows_passed:,}"
    )

if total_rows_passed == 0:
    conn.close()
    raise ValueError(
        "No sanity-passed rows were found in the Step 11 output."
    )


# -----------------------------
# 5. Create indexes to speed up grouping
# -----------------------------
print("\nCreating SQLite indexes...")

conn.execute("""
CREATE INDEX IF NOT EXISTS idx_validated_projection_dedup
ON validated_projection_rows (
    Chromosome,
    BED_start_0based,
    BED_end_0based_exclusive,
    Strand,
    Peptide,
    Peptide_label,
    ProteinID,
    Gene_label,
    BED_block_count,
    BED_block_sizes,
    BED_block_starts
)
""")

conn.commit()


# -----------------------------
# 6. Build non-redundant table in SQLite
# -----------------------------
print("\nBuilding non-redundant validated feature table...")

conn.execute("DROP TABLE IF EXISTS nonredundant_validated_peptides")

conn.execute("""
CREATE TABLE nonredundant_validated_peptides AS
SELECT
    Chromosome,
    CAST(BED_start_0based AS INTEGER) AS BED_start_0based,
    CAST(BED_end_0based_exclusive AS INTEGER) AS BED_end_0based_exclusive,
    Strand,
    Peptide,
    Peptide_label,
    ProteinID,
    Gene_label,
    CAST(BED_block_count AS INTEGER) AS BED_block_count,
    BED_block_sizes,
    BED_block_starts,
    GROUP_CONCAT(DISTINCT Source) AS Sources,
    GROUP_CONCAT(DISTINCT Tissue) AS Tissues,
    COUNT(DISTINCT Tissue) AS Tissue_count,
    COUNT(*) AS Observation_count,
    MAX(CAST(BED_score AS INTEGER)) AS Max_BED_score
FROM validated_projection_rows
GROUP BY
    Chromosome,
    BED_start_0based,
    BED_end_0based_exclusive,
    Strand,
    Peptide,
    Peptide_label,
    ProteinID,
    Gene_label,
    BED_block_count,
    BED_block_sizes,
    BED_block_starts
""")

conn.commit()


# -----------------------------
# 7. Export combined table and BED files in chunks
# -----------------------------
print("\nExporting combined non-redundant table and BED files...")

combined_header_written = False
bed6_header_written = False
bed12_header_written = False

total_nonredundant_rows = 0
unique_peptides = set()
unique_proteins = set()
unique_genes = set()
unique_chromosomes = set()
multi_block_count = 0
within_exon_count = 0
bed_labels_with_introns = 0

sql_query = """
SELECT
    Chromosome,
    BED_start_0based,
    BED_end_0based_exclusive,
    Strand,
    Peptide,
    Peptide_label,
    ProteinID,
    Gene_label,
    BED_block_count,
    BED_block_sizes,
    BED_block_starts,
    Sources,
    Tissues,
    Tissue_count,
    Observation_count,
    Max_BED_score
FROM nonredundant_validated_peptides
ORDER BY
    Chromosome,
    BED_start_0based,
    BED_end_0based_exclusive,
    Strand,
    Peptide,
    ProteinID
"""

for chunk_i, nonredundant in enumerate(
    pd.read_sql_query(sql_query, conn, chunksize=chunk_size),
    start=1
):

    nonredundant.insert(
        0,
        "Index",
        range(total_nonredundant_rows + 1, total_nonredundant_rows + len(nonredundant) + 1)
    )

    nonredundant["BED_name"] = (
        nonredundant["Peptide_label"].astype(str) + "|" +
        nonredundant["ProteinID"].astype(str) + "|" +
        nonredundant["Gene_label"].astype(str) + "|" +
        "validated=translation+sanity" + "|" +
        "tissues=" + nonredundant["Tissue_count"].astype(str)
    )

    nonredundant["BED_name"] = nonredundant["BED_name"].apply(clean_bed_name)

    nonredundant["BED_start_0based"] = nonredundant["BED_start_0based"].astype(int)
    nonredundant["BED_end_0based_exclusive"] = nonredundant["BED_end_0based_exclusive"].astype(int)
    nonredundant["BED_block_count"] = nonredundant["BED_block_count"].astype(int)
    nonredundant["Max_BED_score"] = nonredundant["Max_BED_score"].astype(int)

    # Save combined table incrementally
    nonredundant.to_csv(
        combined_table_out,
        index=False,
        mode="a",
        header=not combined_header_written
    )

    combined_header_written = True

    # BED6
    bed6 = nonredundant[[
        "Chromosome",
        "BED_start_0based",
        "BED_end_0based_exclusive",
        "BED_name",
        "Max_BED_score",
        "Strand"
    ]].copy()

    bed6.to_csv(
        combined_bed6_out,
        sep="\t",
        header=False,
        index=False,
        mode="a"
    )

    # BED12
    bed12 = pd.DataFrame({
        "chrom": nonredundant["Chromosome"],
        "chromStart": nonredundant["BED_start_0based"],
        "chromEnd": nonredundant["BED_end_0based_exclusive"],
        "name": nonredundant["BED_name"],
        "score": nonredundant["Max_BED_score"],
        "strand": nonredundant["Strand"],
        "thickStart": nonredundant["BED_start_0based"],
        "thickEnd": nonredundant["BED_end_0based_exclusive"],
        "itemRgb": "0",
        "blockCount": nonredundant["BED_block_count"],
        "blockSizes": nonredundant["BED_block_sizes"],
        "blockStarts": nonredundant["BED_block_starts"]
    })

    bed12.to_csv(
        combined_bed12_out,
        sep="\t",
        header=False,
        index=False,
        mode="a"
    )

    # Summary counters
    total_nonredundant_rows += len(nonredundant)

    unique_peptides.update(nonredundant["Peptide"].dropna().astype(str).unique())
    unique_proteins.update(nonredundant["ProteinID"].dropna().astype(str).unique())
    unique_genes.update(nonredundant["Gene_label"].dropna().astype(str).unique())
    unique_chromosomes.update(nonredundant["Chromosome"].dropna().astype(str).unique())

    multi_block_count += int((nonredundant["BED_block_count"] > 1).sum())
    
    within_exon_count += int((nonredundant["BED_block_count"] == 1).sum())

    bed_labels_with_introns += int(
        nonredundant["BED_name"]
        .astype(str)
        .str.contains("-", regex=False)
        .sum()
    )

    print(
        f"Export chunk {chunk_i}: cumulative non-redundant rows "
        f"{total_nonredundant_rows:,}"
    )


# -----------------------------
# 8. Summary table
# -----------------------------
step13_summary = pd.DataFrame([{
    "Validated_rows_before_deduplication": total_rows_passed,
    "Nonredundant_validated_rows": total_nonredundant_rows,
    "Redundant_validated_rows_removed": total_rows_passed - total_nonredundant_rows,
    "Unique_peptides": len(unique_peptides),
    "Unique_proteins": len(unique_proteins),
    "Unique_gene_models": len(unique_genes),
    "Unique_chromosomes": len(unique_chromosomes),
    "Multi_block_peptides": multi_block_count,
    "Within_exon_peptides": within_exon_count,
    "BED_labels_with_introns": bed_labels_with_introns,
    "Combined_table_file": combined_table_out.name,
    "BED6_file": combined_bed6_out.name,
    "BED12_file": combined_bed12_out.name,
    "SQLite_database": sqlite_db.name
}])

step13_summary.to_csv(step13_summary_out, index=False)

conn.close()

print("\n===== STEP 13 COMBINED VALIDATED BED SUMMARY =====")
print(f"Validated rows before deduplication: {total_rows_passed:,}")
print(f"Non-redundant validated rows: {total_nonredundant_rows:,}")
print(f"Redundant validated rows removed: {total_rows_passed - total_nonredundant_rows:,}")

print(f"\nCombined non-redundant table saved: {combined_table_out}")
print(f"Combined BED6 saved: {combined_bed6_out}")
print(f"Combined BED12 saved: {combined_bed12_out}")
print(f"Step 13 summary saved: {step13_summary_out}")
print(f"Temporary SQLite database saved: {sqlite_db}")

display(step13_summary)
```

    Step 13 input file:
    python_outputs\tables\wheat_projection_translation_validated_sanity_checks_full_step11.csv
    Using gene/model label column: GeneModel
    Using peptide label column: Peptide_intron_gapped
    
    Loading sanity-passed rows into SQLite...
    Chunk 1: cumulative rows read 100,000 | cumulative sanity-passed rows loaded 99,103
    Chunk 2: cumulative rows read 200,000 | cumulative sanity-passed rows loaded 198,145
    Chunk 3: cumulative rows read 300,000 | cumulative sanity-passed rows loaded 297,110
    Chunk 4: cumulative rows read 400,000 | cumulative sanity-passed rows loaded 396,182
    Chunk 5: cumulative rows read 500,000 | cumulative sanity-passed rows loaded 495,323
    Chunk 6: cumulative rows read 600,000 | cumulative sanity-passed rows loaded 594,161
    Chunk 7: cumulative rows read 700,000 | cumulative sanity-passed rows loaded 693,188
    Chunk 8: cumulative rows read 800,000 | cumulative sanity-passed rows loaded 792,324
    Chunk 9: cumulative rows read 900,000 | cumulative sanity-passed rows loaded 891,548
    Chunk 10: cumulative rows read 1,000,000 | cumulative sanity-passed rows loaded 990,802
    Chunk 11: cumulative rows read 1,100,000 | cumulative sanity-passed rows loaded 1,089,962
    Chunk 12: cumulative rows read 1,200,000 | cumulative sanity-passed rows loaded 1,189,050
    Chunk 13: cumulative rows read 1,300,000 | cumulative sanity-passed rows loaded 1,288,243
    Chunk 14: cumulative rows read 1,400,000 | cumulative sanity-passed rows loaded 1,387,148
    Chunk 15: cumulative rows read 1,500,000 | cumulative sanity-passed rows loaded 1,485,985
    Chunk 16: cumulative rows read 1,600,000 | cumulative sanity-passed rows loaded 1,584,882
    Chunk 17: cumulative rows read 1,700,000 | cumulative sanity-passed rows loaded 1,683,861
    Chunk 18: cumulative rows read 1,800,000 | cumulative sanity-passed rows loaded 1,782,920
    Chunk 19: cumulative rows read 1,900,000 | cumulative sanity-passed rows loaded 1,881,887
    Chunk 20: cumulative rows read 2,000,000 | cumulative sanity-passed rows loaded 1,980,697
    Chunk 21: cumulative rows read 2,100,000 | cumulative sanity-passed rows loaded 2,079,622
    Chunk 22: cumulative rows read 2,200,000 | cumulative sanity-passed rows loaded 2,178,640
    Chunk 23: cumulative rows read 2,300,000 | cumulative sanity-passed rows loaded 2,277,678
    Chunk 24: cumulative rows read 2,400,000 | cumulative sanity-passed rows loaded 2,376,733
    Chunk 25: cumulative rows read 2,500,000 | cumulative sanity-passed rows loaded 2,475,723
    Chunk 26: cumulative rows read 2,600,000 | cumulative sanity-passed rows loaded 2,574,929
    Chunk 27: cumulative rows read 2,700,000 | cumulative sanity-passed rows loaded 2,674,151
    Chunk 28: cumulative rows read 2,800,000 | cumulative sanity-passed rows loaded 2,773,059
    Chunk 29: cumulative rows read 2,900,000 | cumulative sanity-passed rows loaded 2,872,135
    Chunk 30: cumulative rows read 3,000,000 | cumulative sanity-passed rows loaded 2,971,428
    Chunk 31: cumulative rows read 3,100,000 | cumulative sanity-passed rows loaded 3,070,748
    Chunk 32: cumulative rows read 3,200,000 | cumulative sanity-passed rows loaded 3,169,833
    Chunk 33: cumulative rows read 3,300,000 | cumulative sanity-passed rows loaded 3,268,969
    Chunk 34: cumulative rows read 3,400,000 | cumulative sanity-passed rows loaded 3,368,066
    Chunk 35: cumulative rows read 3,500,000 | cumulative sanity-passed rows loaded 3,467,214
    Chunk 36: cumulative rows read 3,600,000 | cumulative sanity-passed rows loaded 3,566,383
    Chunk 37: cumulative rows read 3,700,000 | cumulative sanity-passed rows loaded 3,665,497
    Chunk 38: cumulative rows read 3,800,000 | cumulative sanity-passed rows loaded 3,764,679
    Chunk 39: cumulative rows read 3,900,000 | cumulative sanity-passed rows loaded 3,863,736
    Chunk 40: cumulative rows read 4,000,000 | cumulative sanity-passed rows loaded 3,962,693
    Chunk 41: cumulative rows read 4,100,000 | cumulative sanity-passed rows loaded 4,061,690
    Chunk 42: cumulative rows read 4,200,000 | cumulative sanity-passed rows loaded 4,160,797
    Chunk 43: cumulative rows read 4,300,000 | cumulative sanity-passed rows loaded 4,259,839
    Chunk 44: cumulative rows read 4,400,000 | cumulative sanity-passed rows loaded 4,358,967
    Chunk 45: cumulative rows read 4,500,000 | cumulative sanity-passed rows loaded 4,457,786
    Chunk 46: cumulative rows read 4,600,000 | cumulative sanity-passed rows loaded 4,556,649
    Chunk 47: cumulative rows read 4,700,000 | cumulative sanity-passed rows loaded 4,655,597
    Chunk 48: cumulative rows read 4,800,000 | cumulative sanity-passed rows loaded 4,754,683
    Chunk 49: cumulative rows read 4,900,000 | cumulative sanity-passed rows loaded 4,853,691
    Chunk 50: cumulative rows read 5,000,000 | cumulative sanity-passed rows loaded 4,952,890
    Chunk 51: cumulative rows read 5,100,000 | cumulative sanity-passed rows loaded 5,052,029
    Chunk 52: cumulative rows read 5,200,000 | cumulative sanity-passed rows loaded 5,151,156
    Chunk 53: cumulative rows read 5,300,000 | cumulative sanity-passed rows loaded 5,250,257
    Chunk 54: cumulative rows read 5,400,000 | cumulative sanity-passed rows loaded 5,349,355
    Chunk 55: cumulative rows read 5,500,000 | cumulative sanity-passed rows loaded 5,448,421
    Chunk 56: cumulative rows read 5,600,000 | cumulative sanity-passed rows loaded 5,547,603
    Chunk 57: cumulative rows read 5,700,000 | cumulative sanity-passed rows loaded 5,646,659
    Chunk 58: cumulative rows read 5,800,000 | cumulative sanity-passed rows loaded 5,745,807
    Chunk 59: cumulative rows read 5,900,000 | cumulative sanity-passed rows loaded 5,844,747
    Chunk 60: cumulative rows read 6,000,000 | cumulative sanity-passed rows loaded 5,943,707
    Chunk 61: cumulative rows read 6,100,000 | cumulative sanity-passed rows loaded 6,042,704
    Chunk 62: cumulative rows read 6,200,000 | cumulative sanity-passed rows loaded 6,141,780
    Chunk 63: cumulative rows read 6,300,000 | cumulative sanity-passed rows loaded 6,240,726
    Chunk 64: cumulative rows read 6,400,000 | cumulative sanity-passed rows loaded 6,339,436
    Chunk 65: cumulative rows read 6,500,000 | cumulative sanity-passed rows loaded 6,438,416
    Chunk 66: cumulative rows read 6,600,000 | cumulative sanity-passed rows loaded 6,537,413
    Chunk 67: cumulative rows read 6,700,000 | cumulative sanity-passed rows loaded 6,636,451
    Chunk 68: cumulative rows read 6,800,000 | cumulative sanity-passed rows loaded 6,735,610
    Chunk 69: cumulative rows read 6,900,000 | cumulative sanity-passed rows loaded 6,834,809
    Chunk 70: cumulative rows read 7,000,000 | cumulative sanity-passed rows loaded 6,933,962
    Chunk 71: cumulative rows read 7,100,000 | cumulative sanity-passed rows loaded 7,033,103
    Chunk 72: cumulative rows read 7,200,000 | cumulative sanity-passed rows loaded 7,132,162
    Chunk 73: cumulative rows read 7,300,000 | cumulative sanity-passed rows loaded 7,231,302
    Chunk 74: cumulative rows read 7,400,000 | cumulative sanity-passed rows loaded 7,330,455
    Chunk 75: cumulative rows read 7,500,000 | cumulative sanity-passed rows loaded 7,429,442
    Chunk 76: cumulative rows read 7,600,000 | cumulative sanity-passed rows loaded 7,528,490
    Chunk 77: cumulative rows read 7,700,000 | cumulative sanity-passed rows loaded 7,627,591
    Chunk 78: cumulative rows read 7,800,000 | cumulative sanity-passed rows loaded 7,726,410
    Chunk 79: cumulative rows read 7,900,000 | cumulative sanity-passed rows loaded 7,825,347
    Chunk 80: cumulative rows read 8,000,000 | cumulative sanity-passed rows loaded 7,924,564
    Chunk 81: cumulative rows read 8,100,000 | cumulative sanity-passed rows loaded 8,023,675
    Chunk 82: cumulative rows read 8,200,000 | cumulative sanity-passed rows loaded 8,122,670
    Chunk 83: cumulative rows read 8,214,230 | cumulative sanity-passed rows loaded 8,136,687
    
    Creating SQLite indexes...
    
    Building non-redundant validated feature table...
    
    Exporting combined non-redundant table and BED files...
    Export chunk 1: cumulative non-redundant rows 100,000
    Export chunk 2: cumulative non-redundant rows 200,000
    Export chunk 3: cumulative non-redundant rows 300,000
    Export chunk 4: cumulative non-redundant rows 400,000
    Export chunk 5: cumulative non-redundant rows 500,000
    Export chunk 6: cumulative non-redundant rows 600,000
    Export chunk 7: cumulative non-redundant rows 700,000
    Export chunk 8: cumulative non-redundant rows 800,000
    Export chunk 9: cumulative non-redundant rows 900,000
    Export chunk 10: cumulative non-redundant rows 1,000,000
    Export chunk 11: cumulative non-redundant rows 1,100,000
    Export chunk 12: cumulative non-redundant rows 1,200,000
    Export chunk 13: cumulative non-redundant rows 1,300,000
    Export chunk 14: cumulative non-redundant rows 1,400,000
    Export chunk 15: cumulative non-redundant rows 1,500,000
    Export chunk 16: cumulative non-redundant rows 1,600,000
    Export chunk 17: cumulative non-redundant rows 1,700,000
    Export chunk 18: cumulative non-redundant rows 1,800,000
    Export chunk 19: cumulative non-redundant rows 1,900,000
    Export chunk 20: cumulative non-redundant rows 2,000,000
    Export chunk 21: cumulative non-redundant rows 2,100,000
    Export chunk 22: cumulative non-redundant rows 2,200,000
    Export chunk 23: cumulative non-redundant rows 2,300,000
    Export chunk 24: cumulative non-redundant rows 2,400,000
    Export chunk 25: cumulative non-redundant rows 2,500,000
    Export chunk 26: cumulative non-redundant rows 2,600,000
    Export chunk 27: cumulative non-redundant rows 2,700,000
    Export chunk 28: cumulative non-redundant rows 2,800,000
    Export chunk 29: cumulative non-redundant rows 2,900,000
    Export chunk 30: cumulative non-redundant rows 3,000,000
    Export chunk 31: cumulative non-redundant rows 3,100,000
    Export chunk 32: cumulative non-redundant rows 3,138,903
    
    ===== STEP 13 COMBINED VALIDATED BED SUMMARY =====
    Validated rows before deduplication: 8,136,687
    Non-redundant validated rows: 3,138,903
    Redundant validated rows removed: 4,997,784
    
    Combined non-redundant table saved: python_outputs\tables\wheat_all_tissues_nonredundant_validated_peptides_step13.csv
    Combined BED6 saved: python_outputs\bed_validated\FragPipe_allauthors_allsources_alltissues_nonredundant_validated_peptides.bed6
    Combined BED12 saved: python_outputs\bed_validated\FragPipe_allauthors_allsources_alltissues_nonredundant_validated_peptides.bed12
    Step 13 summary saved: python_outputs\tables\wheat_all_tissues_nonredundant_validated_bed_summary_step13.csv
