## Supplementary material for "Community Resource: A Genome-Based Extension of Large-Scale Wheat Proteogenomics": Vincent_wheat-proteogenomics_technical-note_2026-06-16_Suppl-File-S2.txt

### Vincent-Appels 2026 -- Supplementary File S2

####################################################
################# FragPipe Worflow #################
####################################################
Author: Delphine Vincent (March 2026)

-----------------------------------------------------
----------------- Software version ------------------
-----------------------------------------------------
# Version info:
# FragPipe version 24.0
# DIA-Umpire version 2.3.3
# diaTracer version 2.2.1
# MSFragger version 4.4.1
# Crystal-C version 1.5.10
# MSBooster version 1.4.14
# Percolator version 3.7.1
# PTMProphet version 6.3.2
# Metaproteomics version 1.0.1
# Philosopher version 5.1.3-RC9
# PTM-Shepherd version 3.0.11
# IonQuant version 1.11.20
# TMT-Integrator version 6.1.3
# FragPipe-SpecLib version 0.1.58
# DIA-NN version 1.8.2 Beta 8
# Skyline version N/A
# Pandas version 2.3.3
# Numpy version 1.26.4

-----------------------------------------------------
----------------- Workflow params ------------------
-----------------------------------------------------
# Please edit the following path to point to the correct location.
# In Windows, please replace single '\' with '\\'
database.db-path=C\:\\Users\\Delphine_2026\\DB\\iwgsc_refseqv2.1_annotation_200916_HC_LC_pep_cRAP_with_DECOY.fasta

crystalc.run-crystalc=false
database.decoy-tag=decoy_
diann.channel-normalization-strategy=0
diann.cmd-opts=
diann.gene-level-report=false
diann.generate-msstats=true
diann.heavy=
diann.library=
diann.light=
diann.mbr=false
diann.medium=
diann.min-site-prob=0.75
diann.mod-tag=
diann.modified-peptide-level-report=true
diann.peptide-level-report=true
diann.protein-level-report=false
diann.q-value=0.01
diann.quantification-strategy=3
diann.quantification-strategy-2=2
diann.redo-protein-inference=false
diann.run-dia-nn=false
diann.run-dia-plex=false
diann.run-specific-protein-q-value=false
diann.site-level-report=false
diann.unrelated-runs=false
diatracer.corr-threshold=0.3
diatracer.delta-apex-im=0.01
diatracer.delta-apex-rt=3
diatracer.mass-defect-filter=true
diatracer.mass-defect-offset=0.1
diatracer.rf-max=500
diatracer.run-diatracer=false
diatracer.write-intermediate-files=false
diaumpire.AdjustFragIntensity=false
diaumpire.BoostComplementaryIon=false
diaumpire.CorrThreshold=
diaumpire.DeltaApex=
diaumpire.ExportPrecursorPeak=false
diaumpire.Q1=false
diaumpire.Q2=false
diaumpire.Q3=false
diaumpire.RFmax=
diaumpire.RPmax=
diaumpire.RTOverlap=
diaumpire.SE.EstimateBG=false
diaumpire.SE.IsoPattern=
diaumpire.SE.MS1PPM=
diaumpire.SE.MS2PPM=
diaumpire.SE.MS2SN=
diaumpire.SE.MassDefectFilter=false
diaumpire.SE.MassDefectOffset=
diaumpire.SE.NoMissedScan=
diaumpire.SE.SN=
diaumpire.run-diaumpire=false
fpop.coadaptr.fpop.fpop_masses=
fpop.coadaptr.fpop.run-fpop-coadaptr=false
fpop.fragpipe.fpop.fpop-tmt=false
fpop.fragpipe.fpop.label_control=
fpop.fragpipe.fpop.label_fpop=
fpop.fragpipe.fpop.region_size=1
fpop.fragpipe.fpop.run-fpop=false
fpop.fragpipe.fpop.subtract-control=false
fragpipe-config.bin-diann=C\:\\FragPipe\\FragPipe-24.0\\tools\\diann\\1.8.2_beta_8\\windows\\DiaNN.exe
fragpipe-config.bin-python=C\:\\FragPipe\\FragPipe-24.0\\python\\python.exe
fragpipe-config.tools-folder=C\:\\FragPipe\\FragPipe-24.0\\tools
freequant.mz-tol=10
freequant.rt-tol=0.4
freequant.run-freequant=true
ionquant.excludemods=
ionquant.formula=
ionquant.heavy=
ionquant.imtol=0.05
ionquant.intensitymode=2
ionquant.ionfdr=0.01
ionquant.light=
ionquant.locprob=0.75
ionquant.maxlfq=1
ionquant.mbr=1
ionquant.mbrimtol=0.05
ionquant.mbrmincorr=0
ionquant.mbrrttol=1
ionquant.mbrtoprun=10
ionquant.medium=
ionquant.minfreq=0
ionquant.minions=1
ionquant.minisotopes=1
ionquant.minscans=3
ionquant.mztol=10
ionquant.normalization=1
ionquant.peptidefdr=1
ionquant.proteinfdr=1
ionquant.requantify=1
ionquant.rttol=0.4
ionquant.run-ionquant=false
ionquant.tp=0
ionquant.uniqueness=0
ionquant.use-labeling=false
ionquant.use-lfq=false
ionquant.writeindex=0
mbg.allow_chimeric=false
mbg.expand_db=2
mbg.fdr=0.010
mbg.max_glycan_q=0.01
mbg.max_skips=0
mbg.min_glycans=2
mbg.min_psms=5
mbg.residues_to_add=
mbg.run-mbg=false
metaproteomics.cmd-line-opts=
metaproteomics.delta-hyperscore=0.0
metaproteomics.host-name=Homo sapiens
metaproteomics.iterations=3
metaproteomics.min-pept-cnt-per-prot=1
metaproteomics.min-uniq-pept-cnt=3
metaproteomics.min-uniq-pept-cnt-per-prot=1
metaproteomics.qvalue=0.01
metaproteomics.run-metaproteomics=false
msbooster.find-best-im-model=false
msbooster.find-best-rt-model=false
msbooster.find-best-spectra-model=false
msbooster.fragmentation-type=0
msbooster.im-model=DIA-NN
msbooster.koina-url=
msbooster.predict-im=true
msbooster.predict-rt=true
msbooster.predict-spectra=true
msbooster.rt-model=DIA-NN
msbooster.run-msbooster=false
msbooster.spectra-model=DIA-NN
msbooster.spectral-library-path=
msfragger.Y_type_masses=
msfragger.activation_types=all
msfragger.allowed_missed_cleavage_1=10
msfragger.allowed_missed_cleavage_2=2
msfragger.analyzer_types=all
msfragger.calibrate_mass=0
msfragger.check_spectral_files=false
msfragger.clip_nTerm_M=true
msfragger.deisotope=1
msfragger.delta_mass_exclude_ranges=(-1.5,3.5)
msfragger.deneutralloss=1
msfragger.diagnostic_fragments=
msfragger.diagnostic_intensity_filter=0
msfragger.digest-only=false
msfragger.digest_max_length=100
msfragger.digest_min_length=6
msfragger.fragment_ion_series=b,y
msfragger.fragment_mass_tolerance=0.05
msfragger.fragment_mass_units=0
msfragger.group_variable=0
msfragger.intensity_transform=0
msfragger.ion_series_definitions=
msfragger.isotope_error=0/1/2
msfragger.labile_fragment_ion_series=b,y
msfragger.labile_search_mode=off
msfragger.localize_delta_mass=false
msfragger.mass_diff_to_variable_mod=0
msfragger.mass_offsets=0
msfragger.mass_offsets_detailed=
msfragger.max_fragment_charge=10
msfragger.max_variable_mods_combinations=5
msfragger.max_variable_mods_per_peptide=3
msfragger.min_fragments_modelling=2
msfragger.min_matched_fragments=4
msfragger.min_sequence_matches=2
msfragger.minimum_peaks=15
msfragger.minimum_ratio=0.01
msfragger.misc.fragger.clear-mz-hi=0
msfragger.misc.fragger.clear-mz-lo=0
msfragger.misc.fragger.digest-mass-hi=10000
msfragger.misc.fragger.digest-mass-lo=300
msfragger.misc.fragger.enzyme-dropdown-1=trypsin
msfragger.misc.fragger.enzyme-dropdown-2=trypsin
msfragger.misc.fragger.precursor-charge-hi=4
msfragger.misc.fragger.precursor-charge-lo=1
msfragger.misc.fragger.remove-precursor-range-hi=1.5
msfragger.misc.fragger.remove-precursor-range-lo=-1.5
msfragger.misc.slice-db=2
msfragger.num_enzyme_termini=2
msfragger.output_format=pepXML
msfragger.output_max_expect=50
msfragger.output_report_topN=1
msfragger.output_report_topN_dda_plus=5
msfragger.output_report_topN_dia1=5
msfragger.override_charge=false
msfragger.precursor_mass_lower=-10
msfragger.precursor_mass_mode=selected
msfragger.precursor_mass_units=1
msfragger.precursor_mass_upper=10
msfragger.precursor_true_tolerance=20
msfragger.precursor_true_units=1
msfragger.remainder_fragment_masses=
msfragger.remove_precursor_peak=1
msfragger.report_alternative_proteins=false
msfragger.require_precursor=false
msfragger.restrict_deltamass_to=all
msfragger.reuse_dia_fragment_peaks=false
msfragger.run-msfragger=true
msfragger.search_enzyme_cut_1=KR
msfragger.search_enzyme_cut_2=
msfragger.search_enzyme_name_1=Trypsin/P
msfragger.search_enzyme_name_2=null
msfragger.search_enzyme_nocut_1=
msfragger.search_enzyme_nocut_2=
msfragger.search_enzyme_sense_1=C
msfragger.search_enzyme_sense_2=C
msfragger.table.fix-mods=0.0,C-Term Peptide,true,-1; 0.0,N-Term Peptide,true,-1; 0.0,C-Term Protein,true,-1; 0.0,N-Term Protein,true,-1; 0.0,G (glycine),true,-1; 0.0,A (alanine),true,-1; 0.0,S (serine),true,-1; 0.0,P (proline),true,-1; 0.0,V (valine),true,-1; 0.0,T (threonine),true,-1; 57.02146,C (cysteine),true,-1; 0.0,L (leucine),true,-1; 0.0,I (isoleucine),true,-1; 0.0,N (asparagine),true,-1; 0.0,D (aspartic acid),true,-1; 0.0,Q (glutamine),true,-1; 0.0,K (lysine),true,-1; 0.0,E (glutamic acid),true,-1; 0.0,M (methionine),true,-1; 0.0,H (histidine),true,-1; 0.0,F (phenylalanine),true,-1; 0.0,R (arginine),true,-1; 0.0,Y (tyrosine),true,-1; 0.0,W (tryptophan),true,-1; 0.0,B ,true,-1; 0.0,J,true,-1; 0.0,O,true,-1; 0.0,U,true,-1; 0.0,X,true,-1; 0.0,Z,true,-1
msfragger.table.var-mods=15.9949,M,true,3; 42.0106,[^,true,1; 79.96633,STY,false,3; -17.0265,nQnC,false,1; -18.0106,nE,false,1; 4.025107,K,false,2; 6.020129,KR,false,2; 8.014199,K,false,2; 10.008269,R,false,2; 0.0,site_10,false,1; 0.0,site_11,false,1; 0.0,site_12,false,1; 0.0,site_13,false,1; 0.0,site_14,false,1; 0.0,site_15,false,1; 0.0,site_16,false,1
msfragger.track_zero_topN=0
msfragger.use_all_mods_in_first_search=false
msfragger.use_detailed_offsets=false
msfragger.use_topN_peaks=150
msfragger.write_calibrated_mzml=false
msfragger.zero_bin_accept_expect=0
msfragger.zero_bin_mult_expect=1
opair.activation1=HCD
opair.activation2=HCD
opair.allowed_sites=
opair.filterOxonium=true
opair.glyco_db=
opair.max_glycans=2
opair.max_isotope_error=2
opair.min_isotope_error=0
opair.ms1_tol=20
opair.ms2_tol=20
opair.oxonium_filtering_file=
opair.oxonium_minimum_intensity=0.05
opair.reverse_scan_order=false
opair.run-opair=false
opair.single_scan_type=false
peptide-prophet.cmd-opts=
peptide-prophet.combine-pepxml=false
peptide-prophet.run-peptide-prophet=true
percolator.cmd-opts=
percolator.keep-tsv-files=false
percolator.min-prob=0.5
percolator.run-percolator=false
phi-report.dont-use-prot-proph-file=false
phi-report.filter=--sequential --prot 0.01
phi-report.pep-level-summary=true
phi-report.print-decoys=false
phi-report.prot-level-summary=true
phi-report.remove-contaminants=true
phi-report.run-report=true
protein-prophet.cmd-opts=
protein-prophet.run-protein-prophet=true
ptmprophet.cmdline=NOSTACK KEEPOLD STATIC FRAGPPMTOL\=10 EM\=1 NIONS\=b STY\:79.966331,M\:15.9949 MINPROB\=0.5
ptmprophet.override-defaults=false
ptmprophet.run-ptmprophet=false
ptmshepherd.adv_params=false
ptmshepherd.annotate_assigned_mods=false
ptmshepherd.annotation-common=false
ptmshepherd.annotation-custom=false
ptmshepherd.annotation-glyco=false
ptmshepherd.annotation-unimod=true
ptmshepherd.annotation_file=
ptmshepherd.annotation_tol=0.01
ptmshepherd.cap_y_ions=
ptmshepherd.decoy_type=1
ptmshepherd.diag_ions=
ptmshepherd.diagmine_diagMinFoldChange=3.0
ptmshepherd.diagmine_diagMinSpecDiff=25
ptmshepherd.diagmine_fragMinFoldChange=3.0
ptmshepherd.diagmine_fragMinPropensity=12.5
ptmshepherd.diagmine_fragMinSpecDiff=15
ptmshepherd.diagmine_minIonsPerSpec=2
ptmshepherd.diagmine_minPeps=25
ptmshepherd.diagmine_pepMinFoldChange=3.0
ptmshepherd.diagmine_pepMinSpecDiff=25
ptmshepherd.glyco_fdr=0.01
ptmshepherd.glyco_isotope_max=3
ptmshepherd.glyco_isotope_min=-1
ptmshepherd.glyco_lda=false
ptmshepherd.glyco_lda_features=
ptmshepherd.glyco_ppm_tol=50
ptmshepherd.glycodatabase=
ptmshepherd.histo_smoothbins=2
ptmshepherd.iontype_a=false
ptmshepherd.iontype_b=true
ptmshepherd.iontype_c=false
ptmshepherd.iontype_x=false
ptmshepherd.iontype_y=true
ptmshepherd.iontype_z=false
ptmshepherd.localization_allowed_res=all
ptmshepherd.n_glyco=false
ptmshepherd.normalization-psms=true
ptmshepherd.normalization-scans=false
ptmshepherd.output_extended=false
ptmshepherd.peakpicking_mass_units=0
ptmshepherd.peakpicking_minPsm=10
ptmshepherd.peakpicking_promRatio=0.3
ptmshepherd.peakpicking_width=0.002
ptmshepherd.precursor_mass_units=0
ptmshepherd.precursor_tol=0.01
ptmshepherd.print_decoys=false
ptmshepherd.print_full_glyco_params=false
ptmshepherd.prob_mass=0.5
ptmshepherd.remainder_masses=
ptmshepherd.remove_glycan_delta_mass=false
ptmshepherd.run-shepherd=false
ptmshepherd.run_diagextract_mode=false
ptmshepherd.run_diagmine_mode=false
ptmshepherd.run_glyco_mode=false
ptmshepherd.shuffle_decoy_intensities=false
ptmshepherd.spectra_condPeaks=150
ptmshepherd.spectra_condRatio=0.0001
ptmshepherd.spectra_maxPrecursorCharge=4
ptmshepherd.spectra_maxfragcharge=2
ptmshepherd.spectra_ppmtol=20
ptmshepherd.use_glycan_fragment_probs=false
ptmshepherd.use_msfragger_localization=false
ptmshepherd.varmod_masses=
quantitation.run-label-free-quant=false
run-psm-validation=true
run-validation-tab=true
saintexpress.cmd-opts=
saintexpress.max-replicates=10
saintexpress.run-saint-express=false
saintexpress.virtual-controls=100
skyline.fdr=1
skyline.generate-skyline-quant-report=false
skyline.min-site-prob=0.75
skyline.mod-tag=
skyline.run-skyline=false
skyline.skyline=true
skyline.skyline-custom=false
skyline.skyline-custom-path=
skyline.skyline-daily=false
skyline.skyline-fragment-tolerance=10
skyline.skyline-library-product-ions=12
skyline.skyline-mods-mode=Default
skyline.skyline-precursor-tolerance=10
skyline.skyline-rt-tolerance=2
skyline.use-existing-skyline-document=false
speclibgen.convert-pepxml=true
speclibgen.convert-psm=false
speclibgen.easypqp.extras.max_delta_ppm=15
speclibgen.easypqp.extras.max_delta_unimod=0.02
speclibgen.easypqp.extras.max_glycan_qval=1
speclibgen.easypqp.extras.rt_lowess_fraction=0.01
speclibgen.easypqp.fragment.a=false
speclibgen.easypqp.fragment.b=true
speclibgen.easypqp.fragment.c=false
speclibgen.easypqp.fragment.x=false
speclibgen.easypqp.fragment.y=true
speclibgen.easypqp.fragment.z=false
speclibgen.easypqp.im-cal=Automatic selection of a run as reference IM
speclibgen.easypqp.labile_mode=Regular (not glyco)
speclibgen.easypqp.neutral_loss=false
speclibgen.easypqp.rt-cal=noiRT
speclibgen.easypqp.select-file.text=
speclibgen.easypqp.select-im-file.text=
speclibgen.keep-intermediate-files=false
speclibgen.run-speclibgen=false
tab-run.delete_temp_files=false
tab-run.export_matched_fragments=false
tab-run.sub_mzml_prob_threshold=0.5
tab-run.write_sub_mzml=false
tmtintegrator.add_Ref=-1
tmtintegrator.aggregation_method=0
tmtintegrator.allow_overlabel=true
tmtintegrator.allow_unlabeled=true
tmtintegrator.best_psm=true
tmtintegrator.channel_num=TMT-0
tmtintegrator.extraction_tool=IonQuant
tmtintegrator.glyco_qval=-1
tmtintegrator.groupby=0
tmtintegrator.log2transformed=true
tmtintegrator.max_pep_prob_thres=0
tmtintegrator.min_ntt=0
tmtintegrator.min_pep_prob=0.9
tmtintegrator.min_percent=0.05
tmtintegrator.min_purity=0.5
tmtintegrator.min_resolution=45000
tmtintegrator.min_site_prob=-1
tmtintegrator.min_snr=1000
tmtintegrator.mod_tag=none
tmtintegrator.ms1_int=true
tmtintegrator.outlier_removal=true
tmtintegrator.philosopher-msstats=false
tmtintegrator.print_RefInt=false
tmtintegrator.prot_exclude=none
tmtintegrator.prot_norm=0
tmtintegrator.psm_norm=false
tmtintegrator.quant_level=2
tmtintegrator.ref_d_tag=Pool
tmtintegrator.ref_tag=Bridge
tmtintegrator.run-tmtintegrator=false
tmtintegrator.tolerance=20
tmtintegrator.unique_gene=0
tmtintegrator.unique_pep=false
tmtintegrator.use_glycan_composition=false
transfer-learning.credential=
transfer-learning.instrument=Astral
transfer-learning.keep-decoys=false
transfer-learning.max-charge=3
transfer-learning.min-charge=2
transfer-learning.model-path=
transfer-learning.nce=30
transfer-learning.output-format=speclib
transfer-learning.peptides-to-predict=Whole FASTA file
transfer-learning.perform-transfer-learning=true
transfer-learning.predict-im=true
transfer-learning.predict-ms2=true
transfer-learning.predict-rt=true
transfer-learning.predict-spectral-library=true
transfer-learning.run-transfer-learning=false
transfer-learning.spectral-library-path=
workdir=C\:\\Users\\Delphine_2026\\FragPipe_results5
workflow.input.data-type.im-ms=false
workflow.input.data-type.regular-ms=true
workflow.misc.save-sdrf=true
workflow.misc.sdrf-type=Default
workflow.ram=32
workflow.threads=4

-----------------------------------------------------
----------------- MSFragger params ------------------
-----------------------------------------------------
database_name = C:\Users\Delphine_2026\DB\iwgsc_refseqv2.1_annotation_200916_HC_LC_pep_cRAP_with_DECOY.fasta			# Path to the protein database file in FASTA format.
num_threads = 4			# Number of CPU threads to use.

precursor_mass_lower = -10			# Lower bound of the precursor mass window.
precursor_mass_upper = 10			# Upper bound of the precursor mass window.
precursor_mass_units = 1			# Precursor mass tolerance units (0 for Da, 1 for ppm).
data_type = 0			# Data type (0 for DDA, 1 for DIA, 2 for gas-phase fractionation DIA, 3 for DDA+).
precursor_true_tolerance = 20			# True precursor mass tolerance (window is +/- this value).
precursor_true_units = 1			# True precursor mass tolerance units (0 for Da, 1 for ppm).
fragment_mass_tolerance = 0.05			# Fragment mass tolerance (window is +/- this value).
fragment_mass_units = 0			# Fragment mass tolerance units (0 for Da, 1 for ppm).
calibrate_mass = 0			# Perform mass calibration (0 for OFF, 1 for ON, 2 for ON and find optimal parameters, 4 for ON and find the optimal fragment mass tolerance).
use_all_mods_in_first_search = 0			# Use all variable modifications in first search (0 for No, 1 for Yes).
decoy_prefix = decoy_			# Prefix of the decoy protein entries. Used for parameter optimization only.

deisotope = 1			# Perform deisotoping or not (0=no, 1=yes and assume singleton peaks single charged, 2=yes and assume singleton peaks single or double charged).
deneutralloss = 1			# Perform deneutrallossing or not (0=no, 1=yes).
isotope_error = 0/1/2			# Also search for MS/MS events triggered on specified isotopic peaks.
mass_offsets = 0.0			# Creates multiple precursor tolerance windows with specified mass offsets.
mass_offsets_detailed = 			# Optional detailed mass offset list. Overrides mass_offsets if use_detailed_offsets = 1.
use_detailed_offsets = 0			# Whether to use the regular (0) or detailed (1) mass offset list.
precursor_mass_mode = selected			# One of isolated/selected/corrected.

remove_precursor_peak = 1			#  Remove precursor peaks from tandem mass spectra. 0 = not remove; 1 = remove the peak with precursor charge; 2 = remove the peaks with all charge states (only for DDA mode).
remove_precursor_range = -1.500000,1.500000			# m/z range in removing precursor peaks. Only for DDA mode. Unit: Th.
intensity_transform = 0			# Transform peaks intensities with sqrt root. 0 = not transform; 1 = transform using sqrt root.
activation_types = all			# Filter to only search scans of provided activation type(s). Allowed: All, HCD, CID, ETD, ECD.
analyzer_types = all			# Filter to only include scans matching the provided analyzer type(s) in search. Only support the mzML and raw format. Allowed types: all, FTMS, ITMS, ASTMS.
group_variable = 0			# Specify the variable used to decide the PSM group in the group FDR estimation. 0 = no group FDR; 1 = num_enzyme_termini; 2 = PE from protein header.
require_precursor = 0			# If required, PSMs with no precursor peaks will be discarded. For DIA data type only. 0 = no, 1 = yes.
reuse_dia_fragment_peaks = 0			# Allow the same peak matches to multiple peptides. For DIA data type only. 0 = no, 1 = yes.

write_calibrated_mzml = 0			# Write calibrated MS2 scan to a mzML file (0 for No, 1 for Yes).
write_uncalibrated_mzml = 1			# Write uncalibrated MS2 scan to a mzML file (0 for No, 1 for Yes). Only for .raw and .d formats.
write_mzbin_all = 0
mass_diff_to_variable_mod = 0			# Put mass diff as a variable modification. 0 for no; 1 for yes and remove delta mass; 2 for yes and keep delta mass.

localize_delta_mass = 0			# Include fragment ions mass-shifted by unknown modifications (recommended for open and mass offset searches) (0 for OFF, 1 for ON).
delta_mass_exclude_ranges = (-1.5,3.5)			# Exclude mass range for shifted ions searching.
fragment_ion_series = b,y			# Ion series used in search, specify any of a,b,c,x,y,z,Y,b-18,y-18 (comma separated).
ion_series_definitions = 			# User defined ion series. Example: "b* N -17.026548;b0 N -18.010565".

labile_search_mode = off			# type of search (nglycan, labile, or off). Off means non-labile/typical search.
restrict_deltamass_to = all			# Specify amino acids on which delta masses (mass offsets or search modifications) can occur. Allowed values are single letter codes (e.g. ACD) and '-', must be capitalized. Use 'all' to allow any amino acid.
diagnostic_intensity_filter = 0			# [nglycan/labile search_mode only]. Minimum relative intensity for SUM of all detected oxonium ions to achieve for spectrum to contain diagnostic fragment evidence. Calculated relative to spectrum base peak. 0 <= value.
Y_type_masses = 			#  [nglycan/labile search_mode only]. Specify fragments of labile mods that are commonly retained on intact peptides (e.g. Y ions for glycans). Only used if 'Y' is included in fragment_ion_series.
diagnostic_fragments = 			# [nglycan/labile search_mode only]. Specify diagnostic fragments of labile mods that appear in the low m/z region. Only used if diagnostic_intensity_filter > 0.
remainder_fragment_masses = 			# [labile search_mode only] List of possible remainder fragment ions to consider. Remainder masses are partial modification masses left on b/y ions after fragmentation.
labile_fragment_ion_series = b,y			# Labile ion series used in labile search, specify any of a,b,c,x,y,z (comma separated). Replaces intact mods with labile ion(s) for each specified ion type.

search_enzyme_name_1 = Trypsin/P			# Name of the first enzyme.
search_enzyme_cut_1 = KR			# First enzyme's cutting amino acid.
search_enzyme_nocut_1 = 			# First enzyme's protecting amino acid.
search_enzyme_sense_1 = C			# First enzyme's cutting terminal.
allowed_missed_cleavage_1 = 10			# First enzyme's allowed number of missed cleavages per peptide. Maximum value is 5.

search_enzyme_name_2 = null			# Name of the second enzyme.
search_enzyme_cut_2 = 			# Second enzyme's cutting amino acid.
search_enzyme_nocut_2 = 			# Second enzyme's protecting amino acid.
search_enzyme_sense_2 = C			# Second enzyme's cutting terminal.
allowed_missed_cleavage_2 = 2			# Second enzyme's allowed number of missed cleavages per peptide. Maximum value is 5.

num_enzyme_termini = 2			# 0 for non-enzymatic, 1 for semi-enzymatic, and 2 for fully-enzymatic.

clip_nTerm_M = 1			# Specifies the trimming of a protein N-terminal methionine as a variable modification (0 or 1).

# maximum of 16 mods - amino acid codes, * for any amino acid, [ and ] specifies protein termini, n and c specifies peptide termini
variable_mod_01 = 15.9949 M 3
variable_mod_02 = 42.0106 [^ 1
# variable_mod_03 = 79.96633 STY 3
# variable_mod_04 = -17.0265 nQnC 1
# variable_mod_05 = -18.0106 nE 1
# variable_mod_06 = 4.025107 K 2
# variable_mod_07 = 6.020129 KR 2
# variable_mod_08 = 8.014199 K 2
# variable_mod_09 = 10.008269 R 2
# variable_mod_10 = 0.0 site_10 1
# variable_mod_11 = 0.0 site_11 1
# variable_mod_12 = 0.0 site_12 1
# variable_mod_13 = 0.0 site_13 1
# variable_mod_14 = 0.0 site_14 1
# variable_mod_15 = 0.0 site_15 1
# variable_mod_16 = 0.0 site_16 1

allow_multiple_variable_mods_on_residue = 0
max_variable_mods_per_peptide = 3			# Maximum total number of variable modifications per peptide.
max_variable_mods_combinations = 5			# Maximum number of modified forms allowed for each peptide (up to 65534).

output_format = pepXML			# File format of output files (tsv, pin, pepxml, tsv_pin, tsv_pepxml, pepxml_pin, or tsv_pepxml_pin).
output_report_topN = 1			# Reports top N PSMs per input spectrum.
output_max_expect = 50			# Suppresses reporting of PSM if top hit has expectation value greater than this threshold.
report_alternative_proteins = 0			# Report alternative proteins for peptides that are found in multiple proteins (0 for no, 1 for yes).

precursor_charge = 1 4			# Assumed range of potential precursor charge states. Only relevant when override_charge is set to 1.
override_charge = 0			# Ignores precursor charge and uses charge state specified in precursor_charge range (0 or 1).

digest_min_length = 6			# Minimum length of peptides to be generated during in-silico digestion.
digest_max_length = 100			# Maximum length of peptides to be generated during in-silico digestion.
digest_mass_range = 300.0 10000.0			# Mass range of peptides to be generated during in-silico digestion in Daltons.
max_fragment_charge = 10			# Maximum charge state for theoretical fragments to match (1-4).

track_zero_topN = 0			# Track top N unmodified peptide results separately from main results internally for boosting features.
zero_bin_accept_expect = 0			# Ranks a zero-bin hit above all non-zero-bin hit if it has expectation less than this value.
zero_bin_mult_expect = 1			# Multiplies expect value of PSMs in the zero-bin during  results ordering (set to less than 1 for boosting).

check_spectral_files = 0			# Checking spectral files before searching.
minimum_peaks = 15			# Minimum number of peaks in experimental spectrum for matching.
use_topN_peaks = 150			# Pre-process experimental spectrum to only use top N peaks.
min_fragments_modelling = 2			# Minimum number of matched peaks in PSM for inclusion in statistical modeling.
min_matched_fragments = 4			# Minimum number of matched peaks for PSM to be reported.
min_sequence_matches = 2			# [nglycan/labile search_mode only] Minimum number of sequence-specific (not Y) ions to record a match.
minimum_ratio = 0.01			# Filters out all peaks in experimental spectrum less intense than this multiple of the base peak intensity.
clear_mz_range = 0.0 0.0			# Removes peaks in this m/z range prior to matching.

add_Cterm_peptide = 0.0
add_Nterm_peptide = 0.0
add_Cterm_protein = 0.0
add_Nterm_protein = 0.0

add_G_glycine = 0.0
add_A_alanine = 0.0
add_S_serine = 0.0
add_P_proline = 0.0
add_V_valine = 0.0
add_T_threonine = 0.0
add_C_cysteine = 57.02146
add_L_leucine = 0.0
add_I_isoleucine = 0.0
add_N_asparagine = 0.0
add_D_aspartic_acid = 0.0
add_Q_glutamine = 0.0
add_K_lysine = 0.0
add_E_glutamic_acid = 0.0
add_M_methionine = 0.0
add_H_histidine = 0.0
add_F_phenylalanine = 0.0
add_R_arginine = 0.0
add_Y_tyrosine = 0.0
add_W_tryptophan = 0.0
add_B_user_amino_acid = 0.0
add_J_user_amino_acid = 0.0
add_O_user_amino_acid = 0.0
add_U_user_amino_acid = 0.0
add_X_user_amino_acid = 0.0
add_Z_user_amino_acid = 0.0
