## Supplementary material for "Community Resource: A Genome-Based Extension of Large-Scale Wheat Proteogenomics": Vincent_wheat-proteogenomics_technical-note_2026-06-16_Suppl-File-S3.pdf

### Peptide Genomic Coordinate Reconstruction and Validation

#### Overview

The objective of this workflow was to determine the precise genomic coordinates of experimentally identified wheat peptides and represent them as browser-compatible genomic features suitable for visualization in Apollo/JBrowse.

Rather than aligning peptide sequences directly against the genome, the workflow leveraged existing IWGSC RefSeq v2.1 genome annotations to reconstruct peptide genomic coordinates from known protein, transcript, exon, and coding sequence (CDS) relationships.

This approach ensures that peptide projections remain biologically consistent with the underlying wheat genome annotation while preserving exon–intron gene structures.

#### 1. Peptide Genomic Coordinate Reconstruction

##### 1.1. Peptide identification

Raw LC–MS/MS data were processed using FragPipe/MSFragger.

For each peptide-spectrum match (PSM), the workflow produced:

- peptide amino-acid sequence;
- parent protein accession;
- peptide start position within the protein;
- peptide end position within the protein.

Example:

| Protein | Peptide | Protein AA Position |
|---|---|---|
| TraesCS1A03G0001000.1 | AQLVGKPEPTIDE | AA 125–138 |

At this stage, peptide locations were known only within proteins.

##### 1.2. Protein-to-transcript mapping

Each protein accession was linked to its corresponding transcript using the IWGSC RefSeq v2.1 GFF3 annotation.

The GFF3 file provides relationships between:

- genes;

- transcripts;
- exons;
- coding sequences (CDS);
- proteins.

Example:

Protein

↓

Transcript

↓

CDS segments

↓

Chromosomal coordinates

This mapping provides the structural information required to locate protein sequences within the genome.

#### 1.3. Reconstruction of CDS structure

For each transcript, all CDS segments were extracted from the GFF3 annotation.

Example:

| CDS | Chromosome Coordinates |
|---|---|
| CDS1 | 1000–1200 |
| CDS2 | 1500–1700 |
| CDS3 | 2000–2300 |

The CDS segments were then ordered according to transcript structure and strand orientation.

For transcripts located on the negative strand, CDS order was reversed to maintain correct biological translation.

#### 1.4. Conversion of amino-acid positions to nucleotide positions

Peptide coordinates within proteins were converted into CDS nucleotide coordinates.

Because one amino acid corresponds to three nucleotides:

AA start position × 3

AA end position × 3

was used to determine the corresponding coding sequence interval.

Example:

Protein position:

AA 125–138

Converted CDS position:

nt 373–414

The peptide could then be located within the reconstructed transcript CDS.

#### 1.5. Projection onto genomic coordinates

The nucleotide interval corresponding to the peptide was projected onto genomic coordinates using the CDS structure.

If a peptide fell entirely within a single CDS segment, a single genomic interval was generated.

Example:

Peptide

↓

Chr1A:1050–1092

If a peptide crossed an exon boundary, multiple genomic blocks were generated.

Example:

Peptide

↓

Exon 1: Chr1A:1180–1200

Exon 2: Chr1A:1500–1534

This preserves the biological exon–intron structure of the gene.

#### 1.6. BED track generation

Projected genomic intervals were exported as BED6 and BED12 files.

BED12 format was used when peptides spanned multiple exons because it supports multiple genomic blocks within a single feature.

These files were subsequently loaded into Apollo/JBrowse for visualization.

#### 2. Projection Validation

Because peptide genomic coordinates were reconstructed computationally, all projections were subjected to independent validation procedures before inclusion in downstream analyses.

Only projections passing all validation steps were retained.

##### 2.1. Translation Reconstruction Check

**Purpose**

To verify that projected genomic coordinates reproduce the original peptide sequence.

**Procedure**

For each projected peptide:

1. genomic DNA sequence was extracted;
2. exon segments were reconstructed;
3. CDS sequence was assembled;
4. CDS sequence was translated into amino acids;
5. translated sequence was compared with the original FragPipe peptide.

Example:

Original peptide:

AQLVGKPEPTIDE

Translated peptide:

AQLVGKPEPTIDE

Result:

PASS

If the translated peptide differed from the original peptide sequence:

FAIL

This was considered the most important validation step because it directly tested whether the projected genomic coordinates encoded the experimentally observed peptide.

##### 2.2. BED Geometry Check

**Purpose**

To verify that exported genomic intervals formed valid BED features.

**Procedure**

The workflow checked:

- valid chromosome coordinates;
- non-negative positions;
- valid block counts;
- valid block sizes;
- valid block starts;
- consistency between BED fields.

Malformed BED entries were rejected.

This validation ensures that exported tracks can be displayed correctly in genome browsers.

#### 2.3. Chromosome and Strand Consistency Check

**Purpose**

To ensure that peptide projections remained associated with the correct transcript model.

**Procedure**

The workflow verified that:

- projected coordinates occurred on the expected chromosome;
- strand orientation matched the parent transcript;
- exon ordering remained consistent with transcript structure.

Any inconsistencies were flagged as failures.

This validation prevents coordinate assignment errors resulting from transcript reconstruction or strand handling.

#### 2.4. Protein Coordinate Consistency Check

**Purpose**

To verify that genomic projections corresponded to the correct peptide location within the parent protein.

**Procedure**

The workflow compared:

- peptide amino-acid coordinates;
- CDS coordinates;
- reconstructed genomic coordinates.

All coordinate systems had to remain internally consistent.

Failures were recorded whenever projected genomic intervals could not be reconciled with the original peptide location within the source protein.

#### 2.5. Interpretation of Validation Results

A projection was considered valid only when all four validation procedures were passed:

1. Translation reconstruction check;
2. BED geometry check;
3. Chromosome and strand consistency check;
4. Protein coordinate consistency check.

This multi-layer validation framework provides confidence that projected peptide genomic coordinates accurately represent the underlying biological sequence and can be reliably visualized and interpreted within genome browsers.

Projections failing one or more validation checks were excluded from downstream analyses and reported separately in Supplementary Tables S2 and S3.
